# Beyond bacterial paradigms: uncovering the functional significance and first biogenesis machinery of archaeal lipoproteins

**DOI:** 10.1101/2024.08.27.609747

**Authors:** Yirui Hong, Kira S. Makarova, Rachel Xu, Friedhelm Pfeiffer, Mechthild Pohlschroder

## Abstract

Lipoproteins are major constituents of prokaryotic cell surfaces. In bacteria, lipoprotein attachment to membrane lipids is catalyzed by prolipoprotein diacylglyceryl transferase (Lgt). However, no Lgt homologs have been identified in archaea, suggesting the unique archaeal membrane lipids require distinct enzymes for lipoprotein lipidation. Here, we performed *in silico* predictions for all major archaeal lineages and revealed a high prevalence of lipoproteins across the domain Archaea. Using comparative genomics, we identified the first set of candidates for archaeal lipoprotein biogenesis components (Ali). Genetic and biochemical characterization confirmed two paralogous genes, *aliA* and *aliB*, are important for lipoprotein lipidation in the archaeon *Haloferax volcanii*. Disruption of AliA- and AliB-mediated lipoprotein lipidation results in severe growth defects, decreased motility, and cell-shape alterations, underscoring the importance of lipoproteins in archaeal cell physiology. AliA and AliB also exhibit different enzymatic activities, including potential substrate selectivity, uncovering a new layer of regulation for prokaryotic lipoprotein lipidation.

## Introduction

Archaea, once thought to be exclusively associated with extreme environments, are ubiquitous^1^ and important for ecological processes, biotechnology, and human health^2–6^. Despite their morphological similarities with bacteria, archaea and bacteria differ significantly, especially in their membrane lipids. Bacterial membrane lipids have fatty acyl chains linked to glycerol-3-phosphate backbones via ester bonds, whereas archaeal membrane lipids have isoprenoid-based alkyl chains linked to glycerol-1-phosphate backbones via ether bonds^7^. However, both archaea and bacteria use membrane lipids to anchor surface proteins, including prokaryotic lipoproteins^8,9^. Lipoproteins are lipid-anchored surface proteins involved in various cellular processes, ranging from virulence and cell architecture maintenance to nutrient transport and chemotaxis^10–13^. All prokaryotic lipoproteins feature an N-terminal signal peptide followed by a conserved four-residue motif known as the lipobox ([L/V/I]^−3^ [A/S/T/V/I]^−2^ [G/A/S]^−1^ [C]^+1^)^14^. Previous studies have uncovered the essential role of the lipobox in bacterial lipoprotein biogenesis. Depending on the sequence of the signal peptide, lipoprotein precursors are first translocated either in an unfolded state via the general secretory (Sec) pathway or in a folded state via the twin-arginine translocation (Tat) pathway^15^. The bacterial prolipoprotein diacylglyceryl transferase (Lgt) then catalyzes the formation of a thioether bond between the conserved cysteine in the lipobox and a membrane lipid moiety, thus anchoring the lipoprotein to the membrane^16^. Subsequently, lipoprotein signal peptidase (Lsp) cleaves the signal peptide just before the conserved cysteine, yielding the mature lipoprotein^17^.

Lipoproteins are predicted to be major components of archaeal cell surfaces. In the model archaeon *Haloferax volcanii*, 121 of 316 secreted proteins were predicted to be lipoproteins, implying their significant roles in archaeal cell biology^9^. Substitution of the conserved lipobox cysteine with a serine prevented the maturation of several *Hfx. volcanii* lipoproteins and led to protein mislocalization in some cases^18,19^. Furthermore, mass spectrometry analyses of a lipobox-containing halocyanin from *Natronomonas pharaonis* revealed three modifications: N-terminal signal peptide cleavage, attachment of a lipid (diphytanylglycerol diether) to the lipobox cysteine, and N-terminal acetylation^20^, similar to the modifications of bacterial lipoproteins. Despite the accumulating evidence for a lipobox-dependent protein anchoring mechanism in archaea, components underlying such a mechanism have remained enigmatic. Notably, no archaeal homologs of bacterial Lgt or Lsp are identified, suggesting the unique membrane lipids of archaea might require distinct enzymes for lipoprotein biogenesis.

In this study, we aimed to investigate the significance of lipoproteins in archaeal cell biology and to identify key components involved in archaeal lipoprotein biogenesis. Through comparative genomic analyses, we demonstrated the high prevalence and abundance of lipoproteins across the domain Archaea and identified the first set of candidate <u>a</u>rchaeal <u>li</u>poprotein biogenesis components (Ali). Among these, two paralogous proteins, coined AliA and AliB, were particularly promising due to their conserved residues and visual similarities with Lgt. These proteins were further characterized in *Hfx. volcanii* because of its high lipoprotein abundance and genetic tractability. Using genetic and biochemical assays, we confirmed that AliA and AliB are important for archaeal lipoprotein lipidation. Furthermore, deletions of *aliA* and *aliB* significantly impacted archaeal growth, motility, and cell shape, with Δ*aliA* displaying more severe phenotypes than Δ*aliB*. Overall, our findings establish the pivotal roles of lipoproteins in archaeal cell biology and resolve the long-standing search for lipoprotein biogenesis machinery in archaea, thereby opening new avenues for the study of prokaryotic lipoproteins.

## Results

### Lipoproteins are widespread across archaea

Previous *in silico* studies have revealed a high abundance of lipoproteins in several euryarchaeal species^9,19^, but the lipoprotein prevalence across archaea remains unknown. To address this gap, we analyzed 524 archaeal genomes from the Archaeal Clusters of Orthologous Genes (arCOGs) database^21,22^, covering all major archaeal lineages. Putative Sec and Tat lipoproteins as well as other secreted proteins were identified using SignalP 6.0^23^, and the percentage of secreted proteins that were lipoproteins was calculated for each genome. The results show that lipoproteins are not just widespread in Euryarchaeota but also in the DPANN superphylum. Conversely, they are relatively scarce in TACK and Asgard, which are more closely related to eukaryotes where lipobox-containing proteins have not been identified (Fig. 1a, Extended Data Fig. 1 and Supplementary Data 1). In some archaeal species, especially halobacterial species, lipoproteins constitute more than 50% of secreted proteins, implying their significant roles in archaeal physiology. Since both Sec and Tat pathways can be used for archaeal lipoproteins transport^18,19^, we investigated whether different archaeal species show distinct preferences for these pathways. Consistent with previous findings^19^, halobacterial species predominantly use the Tat pathway for lipoprotein translocation (Fig. 1b), likely to avoid protein precipitation in their high-salt cytoplasm. All other archaeal species primarily use the Sec pathway for lipoprotein translocation.

**Fig. 1.**
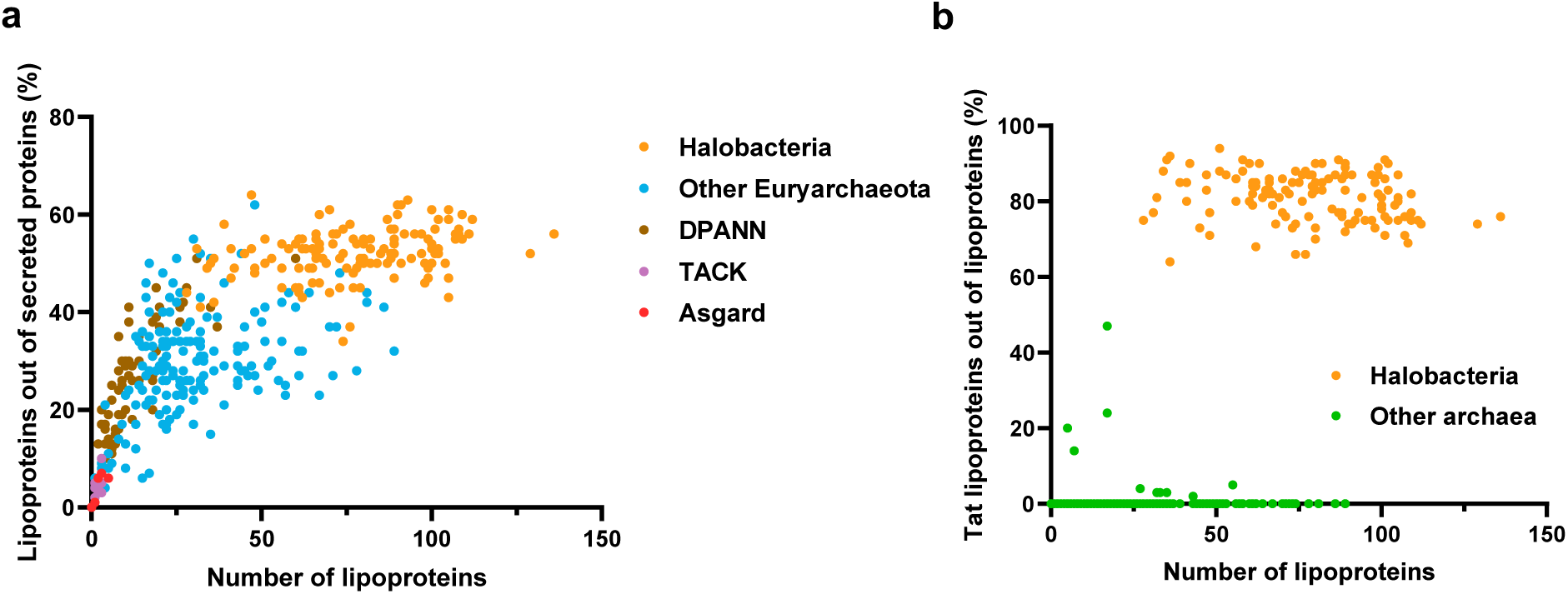
Lipoproteins are widespread across archaea and can be translocated via both Sec and Tat pathways. **a** The percentage of lipoproteins out of total secreted proteins for all archaeal genomes analyzed (n = 524). Each point represents the data for one genome. Euryarchaeota, DPANN, TACK, and Asgard are the four archaeal superphyla. Halobacteria is a class in the Euryarchaeota superphylum. **b** The percentage of Tat lipoproteins out of all lipoproteins for all archaeal genanalyzed (n = 524).

### Computational identification of candidate enzymes for archaeal lipoprotein lipidation

Considering that no archaeal homologs of bacterial Lgt were previously identified, we reasoned that yet uncharacterized archaeal membrane proteins with a phyletic pattern best matching the phyletic pattern of predicted lipoproteins could be promising candidates for further study. Therefore, we calculated a lipoprotein co-occurrence score for each arCOG by comparing its binary phyletic profile with that of lipoproteins (Supplementary Data 2). Among the 25 arCOGs with the highest scores, only two membrane proteins have unassigned functions: arCOG02142, a predicted membrane-associated metalloprotease of the Tiki superfamily^24^, and arCOG02177, an uncharacterized membrane protein (Fig. 2a). arCOG02177 proteins are present in all euryarchaeal lineages with several gaps in Thermoplasmata and Methanobacteria, as well as in the DPANN superphylum with some gaps (Supplementary Data 1 and 3). Multiple alignments of arCOG02177 representatives identified several conserved residues, suggesting potential enzymatic activities of these proteins (Supplementary Data 4). While HHpred search^25^ showed only weak similarity (probability 29%) between arCOG02177 and Lgt (PF01790), an AlphaFold2 model of the *Hfx*. *volcanii* arCOG02177 representative, HVO_2859, showed substantial visual similarity with the *Escherichia coli* Lgt (Fig. 2b). Lgt has two periplasmic subdomains, “arm-1” and “head”^26^, and similar structures are visible in HVO_2859. Moreover, both proteins form pockets with conserved residues, including two conserved arginines and a conserved tyrosine in HVO_2859, which are also found in Lgt and are known to be catalytically important (Fig. 2b)^26,27^. Despite their visual structural and limited sequence similarities, DALI search^28^ did not reveal any structural similarity between these two, possibly due to the different connectivity of their transmembrane segments (Extended Data Fig. 2). A distant paralog of arCOG02177 is encoded in all halobacterial genomes but is classified into a separate arCOG, arCOG02178, due to divergence (Fig. 2a,c and Extended Data Fig. 2,3). Interestingly, both conserved arginine residues in arCOG02177 are replaced by histidines in arCOG02178, suggesting the two paralogs might have different enzymatic activities (Fig. 2b,c and Supplementary Data 4 and 5).

**Fig. 2.**
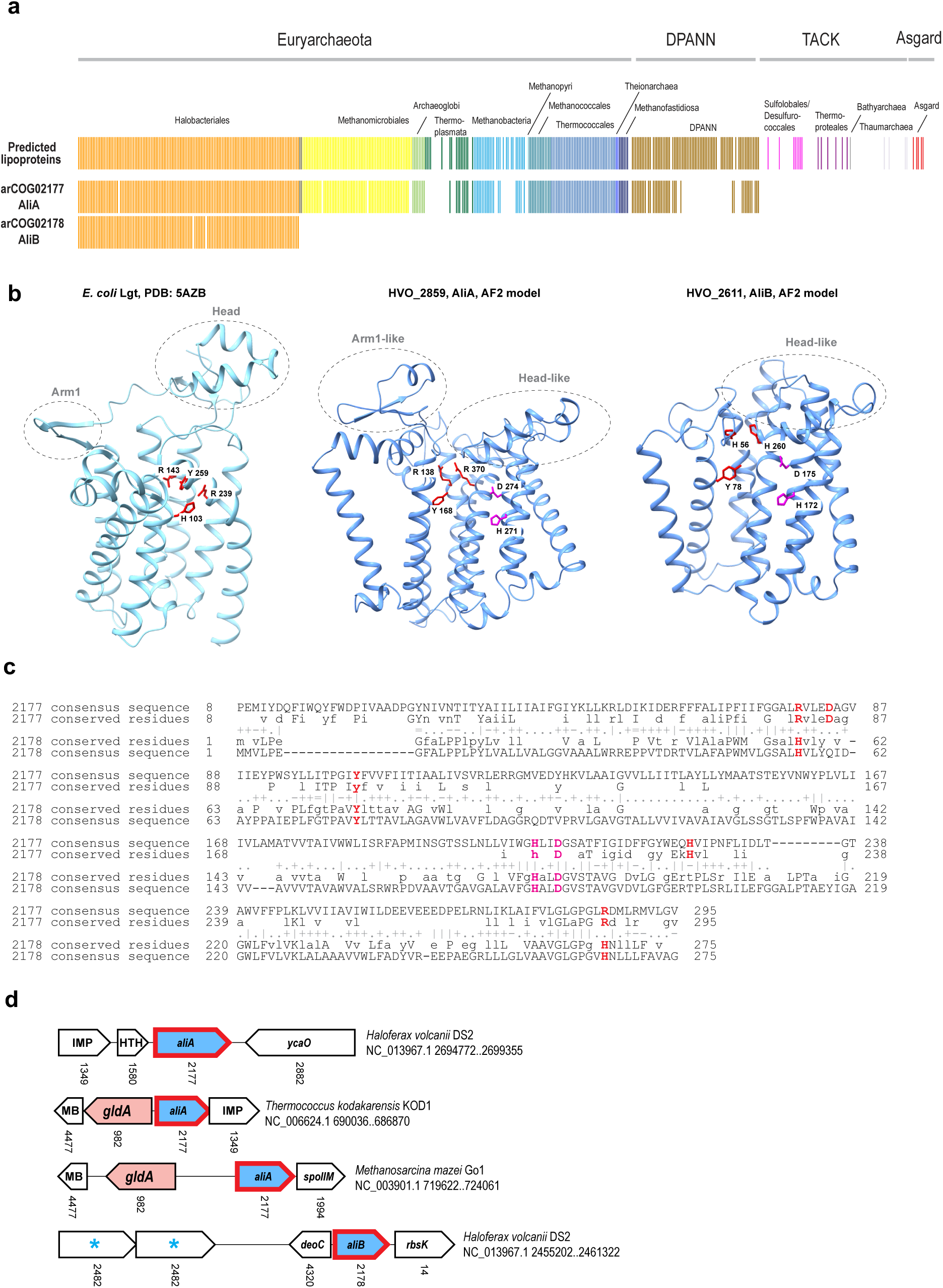
Bioinformatic analyses identified arCOG02177 and arCOG02178 proteins as potential lipoprotein biogenesis components. **a** Phyletic patterns of predicted lipoproteins compared with phyletic patterns of arCOG02177 and arCOG02178. For each genome, gene presence is shown as a vertical bar color-coded according to the major archaeal lineages indicated above. **b** Comparison of *E. coli* Lgt structure (PDB: 5AZB) and AlphaFold2 models of HVO_2859 and HVO_2611. Conserved residues in two different pockets are highlighted in red for one and magenta for the other. Dashed circles indicate similar subdomains between the Lgt structure and AlphaFold2 models. **c** HMM-HMM alignment of arCOG02177 (query) and arCOG02178 (target) showing the conserved residues. Complete alignments are shown in Datasets S4 and S5. The lower case corresponds to less conserved positions. The symbols between the query and the target reflect amino acid similarity between the two multiple alignments as follows: “|”, mostly identical amino acids in respective alignment columns; “+”, very similar amino acids; “.”, positively scored amino acids; “-”, negatively scored amino acids; “=”, very different amino acids. **d** Conserved gene neighborhoods for arCOG02177 and arCOG02178. For each gene neighborhood, the species name, genome partition, and coordinates of the locus are indicated. Genes are depicted by block arrows, with the length roughly proportional to the gene size. arCOG02177 (*aliA*) and arCOG02178 (*aliB*) are shown in blue with a red outline. Lipoproteins are indicated by an asterisk. The genes in the neighborhoods are designated by respective arCOG numbers shown below the arrows, and gene or protein family names are indicated within the arrows. GldA, glycerol dehydrogenase; IMP, inositol monophosphatase family; MB, uncharacterized metal binding protein; DeoC, deoxyribose-phosphate aldolase; SpoIIM, uncharacterized membrane protein, a component of a putative membrane remodeling system; YcaO, ribosomal protein S12 methylthiotransferase accessory factor; RbsK, sugar kinase; HTH, helix-turn-helix protein.

Phylogenetic analyses of arCOG02177 and arCOG02178 representatives showed that major clades are compatible with archaeal taxonomy with limited horizontal transfer (Extended Data, Fig. 3). Such evolutionary behavior is characteristic of proteins involved in important cellular functions^29^. Additionally, gene neighborhood analyses show that arCOG02177 genes are often found nearby, and have the potential to be co-expressed with, inositol monophosphatase family enzymes, predicted metal binding proteins, and glycerol dehydrogenase GldA, a key enzyme of archaeal lipid biosynthesis (Fig. 2d and Supplementary Data 6). arCOG02177 is also associated with arCOG01994, an uncharacterized membrane protein SpoIIM, which was predicted to be involved in membrane remodeling or vesicle formation^30^. arCOG02178 genes are linked to deoxyribose-phosphate aldolase DeoC and are often encoded in the vicinity of predicted lipoproteins (Fig. 2d and Supplementary Data 6). Taken together, these findings suggest that arCOG02177 and arCOG02178 proteins are promising candidates for archaeal lipoprotein biogenesis components.

### AliA and AliB are important for the biogenesis of both Sec and Tat lipoproteins in *Hfx*. *volcanii*

To verify the roles of arCOG02177 and arCOG02178 in lipoprotein biogenesis, we examined their functions *in vivo* using the model archaeon *Hfx. volcanii*. Three predicted lipoproteins including one translocated via the Sec pathway (HVO_1176) and two translocated via the Tat pathway (HVO_B0139 and HVO_1705) were used as substrates (Fig. 3a and Extended Data, Fig. 4). To confirm their identity as lipoproteins, we mutated their lipobox cysteine to a serine (Extended Data, Fig. 4), and tagged both the mutant and wild-type proteins with C-terminal myc tags before overexpressing them in *Hfx. volcanii*. Western blots of cell lysates showed a significantly decreased amount of the three proteins with the C-to-S mutation (Fig. 3b). To determine whether the mutant proteins were degraded or simply released into the supernatant, we performed western blots on concentrated supernatant. Wild-type HVO_1176 and HVO_1705 were present in the supernatant (Extended Data, Fig. 5), presumably due to a phenomenon known as lipoprotein shedding^31^. However, no mutant HVO_1176 and HVO_1705 were detected (Extended Data, Fig. 5), suggesting the mutation might render the proteins unstable. For HVO_B0139, both wild-type and mutant proteins were absent from the supernatant (Extended Data, Fig. 5), reflecting the distinct shedding properties among lipoproteins. Additionally, mutant HVO_1176 and HVO_B0139 migrated more slowly than the mature wild-type proteins (Fig. 3b), suggesting the mutation hindered their maturation. In summary, substituting the lipobox cysteine with a serine disrupted the biogenesis of all three lipoproteins, consistent with the essential role of the cysteine in lipoprotein biogenesis.

**Fig. 3.**
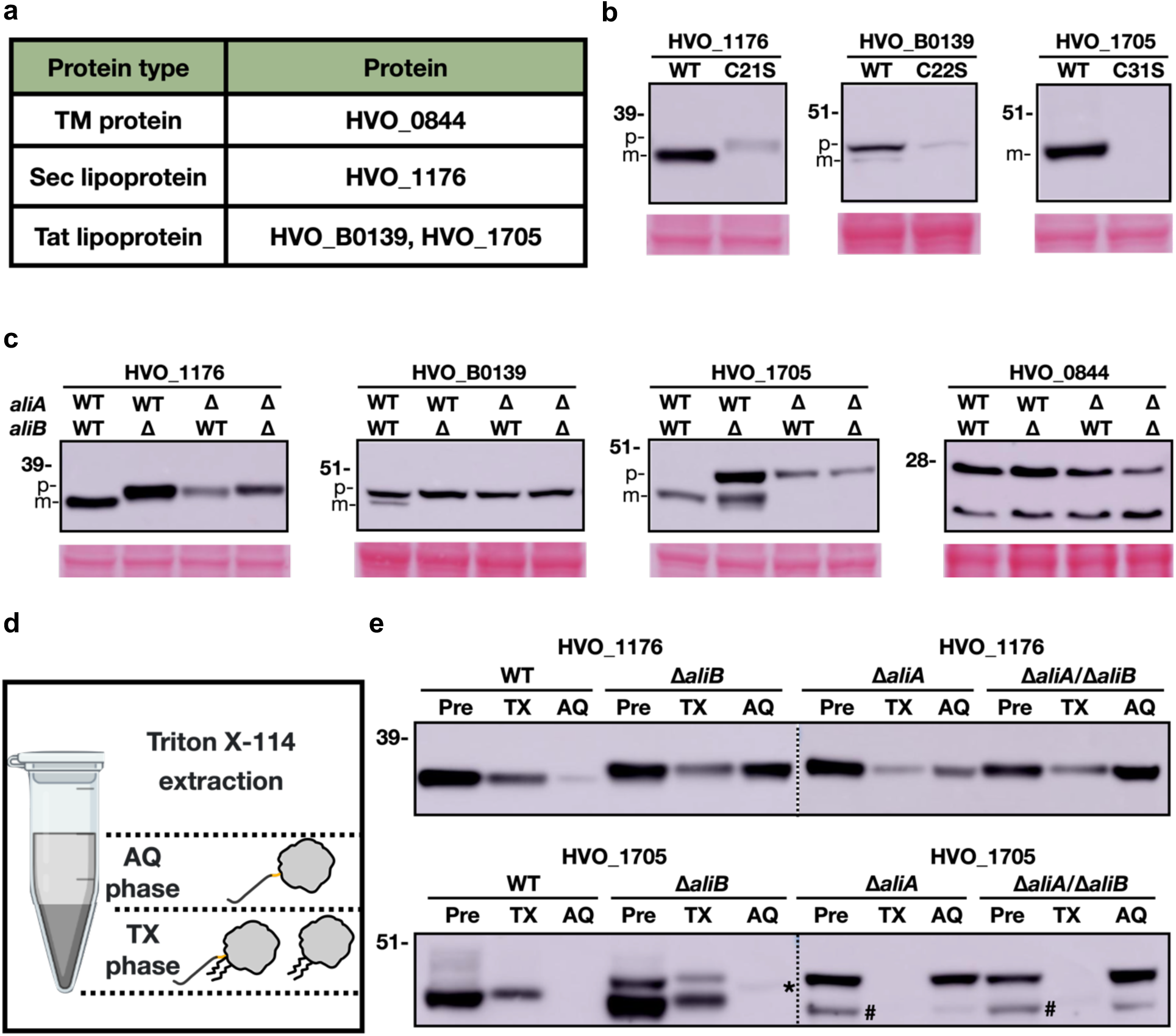
AliA and AliB are involved in Sec and Tat lipoprotein lipidation. **a** Proteins studied in the mobility shift assay. TM, transmembrane. **b** Western blots of overexpressed myc-tagged wild-type lipoproteins and lipoproteins with the cysteine mutated to a serine in *Hfx. volcanii* whole cell lysates. For each lipoprotein, the same amount of total protein was loaded onto the gel across strains. Ponceau S staining of the PVDF membrane is shown at the bottom as a loading control. **c** Western blots of overexpressed myc-tagged lipoproteins in the whole cell lysate of different *Hfx. volcanii* strains. “WT” indicates the presence of wild-type *ali*, and “Δ” indicates the deletion of the wild-type *ali* gene. **d** The anticipated distribution of precursors and mature lipoproteins in the Triton X-114 extraction assay. AQ, aqueous phase; TX, detergent phase. **e** Western blots of samples before and after the Triton X-114 extraction. For each lipoprotein, the same volume of TX and AQ samples were loaded onto the gel. Pre, protein samples before extraction. Dotted lines indicate the border between two different gels. “*” indicates the protein position in the AQ phase of Δ*aliB*. “#”, suspected protein degradation product.

Next, we generated single and double deletion strains of the *Hfx. volcanii* arCOG02177 representative (HVO_2859) and arCOG02178 representative (HVO_2611) to investigate the effect of these deletions on lipoprotein biogenesis. Western blot analyses of myc-tagged HVO_1176 showed that HVO_1176 in the deletion strains migrated more slowly than that in the wild type (Fig. 3c), suggesting a biogenesis defect of HVO_1176 when *hvo_2859* and*/*or *hvo_2611* were deleted. Similarly, although HVO_B0139 appeared as both mature and precursor forms in the wild type, only HVO_B0139 precursors exist in the deletion strains (Fig. 3c). Interestingly, although the maturation of HVO_1705 seemed to be abolished in Δ*hvo_2859*, it was only partially affected in Δ*hvo_2611* (Fig. 3c), implying functional diversification of the two paralogous proteins. The biogenesis defects of these proteins were successfully complemented by *in trans* expression of *hvo*_2859 or *hvo*_2611 (Extended Data, Fig. 6), confirming their roles in lipoprotein biogenesis. As a control, the mobility rate of a transmembrane protein, HVO_0844, was unaffected by the deletions (Fig. 3c), underscoring that the observed mobility shifts in the mutant strains are lipoprotein specific. In summary, deletions of the arCOG02177 representative (HVO_2859) and the arCOG02178 representative (HVO_2611) affect the biogenesis of all three lipoproteins investigated, suggesting both proteins are functional in *Hfx. volcanii* and are important for Sec and Tat lipoprotein biogenesis. Accordingly, we named arCOG02177 and arCOG02178 <u>a</u>rchaeal <u>li</u>poprotein biogenesis components A and B (AliA and AliB), respectively.

### AliA and AliB are involved in lipoprotein lipidation

Based on the results above, we hypothesized that AliA *and* AliB are involved in lipoprotein lipidation. Thus, deletion of *aliA* or *aliB* would prevent lipoproteins from both lipidation and signal peptide cleavage. To test this hypothesis, we analyzed the lipidation status of HVO_1176 and HVO_1705 in different strains using the Triton X-114 extraction assay. HVO_B0139 was excluded from this analysis due to its low expression level and limited quantities of mature proteins. In the assay, the mixture of cell lysates and Triton X-114 solution separates into two phases: lipidated hydrophobic proteins are mainly associated with the detergent phase (TX phase), while non-lipidated hydrophilic proteins remain in the aqueous phase (AQ phase; Fig. 3d)^32,33^. The results indicated that in the wild type, mature HVO_1176 was predominantly associated with the TX phase (Fig. 3e), likely due to the lipid modification. A minor fraction of mature HVO_1176 appeared in the AQ phase (Fig. 3e), possibly due to carryover from the TX phase. Consistent with our prediction that HVO_1176 precursors in Δ*ali* strains lack lipid modifications, these proteins were primarily found in the AQ phase (Fig. 3e), confirming that both AliA and AliB are important for HVO_1176 lipidation. A lesser, yet still notable, amount of these precursors was present in the TX phase, possibly because their highly hydrophobic signal peptide (Extended Data, Fig. 4) was not cleaved in *Δali* strains. Similarly, mature HVO_1705 proteins in both wild type and Δ*aliB* were found in the TX phase (Fig. 3e). In Δ*aliA* strains, HVO_1705 precursors were detected in the AQ phase, suggesting the precursors were non-lipidated and non-cleaved. Interestingly, two forms of HVO_1705 precursors were observed in Δ*aliB*. One form, which was lipidated but non-cleaved, remained in the TX phase, while the other, migrating faster, likely existed in a non-lipidated and non-cleaved state in the AQ phase (Fig. 3e). Collectively, these results revealed that deletion of *aliA* or *aliB* affects the lipidation of HVO_1176 and HVO_1705, confirming the important roles of AliA and AliB in lipoprotein lipidation. While deletions of AliA and AliB have similar effects on HVO_1176, they exhibit different effects on HVO_1705, suggesting that the two paralogs have distinct enzymatic activities.

### AliA*-* and AliB-mediated lipoprotein biogenesis affect archaeal cell physiology

Sporadic studies have investigated the function of individual archaeal lipoproteins^12,13^, but systematic analyses of lipoprotein functions are still lacking. Here, we performed an *in silico* functional categorization of the 93 predicted lipoproteins in *Hfx. volcanii*, revealing their involvement in essential cellular processes including nutrient transport and metabolism, energy production and conversion, and signal transduction (Fig. 4a and Supplementary Data 7). Given their broad roles, we anticipated that disruption of lipoprotein biogenesis in Δ*ali* strains would affect *Hfx. volcanii* fitness. Consistent with the high number of lipoproteins involved in nutrient uptake (Fig. 4a), Δ*ali* strains had smaller, lighter colonies on semi-defined Hv-Cab agar plates (Fig. 4b) and slower growth rates in the exponential phase in Hv-Cab liquid medium (Fig. 4c) compared to the wild type. The growth defect of Δ*ali* strains were more pronounced in minimal medium with glucose and alanine as the sole carbon and nitrogen sources, respectively (Fig. 4d). Δ*aliB* showed a strong reduction in growth compared to the wild type, while Δ*aliA* and Δ*aliA*/Δ*aliB* exhibited minimal growth, highlighting again the functional distinction between AliA and AliB.

**Fig. 4.**
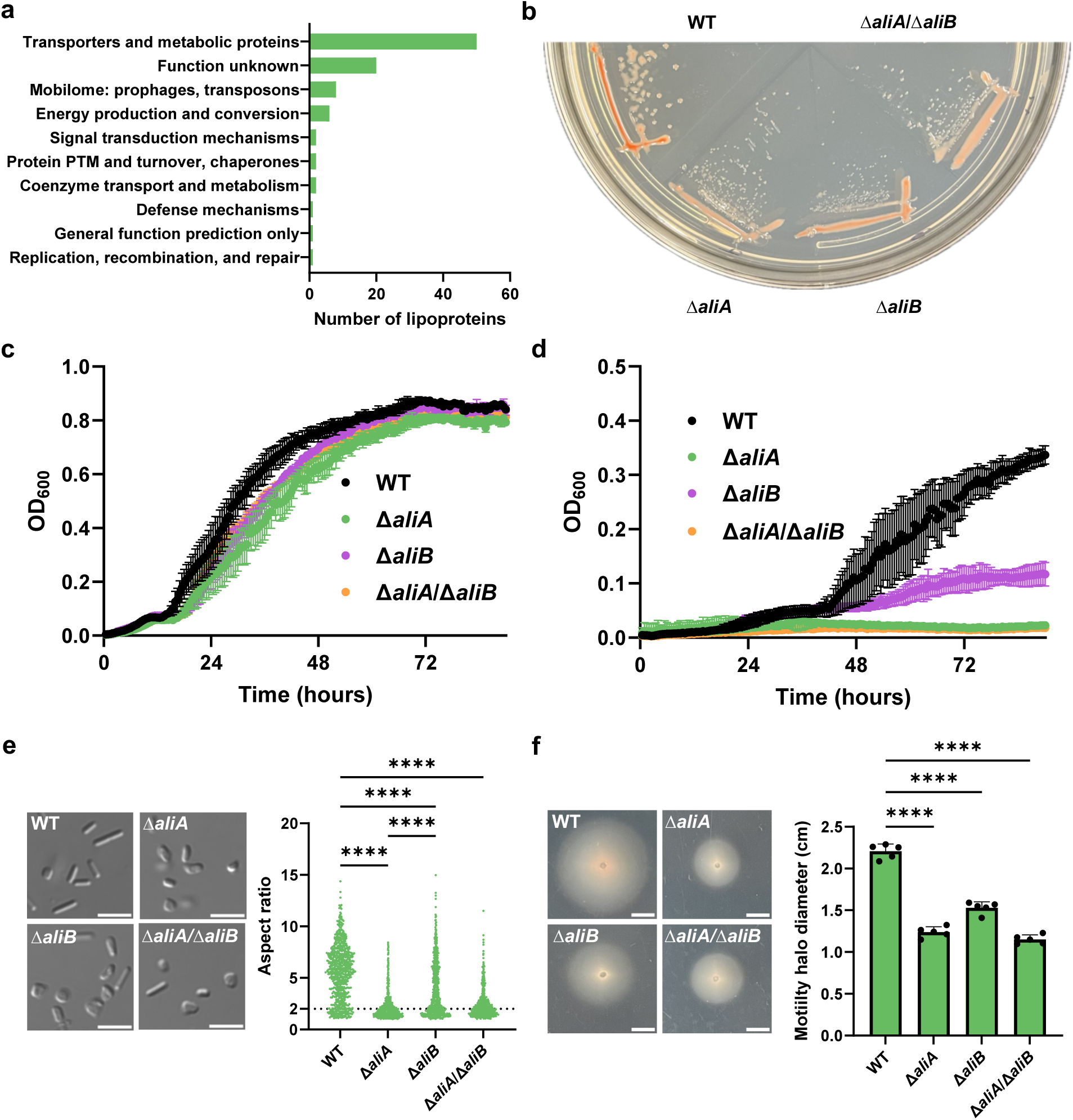
AliA and AliB are important for the growth, cell shape, and motility of *Hfx. volcanii.* **a** Functional classification of 93 predicted *Hfx. volcanii* lipoproteins. PTM, posttranslational modification. **b** Colony morphology of *Hfx. volcanii* wild type and Δ*ali* mutants. Strains were streaked out on Hv-Cab agar plates and incubated at 45 °C for four days prior to imaging. **c** Growth curves of *Hfx. volcanii* wild type and Δ*ali* mutants in Hv-Cab medium. The graph is representative of four biological replicates. **d** Growth curves of *Hfx. volcanii* wild type and Δ*ali* mutants in minimal medium with glucose and alanine as the sole carbon and nitrogen sources, respectively. The graph is representative of four biological replicates. **e** Cell morphology of *Hfx. volcanii* wild type and Δ*ali* mutants (left) in early log and the quantification (right). Cells with aspect ratios < 2 are considered disks and short rods, while others are considered regular rods. Three biological replicates were analyzed for each strain, and 257 cells were analyzed for each biological replicate. Scale bars indicate 5 µm. Data were analyzed using a one-way ANOVA test. *\*\*\*\*P* < 0.0001. **f** Motility halos of *Hfx. volcanii* wild type and Δ*ali* mutants (left) and the quantification (right). Scale bars indicate 0.5 cm. The quantification shows the motility halo diameters and their mean of five biological replicates for each strain. Data were analyzed using a one-way ANOVA test. Error bars represent standard deviations*. ****P* < 0.0001.

Previous studies have shown that wild-type *Hfx. volcanii* transitions from rod-shaped cells in early log to disk-shaped cells in late log when grown in the Hv-Cab medium^34^. While rod-shaped cells are associated with better swimming ability^35^, disk-shaped cells are hypothesized to have better nutrient uptake due to their increased surface-area-to-volume ratio^36^. Given that Δ*ali* strains are likely deficient in nutrient uptake, we examined whether these cells would change their shapes to compensate for this defect. Microscopic analyses showed that early-log wild-type *Hfx. volcanii* culture contained mostly rods with some disks (Fig. 4e). Yet, Δ*ali* cells were primarily disk-shaped in early log (Fig. 4e), suggesting they transitioned to the disk shape earlier than the wild type, presumably to enhance their nutrient uptake abilities. Given the connection between cell shape and motility, we further analyzed the motility of Δ*ali* mutants. All three Δ*ali* mutants showed smaller motility halos compared to the wild type (Fig. 4f), consistent with their early transition to disks. Moreover, deletion of *aliA* resulted in more severe phenotypes, with a smaller motility halo and more disk-shaped cells compared to Δ*aliB* (Fig. 4e,f). In summary, these results represent the first systematic investigation into lipoprotein functions, revealing the crucial role of AliA*-* and AliB-mediated lipoprotein biogenesis in archaeal cell physiology.

## Discussion

Archaeal lipoproteins, despite their high abundance and important roles, have been largely understudied, with no components of their biogenesis pathway identified until now. In this study, using comparative genomics, reverse genetics, and biochemical approaches, we identified the first two components in the pathway, AliA and AliB, which were shown to be crucial for archaeal lipoprotein lipidation.

Our bioinformatic analyses suggest that a family of membrane proteins, arCOG02177, referred to as AliA, exhibits a phyletic pattern closely matching that of predicted lipoproteins across archaea. It has a marginal sequence similarity with bacterial Lgt. Visual examination of the *E. coli* Lgt crystal structure and AliA AlphaFold2 model revealed several common structural features and a remarkable similarity of their conserved residues, including two arginines and an aromatic residue, which are known to be important for Lgt activity^26,27^. Conversely, a DALI search found no structural similarity between the AliA model and Lgt, suggesting that AliA either originated from Lgt by a radical protein rearrangement or has an independent origin. A diverged AliA paralog, arCOG02178 or AliB, possesses a distinct set of putative catalytic residues. While the AliA family is present in Euryarcheota and DPANN and is hypothesized to have been present in the last archaeal common ancestor^30,37^, AliB is limited to halobacterial species. Several archaeal genomes have a substantial number of predicted lipoproteins but lack both AliA and AliB, indicating the potential existence of an alternative lipoprotein lipidation pathway yet-to-be-discovered.

To confirm the involvement of AliA and AliB in lipoprotein biogenesis, we carried out *in vivo* experiments in *Hfx. volcanii* with three predicted lipoproteins as substrates. Substitutions of the conserved cysteine with a serine rendered all three lipoproteins unstable, as shown by their significantly decreased abundance in cell lysates and absence from the supernatant. When overexpressed in Δ*ali* strains, lipoproteins HVO_1176 and HVO_B0139 only existed as precursors, indicating that both AliA and AliB are important for their biogenesis. In contrast, HVO_1705 existed as both mature proteins and precursors in *ΔaliB* but only as precursors in *ΔaliA*, implying a functional difference between AliA and AliB as well as a potential substrate selectivity of AliB. Using the Triton X-114 extraction assay, we demonstrated that AliA and AliB play key roles in lipoprotein lipidation. Both are equally critical for HVO_1176 lipidation, as deletion of either led to the accumulation of non-lipidated HVO_1176. AliA is also vital for HVO_1705 lipidation but deletion of *aliB* only showed minimal effects on HVO_1705 lipidation. Notably, although the three lipoproteins showed a significantly decreased abundance with the cysteine mutation, a similar level of decrease was not seen in Δ*ali* strains. This result suggests that in addition to AliA- and AliB-mediated lipidation, the cysteine is also involved in other processes that maintain lipoprotein stability. It is possible that the mutant lipoproteins were degraded by extracellular proteases, given their increased probability as signal peptidase I substrates with the C-to-S mutation (Extended Data, Table 1). Other cysteine modifications, such as acetylation of its amide group^20^, might also contribute to lipoprotein stability. This strategy is not uncommon; for instance, methylation of a lipidated cysteine has been shown to significantly enhance the half-life of certain proteins^38,39^.

Additionally, disruption of lipoprotein biogenesis by deleting *aliA* or *aliB* affected multiple aspects of *Hfx. volcanii* physiology. The mutant strains formed smaller and lighter colonies on Hv-Cab agar plates and exhibited slower exponential growth in Hv-Cab liquid medium compared to the wild type. However, their growth defect was not severe. Previous studies indicate that some non-lipidated lipoproteins in *Hfx. volcanii* can still be anchored to cell membranes via their hydrophobic signal peptides^19^ and might therefore remain functional. Similarly, non-lipidated lipoproteins in Δ*ali* strains might still support cell growth in semi-defined medium by contributing to the transport and metabolism of certain carbon and nitrogen sources. However, in minimal medium using glucose and alanine as the sole carbon and nitrogen sources, respectively, *Δali* strains showed severe growth defects, likely due to the impaired biogenesis of lipoproteins responsible for glucose and alanine transport or metabolism. *Ali* deletions also affect cell shape and motility. Δ*ali* strains transitioned earlier from rods to disks compared to the wild type, potentially as a compensation mechanism for their defective nutrient uptake capability. Moreover, since no known cell-shape regulatory components are predicted as lipoproteins (Supplementary Data 7), uncharacterized regulatory mechanisms involving lipoproteins might exist and warrant further investigation. Consistent with the association between disk shape and reduced motility, Δ*ali* strains showed smaller motility halos compared to the wild type.

Considering the phenotypic differences of Δ*aliA* and Δ*aliB* in lipoprotein biogenesis and cell physiology, AliA and AliB may have different enzymatic activities and substrate specificities. This hypothesis is supported by the fact that two conserved arginine residues in AliA are replaced by histidines in AliB. In *E. coli* Lgt, the two conserved arginines (R143 and R239) are essential for the lipidation reaction and are proposed to facilitate specific binding of Lgt to the negatively charged membrane lipids^26^. Therefore, AliA and AliB may differ in lipid-binding capabilities, resulting in varying efficiencies of lipoprotein lipidation. Such divergence has been observed in the model bacterium *Mycobacterium smegmatis*^40^, where only one *M. smegmatis* Lgt homolog complemented a *Corynebacterium glutamicum* Δ*lgt* strain^41^. The other homolog carries mutations in several conserved residues and is proposed to either be inactive or have decreased enzymatic activity^40,41^. Despite this finding, closer examinations of the functional differentiation of the two *M. smegmatis* Lgt paralogs are still lacking. Therefore, future structure-function analyses of AliA and AliB using site-directed mutagenesis will provide valuable insights into the lipoprotein biogenesis in both archaea and bacteria. Similarly, *in vitro* lipidation assays, which are currently limited by low protein expression levels, will help determine whether AliA and AliB have stand-alone lipidation activities and different lipid-binding capabilities.

Overall, our study identified and characterized the first two components of the archaeal lipoprotein biogenesis pathway, which are conserved across archaeal species and crucial for archaeal cell biology. The successful identification of AliA and AliB using comparative genomics and sequence analyses also paves the way for identifying and characterizing other components in this pathway, including the downstream lipoprotein signal peptidase, which is currently under investigation. Furthermore, we highlighted the distinct roles of AliA and AliB in lipoprotein lipidation, raising exciting new questions about the delicate regulation of lipoprotein lipidation in both archaea and bacteria.

## Materials and Methods

### Plasmids, strains, primers, and reagents

All plasmids, strains, and primers used in this study are listed in Supplementary Tables 1-3. DNA Phusion Taq polymerase, restriction enzymes, and DNA ligase were purchased from New England BioLabs. The RQ1 RNase-Free DNase was purchased from Promega.

### Sequence comparison, phylogenetic analysis, and gene neighborhood analysis

Secreted proteins encoded in 524 archaeal genomes from the arCOG database were identified using SignalP 6.0^23^. SignalP 6.0 identifies five types of signal peptides, including distinct predictions for lipoproteins secreted via Tat (TATLIPO) and Sec (LIPO) pathways. We considered only confident predictions (probability ≥90%) for all types of signal peptides. The arCOG database^21,22^ that includes annotated clusters of orthologous genes for 524 archaeal genomes covering all major archaeal lineages is available at https://ftp.ncbi.nih.gov/pub/wolf/COGs/arCOG/tmp.ar18/.

HHpred online tool^25^ was used to search for sequence similarity with default parameters against HMM profiles derived from PDB, PFAM, and CDD databases. Muscle5 program^42^ with default parameters was used to construct multiple sequence alignments. For both arCOG02177 and arCOG02178 alignments, a consensus sequence was calculated as described previously^43^. Briefly, each amino acid in the consensus sequence corresponds to the best-scoring amino acid against all amino acids in the respective alignment column, calculated using the BLOSUM62 matrix. The HHalign software^44^ automatically calculates conserved residues. For phylogenetic analysis, poorly aligned sequences or fragments were discarded. Columns in the multiple alignment were filtered for homogeneity value^43^ 0.05 or higher and gap fraction less than 0.667. This filtered alignment was used as an input for the FastTree program^45^ to construct an approximate maximum likelihood phylogenetic tree with the WAG evolutionary model and gamma-distributed site rates. The same program was used to calculate support values. DeepTMHMM^46^ was used to search for transmembrane segments. For genome context analysis and search for putative operons, neighborhoods containing five upstream and five downstream genes were constructed for all identified arCOG02177 and arCOG02178 genes. Structural models for HVO_2859 and HVO_2611 proteins were predicted using ColabFold v1.5.5 (AlphaFold2)^47^ and visualized using UCSF ChimeraX^48^. DALI server^28^ was used to compare predicted models with PDB database.

### Growth conditions

*Hfx. volcanii* strains were grown at 45 °C in liquid (orbital shaker at 250 rpm with an orbital diameter of 2.54 cm) or on solid agar (containing 1.5% [w/v] agar) semi-defined Hv-Cab medium^34^, supplemented with tryptophan (Fisher Scientific) and uracil (Sigma) at a final concentration of 50 μg ml^-1^ unless otherwise noted. Uracil was left out for strains carrying pTA963 or pTA963-based plasmids. 200 μg ml^-1^ tryptophan was used to induce pTA963 expression for western blots, while 50 μg ml^-1^ tryptophan was used in all other experiments. For growth curves in minimal medium, CDM minimal medium was made as previously described^49^, using the same 18% salt water, trace elements, thiamine, and biotin solution as the Hv-Cab medium. Additionally, 20 mM glucose and 25 mM alanine were added as the sole carbon and nitrogen sources, respectively. *E. coli* strains used for cloning were grown at 37 °C in liquid (orbital shaker at 250 rpm with an orbital diameter of 2.54 cm) or on solid agar (containing 1.5% [w/v] agar) NZCYM (RPI) medium, supplemented, if required, with ampicillin at a final concentration of 100 μg ml^-1^.

### Construction of *Hfx. volcanii* overexpression strains

Genes of interest were first amplified by PCR using specific primers that contain artificial restriction sites and then digested using BamHI and EcoRI. The PCR products were then ligated to the BamHI and EcoRI digested expression vector pTA963. The ligation products were later transformed into *E. coli* DH5α cells, and the plasmids were extracted from the DH5α cells using the PureLink Quick Plasmid Miniprep Kit (Invitrogen). Site-directed mutagenesis of *hvo_B0139*, *hvo_1176*, and *hvo_1705* was performed on the DH5α overexpression plasmids using the New England BioLabs Q5 site-directed mutagenesis kit with gene-specific primers. The DH5α plasmids were then transformed into *E. coli* Dam^–^ strain DL739 to get demethylated plasmids. Plasmid sequences were confirmed by Sanger sequencing or whole plasmid sequencing provided by Eurofins Genomics. The verified demethylated plasmids were transformed into *Hfx. volcanii* using the polyethylene glycol (PEG) method^50^.

To construct vectors overexpressing both *aliB* and a lipoprotein gene, the pYH10 (pTA963-based vector expressing the *aliB*-His construct) was linearized with HindIII. The linearized vector was then treated with Klenow DNA Polymerase to remove the overhangs created by the HindIII digestion. In parallel, pYH3 (pTA963-based vector expressing the C-terminally myc-tagged HVO_1176) was digested with PvuII to isolate the *hvo_1176*-myc fragment with the tryptophan-inducible promoter (p.*tnaA*-*hvo_1176-*myc). The PvuII-digested fragment was ligated to the previously linearized blunt-end pYH10, generating the pYH12 vector containing both *aliB-*His and *hvo_1176-*myc under separate tryptophan-inducible promoters. The same cloning strategy was used to generate pYH13 (pTA963 expressing both *aliB-*His and *hvo_1705-*myc). Demethylated plasmid preparation and *Hfx. volcanii* transformations were performed as described above.

Overexpression vectors carrying both *aliA* and a lipoprotein gene could not be isolated from the *E. coli* DH5α strain, possibly due to the potential toxicity of the plasmids to *E. coli* cells. Therefore, we generated the corresponding Gibson assembly^51^ products and directly transformed the products into *Hfx. volcanii*. For example, the p.*tnaA*-*hvo_1176-*myc fragment was amplified by PCR from the pYH3 vector. To facilitate homologous recombination between this fragment and HindIII-digested pYH11 (pTA963-based vector expressing *aliA*), a 22-nucleotide sequence homologous to one blunt end of the linearized pYH11 and a 20-nucleotide sequence homologous to the other blunt end were added to opposite sides of the *p.tnaA*-*hvo_1176*-myc fragment during PCR. The fragment and linearized pYH11 were then added to a homemade Gibson assembly mixture to generate the desired co-expression construct.

### Generation of chromosomal deletions in *Hfx. Volcanii*

Chromosomal deletions were generated by the pop-in/pop-out method as previously described^52^ using H53 as the parent strain and the Hv-Cab medium. Successful gene deletions were first confirmed by colony PCR. The deletion strains were further analyzed by Illumina whole genome sequencing performed by SeqCenter (Pittsburgh, PA, USA) to confirm the complete deletion of the gene as well as search for any secondary genome alterations.

### Immunoblotting

A single colony was used to inoculate a 5 ml liquid culture. At an OD_600_ of 0.8, cells were harvested by centrifugation at 4,300 g for 10 min at 4 °C. The supernatant was transferred into a new tube and centrifuged again at 10,000 g for 10 min. The supernatant was then concentrated using the Amicon Ultra-4 3K centrifugal filter unit at 7,500 g and 4 °C until the final volume was below 250 μl. 2 ml PBS buffer (containing 1 mM AEBSF protease inhibitor from Thermo Scientific) was added to the concentrated supernatant for buffer exchange, and the sample was concentrated again to less than 250 μl. The buffer exchange was repeated once, and the supernatant was concentrated to ∼100 μl and transferred to a clean 1.5 ml tube. The cell pellet was resuspended with 1 ml 18% salt water, made by diluting the 30% salt water used in the Hv-Cab medium. The cells were pelleted again at 4,300 g for 10 min at 4 °C to remove any leftover medium. Cells were then resuspended in 0.25 ml PBS buffer with 1 mM AEBSF and lysed by freezing (with liquid nitrogen) and thawing (at 37 °C) four times. 20 μl RQ1 DNase solution was added to cell lysates, and the mixture was incubated at 37 °C for 30 min to degrade DNA. Unlysed cells were removed by centrifugation at 4,300 g for 10 min at 4 °C, and the supernatant was transferred to a new tube. The protein concentration was measured using the Pierce BCA Protein Assay Kit from Thermo Scientific.

The desired amounts of protein were supplemented with 50 mM dithiothreitol and NuPAGE LDS Sample Buffer (1X). Samples were incubated at 70 °C for 10 min before loaded onto NuPAGE 10%, Bis-Tris gels (1.0 mm x 10 wells) with NuPAGE MOPS SDS Running Buffer. After electrophoresis, proteins were transferred to a polyvinylidene difluoride (PVDF) membrane (Millipore) using a semi-dry transfer apparatus (BioRad) at 15 V for 30 min. Subsequently, the membrane was stained with Ponceau S Staining Solution (Cell Signaling Technology) to verify the successful protein transfer. The membrane was then washed for 10 min twice in PBS buffer and blocked for 1 h in 5% non-fat milk (LabScientific) in PBST buffer (PBS with 1% Tween-20). After blocking, the membrane was washed twice in PBST and once in PBS. For detection of the myc tag, anti-Myc antibody (9E10, UPenn Cell Center Service) was diluted 1,000 times with PBS buffer containing 3% bovine serum albumin, which was then used to incubate the membrane overnight at 4 °C. Subsequently, the membrane was washed twice in PBST and once in PBS, followed by a 45-min incubation at room temperature with the secondary antibody solution (Amersham ECL anti-mouse IgG, horseradish peroxidase-linked whole antibody, from sheep (Cytiva) diluted 10,000 times in PBS containing 10% non-fat milk). After incubation, the membrane was washed three times in PBST and once in PBS. HRP activity was assessed using the Amersham ECL Prime Western Blotting Detection Reagent (GE).

### Triton X-114 extraction

Triton X-114 extraction was performed as previously described^33^ with some modification. 100 μl ice-cold 20% Triton X-114 and 0.9 ml *Hfx. volcanii* protein sample containing ∼500 μg proteins were added to a chilled glass container. The container was put into a glass jar filled with ice to maintain the mixture temperature at 0–4 °C. The mixture was stirred on ice at 340 rpm for 2 h before being transferred to a chilled microtube. The mixture was then centrifuged at 4 °C for 10 min at 15,000 g to remove any precipitation. The supernatant was incubated at 37 °C for 10 min and centrifuged at room temperature for 10 min at 10,000 g. After centrifugation, the mixture was separated into the upper aqueous (AQ) phase and lower detergent (TX) phase. 88.8 μl 20% Triton X-114 was added to the AQ phase, while 0.8 ml of ice-cold PBS buffer was added to the TX phase. After a brief vortex, the mixture was incubated on ice for 10 min and then at 37 °C for 10 min, followed by centrifugation at room temperature for 10 min at 10,000 g. The new TX phase of the AQ phase sample was discarded, and 9 ml of ice-cold methanol was added to the AQ phase. For the TX phase sample, the new AQ phase was discarded, and 0.9 ml ice-cold methanol was added to the TX phase. Both AQ and TX phase samples were stored at −80 °C overnight to precipitate the proteins. The next day, the samples were centrifuged at 4 °C for 10 min (15,000 g for microtubes and 12,000 g for 15 ml Falcon tubes). The sediments of the AQ and TX samples containing the desired proteins were resuspended in the same volume of PBS buffer with 1 mM AEBSF and stored at –20 °C. The same volume of AQ and TX samples were used for western blots.

### Hfx. volcanii growth curve

Colonies for each strain were inoculated in 5 ml Hv-Cab liquid medium and grown until they reached mid-log phase (OD_600_ between 0.3 and 0.8). 1 ml cultures were then harvested via centrifugation (15,800 g, 1 min at room temperature) and washed twice with 18% salt water. Cultures were then diluted to an OD_600_ of 0.01 in a flatbottom polystyrene 96-well plate (Corning) with 200 μl Hv-Cab medium or fresh CDM-based minimal medium supplemented with 20 mM glucose and 25 mM alanine. 200 μl of medium was also aliquoted into each well for two rows of perimeter wells of the plate to help prevent culture evaporation. Growth curves were measured using a BioTek Epoch 2 microplate reader with BioTek Gen6 v.1.03.01 (Agilent). Readings were taken every 30 min with double orbital, continuous fast shaking (355 cycles per min) in between. Readings were taken at a wavelength of 600 nm for about 90 h and then plotted using GraphPad Prism version 10.0.1 for Windows 64-bit.

### Motility assays and motility halo quantification

Motility was assessed on 0.35% agar Hv-Cab medium plates with supplements as required. A toothpick was used to stab-inoculate the agar followed by incubation at 45 °C. Motility assay plates were removed from the incubator after three days and imaged after one day of room temperature incubation. Motility halos were quantified using Fiji (ImageJ)^53^. Images were uploaded to Fiji, and the scale was set based on a 100 mm Petri dish diameter. The statistical significance of halo diameters was assessed with a one-way ANOVA test using GraphPad Prism version 10.0.1 for Windows 64-bit.

### Live-cell imaging and analysis

*Hfx. volcanii* strains were inoculated from a single colony into 5 ml of Hv-Cab liquid medium with appropriate supplementation. Cultures were grown until they reached an OD_600_ of 0.07. Three biological replicates of each strain were grown as described above, and for each culture, 1 ml aliquots were centrifuged at 4,900 g for 6 min. Pellets were resuspended in ∼10 μl of medium. 1.5 μl of the resuspension was placed on glass slides (Fisher Scientific), and a glass coverslip (Globe Scientific Inc.) was placed on top. Slides were visualized using a Leica Dmi8 inverted microscope attached to a Leica DFC9000 GT camera with Leica Application Suite X (version 3.6.0.20104) software, and both brightfield and differential interference contrast (DIC) images were captured at 100x magnification. Brightfield images were quantified using CellProfiler^54^. Brightfield images were uploaded to CellProfiler, and parameters were set for each set of images (strain and replicate) based on specific image conditions to eliminate noise and maximize the number of identified cells. The CellProfiler pipeline used to analyze and quantify the images can be found at https://doi.org/10.5281/zenodo.8404691. Cell-specific data for each image set were exported, and aspect ratio was calculated. The statistical significance of aspect ratio comparisons between strains was assessed with a one-way ANOVA test using GraphPad Prism 10.0.1 for Windows 64-bit.

## Acknowledgments

We thank Dr. Yuri Wolf for suggestions on the design of Fig. 1. We are grateful to Dr. Fevzi Daldal at the University of Pennsylvania for providing anti-myc antibodies for initial tests and offering insightful suggestions on the Triton X-114 experiments. We also extend our thanks to the Maupin-Furlow lab at the University of Florida for their suggestions on the growth curve experiment using minimal medium. Additionally, we acknowledge Dr. Farid Halim at the University of Minnesota for his guidance on constructing dual expression vectors and the Bisson lab at Brandeis University for their advice on the Gibson assembly. Lastly, we thank Pohlschroder lab members for reading the manuscript and providing valuable feedback.

Y.H., R.X., and M.P. were supported by the National Science Foundation Grant NSF-MBC2222076 and the University of Pennsylvania Research Fund grant. Y.H. was additionally supported by the Teece Dissertation Research Award and the SASGov Research Grant at the University of Pennsylvania. K.S.M. was supported by intramural funds of the US Department of Health and Human Services (National Institutes of Health, National Library of Medicine).

## Author Contributions

Y.H., R.X., K.S.M., F.P., and M.P. designed research; Y.H., R.X., K.S.M., F.P., and M.P. performed research; Y.H., R.X., K.S.M., F.P., and M.P. analyzed data; Y.H., R.X., K.S.M., F.P., and M.P. wrote the manuscript.

## Competing Interests statement

The authors declare no competing interest.

---

**Extended Data Fig. 1.**
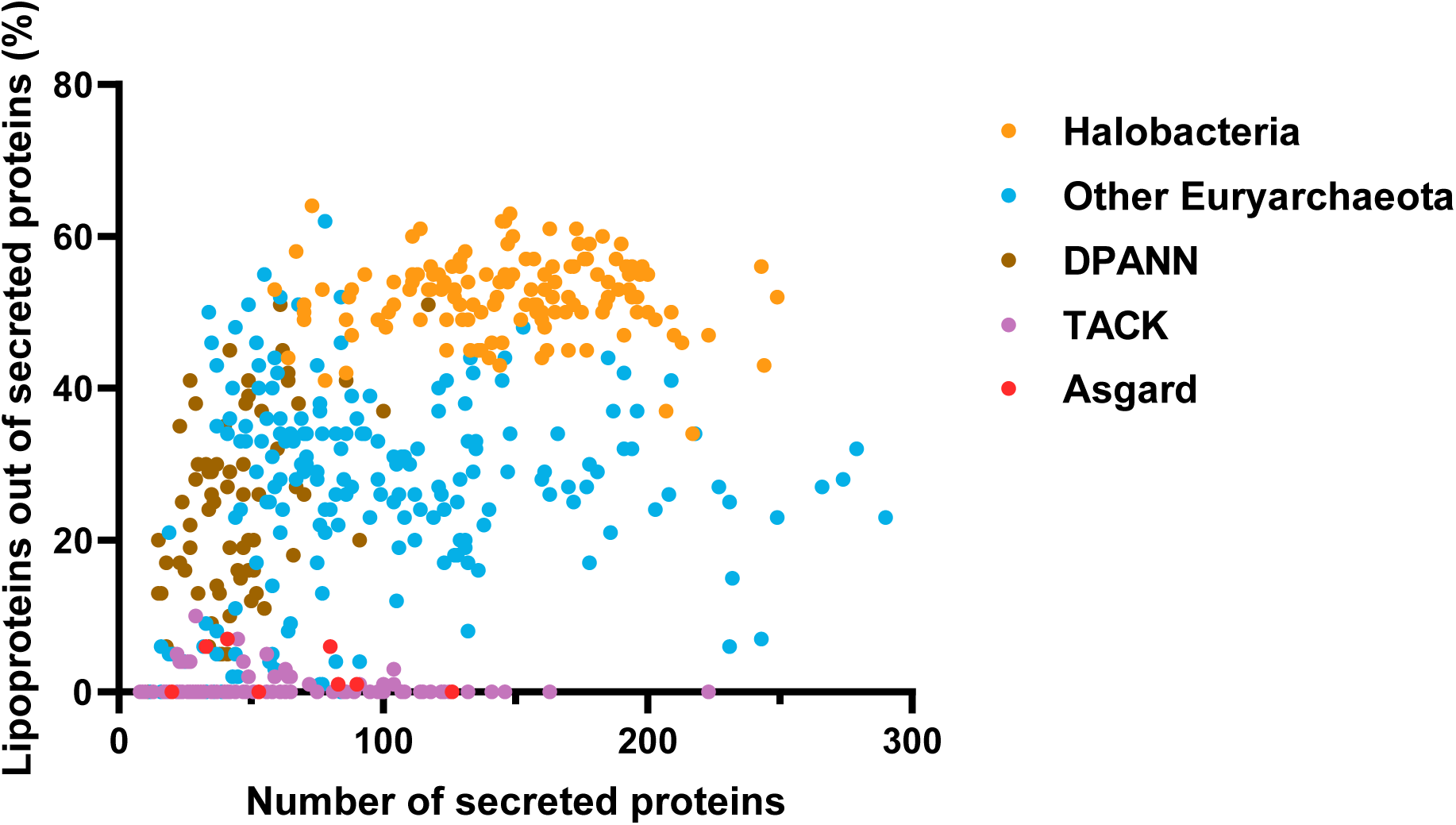
The percentage of lipoproteins out of total secreted proteins as well as the number of secreted proteins for all archaeal genomes analyzed (n = 524). Each point represents the data for one genome. Euryarchaeota, DPANN, TACK, and Asgard are the four archaeal superphyla. Halobacteria is a class in the Euryarchaeota superphylum.

**Extended Data Fig. 2.**
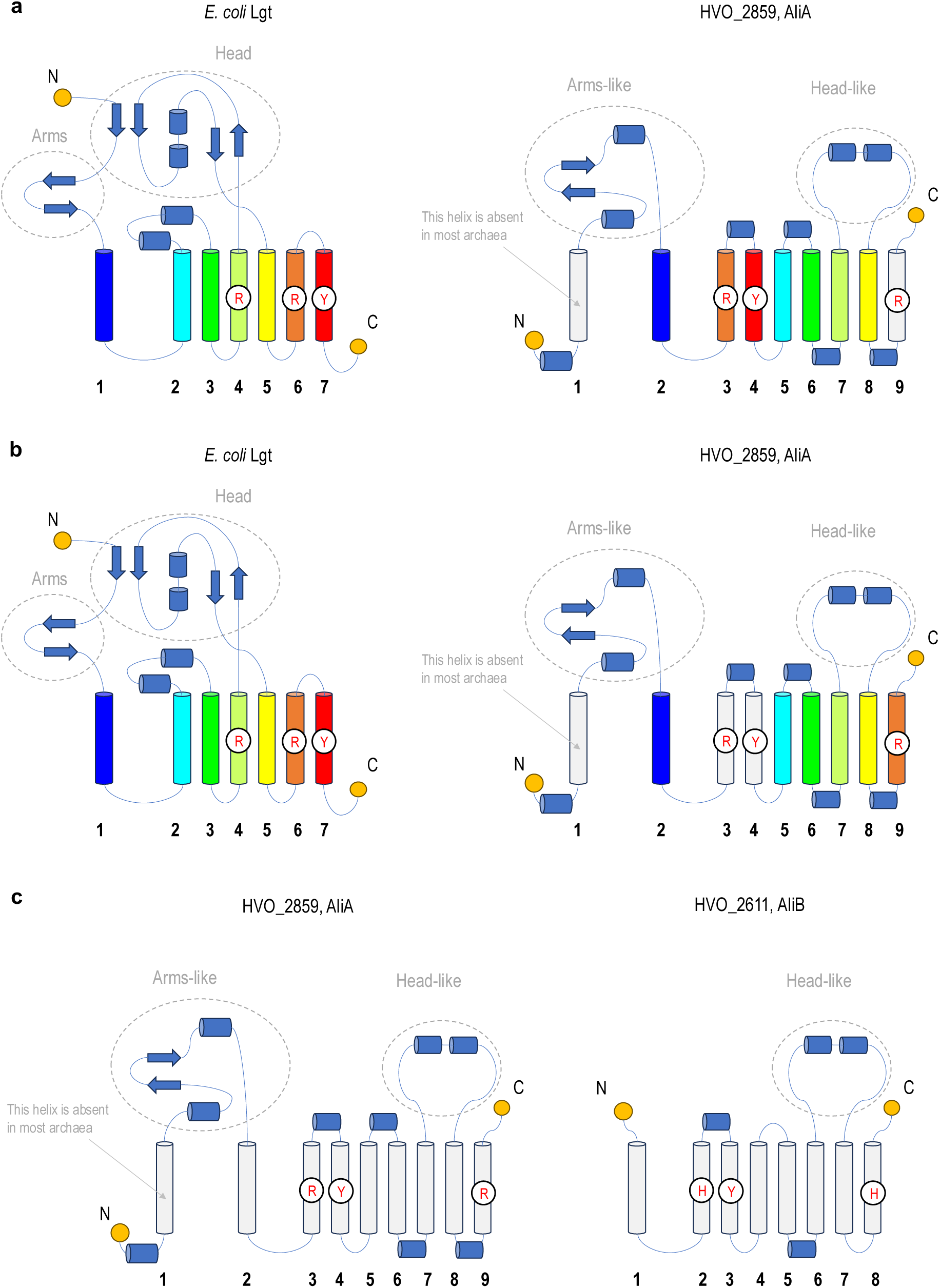
Topology comparison of *E. coli* Lgt (PDB: 5AZB) and AlphaFold2 models of HVO_2859 (AliA) and HVO_2611 (AliB). **a,b** Two possible scenarios for the origin of AliA from Lgt, with preserved connectivity of arms and head subdomains in both cases. Potentially homologous transmembrane helices are shown by the same colors in Lgt and AliA. In **a**, AliA evolves through circular permutation of the orange and red transmembrane helices, along with the emergence of a new C-terminal helix containing a catalytic arginine. In **b**, the orange helix is conserved, and AliA has two new transmembrane helices that emerged following the blue helix, with a catalytic arginine and tyrosine. Conserved residues are indicated by circle and highlighted in red. Dashed circles indicate similar subdomains between Lgt and AliA/AliB. Helices are represented as cylinders, and β-strands as arrows. **c** Topology comparison between AliA and AliB.

**Extended Data Fig. 3.**
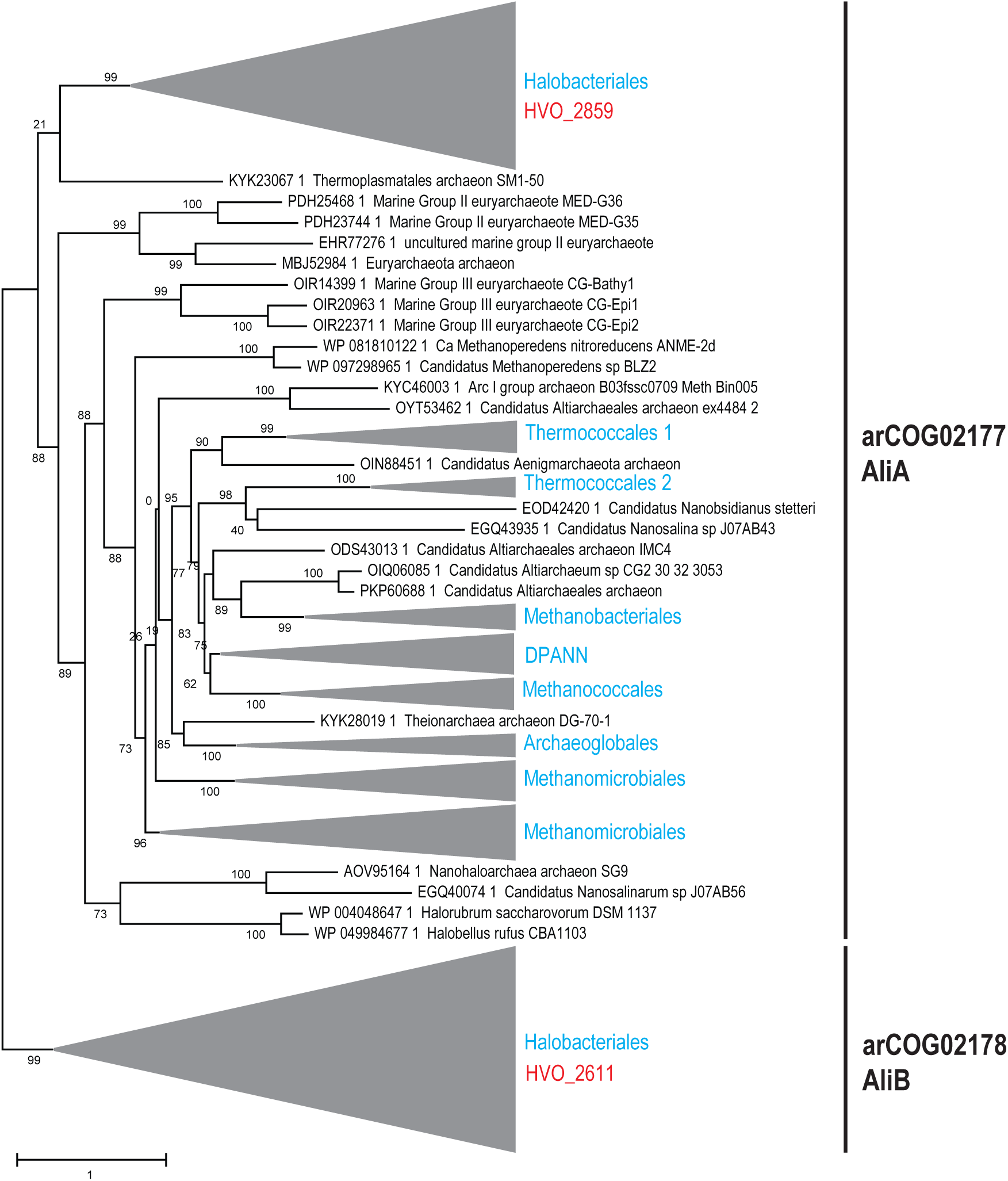
Phylogenetic analysis of AliA and AliB families. Approximate maximum likelihood unrooted phylogenetic tree was built for multiple alignment of 173 arCOG02177 representatives and 70 arCOG02178 representatives. One representative was selected from each MMseqs2^55^ cluster built with 0.8 cutoff for sequence similarity for a complete set of arCOG02177 and arCOG02178 amino acid sequences. FastTree program was also used to calculate support values. The scale bar shows the number of substitutions per site. Several branches are collapsed. Monophyletic archaeal taxa are indicated for collapsed branches on the right. Major branches corresponding to arCOG02177 and arCOG02178 are indicated by vertical bars. *Hfx. volcanii* locus IDs are indicated next to the clades to which they belong.

**Extended Data Fig. 4.**
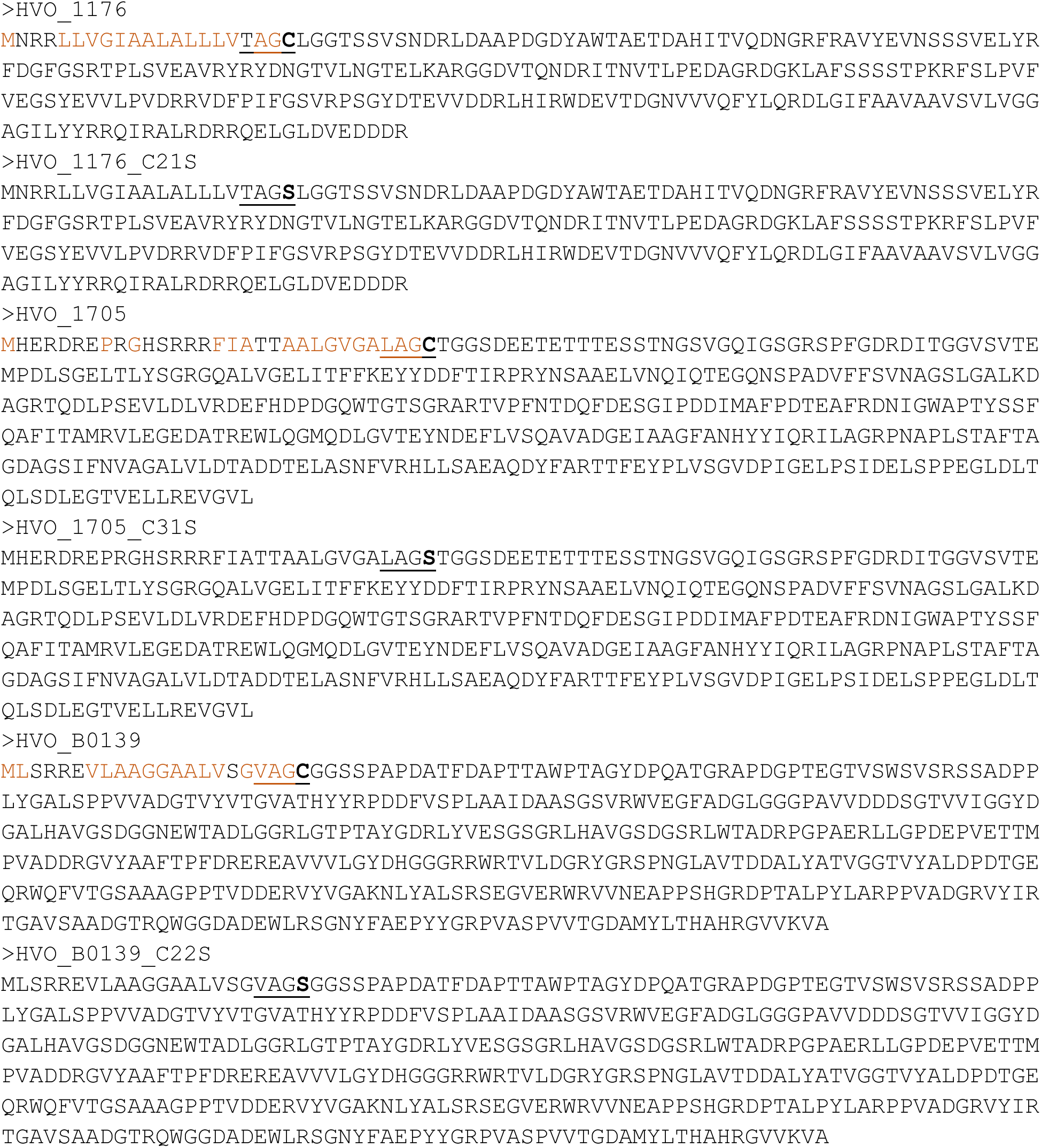
Protein sequences of the wild-type and mutant lipoproteins used in this study. The lipobox sequence is underlined, and the mutation site is indicated in bold. Hydrophobic residues in wild-type proteins are highlighted in orange.

**Extended Data Fig. 5.**
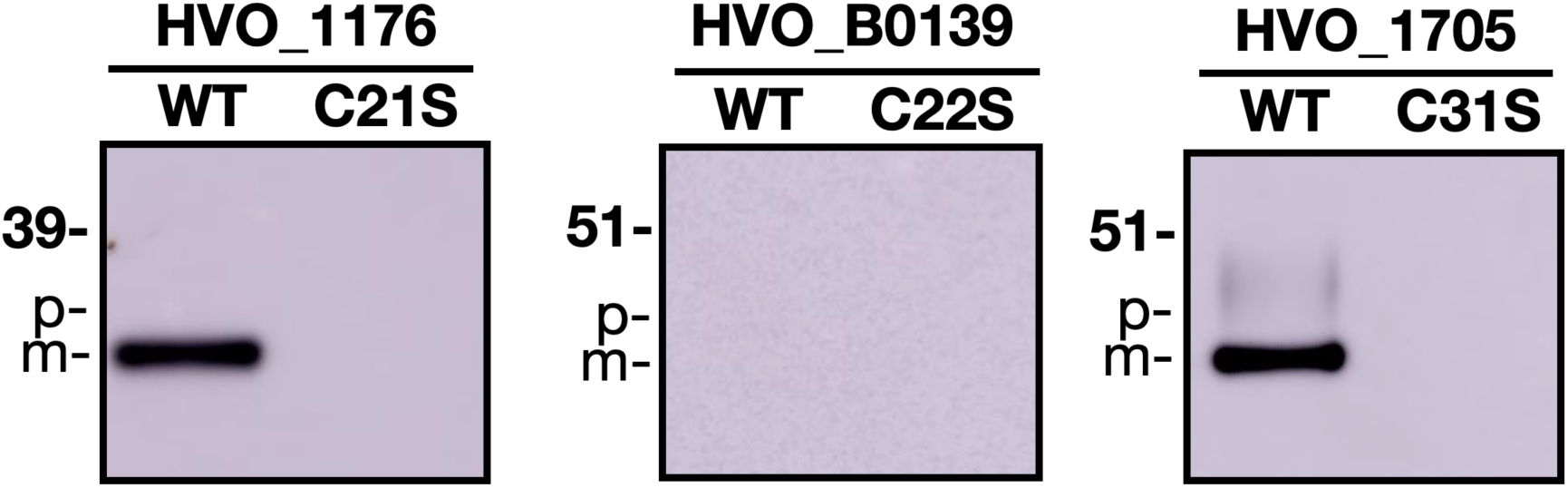
Western blots of overexpressed wild-type and mutant myc-tagged lipoproteins in *Hfx. volcanii* supernatant. The lipobox cysteine was mutated to a serine in the mutant lipoproteins. For each lipoprotein, the same amount of supernatant protein was loaded onto the gel across strains. P, precursors; M, mature proteins.

**Extended Data Fig. 6.**
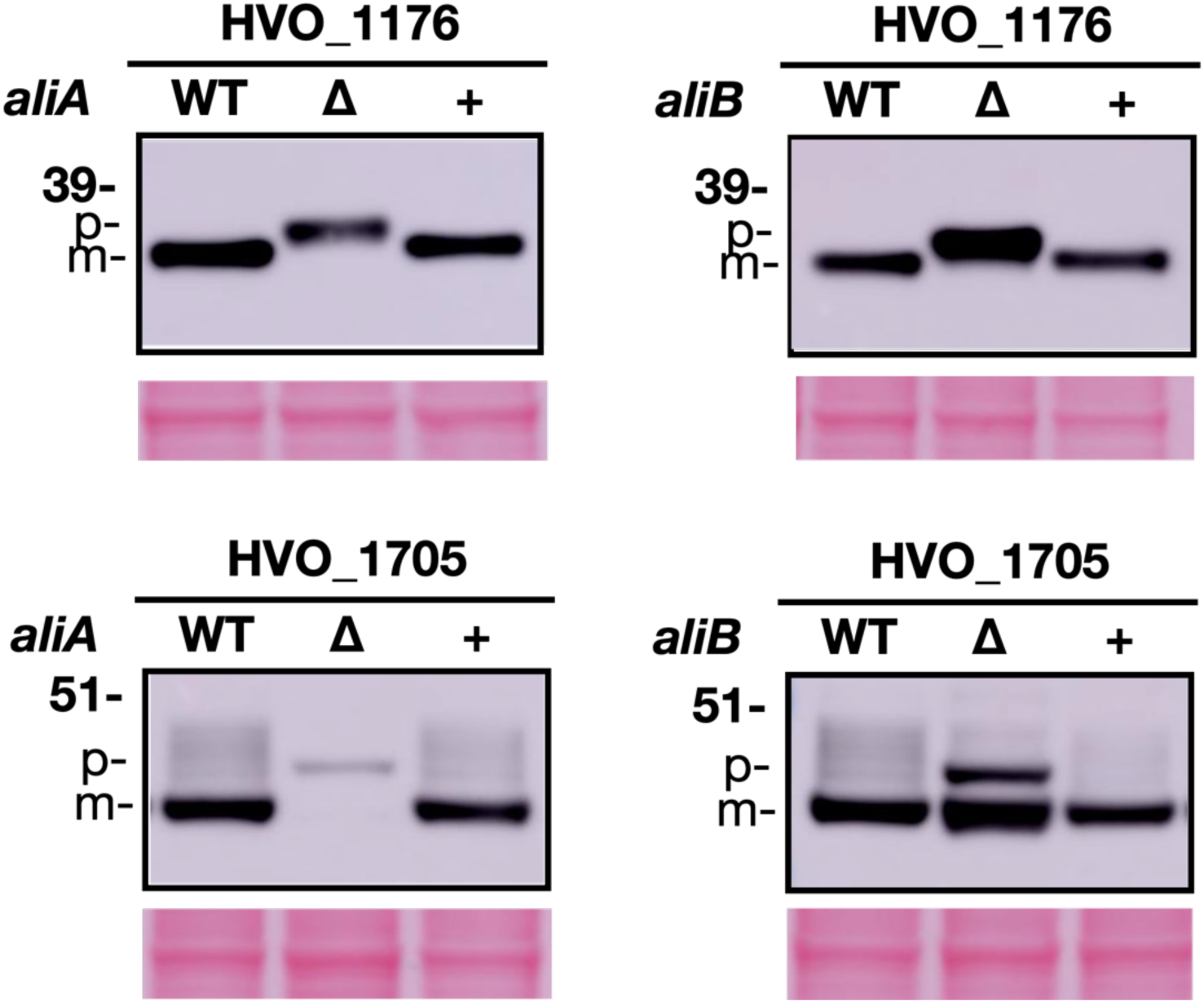
Western blots of overexpressed myc-tagged lipoproteins in cell lysates of wild-type, Δ*ali*, and Δ*ali* complementation strains. For each lipoprotein, the same amount of total protein was loaded onto the gel across strains. Ponceau S staining of the PVDF membrane is shown at the bottom as a loading control. “WT”, the presence of wild-type *ali* gene; “Δ” indicates the deletion of the wild-type *ali* gene; “+” indicates the *in trans* expression of the wild-type *ali* gene in Δ*ali* strains.

**Extended Data Table 1.** SignalP 6.0 predictions of wild-type lipoproteins and mutant lipoproteins. The probability of the most confident prediction(s) for each protein is highlighted in red.

|  | HVO_1165 |  | HVO_B0139 |  | HVO_1705_WT |  |
| --- | --- | --- | --- | --- | --- | --- |
| **Protein** | WT | C21S | WT | C22S | WT | C31S |
| **Prediction** | LIPO | LIPO | TATLIPO | TAT | TATLIPO | TAT |
| OTHER | 0 | 0.107892 | 0 | 0.000477 | 0 | 0.000088 |
| SP(Sec/SPI) | 0 | 0.405326 | 0 | 0.000234 | 0.000001 | 0.000077 |
| LIPO(Sec/SPII) | 1 | 0.483618 | 0.006148 | 0.000524 | 0.000842 | 0.000072 |
| TAT(Tat/SPI) | 0 | 0.001651 | 0.000056 | 0.810492 | 0.000834 | 0.626158 |
| TATLIPO(Tat/SPII) | 0 | 0.000768 | 0.993795 | 0.188279 | 0.998305 | 0.373593 |
| PILIN(Sec/SPIII) | 0 | 0.000721 | 0 | 0.000004 | 0 | 0.000001 |

## Notes

### Competing Interest Statement

The authors have declared no competing interest.

