## Supplementary Data 4 for "Beyond bacterial paradigms: uncovering the functional significance and first biogenesis machinery of archaeal lipoproteins"

**Supplementary Data 4. Multiple alignment of arCOG02177 proteins.** The alignment was colored using http://www.bioinformatics.org/sms2/color_align_cons.html tool with default parameters for amino acid grouping and consensus 90%. The characterized protein from *Hfx. volcanii* (WP_049914905, HVO_2859, AliA) is highlighted with red font, and potential active-site residues in that protein are highlighted in yellow.

```
WP_048201856_1 hypot -----------------------------------------MLIKKKIKDCLSD---KFKFINFWYIFNH------------------------------ 26
WP_048150344_1 hypot ---------------------------------------------------------------MWAKMSK------------------------------ 7
AOV95360_1 hypotheti ---------------------------------------------------------------------------------------------------- 0
EOD42420_1 Uncharact -------------------------------------------------MNLYQ----IFYQYFIYPVIGYQY--------------------------- 20
AOV95164_1 hypotheti ---------------------------------------------------MLE----FIWKYLAGPVIA-DARNAETAVWNGVT--------------- 29
EGQ40074_1 putative  ------------------------------------------------MREVVSE----FLWRFLAGPVVA-DALNQGSATWQGVT--------------- 33
KYK23067_1 hypotheti ---------------MNIPTTMQRYKNIVLFCILFGSLTIIVIGYLLAPTVIYD---QWIWKYYWGPVVA---------------------------DAA 55
WP_084383883_1 DUF63 ---------MSTVTRRVETALPDTGIREWWALYLLAQVVLVGVALLAFPSLVYD---RFVWQYLWGPVVA---------------------------DAA 61
WP_049984677_1 DUF63 --------MSSYFSKQTVEALPETGSKEWWLLYLLAPIVLGAISMLVFPALVYD---RFIWQYLWGPVVA---------------------------DAA 62
WP_103428047_1 hypot ---------MSTVNERVETVLPDTGTREWWALYLLAPVVLVGLGILAFPTLVYD---RFIWQYLWGPVVA---------------------------DAA 61
WP_004048647_1 DUF63 ---------MSTVTRRVETALPETGSREWWALYLLAPPVLIGAGILAFPGLVFD---QFIWQYLWGPVVA---------------------------DAA 61
WP_004594600_1 DUF63 ---------MSTVTRRVETVLPETGSREWWALYLLAPLVLLGGGILVFPTLVYD---RFIWQYLWGPVVA---------------------------DAA 61
WP_004594600_1 DUF63 ---------MSTVTRRVETVLPETGSREWWALYLLAPLVLLGGGILVFPTLVYD---RFIWQYLWGPVVA---------------------------DAA 61
WP_050050397_1 DUF63 ---------MSTVTRRVETVLPETGSREWWALYLLAPLVLLGGGILVFPTLVYD---RFIWQYLWGPVVA---------------------------DAA 61
WP_004594600_1 DUF63 ---------MSTVTRRVETVLPETGSREWWALYLLAPLVLLGGGILVFPTLVYD---RFIWQYLWGPVVA---------------------------DAA 61
WP_050050397_1 DUF63 ---------MSTVTRRVETVLPETGSREWWALYLLAPLVLLGGGILVFPTLVYD---RFIWQYLWGPVVA---------------------------DAA 61
WP_049983690_1 DUF63 ---------MSTVTHRVETALPATGTREWWVLYLLAPVVLVGAALLAFPTLVYD---RFVWQYLWGPVVA---------------------------DAA 61
WP_103428162_1 hypot ---------MSTVTRRVETVLPDTGTREWWVLFLLAPVALIGAVLLAFPTLVYD---RFVWQYLWGPVVA---------------------------DAA 61
WP_009378268_1 DUF63 ---------MSTVTRRVETALPDTGTREWWVLYLLAPVVLIGVALFAFPTLIYD---RFVWQYLWGPVVA---------------------------DAA 61
WP_015763176_1 DUF63 --------MDAVARLESRLDGRD--PLGLWVASLVGLIGLFSGAALVATQTVWD---RFVWQYFWGPVAA-DAQAARCALREGSA-VELVFSADACAQA- 84
WP_018259238_1 DUF63 --------MDAVARLESRLDGRD--PLRLWVASLVGLVGLFTGAALVATQTVWD---RFVWQYFWGPVAA-DAQAARCALREGSA-VELVFSADACAQA- 84
WP_004592615_1 DUF63 --------MET---LERAVDEYG--PGQLWVGSFVAVLLVAIGAVVAAPQAVWD---RFLWQYFWGPVYA-DAKSASCAEMTASG-PQPVY--EGCKAA- 79
WP_004518092_1 DUF63 --------MET---LERAVDEYG--PGQLWVGSFVAVLLVAVGAVVAAPQIVWD---RFLWQYFWGPVYA-DAKSASCAEMTASG-PQPLY--DGCRAA- 79
WP_014040450_1 DUF63 --------MET---FERAVDEYG--PGQLWVGSFAAVLLVAVGAVMAAPQAVWD---RFLWQYFWGPVYA-DAKAASCAEMTASG-PQPLY--DGCRAA- 79
WP_014040450_1 DUF63 --------MET---FERAVDEYG--PGQLWVGSFAAVLLVAVGAVMAAPQAVWD---RFLWQYFWGPVYA-DAKAASCAEMTASG-PQPLY--DGCRAA- 79
WP_008309018_1 DUF63 --------MET---FERAVDEYG--PGQLWVGSFVAVLLVAVGAVMAAPKAVWD---RFLWQYFWGPVYA-DAKAASCAEMTASG-PQPLY--DGCRAA- 79
WP_014040450_1 DUF63 --------MET---FERAVDEYG--PGQLWVGSFAAVLLVAVGAVMAAPQAVWD---RFLWQYFWGPVYA-DAKAASCAEMTASG-PQPLY--DGCRAA- 79
WP_014040450_1 DUF63 --------MET---FERAVDEYG--PGQLWVGSFAAVLLVAVGAVMAAPQAVWD---RFLWQYFWGPVYA-DAKAASCAEMTASG-PQPLY--DGCRAA- 79
WP_005534235_1 DUF63 --------MNT---FERAVDDYG--PGQLWVGSFVAVLLVAVGAVVAAPQAVWD---RFLWQYFWGPVYA-DAKSASCAEMTASG-PQPVY--EGCKAA- 79
WP_004961153_1 DUF63 --------MDT---LERAVDEYG--PGQLWVGSFVAVLLVAVGAVTAAPQAVWD---RFLWQYFWGPVYA-DAKSASCAEMTASG-PQPLY--DGCTAA- 79
WP_004961153_1 DUF63 --------MDT---LERAVDEYG--PGQLWVGSFVAVLLVAVGAVTAAPQAVWD---RFLWQYFWGPVYA-DAKSASCAEMTASG-PQPLY--DGCTAA- 79
WP_101350151_1 hypot --------MET---FERAVDEYG--PGQLWVGSFVAVLLVAVGAVMAAPQAVWD---RFLWQYFWGPVYA-DAKSASCAEMTASG-PQPLY--DGCRAA- 79
WP_053968273_1 DUF63 --------MET---FERAVDEYG--PGQLWVGSFVAVLLVAVGAVMAAPQAVWD---RFLWQYFWGPVYA-DAKSASCAEMTASG-PQPLY--DGCSAA- 79
WP_058995687_1 DUF63 --------MET---FDRAVDEYG--PGQLWVGSFVAVLLVAVGAVMAAPQAVWD---RFLWQYFWGPVYA-DAKSASCAEMTASG-PQPLY--EGCKAA- 79
WP_015790673_1 DUF63 ----------------MDETGFDLTPGRAWLAVFAAGVVTIVGASIAFTRTVWD---EFIWQYFWGPVYA-DAHNAGCAIKEGGQ---TLLGGSGFAENC 77
WP_008524115_1 DUF63 ----------------MDETGFDLTPGRAWLAVFAAGVVAVVGGSIAFTRTVWD---EFIWQYFWGPVYA-DAYNAGCAIKAGGE---TLLGGSGFAENC 77
WP_075936143_1 DUF63 --------------MDSPRSWS---RAQAWLATFTTALLAFGFGVLLAPELVWD---RFLWHYFWGPVYA-DANNAVCVVMDGGT-TELFYSTVACQDA- 77
WP_020446311_1 DUF63 --------------MDDLFDRID--PEKAWALVVAGLALAGTLGSLLFPKQVYD---QWIWKYYWGPVAA-DGNGAKCAVRESGD-VTYYSDSAQCAGA- 78
WP_049898450_1 DUF63 --------------MAAIADRVD--PVRGWLAAAIAAVIVVAGSAVAFPQRVYS---GFLWRYFWGPIDA-DAHNAACAVRTDGV-VRRFAEESACQAA- 78
WP_006077061_1 DUF63 --------------MAAIADRVD--PVRGWIAAALAVVLVVAGSAVTFPQRVYT---EFLWQYFWGPIDA-DAHNAVCSVRVDGV-VSRLGEANACQAA- 78
WP_049996700_1 DUF63 --------------MAAIADWTD--PARTWLAAAVAGTVAIIAAALAFPRAVYT---GFIWRYFWGPIDA-DAHNAACAVRADGV-VRRFAESETCRAA- 78
EMA38628_1 hypotheti --------------MAAIADRTD--PVRTWLAAAVVGVVAIAGASLAFPKAVYT---NFVWQYFWGPIDA-DAHNAVCAIRANGM-VERLGTADACQVA- 78
WP_049992561_1 DUF63 ---------------MDVIDRIG--AVRLWVAAFLASVAGLATGSTLAHRTVYD---GFIWRYFWGPVAA-DAHNAQCAINDGES-VTYGYADAVCTGV- 77
WP_010903025_1 DUF63 ------------MTFPDTT------AERAWTAAVGCVVVALVGGAVVFPDTVYG---GFVWQYFWGPVVA-DAQGAHCAAWNGGA-VELLATKGACAGA- 76
WP_009760947_1 DUF63 ------------MSFPETD------RERTWTAVVGLALVALVVGSLLLRDTVYG---GFIWHYFWGPVYA-DAQGAQCAVWNGGN-VELLSSTAACAAA- 76
WP_059057097_1 DUF63 ------------MTFPETD------RERAWTAVVGVAVLALVGGSLLFPDTVYG---DFVWHYFWGPVVA-DANGAQCAVWNGGA-VQHVYSSTACATA- 76
WP_058983522_1 DUF63 ------------MTFPETD------RERAWTAVVGVALAALVAGSILFPDSVYG---GFVWHYFWGPVVA-DANGAQCAAWNGGA-VRLLYNSSACTAA- 76
WP_071932813_1 DUF63 ------------MAFPSTR------IERLWTGAVGTILAVLVGGSVLFPERVYG---GFVWQYFWGPVAA-DAHGAQAAAYNGGD-PIFFDSVAGASGV- 76
WP_050048584_1 DUF63 -------------MFPETT------TERAWTAAVIGIVAALVGGSLLFPDLVYG---SFVWQYFWGPVYA-DAHNAACAAWNGGT-PQLYASSGACEAA- 75
WP_014051341_1 DUF63 --------------MSSLQERLGVAPERLWLAAFGSLVLLLGGGSLLFPETVYD---GFLWHYFWGPVQA-DAHNAVCAVRPGST-VEYLYTASECAAA- 80
WP_079233299_1 DUF63 --------------MFSLRERADENPERLWAGTVGALLVALVGGSLIAPETVYD---GFIWHYFWGPVQA-DAHNAVCAIRPGST-VEYLYSSSACQAA- 80
WP_053948572_1 DUF63 --------------MFSLREWADGNPERLWAGVVGTLLVALVGGSILAPETVYD---GFIWHYFWGPVQA-DAHNAVCAVRPGST-VEYLYSTSACQAA- 80
WP_049980835_1 DUF63 --------------MFSLRERVDENPARAWAAVVGALLAALIGGSILFPETVYD---GFVWHYFWGPVQA-DAHNAVCAVRPGST-VEYLYSTSACQGA- 80
KPN31749_1 hypotheti -------------------------------------------------------------------MQA-DAHNAVCAVRPGST-VEYLYSSSACQAA- 30
AGB16040_1 putative  --------------MQQYIDRYG--AERIWAATVVVVFGGLALLAALFPQQVYV---EFIWEYFWGPVVA-DANNWNCVAWAGGE-------VQSCDVA- 72
WP_007696755_1 DUF63 --------------MQQYIDRFG--AERIWAATVVVVFGGLALLAALFPQQVYV---EFIWEYFWGPVVA-DANGWNCVAWAGGE-------VQSCDVA- 72
WP_015322904_1 DUF63 --------------MDEYIERFG--AVRIWAATVLVLAAAIVALAVAFPQRVYV---DLIWQYYWGPVVA-DAHGWNCVAWAGGA-------EIPCNEA- 72
WP_076581869_1 DUF63 --------------MDEYIERYG--AERLWVATVVLLAGAVALAAFLFPQRVYV---DFIWQYYWGPVVA-DAHGWNCVTWADGD-------QLECGEV- 72
WP_005559622_1 DUF63 --------------MDEYIERFG--PVKIWAATVIVLAVAIAALAVAFPQRVYV---DLIWQYYWGPVVA-DAHGWNCVAWADGA-------ELPCNEA- 72
WP_006067640_1 DUF63 --------------MDEFIERYG--PERVWAATVVLLVAGVTLAAWVFPQRVYV---DIIWQYFWGPVVA-DAHGWGCIAWAGGE-------QIHCNEA- 72
WP_008164554_1 DUF63 --------------MDEFIERYG--PERVWAATVVLLAVGVTLAAWVFPQRVYV---DIIWQYFWGPVVA-DAHGWSCIAWAGGE-------QIHCNEA- 72
WP_006088112_1 DUF63 --------------MDEYIERYG--AEKVWAAAVVLIFGGVAIAALLFPQRVYV---DFIWQYYWGPMVA-DAHGWGEVAWAGGE-------QINANEA- 72
WP_012943247_1 DUF63 --------------MDEYIERYG--AERLWTATVALLAAAVALAAVLFPQRIYV---DLIWQYYWGPVVA-DAHGWSCVAWADGN-------PVHCSEV- 72
WP_008895026_1 DUF63 --------------MDEYIERYG--AERIWAAAVVLLAAGVALAASLFYQPVYV---DLIWQYYWGPVVA-DAHGWSCIAWADGQ-------QIHCSNA- 72
WP_098727043_1 DUF63 --------------MDDVIDRFG--AERVWVATVASLAAAVVLGALLFPQRVYV---EIIWQYFWGPVVA-DAHSWNCATLVDGQ-------AVSCAEA- 72
WP_049990861_1 DUF63 --------------MDDFIDRYG--AERVWAATVASLATIVVLGAVLFPQRVYV---EIIWQYFWGPVVA-DAHSEPCLAWADGQ-------QVPCGDN- 72
WP_008013001_1 DUF63 --------------MDDFIDRYG--AERVWVATVASLAAIVVLGAVLFPQRVYV---EFIWQYFWGPVVA-DAHSWSCVAWADGQ-------QVACGQA- 72
WP_006180566_1 DUF63 --------------MDDFIDRFG--AERVWAATVATLAVAVTLGAVIFPQRVYV---EFIWQYFWGPVVA-DAHSWNCVAWAGGE-------QVPCTEA- 72
WP_006650865_1 DUF63 --------------MDDFIDRFG--AERVWAATVATLAVVVALGAVIFPQRVYV---EFIWQYFWGPVVA-DAHSWNCVAWAGGE-------QVPCTEA- 72
WP_066301295_1 DUF63 --------------MDDFIDRFG--AERVWAATVATLVAVVALGAVVFPQRVYV---EFIWQYFWGPVVA-DAHSWNCVAWAGGE-------QVPCTEA- 72
WP_049966804_1 DUF63 --------------MDDFIDRFG--AERVWAATVATLAVVVALGAVIFPQRVYV---EFIWQYFWGPVVA-DAHSWNCVAWAGGE-------QVPCTEA- 72
WP_076145380_1 DUF63 --------------MDDFIDRFG--AERVWAATVATLVAVVALGAVVFPQRVYV---EFIWQYFWGPVVA-DAHSWNCVAWAGGE-------QVPCNEA- 72
WP_097378845_1 DUF63 --------------MDDFIDRVG--PERVWALTVASLAAIVALGAVLFPQRVYV---EFIWQYFWGPVVA-DAHSWNCVAWANGQ-------EIACSEA- 72
WP_008452261_1 DUF63 --------------MDDFIDRFG--AERVWAATVASLAAVVVLGAVLFPQRVYV---DFIWQYFWGPVVA-DAHSWECVAWAGGQ-------ELPCSEA- 72
WP_008452261_1 DUF63 --------------MDDFIDRFG--AERVWAATVASLAAVVVLGAVLFPQRVYV---DFIWQYFWGPVVA-DAHSWECVAWAGGQ-------ELPCSEA- 72
WP_006432643_1 DUF63 --------------MDDFIDRFG--AERVWAATVASLAAVVVLGAVLFPQRVYV---DIIWQYFWGPVVA-DAHSWNCVAWAGGQ-------ELPCGEA- 72
WP_008452261_1 DUF63 --------------MDDFIDRFG--AERVWAATVASLAAVVVLGAVLFPQRVYV---DFIWQYFWGPVVA-DAHSWECVAWAGGQ-------ELPCSEA- 72
WP_007109662_1 DUF63 --------------MDDFIDRFG--AERVWAATVASLAAAVVLGAVLFPKRVYV---DFIWQYFWGPVVA-DAHSWECVAWAGGQ-------ELPCGEA- 72
WP_049952821_1 DUF63 --------------MDDFIDRYG--PVRVWAATVALLASAITLGAILFPQRVYV---DIIWHYFWGPVVA-DAHGWSCVTWAGGE-------QLECAAA- 72
WP_086889541_1 DUF63 --------------MYDYVERYG--PVKVWAATVLVLAVGIAAAAMAFPQQVYR---DLIWQYYWGPVVA-DAHGWGCIDWAGGQ-------ELHCTEAA 73
```

```
WP_005580009_1 DUF63 --------------MYEYIERYG--PERVWAVAVLVLAVGIALLAVRFPQRVYV---DLIWQYYWGPVVA-DAHGWGCVDWAGGD-------PMACNAAI 73
WP_049927356_1 DUF63 --------------MYERIERYG--PLKVWAAAVLALAAAVALAAYAFPQRVYI---DIIWQYYWGPVVA-DAHGWSEVAWAGGE-------QIEGARA- 72
WP_087714455_1 DUF63 --------------MYEYIERYG--PERVWAATVLVLAAGVALAAFLFPNRVYV---DLIWQYYWGPVVA-DAHGWGEIAWAGGE-------QIHAAEA- 72
WP_013878438_1 DUF63 --------------MYESIERYG--PERVWAAAVVLLVAGIAAAATAFPRQVYV---EFIWQYYWGPVVA-DAHSWSCVAWAGSE-------QIPCSQA- 72
WP_049921111_1 DUF63 --------------MYEYVERYG--PERIWAATVALLAVAVTLAAITFPQRIYI---DIIWQYFWGPVVA-DAQDMNCIAWAGGE-------QVPCGEA- 72
WP_007142789_1 DUF63 --------------MYESVERYG--PERVWAAAVLVLASAVILLAARFPQRVYV---DIIWQYYWGPVVA-DAHGWSCVAWAGGE-------QVPCGEA- 72
WP_006651941_1 DUF63 --------------MYETVERYG--PLKVWALTVVALAAAVIFAATAFPERVYV---DIIWQYYWGPVVA-DAHGWSEVAWAGGE-------QIPASEA- 72
WP_004216437_1 DUF63 --------------MYETVERYG--PLKVWALTVVALAAAVIFAATAFPERVYV---DIIWQYYWGPVVA-DAHGWSEVAWADGE-------QIPASEA- 72
WP_071402513_1 DUF63 --------------MYETVERYG--PLKVWALTVVALAAAVIFAATAFPERVYV---DIIWQYYWGPVVA-DAHGWSEVAWAGGE-------QIPASEA- 72
WP_006666462_1 DUF63 --------------MYESIERYG--PLKVWALTVVTLVAAVALAATAFYQPVYV---DIIWQYFWGPVVA-DAHGWNCVAWAGGE-------QLPCGEA- 72
WP_049904535_1 DUF63 --------------MYESIERYG--PLKVWALTVVTLIAAVALAATAFYQPVYV---DIVWQYFWGPVVA-DAHGWNCVAWAGGE-------QLPCGEA- 72
WP_006824405_1 DUF63 --------------MYESIERYG--PLKVWALTVVTLVAVVALAATAFYQQVYV---DIIWQYFWGPVVA-DAHGWNCVAWAGGE-------QLRCGEA- 72
WP_011323802_1 DUF63 ------------MDSSD------FDLARMWLAVFVGGIVAIVAAAAAFPRRVYN---GFLWQYFWGPVDA-DAHGAVCAVRSGGT-TERLYSRAACSGA- 76
WP_015409719_1 DUF63 ------------MQLPD-----GVDPVRAWLATCAGGIALLGGGAVAFPGRVYD---GFLWRYFWGPVDA-DAHGASCAVRSDGT-TERLFDGLACSNA- 77
WP_006883947_1 DUF63 ------------MENEQVLT------ERTWLGAFVALVAALAVGSLVATERVYD---RFIWRYFWGPIYS-DANNARCAVLTGDG-IDLLGSTAACQTA- 76
WP_077207545_1 DUF63 ---------------MQVFERM--DAGRAWTAAALAGIAALVLGSLVFYDTVYR---GFVWHYFWGPVYA-DAHNAVAAVHQGDS-VELLYSEAARQQA- 77
ESS12903_1 putative  --------------MSSAERLSGISPETAWAAAVASVATIMVGGALALPRLVWD---QFLWRYFWGPVIA-DANGAVCAIRAGGQ-TRFGASAADCTSA- 80
WP_008417361_1 DUF63 --------------MNDLSARI--DPARAWVAAALAALAALVVGSVVFPRRVYD---GFFWRYFWGPVVA-DGTGSTCARRVGGE-TRLLDSAVECQQA- 78
WP_066381608_1 DUF63 --------------MDDVSDRI--DPARAWVAVALTVLTALVVGAFAFPRQVYD---GFLWRYFWGPVVA-DGTGSACAQRIDGR-T-VLDG--DC--L- 73
WP_049947642_1 DUF63 ------------MSAASLADRLGTTPVRAYLLAVGAAVLALAAGCVLFPETVYD---GFVWHYFWGPVQA-DANSALCAVREGAG-TTYLYSQSECAAA- 82
ESS05953_1 putative  ---------------MSIASRVEADPERAWVAAALAVLLAVAGGSVAFPRAVYD---GFVWQYFWGPVAA-DAHAATCAVRAGGT-TTYLDAAGSCTSA- 79
WP_096390051_1 DUF63 -------------MSDRLDERLDVSPETAWLATVVSIVIGVGAGAVAFPRTVYD---GFLWRYFWGPVVA-DGEGAACAVRRGGVTEYLYSSDACAEAA- 82
WP_021072749_1 DUF63 -------------MSDWLDDRLDVSPEVAWIATLISIVVGLTAGAALFPRTVYD---GFLWRYFWGPVVA-DGEGAACAVRRGGGTEYLYSSDACAEAA- 82
WP_049982791_1 DUF63 -------------MTTD-GARVGLSPERAWATAVGVVAAVLAVGSAAFPRIVYD---RFLWRYFWGPVAA-DGQGAQCAVRDADG-TTLLDSTAACAEA- 80
WP_008585990_1 DUF63 -------------MSTAVADRFDLSPERAWAAVVGGVTALLAVGSVVFTRVVYD---RFLWRYFWGPVVA-DGEGAQCAVRNPDGTTDLLGGEAACQSA- 82
WP_006628721_1 DUF63 -------------MST--ADRFDLTPEQAWAAVVGGVTALLVVGSVVSPRVVYD---RFLWRYFWGPVVA-DGEGAQCAVRSAGGTTELIGSEAACQSA- 80
WP_006113218_1 DUF63 -------------MSTGVADGFDPSPERAWAAVVGGVTALLAIGSVVFPRVVYD---RFLWRYFWGPVVA-DGEGAQCAVRDAGG-TELLGSSAACQSA- 81
WP_049930034_1 DUF63 -------------MSTAVADGLDLSPERAWAAVVGGVTALLAIGSVVFPRVVYD---RFLWRYFWGPVVA-DGEGAQCAVREAGGTTELLGSSAACQSA- 82
WP_049906047_1 DUF63 -------------MSTGVADGFDPSPERAWAAVVGGITALLAVGSLVFPRVVYD---RFLWRYFWGPVVA-DGEGAQCAVREVGGTTELLGSSAACQSA- 82
WP_044965494_1 DUF63 -------------MSTGVADGFDPSPERAWAAVVGGVTALLAIGSVVFPRVVYD---RFLWRYFWGPVVA-DGEGAQCAVREPGGTTELLGSSGACQAA- 82
WP_096393195_1 DUF63 -------------MSTGVADGFDPSPERAWAAVVGGVTALLAVGSVVFPRVVYD---RFLWRYFWGPVVA-DGEGAQCAVREAGGTTELLGSSAACQAA- 82
WP_049908400_1 DUF63 -------------MSTGVANGFDPSPERAWAAVVGGVTALLAIGSVVFPRVVYD---RFLWRYFWGPVVA-DGEGAQCAVREAGGTTELLGSSAACQAA- 82
WP_049908585_1 DUF63 -------------MSTGVADGFDPSPERAWAAVVGGVTALLAIGSVVFPRVVYD---RFLWRYFWGPVVA-DGEGAQCAVREAGGTTELLGSSAACQAA- 82
WP_049902631_1 DUF63 -------------MSTGVADGFDPSPERAWAAVVGGVTALLAIGSVVFPRVVYD---RFLWRYFWGPVVA-DGEGAQCAVREAGGTTELLGSSAACQAA- 82
WP_049983668_1 DUF63 -------------MSTGVADGFDPSPERAWAAVVGGVTALLAVGSVVFPRVVYD---QFLWRYFWGPVVA-DGEGAQCAVRDAGG-TELLGSTAACQAA- 81
WP_053772267_1 DUF63 -------------MSTGLADGFDPSPERAWAAVVGGVTALLAVGSVVFPRVVYD---RFLWRYFWGPVVA-DGEGAQCAVRDAGG-TELLDSTAACEAA- 81
WP_007999151_1 DUF63 -------------MSTGFGDR-DLSPERAWAAVVSAITVLLALGSIVFPRVVYD---RFIWQYFWGPVVA-DGQGAQCAVRDGGG-TELLTSSAACSAA- 80
WP_015909917_1 DUF63 -------------MSTRVEERFGLSPERAWAAVVGAVAALLAIGSAVFPRIVYD---RFIWQYFWGPVAA-DGQGAQCAIHDAGG-TELLGSSSACTAA- 81
WP_004050594_1 DUF63 -------------MSTRVEERFGLSPERAWAAVVGAVAVLLAVGSAVFPRIVYD---RFLWQYFWGPVAA-DGQGAQCAVRDAGG-TELLGSSSACSAA- 81
WP_095636035_1 DUF63 -------------MSTRVEGRFGLSPERAWAAVVGAVTALLAVGSVVFPRIVYD---RFLWRYFWGPVAA-DGQGAQCAVRDASG-TELLGSSSACSAA- 81
WP_008003811_1 DUF63 -------------MNTRVEERFGLSPERAWAAVVGAVTVILAVGSAVFPRIVYD---RFLWRYFWGPVAA-DGQGAQCAVRDAGG-TELLGSSSACTAA- 81
WP_066416034_1 DUF63 -------------MSARFEDRLGLSAERAWATAVAAVSAVLVVGSVLLPRVVYD---RFLWRYFWGPVAA-DGQVAQCAVRPADGGTELLTSAAACADA- 82
ESS03170_1 putative  -------------MSVRAADRFGLSPERAWGTAVAVVTGLLVVGSIVFPRVVYD---RFLWRYFWGPVAA-DGQGARCAVRTA-GGTELLSSASACAEA- 81
WP_089671383_1 DUF63 --------------MAAVSERLDTSSERLWIGTGSVVALVLLVGSLLFRELVYG---RFLWQYFWGPVVA-DGNGANCAIRSGGT-VDFGGSASACTQA- 80
ERH07656_1 putative  --------------MAGLSARVDTDPERLWLGGVGTLTALLVFGSLLARELVYD---RFIWQYFWGPVAA-DGNGARCAVNRAGD-VSYLFSTAECVAA- 80
ESS10096_1 putative  --------------MAGLSARVDTDPERLWLGGVGTLTALLVFGSLLARELVYD---RFIWQYFWGPVAA-DGNGARCAVNRAGD-VSYLFSTAECVAA- 80
ESS07824_1 putative  --------------MAAVTDRVDAEPERLWLATFGVLTAALVGGSLLFRERVWD---RFLWQYFWGPVAA-DGNGAVCAVNGASG-----VEYLASTAAC 77
ERH05689_1 putative  --------------MAAVTDRVDAEPERLWLATFGVLTAALVGGSLLFRERVWD---RFLWQYFWGPVAA-DGNGAACAVNSASG-VEYLASTAACTTS- 80
ERH02266_1 putative  --------------MAAVTDRVDAEPERLWLATFGVLTAALVGGSLLFRERVWD---RFLWQYFWGPVAA-DGNGAACAVNSASG-VEYLASTAACTTS- 80
ESS07825_1 putative  ---------------------------------------------------------------------------------------------------- 0
WP_049970017_1 DUF63 --------------MDTASE--GIDNERTWAGAIAAAIVALVGGALAFPKQVYD---GFIWHYFWGPVVA-DAKNAPCAVREDGT-TRLVSDAGTCATL- 78
WP_007979289_1 DUF63 --------------MDTASE--RLGNERAWAGAVATAVVALVGGALVFPKWVYD---GFIWHYFWGPVVA-DAKGAACAVREGGT-TRFIDSASACASA- 78
WP_007977745_1 DUF63 --------------MDTASE--EADNERLWMGAVAAAIVALVGGALAFPKQVYD---GFIWHYFWGPVLA-DANNASCAVRDGGT-TKYIYDAATCATA- 78
WP_014556445_1 DUF63 --------------MSTVAARLDTSPERLWVGTVSTTVAVLVIGSLVAPETIYD---SFLWHYFWGPVQA-DANSAVCAVRPGST-VQYLTSTTACANA- 80
WP_049935775_1 DUF63 --------------MATVAERLDVEPERLWAGSVLALLAALIGGSLLFPRTVYD---GFVWHYFWGPVQA-DANAAICAVREGGA-TRYLYESAACSAA- 80
WP_008325149_1 DUF63 --------------MATVAERTGLDPERLWGVGVVAILVALVGGSLAFPRVVYD---GFIWHYFWGPVQA-DANSAVCATRAEGI-TQYLDSASACAAA- 80
WP_007543631_1 DUF63 --------------MATVAERTGLDPERLWGVGVVAILVALVGGSLAFPRVVYD---GFIWHYFWGPVQA-DANSAVCATRAGGV-TQYLDSASACAAA- 80
WP_049905041_1 DUF63 --------------MATVAERMGLDPERLWGLGVAAILVALVGGSLAFPRVVYD---GFIWHYFWGPVQA-DANSAVCAIRANGA-TQYLDTASACAAA- 80
WP_049913563_1 DUF63 --------------MATVAERMGLDPERLWGLGVAAILVALVGGSLAFPRVVYD---GFIWHYFWGPVQA-DANSAVCAVRANGV-TQYLDTASACAAA- 80
WP_049967851_1 DUF63 --------------MATVAERIGLDPERLWGLGVAAILVALVGGSLAFPRVVYD---GFIWHYFWGPVQA-DANSAVCAVRANGV-TQYLDTASACAAA- 80
WP_049914905_1 DUF63 --------------MATVAERMGLDSERLWGLGVAAILVALVGGSLAFPRVVYD---GFVWHYFWGPVQA-DANSAVCAVRAGGV-TQYLDTASACTAA- 80
WP_049916430_1 DUF63 --------------MATVAERMGLDSERLWGLGVAAILVALVGGSLAFPRVVYD---GFIWHYFWGPVQA-DANSAVCAVRADGV-TQYLDTASACTAA- 80
WP_049896892_1 DUF63 --------------MATVAERMGLDSERLWGLGVAAILVALVGGSLAFPRVVYD---GFIWHYFWGPVQA-DANSAVCAVRAGGV-TQYLDTASACAAA- 80
WP_049896892_1 DUF63 --------------MATVAERMGLDSERLWGLGVAAILVALVGGSLAFPRVVYD---GFIWHYFWGPVQA-DANSAVCAVRAGGV-TQYLDTASACAAA- 80
WP_049896892_1 DUF63 --------------MATVAERMGLDSERLWGLGVAAILVALVGGSLAFPRVVYD---GFIWHYFWGPVQA-DANSAVCAVRAGGV-TQYLDTASACAAA- 80
WP_058828253_1 DUF63 --------------MATVAERMGLDPERLWGLGVAAILVALVGGSLAFPRVVYD---GFIWHYFWGPVQA-DANSAVCAVRANGV-TQYLDTASACAAA- 80
WP_058568959_1 DUF63 --------------MATVAERMGLDSERLWGLGVVAILVALVGGSLAFPRVVYD---GFIWHYFWGPVQA-DANSAVCAVRANGV-TQYLDTASACAAA- 80
WP_049917947_1 DUF63 --------------MATVAERMGLDPERLWGLGVAAILVALVGGSLAFPRVVYD---GFIWHYFWGPVQA-DANSAVCAVRANGV-TQYLDTASACAAA- 80
WP_049920348_1 DUF63 --------------MATVAERMGLDPERLWGLGVAAILVALVGGSLAFPRVVYD---GFIWHYFWGPVQA-DANSAVCAVRAGGV-TQYLDTASACAAA- 80
WP_089777545_1 DUF63 --------------MATVAERMGLDPERLWGLGVAAILVALVGGSLAFPRVVYD---GFIWHYFWGPVQA-DANSAVCAVRANGV-TQYLDTASACAAA- 80
WP_008320633_1 DUF63 --------------MATVAERVGLDPERLWGVGVVALLVALVGGSLAFPRVVYD---GFIWHYFWGPVQA-DANSAVCAVRAGGV-TQYLDSASACAAA- 80
WP_004060355_1 DUF63 --------------MATVAERVGLDPERLWGVGVVTLLVALVGGSLAFPRVVYD---GFIWHYFWGPVQA-DANSAVCAARSGGV-TQYLDSASACAAA- 80
WP_103426078_1 hypot --------------MATVSDRVGLDPERLWGALVAVLAAALVGGSVAFPRVVYD---RFVWQYFWGPVEA-DAQSAVCAVRTAAG-TEYLGSASACEAA- 80
WP_009367433_1 DUF63 --------------MATVSERLGVDSERLWGGTILALLVALVGGSLLLPRIVYD---NFIWHYFWGPVEA-DANSAVCAVRQGGV-TEYLSSTSACAAA- 80
WP_013440552_1 DUF63 --------------MATVAERVGMDPERLWGSAMVAVLVALIGGSLAFPELVYD---RFIWHYFWGPVQA-DANSAVCAVRTGGV-TQYLTSASACAEA- 80
WP_049916626_1 DUF63 --------------MATVAERVGADPERLWGGTALVALVALVGGSIAFPELVYD---RFIWHYFWGPVQA-DANSAVCAVRPGGV-TEYLSSASACAAA- 80
WP_058582837_1 DUF63 --------------MATVAERVEADPERTWLGTMAVVLTALVGGSLLFPRIVYD---GFLWHYFWGPVVA-DGEGAVCAVRSGGT-TQLLDSAAACDAA- 80
WP_101298124_1 hypot --------------MATVAERVGVAPERLWGGTVLALLLALVGGSIAFPRAVYD---GFLWRYFWGPVVA-DGNGAVCAVRSGGT-TRLLNSQSACAAA- 80
PIN95153_1 hypotheti --------------------------------------------------------------MALTPIET------------------------------ 8
PIU22108_1 hypotheti ----------------------------------------------------MG----FFYDSFGKTITE------------------------------ 14
AAR39201_1 NEQ352    ---------------------------------------------------------------------------------------------------- 0
OIR14399_1 hypotheti -------------------------------MDSQSKTTTSAIGALIIISYFWEPLRDFWYQNFFAPLLA-DARGVPVDGITAT---------------- 52
OIR20963_1 hypotheti -------------------------MEQVQRIQYVIGILVLTILAISFVPQVNS----FWEEYFVSPIQS------------------------------ 41
OIR22371_1 hypotheti -------MEDKPQAAVTRHEGFQIAKISENKYISGAVIAIVLILSIQFIPQVND----FWNEFFVNPIMA------------------------------ 59
EGQ43935_1 putative  -----------------------------------------------MLDVVKA----FLFEKFWLPIID------------------------------ 19
MAG21679_1 hypotheti -------------------------------------------------MDIGE----LLYDIFGKRIAE------------------------------ 17
PIN85618_1 hypotheti --------------------------------------------------MVQD----FLFENFCRPVLD------------------------------ 16
PIN99249_1 hypotheti --------------------------------------------------MVLD----LINEFFVNPITS------------------------------ 16
AJF59838_1 hypotheti ------------------------------------------------MVGFPP----FIQEFFLDPICN------------------------------ 18
WP_042682145_1 DUF63 -----------------------------------------------MLESIRE----FLWIYFIRPMYT------------------------------ 19
WP_013468033_1 DUF63 -----------------------------------------------MLESIKE----FFWIYFIRPMYT------------------------------ 19
WP_055281581_1 DUF63 ------------------------------------------------MESVER----FIWEYFVRPMYT------------------------------ 18
WP_048160382_1 DUF63 ------------------------------------------------MESVER----FLWEHFIRPMYT------------------------------ 18
WP_042701216_1 DUF63 ------------------------------------------------MESVER----FIWEYFIRPMYT------------------------------ 18
WP_004068659_1 DUF63 ------------------------------------------------MESVEK----FIWEYFIRPMYT------------------------------ 18
```

```
WP_058946665_1 DUF63 ------------------------------------------------MESLEG----FIWEYFIRPMYT------------------------------ 18
WP_014835499_1 DUF63 ------------------------------------------------MVDVKE----FLWEYFIRPMYT------------------------------ 18
WP_010884243_1 DUF63 ---------------------------------------------MGIKVNLHQ----FFWEYFIRPMYT------------------------------ 21
WP_013748129_1 DUF63 -------------------------------------------------MNLHE----FLWEYFIRPMYT------------------------------ 17
WP_010867251_1 DUF63 -------------------------------------------------MDVHQ----FLWEYFIRPMYT------------------------------ 17
WP_014733359_1 DUF63 ------------------------------------------------MVDIHQ----FLWEYFIRPMYT------------------------------ 18
WP_068319891_1 DUF63 ------------------------------------------------MFNLRD----FLWNYFIRPMYT------------------------------ 18
WP_068576049_1 DUF63 ------------------------------------------------MFSLHD----FLWNYFIRPMYT------------------------------ 18
WP_014013692_1 DUF63 -----------------------------------------------MLESLWNELWGFMYKYFWEPMFT------------------------------ 23
WP_014789520_1 DUF63 -------------------------------------------MLESLQHEIWN----FLYKYFWEPMFT------------------------------ 23
WP_088180676_1 DUF63 -----------------------------------------------MLESLWNEVWGFIYKYFWEPMFT------------------------------ 23
WP_088864885_1 DUF63 -----------------------------------------------MLESLWNELWGFIYRYFWEPMFT------------------------------ 23
WP_088856755_1 DUF63 -------------------------------------------MLESLWYRIWE----FFYTYFWEPMFT------------------------------ 23
WP_088865981_1 DUF63 -----------------------------------------------MLESLWNEVWGFIYKYFWEPMFT------------------------------ 23
WP_013906334_1 DUF63 -----------------------------------------------MLDVLRD----FLWEYFVRPMYT------------------------------ 19
WP_088862658_1 DUF63 --------------------------------------------MTGIWQAIWE----FFYTYFWHPMFT------------------------------ 22
WP_088882956_1 DUF63 --------------------------------------------MTGIWQSLWE----FFNAYFWEPMFT------------------------------ 22
WP_012571938_1 DUF63 -----------------------------------------------MLESLWN----FIYQYFIEPMYT------------------------------ 19
WP_068663933_1 DUF63 -----------------------------------------------MLEFLWS----FINQYFIEPMYT------------------------------ 19
WP_074631153_1 DUF63 -----------------------------------------------MLESLMD----FLYRYFWEPMFT------------------------------ 19
WP_058939074_1 DUF63 -----------------------------------------------MLESLWNELWGFIYKYFWEPMFT------------------------------ 23
WP_088854681_1 DUF63 -----------------------------------------------MLESVWHEIWWFFHTYFWEPMFT------------------------------ 23
WP_088885426_1 DUF63 -----------------------------------------------MLESLWQ----FLNQYFIEPMYT------------------------------ 19
WP_011249694_1 DUF63 ---------------------------------------------MSVEGAVID----FFYRYFWEPMFT------------------------------ 21
WP_062386972_1 DUF63 ---------------------------------------------MSVEGAVID----FFYRYFWEPMFT------------------------------ 21
WP_010478887_1 DUF63 -----------------------------------------------MLESVWQ----FLNQYFIEPMYT------------------------------ 19
WP_088858605_1 DUF63 -----------------------------------------------MLESVWH----FLNEYFIEPMYT------------------------------ 19
WP_042690028_1 DUF63 ---------------------------------------------MGIEKALWD----FFYNYFWEPMFT------------------------------ 21
WP_048150762_1 DUF63 -----------------------------------------------MLEAIWN----FFNQYFWEPMFT------------------------------ 19
WP_048811076_1 DUF63 -----------------------------------------------MLEAVWK----FFNQYFWEPMFT------------------------------ 19
WP_050003256_1 DUF63 ---------------------------------------------MGAGEALWD----FFYRYFWEPMFT------------------------------ 21
WP_062372100_1 DUF63 -----------------------------------------------MLEAIWN----FFNQYFWEPMFT------------------------------ 19
WP_048165284_1 DUF63 ---------------------------------------------MSISQEIYN----FLWNYFIRPMYT------------------------------ 21
WP_048148750_1 DUF63 ---------------------------------------------MALSQSIYD----FLWNYFIRPMYT------------------------------ 21
OYT53462_1 hypotheti -------------------------------------------------MGV------------------------------------------------ 3
KYC51152_1 hypotheti -------------------------------------------------MGV------------------------------------------------ 3
KYC46003_1 hypotheti -------------------------------------------------MGV------------------------------------------------ 3
KYC48643_1 hypotheti -------------------------------------------------MGV------------------------------------------------ 3
KYC55321_1 hypotheti -------------------------------------------------MGV------------------------------------------------ 3
KYC57927_1 hypotheti -------------------------------------------------MGV------------------------------------------------ 3
KYC57171_1 hypotheti -------------------------------------------------MGV------------------------------------------------ 3
OIO20701_1 hypotheti ---------------------------------------------------MAD----FFTDYFLEPINS------------------------------ 15
PIT83986_1 hypotheti ----------------------------------------------MKCMDLPV----LFEEFFIAPIMQ------------------------------ 20
OIO24760_1 hypotheti ------------------------------------------------MGFIED----FVSEYFVKPLAD------------------------------ 18
OIO24701_1 hypotheti -------------------------------------------------MDFSS----FLDSYFLEPLRR------------------------------ 17
OIO27021_1 hypotheti --------------------------------------------------------MGFVEEHFLNPMRY------------------------------ 14
OIO26762_1 hypotheti ---------------------------------------------------MAD----WIDEFFVAPMRE------------------------------ 15
PIN95811_1 hypotheti ---------------------------------------------------MAD----WIDEFFVAPMRE------------------------------ 15
PIO01637_1 hypotheti ---------------------------------------------------MAD----WIDEFFVAPMRE------------------------------ 15
PIO02820_1 hypotheti --------------------------------------------------------MGFVEEHFLNPMRY------------------------------ 14
PJD01038_1 hypotheti ----------------------------------------------------MG----FVEEHFLNPMRY------------------------------ 14
PIZ91366_1 hypotheti ---------------------------------------------------MAD----WIDEFFVAPMRE------------------------------ 15
WP_013100570_1 DUF63 -----------------------------------------------MLKELKE----FIYKYYIEPAET------------------------------ 19
WP_004590770_1 DUF63 -----------------------------------------------MLDKVYE----FIYKYYIEPAEI------------------------------ 19
WP_048196979_1 DUF63 -----------------------------------------------MKDVIYE----FIYKYYIEPAKL------------------------------ 19
WP_015791538_1 DUF63 -----------------------------------------------MIEEIKT----FIYKYYIEPAEK------------------------------ 19
WP_048202292_1 DUF63 -----------------------------------------------MIEKIRD----FIYKYYIEPAEK------------------------------ 19
WP_012981280_1 DUF63 -----------------------------------------------MIQEIKE----FIYKYYIEPAEK------------------------------ 19
WP_064496496_1 DUF63 -----------------------------------------------MIQEIKN----FIYKYYIEPAEK------------------------------ 19
WP_011972741_1 DUF63 ---------------------------------------------MDIAAYISG----FINKYYINPIYN------------------------------ 21
WP_013798289_1 DUF63 -----------------------------------------------MLSAIKE----FIYEYYIKPMIE------------------------------ 19
WP_013181100_1 DUF63 ---------------------------------------------MDIIQIIDN----FIYEHYIYPIEA------------------------------ 21
WP_013867628_1 DUF63 ---------------------------------------------MDIKETFSE----FVYKYYIEPIKL------------------------------ 21
WP_018153391_1 DUF63 ---------------------------------------------MDIKNTVLE----FINKYYIYPIHA------------------------------ 21
WP_011170272_1 DUF63 ---------------------------------------------MNGMLLIRE----FIYRYYIYPIDT------------------------------ 21
WP_012066151_1 DUF63 ---------------------------------------------MDIMLMIKD----FIYRYYIYPIEA------------------------------ 21
ODS43013_1 hypotheti -----------------------------------------------MIETVWN----FIHKYYIVP--------------------------------- 16
OIQ06085_1 hypotheti -----------------------------------------------MSDIIND-----KINFIINEFTS------------------------------ 18
PIN67069_1 hypotheti -----------------------------------------------MSDIIND-----KINFIINEFTS------------------------------ 18
PIV28099_1 hypotheti -----------------------------------------------MSDIIND-----KINFIINEFTS------------------------------ 18
PJC13070_1 hypotheti -----------------------------------------------MSDIIND-----KINFIINEFTS------------------------------ 18
PIZ29927_1 hypotheti -----------------------------------------------MSDIIND-----KINFIINEFTS------------------------------ 18
PKP60688_1 hypotheti -----------------------------------------------MSDILND-----GINFIINEFTR------------------------------ 18
WP_012956954_1 DUF63 ------------------------------------------------MATVDS----FLPDFIQTTFFS------------------------------ 18
WP_067148430_1 DUF63 ------------------------------------------------MATVDT----FIPDIIQTTFFS------------------------------ 18
WP_080460538_1 DUF63 -------------------------------------------------MSIESSFINFLSIYFFS---------------------------------- 17
WP_010877090_1 DUF63 --------------------------------------------------MISE-----IIEFLESNFFY------------------------------ 15
WP_013294901_1 DUF63 --------------------------------------------------MISE-----LQKFLESNFFY------------------------------ 15
WP_010877090_1 DUF63 --------------------------------------------------MISE-----IIEFLESNFFY------------------------------ 15
BAZ99473_1 hypotheti --------------------------------------------------MISE-----IIEFLESNFFY------------------------------ 15
PKL66404_1 hypotheti ---------------------------------------MQAEIGISFSNSVSQ----FIQDNFLY---------------------------------- 23
WP_023991171_1 DUF63 -------------------------------------------------MGIDS-----IIQYIQENFVY------------------------------ 16
WP_048072898_1 DUF63 -------------------------------------------------MGIDS-----IIQYIQENFVY------------------------------ 16
WP_100905180_1 hypot --------------------------------------------------MVLD----QITQYIQENFIY------------------------------ 16
WP_100907231_1 hypot --------------------------------------------------MVLD----QITQYIQENFIY------------------------------ 16
WP_100907231_1 hypot --------------------------------------------------MVLD----QITQYIQENFIY------------------------------ 16
WP_048081925_1 DUF63 -----------------------------------------------MLSDISE----FIQQNFLYT--------------------------------- 16
WP_048081925_1 DUF63 -----------------------------------------------MLSDISE----FIQQNFLYT--------------------------------- 16
WP_069585525_1 DUF63 -----------------------------------------------MLSDISE----FIQQNFLYT--------------------------------- 16
WP_069585525_1 DUF63 -----------------------------------------------MLSDISE----FIQQNFLYT--------------------------------- 16
WP_013643671_1 DUF63 --------------------------------------------------MIPQ-----IMQYIRENFIY------------------------------ 15
WP_013824571_1 DUF63 -----------------------------------------------MFDQISQ----FVRQNFIY---------------------------------- 15
WP_048192110_1 DUF63 -------------------------------------------MIFDFFQGILE----YIRQNFVY---------------------------------- 19
WP_081810122_1 DUF63 -------------------------MTLPSKIRLSSSMGSAEDERFRMMQNLID----FINKNYIEGIIN------------------------------ 41
WP_097298965_1 DUF63 -------------------------------------------------MSLID----FINRNFIEGIVN------------------------------ 17
```

```
PIO000292_1 hypotheti --------------------------------------------------MVFDSIVQFVYHYFWRPYEQ------------------------------ 20
WP_011833679_1 DUF63 ---------------------------------------------------------MFNELYAWIVELE------------------------------ 13
WP_042698332_1 DUF63 --------------------------------------------------MLNE-----LYKWIVELEGS------------------------------ 15
WP_042698332_1 DUF63 --------------------------------------------------MLNE-----LYKWIVELEGS------------------------------ 15
WP_011448164_1 DUF63 --------------------------------------------------MIRE----FIYKYYIDPIRY------------------------------ 16
WP_007314214_1 DUF63 ----------------------------------------------MYPEVIGD----FIEKYYIDPIRY------------------------------ 20
WP_042705632_1 DUF63 --------------------------------------------------MLRD----FLYKYYVDPILN------------------------------ 16
WP_013329717_1 DUF63 --------------------------------------------------MIGD----FIYKYYVGPIVN------------------------------ 16
WP_004077722_1 DUF63 --------------------------------------------------MIGD----FIYKYYVGPIVN------------------------------ 16
WP_048150927_1 DUF63 --------------------------------------------------MIGD----FIYKYYVGPIVN------------------------------ 16
WP_012107531_1 DUF63 --------------------------------------------------MISD----FLYKYYIDPVKY------------------------------ 16
PKL70081_1 hypotheti --------------------------------------------------MISD----FFYKYYIDPVRY------------------------------ 16
WP_015285007_1 DUF63 --------------------------------------------------MISD----FIYKYYIDPIRL------------------------------ 16
PKL64570_1 hypotheti --------------------------------------------------MVSD----FIYKYYIGPITQ------------------------------ 16
WP_015286404_1 DUF63 --------------------------------------------------MIAE----FITKYYIDPIRY------------------------------ 16
WP_012617618_1 DUF63 --------------------------------------------------MIRE----FIYKYYIDPIRY------------------------------ 16
WP_014867374_1 DUF63 --------------------------------------------------MIRE----FLYKYYIDPIRY------------------------------ 16
CVK33108_1 conserved --------------------------------------------------MIRE----FLYKYYIDPIRY------------------------------ 16
WP_066956331_1 DUF63 --------------------------------------------------MIRE----FIYKYYIDPIRY------------------------------ 16
WP_011844879_1 DUF63 --------------------------------------------------MIRE----FLYKYYIDPIRY------------------------------ 16
WP_048181868_1 DUF63 --------------------------------------------------MIRE----FLYKYYIDPIRY------------------------------ 16
WP_067073014_1 DUF63 --------------------------------------------------MIRE----FLYKYYIDPIRY------------------------------ 16
PKL62929_1 hypotheti --------------------------------------------------MIRE----FLYKYYIDPIRY------------------------------ 16
WP_004039890_1 DUF63 --------------------------------------------------MIRE----FLYKYYIDPIMY------------------------------ 16
WP_067049225_1 DUF63 --------------------------------------------------MIRE----FLYKYYIDPIRY------------------------------ 16
KYK37884_1 hypotheti --------------------------------------------------MIPE----FIEEYFVDPIKY------------------------------ 16
KYK28019_1 hypotheti --------------------------------------------------MIPE----FIQEYFVDPIKY------------------------------ 16
OYT57835_1 hypotheti -----------------------------------------------MLGQIEN----ILQHYFIEPVET------------------------------ 19
OYT33350_1 hypotheti ----------------------------------------------MDSEIIWN----FIKKYYINSIIY------------------------------ 20
WP_012964745_1 DUF63 -------------------------------------------------MSLWE----FIKKYYIDSIVY------------------------------ 17
WP_048091573_1 DUF63 -------------------------------------------------MGLYE----FIKKYYIDSIVY------------------------------ 17
WP_048096399_1 DUF63 -------------------------------------------------MDVYG----FIKEYFIDSIVY------------------------------ 17
WP_012940099_1 DUF63 --------------------------------------------------MIWD----FIKKYYIDSIVY------------------------------ 16
WP_010877969_1 DUF63 -------------------------------------MVKLFNQLLIFPMDLWE----FIKEYYIDSIVY------------------------------ 29
WP_013682839_1 DUF63 --------------------------------------------------MLWD----FIKKYYIDSIVY------------------------------ 16
WP_015591141_1 DUF63 -------------------------------------------------MGIYE----FIKKYYIDSLVY------------------------------ 17
WP_012035887_1 DUF63 --------------------------------------------MNMLSFDIGE----FVYKYYIHPIVY------------------------------ 22
WP_014404626_1 DUF63 -------------------------------------------------MDIGS----FVGRYYIDPIVY------------------------------ 17
BAI60262_1 conserved -------------------------------------------------MDIGS----FIAKYYIDPIIY------------------------------ 17
WP_042684156_1 DUF63 ---------------------------------------------MWWCGDVVSTLSEFVHRYYIDPIVY------------------------------ 25
OFV68024_1 membrane  ---------------------------------------------------------------------------------------------------- 0
WP_013720253_1 DUF63 ------------------------------------------------MIDLAD----WIYRYYIHPIIY------------------------------ 18
WP_014587756_1 DUF63 ---------------------------------------------------MHP----WIYHHYVEPILY------------------------------ 15
ABK14947_1 Protein o --------------------------------------------MNSLSSQIVH----FVQRYYVEPIIN------------------------------ 22
OKY79140_1 putative  -----------------------------------------------MLDKLIE----FINKYYINPIVQ------------------------------ 19
WP_086637003_1 DUF63 -----------------------------------------------MIDRILE----FIDNYYIQPIVQ------------------------------ 19
WP_048089097_1 DUF63 ----------------------------------------------MITDTVWQ----YIYKYYISGIVN------------------------------ 20
WP_097298272_1 DUF63 ---------------------------------------------MSIIDSVWH----YIDKYYLSGIIN------------------------------ 21
WP_013897788_1 DUF63 --------------------------------------------MNSLIDKIIQ----FLNTYYIDPIRY------------------------------ 22
WP_015323982_1 DUF63 --------------------------------------------MNSYIDKVSG----FINTYYIDPILH------------------------------ 22
WP_011499115_1 DUF63 --------------------------------------------MCPYVDKILE----FVNRYYLEPIFQ------------------------------ 22
WP_013037846_1 DUF63 --------------------------------------------MNTFTDKILQ----FVNKYYIEPIIY------------------------------ 22
WP_048205640_1 DUF63 --------------------------------------------MCPFVDKIMQ----FVNEYYLDPIFQ------------------------------ 22
WP_072561629_1 DUF63 --------------------------------------------MNTFTDKILQ----FVNKYYIEPIIY------------------------------ 22
WP_072360082_1 DUF63 --------------------------------------------MNTFTDKILQ----FVNKYYIDPIIY------------------------------ 22
WP_096711793_1 DUF63 --------------------------------------------MNTLTDKILQ----FVNKYYIDPIIY------------------------------ 22
ODV50598_1 hypotheti --------------------------------------------MNTLTDKILQ----FVNKYYIDPIIY------------------------------ 22
WP_013194042_1 DUF63 --------------------------------------------MNSFIDNIWE----FIQNYYIDPIIF------------------------------ 22
WP_048178067_1 DUF63 --------------------------------------------MSSPLDAISQ----FINTYYLDPILT------------------------------ 22
WP_048127111_1 DUF63 --------------------------------------------MSFLTDNISQ----FINTYYLDPIRG------------------------------ 22
WP_011023594_1 DUF63 ----------------------------------------MSDKMNILTDKISQ----FINTYYLDPIKG------------------------------ 26
WP_048184587_1 DUF63 --------------------------------------------MSIFIDKISQ----FINTYYLDPIKT------------------------------ 22
WP_011032542_1 DUF63 --------------------------------------------MSFLMEKISQ----FINTYYLDPIKG------------------------------ 22
WP_011032542_1 DUF63 --------------------------------------------MSFLMEKISQ----FINTYYLDPIKG------------------------------ 22
WP_048129141_1 DUF63 --------------------------------------------MSILADKISQ----FINTYYLDPIRG------------------------------ 22
WP_048129141_1 DUF63 --------------------------------------------MSILADKISQ----FINTYYLDPIRG------------------------------ 22
WP_048169891_1 DUF63 --------------------------------------------MSYLTDNISQ----FINTYYLDPIRT------------------------------ 22
WP_048137175_1 DUF63 --------------------------------------------MSYLTDNISQ----FINTYYLDPIRT------------------------------ 22
WP_048137175_1 DUF63 --------------------------------------------MSYLTDNISQ----FINTYYLDPIRT------------------------------ 22
WP_048137175_1 DUF63 --------------------------------------------MSYLTDNISQ----FINTYYLDPIRT------------------------------ 22
WP_048137694_1 DUF63 --------------------------------------------MSYLIEKISQ----FINTYYLDPIRG------------------------------ 22
WP_048168126_1 DUF63 --------------------------------------------MSFSIDTISQ----FINTYYLDPIRG------------------------------ 22
WP_048118082_1 DUF63 --------------------------------------------MSFSIDNISQ----FINTYYLDPIRG------------------------------ 22
WP_048158120_1 DUF63 --------------------------------------------MSFSIDNISQ----FINTYYLDPIRG------------------------------ 22
WP_011305329_1 DUF63 --------------------------------------------MSFSIDNISQ----FINTYYLDPIRG------------------------------ 22
WP_054298619_1 DUF63 --------------------------------------------MSFSIDTISQ----FINTYYLDPIRG------------------------------ 22
ALK05385_1 hypotheti --------------------------------------------MSFSIDTISQ----FINTYYLDPIRG------------------------------ 22
WP_015052897_1 DUF63 --------------------------------------------MVPIIDKISQ----LINEYYIDPILH------------------------------ 22
WP_023846134_1 DUF63 ---------------------------------------------MPFIDKVTQ----FINEYYIDPILH------------------------------ 21
OIN88451_1 hypotheti --------------------------------------------------MPLD----FFYRYFVEPIEL------------------------------ 16
PIX50278_1 hypotheti --------------------------------------------------MPLD----FFYRYFVEPIEL------------------------------ 16
PIW41402_1 hypotheti --------------------------------------------------MPLD----FFYRYFVEPIEL------------------------------ 16
PIY35178_1 hypotheti --------------------------------------------------MPLD----FFYRYFVEPIEL------------------------------ 16
PJB74886_1 hypotheti --------------------------------------------------MPLD----FFYRYFVEPIEL------------------------------ 16
PIZ33651_1 hypotheti --------------------------------------------------MPLD----FFYRYFVEPIEL------------------------------ 16
WP_048165029_1 DUF63 -------------------------------------------------MGVGE----IFREYFVNPILY------------------------------ 17
WP_042681046_1 DUF63 -------------------------------------------------MGVSE----IFQEYFVNPIKY------------------------------ 17
WP_013467319_1 DUF63 -------------------------------------------------MGVSE----IFQKYFVNPIKY------------------------------ 17
WP_048152160_1 DUF63 -------------------------------------------------MGAWE----VFQRYFVDPIKY------------------------------ 17
WP_015849008_1 DUF63 -------------------------------------------------MGIYE----ILQKYFIDPIRY------------------------------ 17
WP_004069276_1 DUF63 -------------------------------------------------MGIGE----IFQRYFIDPIRY------------------------------ 17
WP_058946638_1 DUF63 -------------------------------------------------MGIGE----VFQRYFIDPIRY------------------------------ 17
WP_042701551_1 DUF63 -------------------------------------------------MGIGK----IFQRYFIDPIRY------------------------------ 17
WP_055282692_1 DUF63 -------------------------------------------------MGIAE----VFQRYFIDPIKY------------------------------ 17
WP_013906023_1 DUF63 --------------------------------------------------MLEE----FFQRYFIDPIRY------------------------------ 16
```

```
WP_011013158_1 DUF63 --------------------------------------------------MIRE----FFQKYFIDPIKY------------------------------ 16
WP_014733166_1 DUF63 --------------------------------------------------MLKE----FFEKYFINPIKY------------------------------ 16
WP_068322889_1 DUF63 --------------------------------------------------MLKE----FFEKYFVNPIKY------------------------------ 16
WP_068575625_1 DUF63 --------------------------------------------------MLKE----FFEKYFVNPIKY------------------------------ 16
WP_010885960_1 DUF63 --------------------------------------------------MLRD----FFEKYFVNPIKY------------------------------ 16
WP_010867401_1 DUF63 --------------------------------------------------MLGD----FFRKYFVNPIKY------------------------------ 16
WP_013747987_1 DUF63 -------------------------------------------------MGVVD----FFRRYFVDPIKY------------------------------ 17
WP_068664049_1 DUF63 -------------------------------------------------MGLWE----FFYRYFIEPIVN------------------------------ 17
WP_011249739_1 DUF63 -------------------------------------------------MGVGE----FFYRYFVEPIKY------------------------------ 17
WP_062386854_1 DUF63 -------------------------------------------------MGVEE----FFYRYFVEPIKY------------------------------ 17
WP_050003781_1 DUF63 -------------------------------------------------MGLEE----FFQRYFIDPIKY------------------------------ 17
WP_042690699_1 DUF63 -------------------------------------------------MGLYE----FFYRYFVEPIKY------------------------------ 17
WP_088885212_1 DUF63 -------------------------------------------------MGFQE----FFQRYFVDPIKY------------------------------ 17
WP_010478519_1 DUF63 -------------------------------------------------MGLQE----FFQRYFVDPIKY------------------------------ 17
WP_088858240_1 DUF63 -------------------------------------------------MGAGE----FFQRYFVDPIKY------------------------------ 17
WP_015859016_1 DUF63 -------------------------------------------------MGLQE----FFQRYFVDPIKY------------------------------ 17
WP_014121940_1 DUF63 -------------------------------------------------MGIQE----FFQRYFVDPIKY------------------------------ 17
WP_062373911_1 DUF63 -------------------------------------------------MGLQE----FFQKYFIDPIKY------------------------------ 17
WP_088882924_1 DUF63 -------------------------------------------------MGLHE----FFQDYFVRPITE------------------------------ 17
WP_088862911_1 DUF63 -------------------------------------------------MGLQG----FFHRYFIEPIQQ------------------------------ 17
WP_012572006_1 DUF63 -------------------------------------------------MGLYE----FFYRYFIEPIIN------------------------------ 17
WP_088854707_1 DUF63 -------------------------------------------------MGLYE----FFYKYFVLPIKE------------------------------ 17
WP_014789474_1 DUF63 -------------------------------------------------MGLYE----FFYEYFVLPIKE------------------------------ 17
WP_088180728_1 DUF63 -------------------------------------------------MGLYE----FFYEYFIKPIQE------------------------------ 17
WP_088864937_1 DUF63 -------------------------------------------------MGLYE----FFYEYFIRPIQE------------------------------ 17
WP_055429686_1 DUF63 -------------------------------------------------MGLYE----FFYEYFIRPIQE------------------------------ 17
WP_088865932_1 DUF63 -------------------------------------------------MGLYE----FFYEYFIKPIQE------------------------------ 17
WP_014013642_1 DUF63 -------------------------------------------------MGLYE----FFYEYFIRPIQE------------------------------ 17
WP_088856804_1 DUF63 -------------------------------------------------MGLYQ----FFYEYFVVPIKE------------------------------ 17
WP_058939475_1 DUF63 -------------------------------------------------MGLYE----FFYEYFIRPIQE------------------------------ 17
EHR77276_1 conserved -MSTNALEEHLENAEPALPDAEYSVYEKIALWAVLVLIAAVVSGLVLANETVWD---EGLKPIVWDPVTK------------------------------ 66
OUV40025_1 hypotheti MATLESIDEVLGTHQPALPSTRLSMVEQTLTRLLLFLIIGVAIGLLLMPEAIWD---DGLRPIIWEPIQQ------------------------------ 67
MBJ52984_1 hypotheti ------MKEKYFTKSFILKIKKFNTFELIAICFVLINIIILGFGSLIFTESIWN---QYVYPNIWEPIVN------------------------------ 61
PDH23744_1 hypotheti -----MAIRRPVNNPLDEAHPYETWAINALLAVFL-LIFLGLGLEWAGIENFLS---ESLNEYFIDPIKG-ES----T---------------------- 64
PDH25468_1 hypotheti -----MLARTPMMNHLEEWYDYEQTVIKALSAIMTLFFAGFALHELDIENPLSD----FVYQYYLDPIIG------------------------------ 61

WP_048201856_1 hypot ---------------------YYFIQTSPYY--------NGYTKKTLIIWLG---------FAPFLTSFLILLVFGENTVKIPLD--------------- 73
WP_048150344_1 hypot ------------A----G---FFLLSLS----LSLLLTLLIIMS---TTRLD------LTTVLVFNLLSIPFIAIG-GLLVEKGD-----ACPFV----- 64
AOV95360_1 hypotheti ---------------------MTATNIALDS-IILAYTILLTSYTIKKRRKK----QVHSINRYFTAVLIPYIILI-----SLTS-----A---N----- 56
EOD42420_1 Uncharact -----------------Q---YNIYNTFLYG-LLFVVGFILFYY-LFVKNKK------VIIDRYFLLDLIILTTIF-IIPRVNID-----L---G----- 78
AOV95164_1 hypotheti ----------AHT----G---YNLFNTVAWA-LIAVTAILLIRK-E-FNKRE------IQLKTQTAISAIPFILLG-GILRFLED-----A---A----- 89
EGQ40074_1 putative  ----------AYT----G---YNPVNTGVWV-GIALVCLGGAKR-L-ANEYN------ISYEPSTVVYASPFVMMG-GVLRFIED-----A---A----- 93
KYK23067_1 hypotheti GHTVSHNGVIAQE----G---YTLVSEITYG-IILICALYGLYK-L-LKKLE------IRIDWYFCLALFPYILFG-PVTRVLED-----T---N----- 125
WP_084383883_1 DUF63 GQPVTHEGIRAVR----G---YNAVNTMTYL-AVVVYSLPGLRA-Y-LDALD------VSFDARLAYGFAPIIVAG-GAMRALED-----I---G----- 131
WP_049984677_1 DUF63 GRPVVHDGVRAVT----G---YNIVNTATYL-AAVGYSFPGLRA-Y-LRYLN------VTFDARLAYGFAPIIIAG-GAMRALED-----I---G----- 132
WP_103428047_1 hypot SQPVTHEGIRAVS----G---YNAVNTVTYL-AAVGYSLPGLRA-Y-LDQLD------VTFDARLAYGFAPIIVAG-GAMRALED-----I---G----- 131
WP_004048647_1 DUF63 SEPVMHEGIRAVR----G---YNAVNTVTYL-AAVVYSLPGLRV-Y-LDQLD------VTFDARLAYGFAPIIIAG-GAMRALED-----I---G----- 131
WP_004594600_1 DUF63 GQPVTHEGVRAVQ----G---YNAVNTVIYL-AAVVYSLPGLRA-Y-LDQLD------VTFDARLAYGFAPIIIAG-GAMRALED-----L---G----- 131
WP_004594600_1 DUF63 GQPVTHEGVRAVQ----G---YNAVNTVIYL-AAVVYSLPGLRA-Y-LDQLD------VTFDARLAYGFAPIIIAG-GAMRALED-----L---G----- 131
WP_050050397_1 DUF63 GQPVTHEGIRAVQ----G---YNAVNTVIYL-AAVVYSLPGLRA-Y-LDQLD------VTFDARLAYGFAPIIIAG-GAMRALED-----L---G----- 131
WP_004594600_1 DUF63 GQPVTHEGVRAVQ----G---YNAVNTVIYL-AAVVYSLPGLRA-Y-LDQLD------VTFDARLAYGFAPIIIAG-GAMRALED-----L---G----- 131
WP_050050397_1 DUF63 GQPVTHEGIRAVQ----G---YNAVNTVIYL-AAVVYSLPGLRA-Y-LDQLD------VTFDARLAYGFAPIIIAG-GAMRALED-----L---G----- 131
WP_049983690_1 DUF63 GQPVTHEGIRAVR----G---YNAVNTVTYL-AAVVYSLPGLRA-Y-LDSLD------VTYDARLAYGFAPIIVAG-GAMRALED-----I---G----- 131
WP_103428162_1 hypot GQPVTHEGIRAVS----G---YNAVNTVTYL-AAVVYSLPGLRA-Y-LDALN------VSFDTRLAYGFAPIIVAG-GAMRALED-----I---G----- 131
WP_009378268_1 DUF63 GQPVTHEGIRAVS----G---YNAVNTVTYL-AAVVYSLPGLRA-Y-LDALN------VSFDTRLAYGFAPIIVAG-GAMRALED-----I---G----- 131
WP_015763176_1 DUF63 --RATGTAIYAEP----G---YTIVSEVGYM-LVLVYMLVGVYL-F-LQRYE------LVTDIDLYYPLVPFMLLG-GSLRVVED-----A---T----- 152
WP_018259238_1 DUF63 --QASGTAVYAEP----G---YTIISEIGYM-LVLVYMLVGVYL-F-LQRYD------LVTDIDLYYPLVPFMLLG-GSLRVVED-----A---T----- 152
WP_004592615_1 DUF63 --IQEGRLI-AEP----G---YTIVSEIGYM-LILVYMLVGVYF-L-LERLD------IAEDPKLYFAFVPFMLLG-GALRTVED-----A---T----- 146
WP_004518092_1 DUF63 --IQEGRLI-AEP----G---YTIVSEVGYM-LILVYMLVGVYF-L-LERLD------IAEDPELYFAFVPFMLLG-GALRTVED-----A---T----- 146
WP_014040450_1 DUF63 --IQEGRLI-AEP----G---YTIVSEIGYM-LILVYMLVGVYF-L-LERLD------IAEDPKLYFAFVPFMLLG-GALRTVED-----A---T----- 146
WP_014040450_1 DUF63 --IQEGRLI-AEP----G---YTIVSEIGYM-LILVYMLVGVYF-L-LERLD------IAEDPKLYFAFVPFMLLG-GALRTVED-----A---T----- 146
WP_008309018_1 DUF63 --IQEGRLI-AEP----G---YTIVSEIGYM-LILVYMLVGVYF-L-LERLD------IAEDPKLYFAFVPFMLLG-GALRTVED-----A---T----- 146
WP_014040450_1 DUF63 --IQEGRLI-AEP----G---YTIVSEIGYM-LILVYMLVGVYF-L-LERLD------IAEDPKLYFAFVPFMLLG-GALRTVED-----A---T----- 146
WP_014040450_1 DUF63 --IQEGRLI-AEP----G---YTIVSEIGYM-LILVYMLVGVYF-L-LERLD------IAEDPKLYFAFVPFMLLG-GALRTVED-----A---T----- 146
WP_005534235_1 DUF63 --IQEGRLI-AEP----G---YTIVSEIGYM-LILVYMLVGVYF-L-LERLD------IAEDPKLYFAFVPFMLLG-GALRTVED-----A---T----- 146
WP_004961153_1 DUF63 --IQEGRLI-AEP----G---YTIVSEIGYM-LILVYMLVGVYF-L-LERLD------IAEDPKLYFAFVPFMLLG-GALRTVED-----A---T----- 146
WP_004961153_1 DUF63 --IQEGRLI-AEP----G---YTIVSEIGYM-LILVYMLVGVYF-L-LERLD------IAEDPKLYFAFVPFMLLG-GALRTVED-----A---T----- 146
WP_101350151_1 hypot --IQDGRLI-AEP----G---YTIVSEVGYM-LILVYMLVGVYF-L-LERLD------IAEDPKLYFAFVPFMLLG-GALRTVED-----A---T----- 146
WP_053968273_1 DUF63 --IQEGRLI-AEP----G---YTIVSEVGYM-LILVYMLVGVYF-L-LERLD------IAEDPKLYFAFVPFMLLG-GALRTVED-----A---T----- 146
WP_058995687_1 DUF63 --IQEGRLI-AEP----G---YTIVSEIGYM-LILVYMLVGVYF-L-LERLD------IAEDPKLYFAFVPFMLLG-GALRTVED-----A---T----- 146
WP_015790673_1 DUF63 GIAVEQGHIVAEP----G---YTLVSEAGYM-VLGLFFLIGVYL-L-LKNLD------IDLDREFFYALFPFMLFG-GVVRVVED-----ASDAA----- 150
WP_008524115_1 DUF63 GIAAQQGHIVAEP----G---YTLVSEAGYM-IVGLFFLIGVYL-L-LSNLD------IDLDKAFFYALFPFMLFG-GALRVLED-----ATDAA----- 150
WP_075936143_1 DUF63 -----SGIV-AEP----G---YTLVSEAGYA-ISLVFFLVGVLT-L-LHRLE------IGDTKQLFLALVPFMFFG-GALRVIED-----A---N----- 141
WP_020446311_1 DUF63 -----SGIV-AEP----G---YTVISTISYA-LLLVLLLAGVYF-L-VERLD------IDAKVTALYALFPWVLFG-GALRTVED-----A---S----- 142
WP_049898450_1 DUF63 --AAQGAVV-ARP----G---YTIVSEVGYA-LTLLFMLIGVLL-L-IERLG------VGTDRRLLYALLPFALFG-GALRVVED-----A---N----- 145
WP_006077061_1 DUF63 --AAQGAVV-AEP----G---YTIISEIGYA-LTLLFMLIGVLL-L-IKRLG------VGTDRRLLYALLPFTLFG-GALRVVED-----A---N----- 145
WP_049996700_1 DUF63 --AAQGFTV-AEP----G---YTLVSEAGYA-IVLLFMLAGVLL-L-VRRLG------IGTDRRLFFALIPFTLFG-GALRVVED-----A---N----- 145
EMA38628_1 hypotheti --ASRGFTV-AEP----G---YTFVSEAGYA-LVLLFMLAGILL-L-VRRLG------IGTDRRVVFALIPFVLFG-GALRTVED-----A---A----- 145
WP_049992561_1 DUF63 ---PSDAFV-ATP----G---YTAVSEVGYM-IVGLFFLLGVIL-L-LRRLN------LGHNRNLFFGLVPFMLFG-GVLRVVED-----A---N----- 143
WP_010903025_1 DUF63 -----GGPV-AYP----G---YTLVSEVGYA-ATLLGALVGVHF-L-LERLR------IADSLALVYALTPFVLLG-GVVRVVED-----A---N----- 140
WP_009760947_1 DUF63 -----TEPV-AYP----G---YTLISEAGYA-VTLLASLIGVSF-L-LDHMR------VAEKTKFLYALFPFVLFG-GALRVVED-----A---N----- 140
WP_059057097_1 DUF63 -----TEPV-AYP----G---YTLVSEVGYA-VTLLVSLVGVTF-L-LDHLE------VGEDLRLLYALFPFVLLG-GALRVVED-----A---N----- 140
WP_058983522_1 DUF63 -----AEPV-AYP----G---YTLVSEVGYA-VTLLVSLVGVTF-L-LDHLE------VGEDLDLLYALFPFVLLG-GALRVVED-----A---N----- 140
WP_071932813_1 DUF63 -----AGPI-AYP----G---YTIVSEIGYA-ITLLVALAGIVF-L-LDRLD------FEERPQLIYPLLPFMLFG-GALRVVED-----A---H----- 140
WP_050048584_1 DUF63 -----TGPT-AYP----G---YTVVSEIGYG-LTLIGSLSGLVF-L-LRRLD------IGDRIAFIYPLVPFMLFG-GALRVVED-----A---M----- 139
WP_014051341_1 DUF63 -----AEPV-AEP----G---YTLVSEAGYV-VALLIGISGTVL-L-LENLD------VGEEARFFWAMVPFMLLG-SALRVVED-----A---H----- 144
WP_079233299_1 DUF63 -----AEPV-AYP----G---YTVVSEIGYI-VSLLLTITGVVM-L-LERLG------IGQRRGLFWAMVPFMLLG-GALRTVED-----A---H----- 144
WP_053948572_1 DUF63 -----AEPV-AYP----G---YTVVSEIGYI-VSLLLTITGVVL-L-LERLG------IGEERGLFWAMVPFMLLG-GALRTVED-----A---H----- 144
WP_049980835_1 DUF63 -----AEPV-AEP----G---YTVVSEIGYI-VSLLITITGVVL-L-LDYLD------IGEDRGLFWAMVPFMLLG-GALRTVED-----A---H----- 144
KPN31749_1 hypotheti -----AEPV-AEP----G---YTVVSEIGYI-VALLITITGVVL-L-LEYLN------IGEDRGLFWAMVPFMFLG-GALRTVED-----A---H----- 94
AGB16040_1 putative  --GPDAGPT-ASP----G---YTFTSYAGYI-PTLLLLLTGFIF-A-IERLD------IERYRAGFWGLFPFMLFG-GALRTVED-----A---N----- 139
WP_007696755_1 DUF63 --GSGAGPT-ASP----G---YTVTSYAGYI-PTLLLLLTGFIF-V-IDRLD------IERYRAGFWGLFPFMLFG-GALRAVED-----A---N----- 139
WP_015322904_1 DUF63 --GTGAGPT-AEP----G---YTAVSYAGYI-PTLLLLLIGIYF-I-IDWLD------IERYRAGFFALFPFMLFG-GALRTVED-----A---N----- 139
WP_076581869_1 DUF63 --TGDDGPT-ASP----G---YTAVSYAGYI-PTLILMAVGILF-A-IRRLD------IDRYRAGFFALFPFMLFG-GALRTVED-----A---N----- 139
WP_005559622_1 DUF63 --GANAGPT-AEP----G---YTAISYAGYI-PTLLLLLVGIYF-I-IDWLE------IERYRAGFFALFPFMLFG-GALRTVED-----A---N----- 139
```

```
WP_006067640_1 DUF63 --TGDFGPT-AAP----G---YTAISYA------XILMAIGVLF-V-LRRLE------IGSFRGGFFALFPFMLFG-GALRTVED-----A---N----- 134
WP_008164554_1 DUF63 --TGDFGPT-AAP----G---YTAISYAGYI-PTLILMAIGILF-V-LRRLE------IGSFRGGFFALFPFMLFG-GTLRTVED-----A---N----- 139
WP_006088112_1 DUF63 --GPDDGPF-ASP----G---YTVVSYAGYI-PTLILMAIGILF-A-IRRLD------IDRYRAGFFALFPFMLFG-GALRTVED-----A---N----- 139
WP_012943247_1 DUF63 --GPNAGPT-AEP----G---YTFVSYAGYI-PTLVLMAIGILF-L-VRRLE------IERYRAGFFALFPFMLFG-GALRVVED-----T---S----- 139
WP_008895026_1 DUF63 --GANAGPT-AEP----G---YTFVSYAGYI-PTLILMAIGVLF-L-VRRLE------IERYRAGFFALFPFMLFG-GALRVVED-----T---S----- 139
WP_098727043_1 DUF63 --PADASPT-AEP----G---YTFVSYAGYI-PTLVLMLIGVIF-L-VRRLE------IERYRAGFFALFPFMLFG-GALRVVED-----A---N----- 139
WP_049990861_1 DUF63 --V--SGPT-AEP----G---YTFVSYAGYI-PTLVLLLIGVIF-L-VRRLE------IERYRAGFFALFPFMLFG-GALRVVED-----V---N----- 137
WP_008013001_1 DUF63 --GPNAGPT-AEP----G---YTFVSYAGYI-PTLVLLLVGVIF-I-LRRLE------IERFRTGFFALFPFMLFG-GALRVVED-----A---N----- 139
WP_006180566_1 DUF63 --AANAGPT-AEP----G---YTFVSYAGYI-PTLVLLLIGIIF-L-VRRLE------IERYRAGFFALFPFMLFG-GALRVVED-----A---N----- 139
WP_006650865_1 DUF63 --ATNAGPT-AEP----G---YTFVSYAGYI-PTLVLLLIGIIF-L-VRRLE------IERYRAGFFALFPFMLFG-GALRVVED-----A---N----- 139
WP_066301295_1 DUF63 --ATDAGPT-AEP----G---YTFVSYAGYI-PTLVLLLIGIIF-L-VRRLG------IERYRAGFFALFPFMLFG-GALRVVED-----A---N----- 139
WP_049966804_1 DUF63 --ATNAGPT-AEP----G---YTFVSYAGYI-PTLVLLLIGIIF-L-VRRLE------IERYRAGFFALFPFMLFG-GALRVVED-----A---N----- 139
WP_076145380_1 DUF63 --AADAGPT-AEP----G---YTFVSYAGYI-PTLVLLLIGIIF-L-VRRLE------IERYRAGFFALFPFMLFG-GALRVVED-----A---N----- 139
WP_097378845_1 DUF63 --GPNAGPT-AEP----G---YTFVSYSGYI-PTLVLLLIGIIF-L-IRRLE------IERYRAGFFALFPFMLFG-GALRVVED-----V---N----- 139
WP_008452261_1 DUF63 --GAGAGPT-AEP----G---YTFVSYGGYI-PTLVLLLTGIIF-L-IRRLE------IERYRAGFFALFPFMLFG-GALRVVED-----V---N----- 139
WP_008452261_1 DUF63 --GAGAGPT-AEP----G---YTFVSYGGYI-PTLVLLLTGIIF-L-IRRLE------IERYRAGFFALFPFMLFG-GALRVVED-----V---N----- 139
WP_006432643_1 DUF63 --GTGAGPT-AEP----G---YTFVSYSGYI-PTLVLLLVGVIF-L-VRRLE------IERYRAGFFALFPFMLFG-GALRVVED-----V---N----- 139
WP_008452261_1 DUF63 --GAGAGPT-AEP----G---YTFVSYGGYI-PTLVLLLTGIIF-L-IRRLE------IERYRAGFFALFPFMLFG-GALRVVED-----V---N----- 139
WP_007109662_1 DUF63 --GAGAGPT-AEP----G---YTFVSYGGYI-PTLVLLLTGIIF-L-IRRLE------IERYRAGFFALFPFMLFG-GALRVVED-----V---N----- 139
WP_049952821_1 DUF63 --PGDAGPT-ASP----G---YTVQSYAGYI-PTLVILLIGIIF-I-IKRLD------IDRYRAGFYGLFPFFLFG-GVLRVVED-----A---N----- 139
WP_086889541_1 DUF63 QEGVATGPT-AAP----G---YTFVSYAGYI-PTLVLFLVGIVF-V-IQRLE------IERYRAGFYALFPFMLFG-GALRVVED-----A---N----- 142
WP_005580009_1 DUF63 SEGVATGPT-ASP----G---YTFVSYAGYI-PALVLGVIGIVF-L-VRRLG------IERYRAGFYALFPFILFG-GALRTVED-----A---N----- 142
WP_049927356_1 DUF63 --SPSDGPF-AEP----G---YTFVSYAGYI-PTLILFVIGIVL-L-IRRLD------IQRYRAGFYGLFPFMLFG-GALRVVED-----S---N----- 139
WP_087714455_1 DUF63 --GPDDGPF-AEP----G---YTFVSYAGYI-PTLILGVIGIIF-L-IQRLE------IERYRAGFYGLFPFMLFG-GALRVVED-----V---N----- 139
WP_013878438_1 DUF63 --GPNAGPT-ASP----G---YTFVSYAGYI-PTLILGVIGVVF-L-IKRLE------IERYRAGFYGLFPFMLFG-GALRVVED-----A---N----- 139
WP_049921111_1 DUF63 --GPNAGPT-AEP----G---YTFVSYAGYI-PTLVLFLTGIVF-L-VGRLD------IDRYRAGFYGLFPFMLFG-GALRVVED-----M---N----- 139
WP_007142789_1 DUF63 ---SGAGPT-AEP----G---YTFVSYAGYI-PTLVLAVIGVVF-L-IKRLE------IERYRAGFYGLFPFMLFG-GALRVVED-----T---N----- 138
WP_006651941_1 DUF63 --GPDAGPT-ASP----G---YTFVSYAGYI-PTLILGVIGIVF-L-IDRLE------IERYRAGFYGLFPFMLFG-GALRVVED-----A---N----- 139
WP_004216437_1 DUF63 --GPDAGPT-ASP----G---YTFVSYAGYI-PTLILGVIGIVF-L-IDRLE------IERYRAGFYGLFPFMLFG-GALRVVED-----A---N----- 139
WP_071402513_1 DUF63 --GPDAGPT-ASP----G---YTFVSYAGYI-PTLILGVIGIVF-L-IDRLE------IERYRAGFYGLFPFMLFG-GALRVVED-----A---N----- 139
WP_006666462_1 DUF63 --GVDAGPT-ASP----G---YTFVSYAGYI-PTLILSVVGVVF-L-VDRLE------IERYRAGFYGLFPFMLFG-GALRVVED-----S---N----- 139
WP_049904535_1 DUF63 --GVDAGPT-ASP----G---YTFVSYAGYI-PTLILSVVGVVF-L-IDRLE------IERYRVGFYGLFPFMLFG-GALRVVED-----S---N----- 139
WP_006824405_1 DUF63 --GVGAGPT-ASP----G---YTFVSYAGYI-PTLILSVVGVVF-L-IDRLE------IERYRAGFYGLFPFMLFG-GALRVVED-----S---N----- 139
WP_011323802_1 DUF63 --E---GFV-AEP----G---YTTVSTVSYA-VVLVFMLIGVLF-L-LRRLD------IEMSPAFYFALFPYMLLG-GALRTVED-----V---N----- 140
WP_015409719_1 DUF63 --E---GIV-AEP----G---YTAVSTVSYA-VVLVFMLVGVLL-L-LRRLD------VRMTPRFYVALFPFMLLG-GALRVVED-----V---N----- 141
WP_006883947_1 DUF63 --T---GVV-AET----G---YTTVSTFGYM-AVLVFMLGGVYL-L-LDRLD------VGGDKRLFLSLVPFMLFG-GAVRVVED-----A---T----- 140
WP_077207545_1 DUF63 --MAAGKVV-AEP----G---YTIVSEVGYM-ILLLFGLIGVLE-L-VRRLG------VGRDRDLFFALVPFMLFG-GALRVVED-----A---N----- 144
ESS12903_1 putative  --A---GVV-AEP----G---YTTVSTVSYG-LVLILLLFGIVF-L-LDRVD------AGSDRGFFYALFPFMLFG-GALRTVED-----A---N----- 144
WP_008417361_1 DUF63 --T---GIV-AEP----G---YTTVSTLSYA-VVLVFALLGVVF-F-LKRLD------IGTRPGFFFALFPFMLFG-GALRTVED-----A---S----- 142
WP_066381608_1 DUF63 --T---GIV-AEP----G---YTTVSTVSYA-IVLVFALLGVVF-F-LRRLD------IGTGQGFFFALFPFMLFG-GALRTVED-----A---S----- 137
WP_049947642_1 DUF63 --A---EPV-AEP----G---YTLVSEVGYV-VVLLVALSGVIL-L-LRKLN------IARDTDFFLALVPFFFFG-GALRTVED-----A---D----- 146
ESS05953_1 putative  --A---GIV-AYP----G---YTLVSEAGYV-VVLLLALSGVVL-L-LRRLR------LGSSRGLVVALLPLVLFG-GALRVVED-----A---T----- 143
WP_096390051_1 DUF63 --AAAGSVV-ASP----G---YTTVSTVSYI-VILLFALIGVLF-L-LRRLG------IGEERRFFFALFPFVLLG-GALRTIED-----A---G----- 149
WP_021072749_1 DUF63 --AAADAVV-ASP----G---YTIVSTVSYI-VILLFALIGVLL-L-LRRLG------IGEERRFFFALFPFVLLG-GALRTIED-----A---G----- 149
WP_049982791_1 DUF63 --VDAGAVV-AFP----G---YTTVSTVSYV-VILLGMLVGVVF-L-LQRFD------IGNELRFFYALFPFMLLG-GTLRTVED-----A---G----- 147
WP_008585990_1 DUF63 --RAAGEVV-ASP----G---YTTLSTVSYV-VILLAMLIGVVF-L-LRRLD------IATELRFFYALFPFMLFG-GAMRTVED-----A---G----- 149
WP_006628721_1 DUF63 --VSAGEVV-ATP----G---YTTFSTVSYV-VILLAMLIGVVF-L-LRRLD------IATELRFFYALFPFMLFG-GAMRTVED-----A---G----- 147
WP_006113218_1 DUF63 --VSAGEVV-ASP----G---YTTFSTVSYV-VILLAMLIGVVF-L-LRRLD------IATELRFFYALFPFMLFG-GAMRTVED-----A---G----- 148
WP_049930034_1 DUF63 --RSAGEVV-AAP----G---YTTLSTVSYV-VILLAMLVGVVF-L-LRRLD------IATELRFFYALFPFMLFG-GAMRTVED-----A---G----- 149
WP_049906047_1 DUF63 --ASAGEVV-ATP----G---YTTFSTVSYV-VILLAMLIGVVF-L-LRRLD------IATELRFFYALFPFMLFG-GAMRTVED-----A---G----- 149
WP_044965494_1 DUF63 --RSAGEVV-AAP----G---YTTVSTVSYV-VILLAMLIGVVF-L-LRRLD------IATELRFFFALFPFMLFG-GAMRTVED-----A---G----- 149
WP_096393195_1 DUF63 --RSAGEVV-ASP----G---YTTFSTVSYV-VILLAMLIGVVF-L-LRRLD------IATELRFFYALFPFMLFG-GAMRTVED-----A---G----- 149
WP_049908400_1 DUF63 --RSAGEVV-ATP----G---YTTFSTVSYV-VILLAMLIGVVF-L-LRRLD------IATELRFFFALFPFMLFG-GAMRTVED-----A---G----- 149
WP_049908585_1 DUF63 --RSAGEVV-ATP----G---YTTFSTVSYV-VILLAMLIGVVF-L-LRRLD------IATELRFFFALFPFMLFG-GAMRTVED-----A---G----- 149
WP_049902631_1 DUF63 --RSAGEVV-ATP----G---YTTFSTVSYV-VILLAMLIGVVF-L-LRRLD------IATELRFFFALFPFMLFG-GAMRTVED-----A---G----- 149
WP_049983668_1 DUF63 --RGAGEVV-AAP----G---YTTFSTVSYV-VILLAMLIGVVF-L-LRRLD------IATELRFFYALFPFMLFG-GAMRTVED-----A---G----- 148
WP_053772267_1 DUF63 --RSAGEVV-AAP----G---YTTFSTVSYV-VILLAMLIGVVF-L-LRRLD------IATELRFFYALFPFMLFG-GAMRTVED-----A---G----- 148
WP_007999151_1 DUF63 --VDGGAVV-AFP----G---YTTVSTVSYV-VILLGMLIGVVF-L-LQRLD------IGNELRFFYALFPFMLLG-GALRTVED-----S---G----- 147
WP_015909917_1 DUF63 --VEAGEAV-AFP----G---YTTVSTVSYV-VILLAMLIGVVF-L-LRRLD------IGTELRFFYALFPFMLLG-GALRTVED-----S---G----- 148
WP_004050594_1 DUF63 --VEAGEVV-AFP----G---YTTVSTVSYV-VILLAMLIGVVF-L-LRRLE------IGTELRFFYALFPFMLLG-GALRTVED-----S---G----- 148
WP_095636035_1 DUF63 --LEAGEVV-AEP----G---YTTVSTVSYV-VILLAMLIGVVF-L-LRRLE------IGTELRFFYALFPFMLLG-GALRTVED-----A---G----- 148
WP_008003811_1 DUF63 --VEAGGVV-AEP----G---YTTVSTVSYV-VILLAMLIGVVF-L-LRRLD------IGTELRFFYALFPFMLLG-GALRTVED-----S---G----- 148
WP_066416034_1 DUF63 --VAAGEVV-AYP----G---YTTVSTVSYI-VVLLAMLIGVVF-L-LRRLD------IATEFRFFYALFPFMLFG-GALRTVED-----A---G----- 149
ESS03170_1 putative  --TAAGGVV-ASP----G---YTTVSTLSYV-VILLGMLIGVVF-L-LRRLD------IATEFRFFYALFPFMLFG-GALRTVED-----A---G----- 148
WP_089671383_1 DUF63 --ETAGHIV-AYP----G---YTLVSEIGYM-LMLLFTLFGVIL-L-LRRLD------IGTDRRFFYALFPFMLLG-GALRTIED-----A---G----- 147
ERH07656_1 putative  --ERAGEIV-AYP----G---YTLVSEVGYV-LVLLVALIGVYL-L-LRRLK------IGDSRSLFYALFPFMLFG-GALRTVED-----A---G----- 147
ESS10096_1 putative  --ERAGEIV-AYP----G---YTLVSEVGYV-LVLLVALIGVYL-L-LRRLK------IGDSRSLFYALFPFMLFG-GALRTVED-----A---G----- 147
ESS07824_1 putative  TTSEAAGEIVAYP----G---YTLVSEVGYI-LILLFALVGVYL-M-LNRLE------IGDSRELFYALVPFILFG-GALRTVED-----A---G----- 147
ERH05689_1 putative  --EAAGEIV-AYP----G---YTLVSEVGYI-LILLFALVGVYL-M-LDRLE------IGDSRELFYALVPFILFG-GALRTVED-----T---G----- 147
ERH02266_1 putative  --EAAGEIV-AYP----G---YTLVSEVGYI-LILLFALVGVYL-M-LDRLE------IGDSRELFYALVPFILFG-GALRTVED-----T---G----- 147
ESS07825_1 putative  -----------------M--------------TLLTLSVGYLTY-I------------------------------------------------G----- 15
WP_049970017_1 DUF63 --P---EPV-AEP----G---YTLVSEIGYA-VTLIVALVGLVF-L-LRRLG------IGRDKSLFYALFPFMLFG-GALRVVED-----A---D----- 142
WP_007979289_1 DUF63 --P---EPV-AVP----G---YTLVSEVGYA-VTLIVALLGVVL-L-LRRLN------VGRDRGLFYALFPFMLFG-GALRVVED-----A---N----- 142
WP_007977745_1 DUF63 --P---GPV-AEP----G---YTLVSEVGYA-VTLIVALMGVVY-L-LRRLN------VGQERRLFYALFPFMLFG-GALRVVED-----A---N----- 142
WP_014556445_1 DUF63 --T---EPV-AYP----G---YTLVSEVGYM-ITLLVALIGVVF-L-LRQLK------IGSDRQLFYALTPFIFFG-GALRVIED-----A---N----- 144
WP_049935775_1 DUF63 --P---EPV-AFP----G---YTLVSEVGYV-VILLIALSGLVF-L-LDRLD------IGSDRDLFFALLPFVFFG-GALRVVED-----V---T----- 144
WP_008325149_1 DUF63 --V---EPV-AYP----G---YTLVSEVGYM-VTLIIALTGVVF-L-LRRLD------IGDDPRFFFSLVPFMFFG-GAFRVIED-----A---N----- 144
WP_007543631_1 DUF63 --A---EPV-AYP----G---YTLVSEVGYM-VTLIIALTGVVF-L-LRRLD------IGDDPRFFFSLVPFMFFG-GAFRVIED-----A---N----- 144
WP_049905041_1 DUF63 --A---EPV-AYP----G---YTLVSEVGYM-VTLVIALSGVVF-L-LRRLD------IGESPRFFYALFPFMLFG-GAFRVVED-----A---N----- 144
WP_049913563_1 DUF63 --A---EPV-AYP----G---YTLVSEVGYM-VTLVIALSGVVF-L-LRRLD------IGERPRFFYALFPFMLFG-GAFRVVED-----A---N----- 144
WP_049967851_1 DUF63 --A---EPV-AYP----G---YTLVSEVGYM-VTLVIALSGVVF-L-LRRLD------IGERPRFFYALFPFMLFG-GAFRVVED-----A---N----- 144
WP_049914905_1 DUF63 --A---EPV-AYP----G---YTLVSEVGYM-VTLVIALSGVVF-L-LRRLD------IGERPRFFYALFPFMLFG-GAFRVVED-----A---N----- 144
WP_049916430_1 DUF63 --A---EPV-AYP----G---YTLVSEVGYM-VTLVIALSGVVF-L-LRRLD------IGERPRFFYALFPFMLFG-GAFRVVED-----A---N----- 144
WP_049896892_1 DUF63 --A---EPV-AYP----G---YTLVSEVGYM-VTLVIALSGVVF-L-LRRLD------IGERPRFFYALFPFMLFG-GAFRVVED-----A---N----- 144
WP_049896892_1 DUF63 --A---EPV-AYP----G---YTLVSEVGYM-VTLVIALSGVVF-L-LRRLD------IGERPRFFYALFPFMLFG-GAFRVVED-----A---N----- 144
WP_049896892_1 DUF63 --A---EPV-AYP----G---YTLVSEVGYM-VTLVIALSGVVF-L-LRRLD------IGERPRFFYALFPFMLFG-GAFRVVED-----A---N----- 144
WP_058828253_1 DUF63 --A---EPV-AYP----G---YTLVSEVGYM-VTLVIALSGVVF-L-LRRLD------IGESPRFFYALFPFMLFG-GAFRVIED-----A---N----- 144
WP_058568959_1 DUF63 --A---EPV-AYP----G---YTLVSEVGYM-VTLVIALSGVVF-L-LRRLD------IGESPRFFYALFPFMLFG-GAFRVVED-----A---N----- 144
WP_049917947_1 DUF63 --A---EPV-AYP----G---YTLVSEVGYM-VTLVIALSGVVF-L-LRRLD------IGERPRFFYALFPFMLFG-GAFRVVED-----A---N----- 144
WP_049920348_1 DUF63 --A---EPV-AYP----G---YTLVSEVGYM-VTLVIALSGVVF-L-LRRLD------IGESPRFFYALFPFMLFG-GAFRVVED-----A---N----- 144
WP_089777545_1 DUF63 --A---EPV-AYP----G---YTLVSEVGYM-VTLVIALSGVVF-L-LRRLD------IGESPRFFYALFPFMLFG-GAFRVVED-----A---N----- 144
WP_008320633_1 DUF63 --A---EPV-AYP----G---YTLVSEVGYM-VTLIIALTGVVF-L-LRRLD------IGEDPRFFFALVPFMFFG-GAFRVVED-----A---N----- 144
WP_004060355_1 DUF63 --A---EPV-AYP----G---YTLVSEVGYM-VTLIIALTGIVF-L-LRRLD------IGEDPRFFFALVPFMFFG-GAFRVVED-----A---N----- 144
WP_103426078_1 hypot --T---GVV-AYP----G---YTLVSEVGYM-VTLLIALTGVVF-L-MRRLD------IGRERRFFYALIPFVFFG-GALRVVED-----A---N----- 144
WP_009367433_1 DUF63 --P---EPV-AYP----G---YTLVSEVGYI-VVLLLALIGIVF-L-LRRLD------IGTDREFFFALFPFMLFG-GALRVVED-----A---N----- 144
```

```
WP_013440552_1 DUF63 --A---EPV-AYP----G---YTLVSEVGYM-LTLLFALTGVVF-L-MRRLD------IGTDRNFFYSLLPFMFFG-GALRVVED-----A---N----- 144
WP_049916626_1 DUF63 --A---EPV-AYP----G---YTLVSEVGYM-LTLLFALTGVVF-L-MRRLD------IGRKRSFFYALLPFMFFG-GALRVVED-----A---N----- 144
WP_058582837_1 DUF63 --T---GFV-AYP----G---YTLVSEAGYV-VTLLLMLGGVVL-L-LRRLN------IGTSRKFFYALFPFMLFG-GALRVVED-----A---G----- 144
WP_101298124_1 hypot --E---GFV-AYP----G---YTLVSEVGYI-VTLLLMLMGVVL-L-LRRLG------IGEDRTFFFALFPFMLFG-GALRVVED-----A---G----- 144
PIN95153_1 hypotheti --------------------------------IALIFAVIGIVK-LLV----------IIIDRRSWF----------PIIRGIYG---------N----- 41
PIU22108_1 hypotheti -----------DA----G---YNIVNTFAYVILALVLFFFVLLP-L-VKRWY------GKLDLHFFLYTLSIIFIG-AMLRVAEE---------S----- 73
AAR39201_1 NEQ352    -----------------M---YTIANSIAYAIIFLFLIPYLLYK-L---GID------VEKETTKFFPLILSFLFI----RVFVD-----I---N----- 53
OIR14399_1 hypotheti ---------------------YNTYNTILYA-FLFYLGFVIIRD-L-LKKWK------VKIDDDFMFATFPLIVAG-GAIRVLED-----A---G----- 108
OIR20963_1 hypotheti -DACEVGDVDCDK----TGAKYNIYNTIAYG-FCFFIFFTIINE-L-LEYWK------IVIDDKFVFNSIPLLILG-GVVRTLED-----A---D----- 113
OIR22371_1 hypotheti -----------DSKGEAGAK-YNSYNTVAYG-LGFFVLFMAINE-L-LTRWK------IELNEKFVFSCIPLLILG-GVARVLED---------A----- 123
EGQ43935_1 putative  -----------------ESIYYNPYNTALYA-IVFGILVIYVLKPLIQKRMN------LKINKEFVLGFTPFIVFG-GAARALKD--------------- 79
MAG21679_1 hypotheti ----------GGQ--------FSPVEYLIYGAIMLALAFFVIFP-V-LDRRG------IKFDLRFAMAIIPFILLG-SLIHVLED-----M---A----- 77
PIN85618_1 hypotheti ---------PSVQ----G---YNIVNTSVYALILFVIVYFWLYP-L-LKKQD------IKFSYGFALALLPFIVFG-SALRVLND-----M---GILHKT 83
PIN99249_1 hypotheti -----------HS----G---YNLVNTLVYAAILLGIAFFVVFP-F-FKKKG------ISFDFDFFKAVFPFIVLG-STIRIFEE---------N----- 75
AJF59838_1 hypotheti -----------YS----G---YNFVNTLTYGIILIGIAFYLVFP-F-FNKRG------IRFDFRFGLAVLAFVLFG-STLRILED-----LKILG----- 81
WP_042682145_1 DUF63 -----------RE----G---YNPYNTLVYA-ILLGLGVIYSYK-YIIKPLK------IKVDEKLFWAVTPMVVFG-ATVRALVD-----G---G----- 79
WP_013468033_1 DUF63 -----------RE----G---YNPYNTLVYA-FLLGLGVIYSYK-YIIKPLK------IKVDEKLFWAVTPMVVFG-ATVRALVD-----G---G----- 79
WP_055281581_1 DUF63 -----------RE----G---YNPYNTLVYA-LVLGFAIIYTYK-WIIRPLR------IRVDEKLFYAVTPMVIFG-STVRALVD-----G---G----- 78
WP_048160382_1 DUF63 -----------RE----G---YNPYNTLVYA-IILGLAIIYTYR-WIIKPLK------IKVDEKLFYAVTPMTVFG-ATVRALID-----G---G----- 78
WP_042701216_1 DUF63 -----------RE----G---YNPYNTLVYA-IILGLAIIYTYR-WIIKPLK------IKVDEKLFYAVTPMVVFG-ATVRALVD-----G---G----- 78
WP_004068659_1 DUF63 -----------RE----G---YNPYNTLVYA-VILGLAIIYTYR-WIIQPLR------IKVDEKLFYAVTPMVVFG-ATVRALVD-----G---G----- 78
WP_058946665_1 DUF63 -----------RE----G---YNPYNTIVYA-VVLGLAIIYTYR-WIIRPLK------IKVDERLFYAVTPMVVFG-STVRALVD-----G---G----- 78
WP_014835499_1 DUF63 -----------RE----G---YNPINTVVYA-LIFGFAVIYTYK-YIIKPLK------IKVDERLFLAVTPMTIFG-ATLRALVD-----G---G----- 78
WP_010884243_1 DUF63 -----------RE----G---YNPINTVVYA-LIFGFAVIYTYK-YIIKPLK------IKVDERLFLAVTPMVIFG-STIRALVD-----G---G----- 81
WP_013748129_1 DUF63 -----------RE----G---YNPINTVVYA-LIFGFAVIYTYK-YIIKPLK------IRVDERLFLAVTPMVIFG-STIRALVD-----G---G----- 77
WP_010867251_1 DUF63 -----------RE----G---YNPINTVVYA-LIFGFAVIYTYK-YVIKPLK------IKVDERLFLAVTPMVIFG-STVRALVD-----G---G----- 77
WP_014733359_1 DUF63 -----------RE----G---YNPINTLVYA-LVFGFAVIYTYR-YIIKPLK------IKVDERLFLAVTPMVIFG-ATVRALVD-----G---G----- 78
WP_068319891_1 DUF63 -----------RE----G---YNPINTVVYA-LIFGFAVIYTYK-YIIKPLK------IKVDERLFLAVTPMVIFG-STVRALVD-----G---G----- 78
WP_068576049_1 DUF63 -----------RE----G---YNPINTVVYA-LIFGFAVIYTYK-YIIKPLK------IRVDERLFLAVTPMVIFG-STVRALVD-----G---G----- 78
WP_014013692_1 DUF63 -----------RS----G---YNPINTLVYA-LMLGLGAIYTYR-YILKPLK------IKIDKTLFMAVTLMVVFG-STVRALVD-----G---G----- 83
WP_014789520_1 DUF63 -----------RS----G---YNPVNTLVYA-LLLGFGAIYTYR-YILKPLR------IKIDRTLFMAVTLMVVFG-STVRALVD-----G---G----- 83
WP_088180676_1 DUF63 -----------RS----G---YNPINTLVYA-FMLGFGAIYTYR-YILKPLK------IKIDRTLFIAVTLMVVFG-STVRALVD-----G---G----- 83
WP_088864885_1 DUF63 -----------RS----G---YNPVNTLVYA-FMLGFGAIYTYR-YILKPLR------IKIDRTLFIAVTLMVVFG-STVRALVD-----G---G----- 83
WP_088856755_1 DUF63 -----------RS----G---YNPINTLVYA-LLLGFGAIYTYR-YILKPLR------IKIDQNMFIAVTLMVIFG-STVRALVD-----G---G----- 83
WP_088865981_1 DUF63 -----------RS----G---YNPINTLVYA-FMLGFGAIYTYR-YILKPLK------IKIDRTLFIAVTLMVVFG-STVRALVD-----G---G----- 83
WP_013906334_1 DUF63 -----------RE----G---YNPVNTLVYA-LIFGFAVIYTYR-YIIRPLR------IKVDERLFLAVTPMVVFG-ATVRALVD-----G---G----- 79
WP_088862658_1 DUF63 -----------RS----G---YNAVNTFVYA-LLFGLGVIYSYK-YIIKPLG------IKVDERLFLAVTPMVVFG-STVRALVD-----G---G----- 82
WP_088882956_1 DUF63 -----------RS----G---YNAINTLVYA-LLLGLGVIYSYR-YIIKPLG------IKVDGRLYLAVTPMVVFG-ATVRALVD-----G---G----- 82
WP_012571938_1 DUF63 -----------RS----G---YNPYNTLVYA-LLLGFGIIYSYK-YIIKPLK------IRIDERLFLAVTPMVVFG-ATVRALVD-----G---G----- 79
WP_068663933_1 DUF63 -----------RS----G---YNPYNTFVYA-LLLGFGIIYSYKHI-IKPLK------IRVDERIFLAVTPMVVFG-ATVRALVD-----G---G----- 79
WP_074631153_1 DUF63 -----------RS----G---YNPINTFVYA-FLLGLGVIYSYK-YIIKPLR------IKVDERLFLAVTPMVIFG-STVRALVD-----G---G----- 79
WP_058939074_1 DUF63 -----------RS----G---YNPVNTLVYA-LLLGFGVIYSYK-YIIKPLR------IKVDERLFLAVTPMVIFG-STVRALVD-----G---G----- 83
WP_088854681_1 DUF63 -----------RS----G---YNPVNTFVYA-FLLGLGVIYSYR-YIIKPLK------IPVDERLFWAVTPMVVFG-ATVRALVD-----G---G----- 83
WP_088885426_1 DUF63 -----------RE----G---YNPVNTFVYA-FLFGLGVTYSYR-YIIRPLK------IKVDERLFWAVTPMVVFG-ATVRALVD-----G---G----- 79
WP_011249694_1 DUF63 -----------RS----G---YNAVNTFVYA-LLFGLGVIYTYK-WIIKPLK------IPVDERLFWAVTPMVVFG-ATVRALVD-----G---G----- 81
WP_062386972_1 DUF63 -----------RS----G---YNAVNTFVYA-LLFGLGVIYTYK-WIIKPLK------IPVDERLFWAVTPMVVFG-ATVRALVD-----G---G----- 81
WP_010478887_1 DUF63 -----------RE----G---YNAVNTFVYA-LLFGLGVIYSYR-YIIKPLR------IKVDGRLFWAVTPMVVFG-ATVRALVD-----G---N----- 79
WP_088858605_1 DUF63 -----------RS----G---YNAINTFVYA-LLFGLGVIYSYR-YIIKPLR------IKVDERLFWAVTPMVVFG-STVRALVD-----G---G----- 79
WP_042690028_1 DUF63 -----------RS----G---YNAVNTFVYA-LLFGLGVIYSYR-YIIKPLK------IKVDERLFWAVTPMVVFG-ATVRALVD-----G---G----- 81
WP_048150762_1 DUF63 -----------RS----G---YNAINTLVYA-VLFGLGVTYSYR-YIIKPLG------IKVDERLFWAVTPMVVFG-ATVRALVD-----G---G----- 79
WP_048811076_1 DUF63 -----------RS----G---YNPINTLVYA-LLFGLGVVYSYR-YIIKPLK------IKVDERLFWAVTPMVVFG-ATLRALVD-----G---G----- 79
WP_050003256_1 DUF63 -----------RS----G---YNAVNTFVYA-LLFGLGVIYSYR-YIIKPLK------IKVDERLFWAVTPMVIFG-STVRALVD-----G---G----- 81
WP_062372100_1 DUF63 -----------RS----G---YNAVNTFVYA-LLFGLGVIYSYR-YIIKPLK------IKVDERLFWAVTPMVVFG-STVRALVD-----G---G----- 79
WP_048165284_1 DUF63 -----------RE----G---YNPYNTLVYG-FLLGLGIILTYDHI-IKRFK------IKVDERLIYATAPMVVFG-STMRALVD---------G----- 80
WP_048148750_1 DUF63 -----------RE----G---YNPYNTALYG-FLLGVGVILTYDHI-IKRFN------IKVDERLFYAVIPMVLFG-STTRALVD---------G----- 80
OYT53462_1 hypotheti ---------------------YSVTDYIIYL-LILIIVLDLMRR-FLFSY--------IKLDRKFIILISPYMFFG-IFVRLLAD-----V---G----- 58
KYC51152_1 hypotheti ---------------------YQPQDYVMYIGILLVLLGLALKF-L-FNK--------IKIDKTFIIAISPYMIMA-IALRVLAD-----V---G----- 58
KYC46003_1 hypotheti ---------------------YQPQDYVMYIGILLVLLGLALKF-L-FNK--------IKIDKTFIIAISPYMIMA-IALRVLAD-----V---G----- 58
KYC48643_1 hypotheti ---------------------YQPQDYVMYIGILLVLLGLALKF-L-FNK--------IKIDKTFIIAISPYMIMA-IALRVLAD-----V---G----- 58
KYC55321_1 hypotheti ---------------------YNPQDYFIYIGILLVLLTFALKF-L-FNK--------IKIDKTFIVAISPYMILA-IAIRVLTD-----V---GFYERS 63
KYC57927_1 hypotheti ---------------------YNPQDYFIYIGILLVLLTFALKF-L-FNK--------IKIDKTFIVAISPYMILA-IAIRVLTD-----V---G-FYER 62
KYC57171_1 hypotheti ---------------------YNPQDYFIYIGILLVLLTFALKF-L-FNK--------IKIDKTFIVAISPYMILA-IAIRVLTD-----V---G-FYER 62
OIO20701_1 hypotheti -----------HA----G---YNLVNTLAYA-LIALLFAYGIYL-A-LKKYK------VRVGKEFFLAVISFVLLG-STMRVVSD-----SVDTG----- 77
PIT83986_1 hypotheti -----------KS----G---YNIVNTFAYA-LAALCALFAVWK-I-LEWKK------INIGKEFWAGAVGFTFFG-SSLRVVID-----AVDSGAMANA 87
OIO24760_1 hypotheti -------PTNTAP--------YNAVNTLVFA-ALALAAAYLIFK-G-LKKFG------IKIDEELVKSLVPFILLG-SAVRAVVD-----A---GILPRV 85
OIO24701_1 hypotheti -------PGEAAP--------YNWANTLAYA-VAALVATFAIFK-G-LQRLK------IKIDGKFFAAILPFVLFG-SALRVAQD-----A---G----- 79
OIO27021_1 hypotheti -------PEAYAP--------YNWVNTLVYA-GLAVVAACALYK-L-LKHYG------FNPDEKLAKAVLPFVLFG-SALRVITD-----A---G----- 76
OIO26762_1 hypotheti -------PSAYAP--------YNWVNTLAYA-LVALVAAYIIYR-L-LRRFK------IAVDENLLKTVLPFVVFG-SALRVVQD-----A---N----- 77
PIN95811_1 hypotheti -------PSAYAP--------YNWVNTLAYA-LVALVAAYIIYR-L-LRRFK------IAVDENLLKTVLPFVVFG-SALRVVQD-----A---N----- 77
PIO01637_1 hypotheti -------PSAYAP--------YNWVNTLAYA-LVALVAAYIIYR-L-LRRFK------IAVDENLLKTVLPFVVFG-SALRVVQD-----A---N----- 77
PIO02820_1 hypotheti -------PEAYAP--------YNWVNTLVYA-GLAVVAACALYK-L-LKHYG------FNPDEKLAKAVLPFVLFG-SALRVITD-----A---G----- 76
PJD01038_1 hypotheti -------PEAYAP--------YNWVNTLVYA-GLAVVAACALYK-L-LKHYG------FNPDEKLAKAVLPFVLFG-SALRVITD-----A---G----- 76
PIZ91366_1 hypotheti -------PSAYAP--------YNWVNTLAYA-LVALVAAYIIYR-L-LRRFK------IAVDENLLKTVLPFVVFG-SALRVVQD-----A---N----- 77
WP_013100570_1 DUF63 -----------GS----G---YNLVQEITYG-LILALSLYLFYK-G-LKKLN------IKIDEKFAIPGVIFTILI-ALMRALVD-----L---G----- 78
WP_004590770_1 DUF63 -----------GS----G---YNIIQEITYG-AILAISLYLFYK-A-LIRLN------VKIDEKFAIPSIVFSILI-ALIRALVD-----L---G----- 78
WP_048196979_1 DUF63 -----------GT----G---YNIIQEITYG-TILAVFLYLFYK-A-LRKLN------IKIDEKFAIPGIVFSVLI-ALIRALVD-----L---G----- 78
WP_015791538_1 DUF63 -----------GT----G---YNLIQEITYG-IILALALYLFYK-A-LRKLN------INIDEKFAIPGIVFTVLI-ALMRALVD-----C---G----- 78
WP_048202292_1 DUF63 -----------GT----G---YNLIQEITYG-IILALALYLFYK-A-LRKLN------INIDEKFAIPGIVFTVLI-ALMRALVD-----C---G----- 78
WP_012981280_1 DUF63 -----------GT----G---YNIVQEITYG-IILALALYLFYK-A-LRKLN------INIDERFAIPGIVFTVLI-ALMRALVD-----C---G----- 78
WP_064496496_1 DUF63 -----------GT----G---YNIVQEITYG-IILALALYLFYK-A-LRKLN------INIDEKFAIPGIVFTVLI-ALMRALVD-----C---G----- 78
WP_011972741_1 DUF63 -----------QS----G---YTIIQEITYG-ILLFVMVYIFYR-A-CKSLK------IEIDSKFALTTVFYVILI-SLLRSMTD-----A---G----- 80
WP_013798289_1 DUF63 -----------GS----G---YNLVQEITYG-IFLTFMVYIFYK-A-CVKLR------VAIDEKFAQTTVFYVVLI-SLMRALVD-----A---G----- 78
WP_013181100_1 DUF63 -----------GT----G---YNLVQEFTYG-ILLFVMVYLFYK-I-AQKFQ------IEIDEKFAKVTIVYIILI-TLMRALVD-----A---G----- 80
WP_013867628_1 DUF63 -----------KS----G---YNIIQEITYG-ILLFVMVYLFYK-V-CKLLN------IKINKKFALTTIFYIILI-ALMRALVD-----A---G----- 80
WP_018153391_1 DUF63 -----------KT----G---YNLIQEITYG-TLLFVMVYIFYK-A-CNFLN------IRVDKRFAEVTIFYVILI-TLMRALVD-----A---G----- 80
WP_011170272_1 DUF63 -----------KQ----G---YNLIQEITYG-ILLFVMVYTFYR-M-CLKLK------IAIDRRFAEVTVFYVILI-TLMRALVD-----A---G----- 80
WP_012066151_1 DUF63 -----------KQ----G---YNLIQTATYG-VFLFFMVYLFYK-V-CLKLK------ITVDRQFAEVTVFYIILI-TLMRSLVD-----A---G----- 80
ODS43013_1 hypotheti -----------------G---YDIVDTLTYG-IILGLAVFGIIP-L-LRKYC------VQITKRFLVAVSPFIVFG-ATMRELVY-----R---G----- 73
OIQ06085_1 hypotheti -----------VE--------YTWFNTIMFG-IILVIALFFVIK-L-MKKLK------IEIDKNFVLALIPFILFG-STMRALAD-----A---E----- 76
PIN67069_1 hypotheti -----------VE--------YTWFNTIMFG-IILVIALFFVIK-L-MKKLK------IEIDKNFVLALIPFILFG-STMRALAD-----A---E----- 76
PIV28099_1 hypotheti -----------VE--------YTWFNTIMFG-IILVIALFFVIK-L-MKKLK------IEIDKNFVLALIPFILFG-STMRALAD-----A---E----- 76
PJC13070_1 hypotheti -----------VE--------YTWFNTIMFG-IILVIALFFVIK-L-MKKLK------IEIDKNFVLALIPFILFG-STMRALAD-----A---E----- 76
PIZ29927_1 hypotheti -----------VE--------YTWFNTIMFG-IILVIALFFVIK-L-MKKLK------IEIDKNFVLALIPFILFG-STMRALAD-----A---E----- 76
PKP60688_1 hypotheti -----------VE--------YTWFNTIMFG-IVLIIALFLVIK-L-MKKWN------IEIDKNFVFALLPFILFG-STMRALAD-----A---E----- 76
WP_012956954_1 DUF63 -----------------G---YTIFNTVIYT-LILLIFIIAIIK-M-FKKIK------ID-PISIIYPIIPYIFLG-SLIRALVD-----N---G----- 74
```

```
WP_067148430_1 DUF63 -----------------G---YTIFNTVVYT-LVLLIFILAIIK-M-FKKLE------ID-PLSIFFSIVPFIFLG-CSIRALVD-----N---G----- 74
WP_080460538_1 DUF63 -----------------G---YTIFNTIVYG-IILVLVLFMIIK-I-FKNYK------IN-PTDLIIPLIPFIFLG-SSVRALVD-----N---G----- 73
WP_010877090_1 DUF63 ----------LHP----G---YTPLNTVVFG-IVLGIAVLIILR-M-FRWLK-------KDPGELLVPLLPFIFLG-SGARALVD-----N---G----- 74
WP_013294901_1 DUF63 ----------LHP----G---YTPLNTVVFG-LVLGFAVLIILR-M-FRWLK-------KDPRELLVPLLPFIFLG-SGARALVD-----N---G----- 74
WP_010877090_1 DUF63 ----------LHP----G---YTPLNTVVFG-IVLGIAVLIILR-M-FRWLK-------KDPGELLVPLLPFIFLG-SGARALVD-----N---G----- 74
BAZ99473_1 hypotheti ----------LHP----G---YTPLNTVVFG-IVLGIAVLIILR-M-FRWLK-------KDPGELLVPLLPFIFLG-SGARALVD-----N---G----- 74
PKL66404_1 hypotheti ----------LHP----G---YTLFNTIIFG-LILGLVVMAIIK-L-FKWID-------KDPRDLFIALIPFIFFG-SSARALVD-----N---G----- 82
WP_023991171_1 DUF63 ----------LHP----G---YTTYNTIVFG-IILGLVILLIIR-M-FKWID-------KDPKGLFIPLIPFIFFG-SSARALVD-----N---G----- 75
WP_048072898_1 DUF63 ----------LHP----G---YTTYNTIVFG-IILGLVILLIIR-M-FRWID-------KDPKDLFIPLIPFIFFG-SSARALVD-----N---G----- 75
WP_100905180_1 hypot ----------LHP----G---YTTLNTIVFG-IILGMVILLIIR-M-FRWID-------KDPKDLFISIIPFIFFG-SSARALVD-----N---G----- 75
WP_100907231_1 hypot ----------LHP----G---YTTLNTIVFG-IILGMVILLIIR-M-FRWID-------KDPKDLFIPIIPFIFFG-SSARALVD-----N---G----- 75
WP_100907231_1 hypot ----------LHP----G---YTTLNTIVFG-IILGMVILLIIR-M-FRWID-------KDPKDLFIPIIPFIFFG-SSARALVD-----N---G----- 75
WP_048081925_1 DUF63 -----------HP----G---YTILNTVVFG-IILGIAVILIIK-M-FKYIK-------KDPKDLFIPLVPFIFFG-SSARALVD-----N---G----- 74
WP_048081925_1 DUF63 -----------HP----G---YTILNTVVFG-IILGIAVILIIK-M-FKYIK-------KDPKDLFIPLVPFIFFG-SSARALVD-----N---G----- 74
WP_069585525_1 DUF63 -----------HP----G---YTILNTVVFG-IILGIAVILIIK-M-FKYIK-------KDPKDLFIPLVPFIFFG-SSARALVD-----N---G----- 74
WP_069585525_1 DUF63 -----------HP----G---YTILNTVVFG-IILGIAVILIIK-M-FKYIK-------KDPKDLFIPLVPFIFFG-SSARALVD-----N---G----- 74
WP_013643671_1 DUF63 ----------LHP----G---YTTLNTVVFG-LILGISILLIIK-M-FKYIE-------KDPADLIIPLTPFIFFG-SGARALVD-----N---N----- 74
WP_013824571_1 DUF63 ----------LHP----G---YTLLNTVVFG-IILGISIILLIK-M-FKFLK-------KDPKDLVIPLIPFIFFG-SSARALVD-----N---G----- 74
WP_048192110_1 DUF63 ----------LHP----G---YTILNTVVFG-IILGISILLIIK-M-FKYIK-------KDPADLMIPLIPFIFFG-SSARALVD-----N---N----- 78
WP_081810122_1 DUF63 -----------DT----S---YNHFDMLTYF-IILFAGVYAVLK-L-LNRLK------IEINETFVIATIPYIIMG-SVFRVIED-----A---D----- 100
WP_097298965_1 DUF63 -----------DT----S---YNHFDMITYV-IILFAGVLAITK-L-LNKLR------IKVDEEFVIATIPFIFMG-SVYRVIED-----A---D----- 76
PIO00292_1 hypotheti -----------NL----G---YNYVNTFTYGIILIGLLILGDKV---LKRFN------ITFTRKMAFSLIPFILFG-STVRVLRD-----A---E----- 79
WP_011833679_1 DUF63 ----------GST--------YTVIQTILYA-LLVLLGLYILYR-C-LKKAN------IAIDTPLILTGLTYIILG-GLLRVVED-----T---G----- 72
WP_042698332_1 DUF63 -----------------T---YTVFQTILYA-LLVLLGLYILYR-C-LKKAD------IPIDTPLILTGLSYIILG-GLLRVVED-----T---G----- 72
WP_042698332_1 DUF63 -----------------T---YTVFQTILYA-LLVLLGLYILYR-C-LKKAD------IPIDTPLILTGLSYIILG-GLLRVVED-----T---G----- 72
WP_011448164_1 DUF63 ----------GQP--------YTLVDTLTYA-LILLVCIWLIYR-W-LEANH------ISIDREFVYSLIPWVVFG-GCLRVVED-----T---G----- 75
WP_007314214_1 DUF63 ----------GQP--------YNVVDTTTYA-LVLILAIYIIYR-W-LSRSR-----TILIDGDFVLVTLPYVVMG-GLLRVVED-----T---G----- 80
WP_042705632_1 DUF63 ----------GGA--------YNPVDTLTYA-TILIIAVYLVYR-W-LVRSK------IEINGDFIIALLPFVVLG-ATSRVVED-----T---G----- 75
WP_013329717_1 DUF63 ----------GGA--------YTIVDTLTYA-IILIVSVYLVYK-W-LIRAG------IEIDKDFVLSLIPYIVLG-GFLRVVED-----T---G----- 75
WP_004077722_1 DUF63 -----------GE----A---YTIVDTLTYA-LILILSVYLVYR-W-LERTG------ISIDRDFILAVIPYVVLG-GTLRVVED-----T---G----- 75
WP_048150927_1 DUF63 ----------GGA--------YTIVDTLTYA-IILIVSVYLVYK-W-LLRAG------IEIDKDFVLSLIPYIVLG-GFLRVVED-----T---G----- 75
WP_012107531_1 DUF63 ----------GEP--------YNIVETLTYA-IILIIGVYLLYR-W-FSNSAWLLDHGIKLDSSFILATLPYVVLG-GVLRVIQD-----T---G----- 81
PKL70081_1 hypotheti ----------EQP--------YNAVETITYA-LILIAALYLIYH-W-FKKSN------ISIDGPFVIATLPYVILG-GVLRVVQD-----T---H----- 75
WP_015285007_1 DUF63 ----------GQP--------YNVVDTLTYA-LILVVGVYLLYR-W-LSTSTWLSDIGFTIDATFILSTIPYVILG-GVMRVIED-----T---G----- 81
PKL64570_1 hypotheti -----------GE----A---YNIVDTLTYA-IILVAGVYLLYR-W-LTTSTWLTDIGFTIDQGFILATLPYVILG-GVMRVIED-----T---G----- 81
WP_015286404_1 DUF63 ----------GQP--------YNVVDTLTYA-IILIIAVYLIYR-W-LNSTG------ISVDRKFVLATVPFIVLG-GLLRVVED-----T---G----- 75
WP_012617618_1 DUF63 -----------GG----A---YTIVDTLTYA-LILIVSLYLIYQ-W-LNRTK------ISIDRSFIYSLLPFVVLG-SLLRVIQD-----T---G----- 75
WP_014867374_1 DUF63 -----------GQ----A---YTLVDTLTYA-LILIAAVYIVYR-G-LQRYK------IAVDDELVLATMPFVVLG-GLLRVVED-----T---G----- 75
CVK33108_1 conserved -----------GQ----A---YTLVDTLTYA-LILIAAVYIVYR-G-LQRYK------IAVDDELVLATMPFVVLG-GLLRVVED-----T---G----- 75
WP_066956331_1 DUF63 -----------GE----A---YTLVDTLTYA-LILIAAVYLVYR-G-LRRYK------IAVDDELVLATLPFVVLG-GLLRVVED-----T---G----- 75
WP_011844879_1 DUF63 -----------GE----A---YTLVDTLTYA-LILIAAVYLVYR-G-LRRYR------IAVDEELVLATLPFVVLG-GLLRVVED-----T---G----- 75
WP_048181868_1 DUF63 -----------GE----A---YTLVDTLTYA-LILIAAVYLVYR-G-LRRYR------IAVDDELVLATLPFVVLG-GLLRVVED-----T---G----- 75
WP_067073014_1 DUF63 -----------GE----A---YTLVDTLTYA-LILIAAVYLVYR-G-LRRYG------IAVDEELVLATLPFVVLG-GLLRVVED-----T---G----- 75
PKL62929_1 hypotheti -----------GE----A---YTLVDTLTYA-LILIVAVYLVYR-G-LRRYK------IAIDDELVLATLPFVVLG-GLLRVVED-----T---G----- 75
WP_004039890_1 DUF63 ----------GQP--------YTIVDTLTYA-LILILSVYLVYR-G-LRRFG------IEIDRDFVLATIPFVILG-GLLRVVED-----T---G----- 75
WP_067049225_1 DUF63 ----------GQP--------YTIVDTLTYA-LILIFSVWLVYR-G-LRRFG------IEVDRRFTLSTIPFVVLG-GLLRVVED-----T---G----- 75
KYK37884_1 hypotheti -----------GE----G---YNWVNTLTYG-VILIVAAYVVYQ---YLKTR------VTLDEKFYYSVSTFILVG-ASARVIKD-----M---H----- 74
KYK28019_1 hypotheti -----------GE----G---YNWVNTLTYG-LILIAASYLVYA---YLKTR------IAMDQKFYYSVSTFILVG-ASARVVKD-----M---N----- 74
OYT57835_1 hypotheti -----------GN----G---YNIVNTSAYA-LIFIASLFLLYK-I-FKRER------VKIDKNFLLGVLPYIILG-AVLRVLRD-----A---N----- 78
OYT33350_1 hypotheti -----------KE----G---YNVVNTLTWA-VILIFMVWIIYR-FLENRLS--------IDKTFLFSTIPYVIFG-SSIRIIED-----A---G----- 78
WP_012964745_1 DUF63 -----------KQ----G---YNIVNTLTWA-AILLVAAWLIYR-YLSKRLF--------FDKKFMLYTVPFIFFG-SSVRVVED-----A---G----- 75
WP_048091573_1 DUF63 -----------KQ----G---YNPVNTLTFA-ILLIVSVYLIYR-Y-LSPR-------VRFDVRFALYTFPYILLG-SSTRVVED-----A---G----- 75
WP_048096399_1 DUF63 -----------KE----G---YNPVNTMTFA-AILIVAIYLLYR-Y-LSPR-------VRFDLRFALYTFPYILLG-SSTRIVED-----A---G----- 75
WP_012940099_1 DUF63 -----------KT----G---YNPVNTVTWA-IILILAVYAIYK-YLSKRLS--------FDERFFISNVPYVFFG-SFLRVVED-----A---G----- 74
WP_010877969_1 DUF63 -----------KE----G---YNIVNTLTWA-AILIVAVYALYR-FLGKRLD--------FDEKFVYANIPYILLG-SSVRVVED-----A---G----- 87
WP_013682839_1 DUF63 -----------KQ----G---YNPVNTLTWA-LILVAAVILIYR-T-LKKYD------VKFDERFVAGNIPYVILG-ASVRVVED-----A---G----- 75
WP_015591141_1 DUF63 -----------KQ----G---YNIVNTLTFA-IILILAVVAIYK-YLSKR--------IKFDEKFVIANIPYILLG-SSVRVVED-----A---E----- 75
WP_012035887_1 DUF63 ----------AGE----G---YNPVNTVTYA-IILGISLFAILK-V-MDMLK------VRIDEKFVVSTVPFIIVG-ASLRVVQD-----V--------- 81
WP_014404626_1 DUF63 -----------NE----G---YNAVNTLTYA-IILGISLFAILK-L-MAFLK------LKVDERLIMATSPFIILG-ASLRNLED-----V--------- 75
BAI60262_1 conserved -----------NE----G---YNVVNTLTYA-IILAISLFAILK-L-MALLR------LKIDERLIMAVAPFIILG-ASLRNLED--------------- 74
WP_042684156_1 DUF63 -----------DT----G---YNVVNTLTWA-LVLGLSLFGVYR-L-LKRLG------VRIDSGFILALAPYIAMG-SLLRVVED-----A---R----- 84
OFV68024_1 membrane  ------------------------------------MCLFGLLR-L-LKYFE------IRIDTRFIFSLIPYIFVG-STLRVIED-----A---N----- 42
WP_013720253_1 DUF63 -----------DT----S---YNPVDTITWA-VVLGLAILGLVK-L-FRHYR------LAIDERLVLSTLPYILAG-SSLRVIED---------A----- 76
WP_014587756_1 DUF63 -----------DA----G---YNPVNTVTWA-LILGGMILLILR-L-LRRLE------VSFDERFFLATAPYVLAG-ASLRVVED-----A---E----- 74
ABK14947_1 Protein o -----------DS----G---YNPVNTVTWA-IVLGVFIVLIQR-A-FRRYG------IEVNSRFILATAPYIIAG-SSLRVIED-----A---E----- 81
OKY79140_1 putative  -----------DS----G---YNPVNTLTWG-ILLVLCFFGIIK-L-LDTLD------IKVNRKFVFSLIPFIFVG-STFRVFED-----A---G----- 78
WP_086637003_1 DUF63 -----------DA----G---YNPVNTITWA-ILLVIFLFISLK-I-LQKLE------IKLDRGFIYSLVPFIFIG-SGLRVVED-----A---G----- 78
WP_048089097_1 DUF63 -----------DA----S---YNPVDTITYA-ILLGLSVFGILK-L-LEKLD------IEIDSRFIAAVTPYILAG-SSLRVLED-----L---N----- 79
WP_097298272_1 DUF63 -----------DT----S---YNPVDTLTYA-IVLGISLFGVLK-L-LEKLN------VKVDSRFMISVTPYIIAG-CALRVLED-----S---K----- 80
WP_013897788_1 DUF63 -----------DT----G---YNIVNTLTWA-IILGISIFGVIK-I-LKRLD------IRIDNQFIMALLPFVFAG-ASFRVVED-----A---G----- 81
WP_015323982_1 DUF63 -----------DE----G---YNIINTLTWA-IILGLCIFAVVK-L-LKKLD------IRTDSRFIAAIVPFVLAG-SSMRVLED-----A---G----- 81
WP_011499115_1 DUF63 -----------DS----G---YNVVNTLTWA-LILGICVFAVAW-L-LEKWN------VEVDDKFVFSIIPFILAG-SSLRVIED-----A---G----- 81
WP_013037846_1 DUF63 -----------DS----G---YNPVNTLTWA-LILGVCVFGVIK-L-LDRMK------VEVDERFIYSIIPFVLAG-SSLRVLED-----A---N----- 81
WP_048205640_1 DUF63 -----------DS----G---YNPVNTLTWA-LILGICVFAVAK-L-LDKWK------VEVDERFVFSIIPFILAG-SSLRVMED-----A---G----- 81
WP_072561629_1 DUF63 -----------DS----G---YNPVNTLTWA-LILGACVFGVIK-L-LDRMK------VEVDERFIFSIIPFILAG-SSLRVLED-----A---N----- 81
WP_072360082_1 DUF63 -----------DS----G---YNPVNTLTWA-LILGACVFGVIK-L-LDRMK------VEVDERFIFSIIPFVLAG-SSLRVLED-----A---N----- 81
WP_096711793_1 DUF63 -----------DS----G---YNPVNTLTWA-LILGACVFGVIK-L-LDRMK------VEVDERFILTIIPFILAG-SSLRVLED-----A---N----- 81
ODV50598_1 hypotheti -----------DS----G---YNPVNTLTWA-LILGACVFGVIK-L-LDRMK------VEVDERFILTIIPFILAG-SSLRVLED-----A---N----- 81
WP_013194042_1 DUF63 -----------DT----G---YNPVNTITWA-IFLGIFLFGVIK-L-FKRLN------ISVDDRFILSIIPFILAG-SSLRVVED-----A---G----- 81
WP_048178067_1 DUF63 -----------DA----G---YNPVNTFTWA-VVLGICIFGIFR-L-LEKFN------VKVNSRFILSIIPFVLAG-SSLRVLED-----S---P----- 81
WP_048127111_1 DUF63 -----------DE----G---YNLVNTITWA-VVLGICIFGVFR-L-LKKMD------IKINPSFILSILPFVLAG-SSLRVLED-----S---P----- 81
WP_011023594_1 DUF63 -----------DE----G---YNLVNTFTWA-IVLGICIFGIFK-L-LKKLE------VKITPEFILSILPFVLAG-SSLRVLED---SPA---G----- 87
WP_048184587_1 DUF63 -----------DS----G---YNIVNTFTWA-IVLGICIFGIFK-L-LKKME------IKITPEFILSILPFVLAG-SSLRVIED-----S---P----- 81
WP_011032542_1 DUF63 -----------DE----G---YNIVNTFTWA-VVLGICIFGVFR-L-LEKME------VKITPKFILSLLPFVLAG-SSLRVLED---SPA---N----- 83
WP_011032542_1 DUF63 -----------DE----G---YNIVNTFTWA-VVLGICIFGVFR-L-LEKME------VKITPKFILSLLPFVLAG-SSLRVLED---SPA---N----- 83
WP_048129141_1 DUF63 -----------DE----G---YNLVNTFTWA-VVLGICIFGVFR-L-LNKLE------VKITPRFILSLLPFVLAG-SSLRVLED---SPA---G----- 83
WP_048129141_1 DUF63 -----------DE----G---YNLVNTFTWA-VVLGICIFGVFR-L-LNKLE------VKITPRFILSLLPFVLAG-SSLRVLED---SPA---G----- 83
WP_048169891_1 DUF63 -----------DD----G---YNLVNTFTWA-VVLGICIFGVFR-L-LEKLE------VKITPRFILSILPFVLAG-SSLRVLED---SPA---G----- 83
WP_048137175_1 DUF63 -----------DD----G---YNLVNTFTWA-VVLGICIFGVFR-L-LEKLE------VKITPRFILSILPFVLAG-SSLRVLED---SPA---G----- 83
WP_048137175_1 DUF63 -----------DD----G---YNLVNTFTWA-VVLGICIFGVFR-L-LEKLE------VKITPRFILSILPFVLAG-SSLRVLED---SPA---G----- 83
WP_048137175_1 DUF63 -----------DD----G---YNLVNTFTWA-VVLGICIFGVFR-L-LEKLE------VKITPRFILSILPFVLAG-SSLRVLED---SPA---G----- 83
WP_048137694_1 DUF63 -----------DE----G---YNLVNTFTWA-VILGICIFGIFR-L-LEKLE------VKITPRFILSILPFVLAG-SSLRVLED---SPA---G----- 83
WP_048168126_1 DUF63 -----------DE----G---YNIVNTFTWA-VVLGICIFGIYR-L-LEKLE------VKITPKFIISLLPFVLAG-SSLRVIED-----S---P----- 81
WP_048118082_1 DUF63 -----------DE----G---YNTVNTITWA-IILGICIFGVFR-L-LEKLE------VKITPRFIASVLPFVLAG-SSLRVIED---SPA---G----- 83
WP_048158120_1 DUF63 -----------DE----G---YNTVNTITWA-IILGICIFGVFR-L-LEKLE------VKITPRFIASVLPFVLAG-SSLRVIED---SPA---G----- 83
```

```
WP_011305329_1 DUF63 -----------DE----G---YNTVNTVTWA-IILGICIFGVFR-L-LEKLE------VKITPRFIASVLPFVLAG-SSLRVIED---SPA---G----- 83
WP_054298619_1 DUF63 -----------DE----G---YNIVNTFTWA-VVLGICIFGIYR-L-LEKLE------VKITPRFIASLLPFVLAG-SSLRVIED---SPA---G----- 83
ALK05385_1 hypotheti -----------DE----G---YNIVNTFTWA-VVLGICIFGIYR-L-LEKLE------VKITPKFIISLLPFVLAG-SSLRVIED---SPA---G----- 83
WP_015052897_1 DUF63 -----------DS----G---YNIVNTITWA-IILGICIFGIVK-L-LKRMN------VNITDRLTASVIPFILAG-SSLRVIED-----T---G----- 81
WP_023846134_1 DUF63 -----------DS----G---YNIVNTITWA-IVLGICIFGVVK-L-LKKLD------IEIDDRFTVSVIPFILAG-SSMRVVED-----T---G----- 80
OIN88451_1 hypotheti -----------GY----G---YNPVNTLAYA-LLFVSLTYLLFV-F-LKKSG------IKTDKRFVLAVAPWIIFG-ILLRILED-----M---G----- 75
PIX50278_1 hypotheti -----------GY----G---YNPVNTLAYA-LLFVSLTYLLFV-F-LKKSG------IKTDKRFVLAVAPWIIFG-ILLRILED-----M---G----- 75
PIW41402_1 hypotheti -----------GY----G---YNPVNTLAYA-LLFVSLTYLLFV-F-LKKSG------IKTDKRFVLAVAPWIIFG-ILLRILED-----M---G----- 75
PIY35178_1 hypotheti -----------GY----G---YNPVNTLAYA-LLFVSLTYLLFV-F-LKKSG------IKTDKRFVLAVAPWIIFG-ILLRILED-----M---G----- 75
PJB74886_1 hypotheti -----------GY----G---YNPVNTLAYA-LLFVSLTYLLFV-F-LKKSG------IKTDKRFVLAVAPWIIFG-ILLRILED-----M---G----- 75
PIZ33651_1 hypotheti -----------GY----G---YNPVNTLAYA-LLFVSLTYLLFV-F-LKKSG------IKTDKRFVLAVAPWIIFG-ILLRILED-----M---G----- 75
WP_048165029_1 DUF63 -----------NT----G---YNIVNTLTYA-IILGLAAMGIFK-V-LKKLN------VKYDNAFFRALIPYMILG-AFSRALTD-----A---T----- 76
WP_042681046_1 DUF63 -----------NT----G---YNVVNTLTYA-IILGIAALAVYK-I-LKRLG------IKYDNAFFRALIPYMIFG-AFTRALTD-----A---T----- 76
WP_013467319_1 DUF63 -----------NT----G---YNVVNTLTYA-IILGIAALGVYK-I-LKRLG------IKYDNAFFRALIPYMIFG-AFTRALTD-----A---T----- 76
WP_048152160_1 DUF63 -----------NE----G---YNTVNTLTYA-VILGLAALGVYK-I-LKRLG------IKYDKAFFRALIPYMILG-AFGRALTD-----A---G----- 76
WP_015849008_1 DUF63 -----------NQ----G---YNIVNTLTYA-IILGIAALVVYK-V-LKKLG------IKYDNAFFRALIPYMILG-AFGRALTD-----A---T----- 76
WP_004069276_1 DUF63 -----------NQ----G---YNVVNTLTYA-IILGLAALGVYK-V-LKKLG------IKYDNAFFRALIPYMILG-AFGRALTD-----A---T----- 76
WP_058946638_1 DUF63 -----------NQ----G---YNVVNTLTYA-IILGLAALGVYR-V-LKKLG------IEYDNAFFRALIPYMILG-AFGRALTD-----A---T----- 76
WP_042701551_1 DUF63 -----------NE----G---YNIVNTLTYA-IILGLAALGVYK-I-LKKLG------IKYDNAFFRALIPYMILG-AFGRALTD-----A---T----- 76
WP_055282692_1 DUF63 -----------NQ----G---YNIVNTLTYA-LILGLASLGVYK-I-LKKLN------IKYDNAFFRALMPYMILG-AFGRALTD-----A---T----- 76
WP_013906023_1 DUF63 -----------NT----G---YNVVNTLTYA-IILGVATLFVYR-V-LNRLG------IKYDNAFFRALIPYMVFG-AFTRALTD-----A---G----- 75
WP_011013158_1 DUF63 -----------NT----G---YNPVNTLVYA-IILGIATLLVYK-V-LKRLK------IEINNAFFRALIPYMIFG-AFTRALTD-----A---G----- 75
WP_014733166_1 DUF63 -----------NT----G---YNPVNTLVYA-IILGVATLVVYK-V-LKKLK------IEIDNAFFRALIPYMVFG-AFTRALTD-----A---G----- 75
WP_068322889_1 DUF63 -----------NT----G---YNPVNTLVYA-IILGIATLIVYK-V-LKRMK------IKIDNAFFRALIPYMVFG-AFTRALTD-----A---G----- 75
WP_068575625_1 DUF63 -----------NT----G---YNPVNTLVYA-IILGIATLIVYK-V-LKKMK------IKIDKAFFRALIPYMVFG-AFTRALTD-----A---G----- 75
WP_010885960_1 DUF63 -----------NT----G---YNPVNTLTYA-IILGIATILVYK-G-LKKLK------IKIDNAFFRALVPYMIFG-AFTRALTD-----A---S----- 75
WP_010867401_1 DUF63 -----------NT----G---YNPVNTLTYA-VILGLATILVYK-V-LKRLR------IKIDNAFFRALIPYMIFG-AFTRALTD-----A---G----- 75
WP_013747987_1 DUF63 -----------NT----G---YNPVNTLTYA-IILGLATLLVYK-V-LKKLR------IKIDNAFFRALIPYMIFG-AFTRALTD-----A---G----- 76
WP_068664049_1 DUF63 -----------NE----G---YNVVNTLTYA-VILGIGVVVVYR-A-LKRAG------ISTDRQFFKALLPYIALG-ALMRAMTD-----A---T----- 76
WP_011249739_1 DUF63 -----------NQ----G---YNPVNTLVYA-VILGIMVLLLYR-I-LKRMG------IRIDDRFFLSLMPYIVLG-PLMRAITD-----E---G----- 76
WP_062386854_1 DUF63 -----------NQ----G---YNPVNTLVYA-IILGVMVLLLYK-M-LKRMG------IKVDERFFLALVPYIVLG-PLMRAITD-----K---G----- 76
WP_050003781_1 DUF63 -----------NQ----G---YNAVNTLVYS-IILGIAVLLLHK-M-LKRMG------IKVDERFFVALMPYIILG-PLMRSITD-----I---G----- 76
WP_042690699_1 DUF63 -----------NQ----G---YNVVNTLVYA-LILGVAVLLLYK-M-LKRMR------VKVDERFFVALMPYIILG-PLMRSMTD-----I---G----- 76
WP_088885212_1 DUF63 -----------NQ----G---YNPVNTTVYA-IILGIAVLLLHR-M-LKRMK------IRVDERFFVALIPYIFLG-PLMRAMTD-----V---G----- 76
WP_010478519_1 DUF63 -----------DQ----G---YNPVNTTVYA-IILGIAVLLLYR-M-LKRMG------IRVDERFFVALMPYIFLG-PLMRAMTD-----V---G----- 76
WP_088858240_1 DUF63 -----------NQ----G---YNPVNTTVYA-IILGIAVLLLYR-L-LKKLG------IRVNETFFVSLMPYIFLG-PLMRAMTD-----V---G----- 76
WP_015859016_1 DUF63 -----------NQ----G---YNPVNTLVYA-IILGIAVLLLYR-F-LKRLG------VKVDERFFVALMPYIFLG-PLMRAMTD-----V---G----- 76
WP_014121940_1 DUF63 -----------NQ----G---YNPVNTLVYA-IILGIAVLLLYR-F-LKRLG------VKVDERFFVALMPYIFLG-PLMRAMTD-----V---G----- 76
WP_062373911_1 DUF63 -----------NQ----G---YNPVNTLTYA-VILGIAVLLLYR-F-LKRLN------VRVDERFFVALMPYIFLG-PLMRAITD-----I---G----- 76
WP_088882924_1 DUF63 -----------NQ----G---YNPVNTLVYA-LILGVAVILLYK-M-LKRMG------IRVDDRFFRALMPYVILG-PLMRSMTD-----T---G----- 76
WP_088862911_1 DUF63 -----------NQ----G---YNPVNTLTYA-VILGVAVILIYK-M-LKRMG------IRVDDRFFKALIPYIILG-PLMRSMTD-----V---G----- 76
WP_012572006_1 DUF63 -----------NE----G---YNIVNTFVYA-IILGIAVLLLYR-L-LKKMG------IKIDEHFFKALIPYMILG-PLMRAMTD-----V---G----- 76
WP_088854707_1 DUF63 -----------NQ----G---YNPVNTVTYA-IILGIAVILLYK-M-LKRMG------IRVDDRFFKALIPYIILG-PLMRGMTD-----V---G----- 76
WP_014789474_1 DUF63 -----------NQ----G---YNPVNTIAYA-IILGIAVLLLYR-M-LKRME------IKVDDRFFRALIPYIILG-PLMRSMTD-----V---G----- 76
WP_088180728_1 DUF63 -----------NQ----G---YNPVNTVVYA-IILGVAVILLYK-M-LKRMG------IKVDDRFFKALIPYIILG-PLMRSMTD-----V---G----- 76
WP_088864937_1 DUF63 -----------NQ----G---YNPVNTVVYA-IILGIAVILLYR-M-LKRME------IKVDDRFFRALIPYIILG-PLMRSMTD-----V---G----- 76
WP_055429686_1 DUF63 -----------NQ----G---YNPVNTVVYA-VILGVAVIILYR-M-LKRMG------IKVDERFFSALIPYIILG-PLMRSMTD-----V---G----- 76
WP_088865932_1 DUF63 -----------NQ----G---YNPVNTVVYA-IILGVAVILLYK-M-LKRMG------IKVDDRFFKALIPYIILG-PLMRSVTD-----V---G----- 76
WP_014013642_1 DUF63 -----------NQ----G---YNPVNTVVYA-IILGIAVLLLYR-M-LKRME------IRVDGRFFKALIPYIILG-PLMRSMTD-----V---G----- 76
WP_088856804_1 DUF63 -----------NQ----G---YNPVNTIVYA-IILGIAVILLYK-M-LKRMD------IRVEGRFFKALIPYIILG-PLMRSMTD-----V---G----- 76
WP_058939475_1 DUF63 -----------NQ----G---YNPVNTVVYA-IILGIAVLLLYK-M-LKRME------IRVDGRFFKALVPYIVLG-PLMRSMTD-----V---G----- 76
EHR77276_1 conserved -----DAGAAGDA----G---YSPENTTIYT-GSMLVSVIILQA-L-FRKAN------VPCDDKMTLALIAWVCLA-PVMRVLED-----A---D----- 131
OUV40025_1 hypotheti -----DAGAQGDA----G---YSYQNTAIYT-FGLLASVVVFQA-L-FRTLQ------LPADDKMMVALIAWVCLA-PILRVLED-----A---D----- 132
MBJ52984_1 hypotheti ------DATVGDS----G---YNTHNTILFA-VLLFAFVISFSG-I-FRIFN------LPVRYDSIIAFIPWVILA-ATIRVLED-----A---E----- 125
PDH23744_1 hypotheti -----------------GDSGYNVVNTMTYA-IVLGLFVVALSA-W-LRRLG------IDPTDTSLLALLPFITWA-VFGEVAEDAEMFGA---G----- 129
PDH25468_1 hypotheti -------ESTGDS----G---YNMVNTMTYG-IVLAMFAIALSG-W-LRHLG------IDGSDMTLLALLPFVLWA-ALGEVVED-----A---Q----- 124

WP_048201856_1 hypot -------------------------------------------------------------VKSIIHL-------------------------------- 80
WP_048150344_1 hypot --V------------------------------ASIFIP-----F--ATL--------LITLTSGIFQ-------------------------------- 85
AOV95360_1 hypotheti --I-----------------------------------A-----A--QKA-----------VAASITA-------------------------------- 69
EOD42420_1 Uncharact --Y-----------------------------TI----------K--TYL--------L--FTPLLEL-------------------------------- 93
AOV95164_1 hypotheti --V------------------------------LPFI-------I--RPL--------A--ITPIIYI-------------------------------- 106
EGQ40074_1 putative  ---------------------------------AVPFEA--------RLL--------L--VTPVIYF-------------------------------- 110
KYK23067_1 hypotheti ---------------------------------YFSD-P-----F--VYW--------F--ISPLIYV-------------------------------- 142
WP_084383883_1 DUF63 --------------------------------LLGDYSV-----W-------------F--ITPSIYF-------------------------------- 147
WP_049984677_1 DUF63 ---------------------------------LLGD-------Y--AVL--------F--ITPTIYL-------------------------------- 148
WP_103428047_1 hypot --------------------------------ILGDY----------AVL--------F--ITPSIYL-------------------------------- 147
WP_004048647_1 DUF63 --------------------------------LLGDYAV-----W-------------F--ITPSIYF-------------------------------- 147
WP_004594600_1 DUF63 --------------------------------LLGDYAV-----W-------------F--ITPSIYI-------------------------------- 147
WP_004594600_1 DUF63 --------------------------------LLGDYAV-----W-------------F--ITPSIYI-------------------------------- 147
WP_050050397_1 DUF63 --------------------------------LLGDYAV-----W-------------F--ITPSIYI-------------------------------- 147
WP_004594600_1 DUF63 --------------------------------LLGDYAV-----W-------------F--ITPSIYI-------------------------------- 147
WP_050050397_1 DUF63 --------------------------------LLGDYAV-----W-------------F--ITPSIYI-------------------------------- 147
WP_049983690_1 DUF63 --------------------------------LLGDY----------AVL--------F--ITPSIYF-------------------------------- 147
WP_103428162_1 hypot --L------------------------------LGDY-------A--VWF-----------ITPSIYF-------------------------------- 147
WP_009378268_1 DUF63 --------------------------------LLGDYAV-----W-------------F--ITPSVYF-------------------------------- 147
WP_015763176_1 DUF63 --D-----------------AAV--RAGVD--PAISY-P-----L--NSL--------L--VSPVIYF-------------------------------- 179
WP_018259238_1 DUF63 --D-----------------AAV--RAGVD--PAISY-P-----L--NSL--------L--VSPVIYF-------------------------------- 179
WP_004592615_1 DUF63 --D-----------------RAV--DAGVT--PIVEY-P-----L--SSL--------I--ISPVIYG-------------------------------- 173
WP_004518092_1 DUF63 --D-----------------RAV--DAGVS--PIVEY-P-----L--SSL--------I--ISPVIYG-------------------------------- 173
WP_014040450_1 DUF63 --D-----------------RAV--DAGVS--PIVEY-P-----L--SSL--------I--ISPVIYG-------------------------------- 173
WP_014040450_1 DUF63 --D-----------------RAV--DAGVS--PIVEY-P-----L--SSL--------I--ISPVIYG-------------------------------- 173
WP_008309018_1 DUF63 --D-----------------RAV--DAGVS--PIVEY-P-----L--SSL--------I--ISPVIYG-------------------------------- 173
WP_014040450_1 DUF63 --D-----------------RAV--DAGVS--PIVEY-P-----L--SSL--------I--ISPVIYG-------------------------------- 173
WP_014040450_1 DUF63 --D-----------------RAV--DAGVS--PIVEY-P-----L--SSL--------I--ISPVIYG-------------------------------- 173
WP_005534235_1 DUF63 --D-----------------RAV--DAGVT--PIVEY-P-----L--SSL--------I--ISPVIYG-------------------------------- 173
WP_004961153_1 DUF63 --D-----------------RAV--DAGVS--PIVEY-P-----L--SSL--------I--ISPVIYG-------------------------------- 173
WP_004961153_1 DUF63 --D-----------------RAV--DAGVS--PIVEY-P-----L--SSL--------I--ISPVIYG-------------------------------- 173
WP_101350151_1 hypot --D-----------------RAV--DAGVT--PIVEY-P-----L--SSL--------I--ISPVIYG-------------------------------- 173
WP_053968273_1 DUF63 --D-----------------RAV--DAGVT--PIVEY-P-----L--SSL--------I--ISPVIYG-------------------------------- 173
WP_058995687_1 DUF63 --D-----------------RAV--DAGVT--PIVEY-P-----L--SSL--------I--ISPVIYG-------------------------------- 173
WP_015790673_1 DUF63 --V------------------------EAGVEPVLSY-P-----F--NTL--------F--ISPLVYF-------------------------------- 174
WP_008524115_1 DUF63 --V------------------------DAGVEPVISY-P-----F--NTL--------F--ISPIIYF-------------------------------- 174
WP_075936143_1 DUF63 --D-----------------SI---PEGAA--QILAY-P-----W--NTL--------I--ISPIIYF-------------------------------- 167
WP_020446311_1 DUF63 --V-----------------ALL--SATGD--PAIPF-P-----W--TAA--------I--ISPFIYV-------------------------------- 169
```

```
WP_049898450_1 DUF63 --D-----------------AV---PDSAT--AAISY-P-----A--NAL--------I--ISPVIYL-------------------------------- 171
WP_006077061_1 DUF63 --D-----------------AV---PADAT--AAISY-P-----T--NTL--------I--ISPIIYV-------------------------------- 171
WP_049996700_1 DUF63 --D-----------------AV---PADAT--AAISY-P-----A--NAL--------I--ISPIIYL-------------------------------- 171
EMA38628_1 hypotheti --D-----------------VM---PASVA--VGIDY-P-----A--SAL--------I--ISPVIYV-------------------------------- 171
WP_049992561_1 DUF63 --D-----------------SVMGPNFGGE--ALIPY-P-----T--NTL--------L--ISPIIYF-------------------------------- 172
WP_010903025_1 DUF63 --N-----------------AAE--LFGTTGQPFLSY-P-----A--NTL--------I--ISPVIYV-------------------------------- 169
WP_009760947_1 DUF63 --N-----------------AAE--LFGNT-EPFLDY-P-----L--NAL--------I--ISPVIYF-------------------------------- 168
WP_059057097_1 DUF63 --D-----------------AAA--AAPGV-EQFLNF-P-----V--NTL--------I--ISPVIYF-------------------------------- 168
WP_058983522_1 DUF63 --D-----------------AAA--AAPGV-EQFLDY-P-----A--NTL--------I--ISPVIYF-------------------------------- 168
WP_071932813_1 DUF63 --D-----------------TLA--V--GA--GLIEF-P-----W--IAL--------F--ISPFIYV-------------------------------- 165
WP_050048584_1 DUF63 --D-----------------SVP--A--GV-EAAIQY-P-----W--NTL--------I--ISPLIYF-------------------------------- 165
WP_014051341_1 DUF63 --N-----------------AM------SD-GGGLTY-P-----L--NTL--------F--ISPVIYV-------------------------------- 168
WP_079233299_1 DUF63 --N-----------------AI------GD--SWFDY-P-----L--NTL--------L--ISPIIYF-------------------------------- 167
WP_053948572_1 DUF63 --N-----------------AM------GD--SWFDY-P-----L--NTL--------L--ISPIIYF-------------------------------- 167
WP_049980835_1 DUF63 --N-----------------AI------GD--SWFDY-P-----L--NTL--------L--ISPIIYV-------------------------------- 167
KPN31749_1 hypotheti --N-----------------AI------GD--SWFDY-P-----L--NTL--------L--ISPTIYV-------------------------------- 117
AGB16040_1 putative  --V-----------------AVH--EATGD--AALSL-P-----W--QAL--------I--ISPFIYV-------------------------------- 166
WP_007696755_1 DUF63 --V-----------------KVA--EATGD--AALSL-P-----W--QAL--------L--ISPFTYV-------------------------------- 166
WP_015322904_1 DUF63 --V-----------------AAY--RDTGE--LAIEL-P-----W--SGF--------L--ISPLIYV-------------------------------- 166
WP_076581869_1 DUF63 --V-----------------AAY--GATGD--LAIQL-P-----W--SGF--------L--ISPLIYF-------------------------------- 166
WP_005559622_1 DUF63 --V-----------------AAY--RDTGE--LAIEL-P-----W--SGF--------L--ISPLIYV-------------------------------- 166
WP_006067640_1 DUF63 --V-----------------AAY--RETGE--LAIQL-P-----W--SGF--------L--ISPLIYF-------------------------------- 161
WP_008164554_1 DUF63 --V-----------------ATY--RETGE--LAIQL-P-----W--SGF--------L--ISPLIYF-------------------------------- 166
WP_006088112_1 DUF63 --V-----------------AAY--RETGE--LAIQL-P-----W--SGF--------L--ISPLIYF-------------------------------- 166
WP_012943247_1 DUF63 --V-----------------AAY--RETGE--LMMEL-P-----W--IGF--------L--ISPLIYF-------------------------------- 166
WP_008895026_1 DUF63 --V-----------------AAY--RQTGE--LMMEL-P-----W--IGF--------L--ISPLIYF-------------------------------- 166
WP_098727043_1 DUF63 --A-----------------AAF--VYTGE--MAIDL-P-----W--SGF--------I--ISPLIYF-------------------------------- 166
WP_049990861_1 DUF63 --A-----------------TAY--QATNE--MAIQL-P-----W--SGF--------I--ISPLIYF-------------------------------- 164
WP_008013001_1 DUF63 --A-----------------ASY--AYSGE--MAIQL-P-----W--SGF--------I--ISPLIYF-------------------------------- 166
WP_006180566_1 DUF63 --A-----------------AAY--EYAGE--MAIQL-P-----W--SGF--------I--ISPLIYF-------------------------------- 166
WP_006650865_1 DUF63 --A-----------------AAY--EYTGE--MAIQL-P-----W--SGF--------I--ISPLIYF-------------------------------- 166
WP_066301295_1 DUF63 --A-----------------AAY--EYSGE--MAIQL-P-----W--SGF--------I--ISPLIYF-------------------------------- 166
WP_049966804_1 DUF63 --A-----------------AAY--EYTGE--MAIQL-P-----W--SGF--------I--ISPLIYF-------------------------------- 166
WP_076145380_1 DUF63 --A-----------------AAY--EYSGE--MAIQL-P-----W--SGF--------I--ISPLIYF-------------------------------- 166
WP_097378845_1 DUF63 --A-----------------TAF--AATGE--MSIPL-P-----W--SGF--------I--ISPLIYF-------------------------------- 166
WP_008452261_1 DUF63 --A-----------------TAF--EATGE--MAIHL-P-----W--SGF--------I--ISPLIYF-------------------------------- 166
WP_008452261_1 DUF63 --A-----------------TAF--EATGE--MAIHL-P-----W--SGF--------I--ISPLIYF-------------------------------- 166
WP_006432643_1 DUF63 --A-----------------TAF--EATGE--MAIQL-P-----W--SGF--------I--ISPLIYF-------------------------------- 166
WP_008452261_1 DUF63 --A-----------------TAF--EATGE--MAIHL-P-----W--SGF--------I--ISPLIYF-------------------------------- 166
WP_007109662_1 DUF63 --A-----------------TAF--EATGE--MAIQL-P-----W--SGF--------I--ISPLIYF-------------------------------- 166
WP_049952821_1 DUF63 --V-----------------ATY--TATGE--LPIQM-P-----W--NGI--------L--ISPLIYF-------------------------------- 166
WP_086889541_1 DUF63 --V-----------------AAF--RDTGE--LAIEL-P-----W--SGF--------L--ISPLIYF-------------------------------- 169
WP_005580009_1 DUF63 --V-----------------AAF--RHTGE--LAIEL-P-----W--SGF--------L--ISPLIYV-------------------------------- 169
WP_049927356_1 DUF63 --V-----------------AAY--RETGE--LAIEL-P-----W--SGL--------L--ISPLIYF-------------------------------- 166
WP_087714455_1 DUF63 --V-----------------AAY--RDTGE--LAIEL-P-----W--SGF--------L--ISPLIYF-------------------------------- 166
WP_013878438_1 DUF63 --A-----------------AAY--RATGT--LAIDL-P-----W--SGF--------I--ISPLIYF-------------------------------- 166
WP_049921111_1 DUF63 --V-----------------AAY--RVTGE--LAIQL-P-----W--VGI--------L--ISPLIYF-------------------------------- 166
WP_007142789_1 DUF63 --V-----------------AAF--RYTGE--LAIDL-P-----W--VGF--------L--ISPLIYF-------------------------------- 165
WP_006651941_1 DUF63 --V-----------------AAY--RETGE--LAIQL-P-----W--SGL--------L--ISPLIYF-------------------------------- 166
WP_004216437_1 DUF63 --V-----------------AAY--RETGE--LAIQL-P-----W--SGL--------L--ISPLIYF-------------------------------- 166
WP_071402513_1 DUF63 --V-----------------AAY--RETGE--LAIQL-P-----W--SGL--------L--ISPLIYF-------------------------------- 166
WP_006666462_1 DUF63 --V-----------------AAY--RATGE--LPIQL-P-----W--SGI--------I--ISPLIYF-------------------------------- 166
WP_049904535_1 DUF63 --V-----------------AAY--RATGE--LPIQL-P-----W--SGI--------I--ISPLIYF-------------------------------- 166
WP_006824405_1 DUF63 --V-----------------AAY--RATGE--LPIQL-P-----W--SGI--------I--ISPLIYF-------------------------------- 166
WP_011323802_1 DUF63 --V-----------------AFR--GEAVG--MLIPY-P-----A--VAL--------I--ISPFIYF-------------------------------- 167
WP_015409719_1 DUF63 --V-----------------TFL--RAGVD--TLIPY-P-----P--VAL--------I--ISPFIYF-------------------------------- 168
WP_006883947_1 DUF63 --D-----------------AAV--AAGTD--PVISY-P-----L--NVL--------F--ISPVIYV-------------------------------- 167
WP_077207545_1 DUF63 --D-----------------AV---PEGVS--QAIAY-P-----W--NSL--------I--ISPVIYF-------------------------------- 170
ESS12903_1 putative  --V-----------------ELL--REGV---SLLPL-P-----W--TAL--------I--ISPFIYF-------------------------------- 170
WP_008417361_1 DUF63 --I-----------------ATL--RATGE--PAIAF-P-----L--SGL--------F--ISPFIYF-------------------------------- 169
WP_066381608_1 DUF63 --V-----------------ALL--RTTGE--PAISF-P-----L--SGL--------F--ISPFIYF-------------------------------- 164
WP_049947642_1 DUF63 --D-----------------AM-----AEA--ALFPY-P-----W--NTL--------L--ISPVIYF-------------------------------- 170
ESS05953_1 putative  --D-----------------TG----------ELLTY-P-----L--NTL--------V--ISPVIYV-------------------------------- 164
WP_096390051_1 DUF63 --V-----------------AAL--DAGGQ--HLIPA-P-----W--TAL--------F--ISPFIYV-------------------------------- 176
WP_021072749_1 DUF63 --V-----------------AAL--DAGGQ--HLIPA-P-----W--TAL--------F--ISPFIYV-------------------------------- 176
WP_049982791_1 DUF63 --V-----------------AAM--DAGVE--PLIPF-P-----A--SAL--------L--ISPFIYF-------------------------------- 174
WP_008585990_1 DUF63 --V-----------------AAT--AAGVD--PLVPF-P-----A--SAV--------L--ISPFIYF-------------------------------- 176
WP_006628721_1 DUF63 --V-----------------AAT--AAGVE--PLISF-P-----A--SAL--------L--ISPFIYF-------------------------------- 174
WP_006113218_1 DUF63 --V-----------------AAT--AAGVE--PLIPF-P-----A--SAV--------L--ISPFIYF-------------------------------- 175
WP_049930034_1 DUF63 --V-----------------AAT--AAGAE--PLIPF-P-----A--SAV--------L--ISPFIYF-------------------------------- 176
WP_049906047_1 DUF63 --V-----------------AAT--DAGVE--PLIAF-P-----A--SAI--------L--ISPFIYF-------------------------------- 176
WP_044965494_1 DUF63 --V-----------------AAT--DAGVE--PLIAF-P-----A--SAV--------L--ISPFIYF-------------------------------- 176
WP_096393195_1 DUF63 --V-----------------AAT--DAGVE--PLIAF-P-----A--SAV--------L--ISPFIYF-------------------------------- 176
WP_049908400_1 DUF63 --V-----------------AAT--DAGVE--PLISF-P-----A--SAV--------L--ISPFIYF-------------------------------- 176
WP_049908585_1 DUF63 --V-----------------AAT--DAGVE--PLIPF-P-----A--SAV--------L--ISPFIYF-------------------------------- 176
WP_049902631_1 DUF63 --V-----------------AAT--DAGVE--PLIPF-P-----A--SAV--------L--ISPFIYF-------------------------------- 176
WP_049983668_1 DUF63 --V-----------------AAT--AAGVE--PLIAF-P-----A--SAV--------L--ISPFIYF-------------------------------- 175
WP_053772267_1 DUF63 --V-----------------AAT--AAGVD--PLIAF-P-----A--SAV--------L--ISPFIYF-------------------------------- 175
WP_007999151_1 DUF63 --I-----------------AAL--RAGVE--PLIPF-P-----A--SAL--------L--ISPFIYF-------------------------------- 174
WP_015909917_1 DUF63 --V-----------------AAM--RAGVE--PLIPF-P-----A--SAL--------L--ISPFIYF-------------------------------- 175
WP_004050594_1 DUF63 --V-----------------AAM--RAGVE--PLIPF-P-----V--SAV--------L--ISPFIYF-------------------------------- 175
WP_095636035_1 DUF63 --I-----------------AAM--RAGVE--PLIPF-P-----A--SAL--------L--ISPFIYF-------------------------------- 175
WP_008003811_1 DUF63 --V-----------------AAM--RAGVE--PLIPF-P-----A--SAL--------L--ISPFIYF-------------------------------- 175
WP_066416034_1 DUF63 --I-----------------AAI--RAGVD--PLIPF-P-----W--SAL--------I--ISPFIYF-------------------------------- 176
ESS03170_1 putative  --I-----------------SAL--DAGAD--PLISF-P-----L--SAL--------I--ISPFIYF-------------------------------- 175
WP_089671383_1 DUF63 --I---------------------SAMRFGAEPLIDF-P-----W--SAV--------I--ISPFIYF-------------------------------- 174
ERH07656_1 putative  --I---------------------AAITAGAEPLIPF-P-----W--SAL--------I--ISPFIYL-------------------------------- 174
ESS10096_1 putative  --I---------------------AAITAGAEPLIPF-P-----W--SAL--------I--ISPFIYL-------------------------------- 174
ESS07824_1 putative  --------------------------------------------I--AAL--------DAGVEPLISF-------------------------------- 161
ERH05689_1 putative  --I---------------------AALDAGVEPLISF-P-----L--SAA--------I--ISPFIYG-------------------------------- 174
ERH02266_1 putative  --I---------------------AALDAGVEPLISF-P-----L--SAA--------I--ISPFIYG-------------------------------- 174
ESS07825_1 putative  --L-----------------------------------T-----T--SYA-------------------------------------------------- 21
WP_049970017_1 DUF63 --N-----------------AL---QQVGG--GAIPF-P-----W--NTL--------I--ISPIIYF-------------------------------- 168
WP_007979289_1 DUF63 --D-----------------AL---RQTGA--EAIPF-P-----W--NTL--------I--ISPIIYF-------------------------------- 168
WP_007977745_1 DUF63 --D-----------------AL---GQVGQ--SAIPF-P-----W--NTL--------I--ISPIIYF-------------------------------- 168
```

```
WP_014556445_1 DUF63 --D-----------------AV----PSAD--ALISY-P-----L--NTL--------V--ISPIIYF-------------------------------- 169
WP_049935775_1 DUF63 --D-----------------AA------AG--SPIGY-P-----L--NTL--------I--ISPVIYF-------------------------------- 167
WP_008325149_1 DUF63 --D-----------------TP----GVAD--ALITY-P-----V--NTL--------I--ISPVIYV-------------------------------- 169
WP_007543631_1 DUF63 --D-----------------TP----GVAD--ALITY-P-----V--NTL--------I--ISPVIYV-------------------------------- 169
WP_049905041_1 DUF63 --D-----------------SP----GVAD--ALISY-P-----L--NTL--------V--ISPVIYV-------------------------------- 169
WP_049913563_1 DUF63 --D-----------------SP----GVAD--ALISY-P-----L--NTL--------V--ISPVIYV-------------------------------- 169
WP_049967851_1 DUF63 --D-----------------SP----GVAD--ALISY-P-----L--NTL--------V--ISPVIYV-------------------------------- 169
WP_049914905_1 DUF63 --D-----------------AP----GVAD--ALITY-P-----L--NTL--------V--ISPVIYV-------------------------------- 169
WP_049916430_1 DUF63 --D-----------------AP----GVAD--ALITY-P-----L--NTL--------V--ISPVIYV-------------------------------- 169
WP_049896892_1 DUF63 --D-----------------AP----GVAD--ALITY-P-----L--NTL--------V--ISPVIYV-------------------------------- 169
WP_049896892_1 DUF63 --D-----------------AP----GVAD--ALITY-P-----L--NTL--------V--ISPVIYV-------------------------------- 169
WP_049896892_1 DUF63 --D-----------------AP----GVAD--ALITY-P-----L--NTL--------V--ISPVIYV-------------------------------- 169
WP_058828253_1 DUF63 --D-----------------SP----GVAD--ALISY-P-----L--NTL--------V--ISPVIYV-------------------------------- 169
WP_058568959_1 DUF63 --D-----------------SP----GVAD--ALISY-P-----L--NTL--------V--ISPVIYV-------------------------------- 169
WP_049917947_1 DUF63 --D-----------------AP----GVAD--ALITY-P-----L--NTL--------V--ISPVIYV-------------------------------- 169
WP_049920348_1 DUF63 --D-----------------AP----GVAD--ALITY-P-----L--NTL--------V--ISPVIYV-------------------------------- 169
WP_089777545_1 DUF63 --D-----------------AP----GVAD--ALITY-P-----L--NTL--------V--ISPVIYV-------------------------------- 169
WP_008320633_1 DUF63 --D-----------------TP----GVAD--ALITY-P-----L--NTL--------V--ISPVIYV-------------------------------- 169
WP_004060355_1 DUF63 --D-----------------AP----GVAD--ALITY-P-----V--NTL--------V--ISPVIYV-------------------------------- 169
WP_103426078_1 hypot --D-----------------TP----GTAD--ALISY-P-----L--NTL--------F--ISPVIYF-------------------------------- 169
WP_009367433_1 DUF63 --D-----------------TG----G-AE--ALISY-P-----W--NAL--------V--ISPVIYV-------------------------------- 168
WP_013440552_1 DUF63 --D-----------------TA----AVAE--SIITY-P-----L--NTL--------V--ISPVIYF-------------------------------- 169
WP_049916626_1 DUF63 --D-----------------TP----GAAE--ALISY-P-----W--NAL--------V--ISPVIYF-------------------------------- 169
WP_058582837_1 DUF63 --I-----------------AAM--RAGVD--PAIPF-P-----W--SAL--------I--ISPFIYF-------------------------------- 171
WP_101298124_1 hypot --I-----------------AAM--RAGVD--PAIPF-P-----W--SAL--------I--ISPFIYF-------------------------------- 171
PIN95153_1 hypotheti --------------------------------------P-----K----------------ISSVIVF-------------------------------- 50
PIU22108_1 hypotheti --Y----------------------STLQWFSRSANPLE-----M--GFY--------F--ITPGVYL-------------------------------- 100
AAR39201_1 NEQ352    ------------------------------------LLP-----N--TFF--------T--VTPGVWI-------------------------------- 68
OIR14399_1 hypotheti ---------------------------------HFEE-P-----L--QYF--------F--ISPLIYL-------------------------------- 125
OIR20963_1 hypotheti ---------------------------------AFEP-P-----I--QYI--------M--ISPLIYG-------------------------------- 130
OIR22371_1 hypotheti --D------------------------------TFEP-P-----I--QYF--------F--ISPLIYG-------------------------------- 141
EGQ43935_1 putative  ---------------------------------VKAF-------N--TIL--------L--ETPFIYG-------------------------------- 95
MAG21679_1 hypotheti ---------------------------------LIPTQSANPLEL--GFW--------F--VTPGIWI-------------------------------- 100
PIN85618_1 hypotheti CNI---------------------------------LDP--------GFF--------T--FTPGIWF-------------------------------- 100
PIN99249_1 hypotheti --------------------------YSAVYFSLFER-STDLFSF--GFY--------T--VSPGIYI-------------------------------- 104
AJF59838_1 hypotheti -------------------------------RSCNPFEP--------GFL--------T--VTPGIYV-------------------------------- 100
WP_042682145_1 DUF63 --V---------------------------------LKP--------NPL--------I--LTPGIFF-------------------------------- 94
WP_013468033_1 DUF63 --V---------------------------------LKP--------NPL--------I--LTPGIFF-------------------------------- 94
WP_055281581_1 DUF63 --I---------------------------------LKP--------HPL--------I--LTPGIFF-------------------------------- 93
WP_048160382_1 DUF63 --V---------------------------------LEP--------HPL--------I--LTPGIFF-------------------------------- 93
WP_042701216_1 DUF63 --V---------------------------------LKP--------HPL--------I--LTPGIFF-------------------------------- 93
WP_004068659_1 DUF63 --V---------------------------------LKP--------HPL--------I--LTPGIFF-------------------------------- 93
WP_058946665_1 DUF63 --V---------------------------------LKP--------HPL--------I--LTPGIFF-------------------------------- 93
WP_014835499_1 DUF63 ---------------------------------VVEPNP-----W-------------I--LTPGIFF-------------------------------- 93
WP_010884243_1 DUF63 ---------------------------------VLAPNP-----W-------------I--LTPGIFF-------------------------------- 96
WP_013748129_1 DUF63 ---------------------------------VLNPNP-----W-------------I--LTPGIFF-------------------------------- 92
WP_010867251_1 DUF63 ---------------------------------ILKPNP-----W-------------I--LTPGIFF-------------------------------- 92
WP_014733359_1 DUF63 ---------------------------------VLKPNP-----W-------------I--LTPGIFF-------------------------------- 93
WP_068319891_1 DUF63 ---------------------------------VLKPNP-----W-------------I--LTPGIFF-------------------------------- 93
WP_068576049_1 DUF63 ---------------------------------VLKPNP-----W-------------I--LTPGIFF-------------------------------- 93
WP_014013692_1 DUF63 --I---------------------------------L-P-----Q--NPL--------I--LTPGIFF-------------------------------- 98
WP_014789520_1 DUF63 --I---------------------------------L-P-----Q--NPL--------I--LTPGIFF-------------------------------- 98
WP_088180676_1 DUF63 --I---------------------------------L-P-----Q--NPL--------I--LTPGIFF-------------------------------- 98
WP_088864885_1 DUF63 --I---------------------------------L-P-----Q--NPL--------I--LTPGIFF-------------------------------- 98
WP_088856755_1 DUF63 --I---------------------------------L-P-----E--NPL--------I--LTPGIFF-------------------------------- 98
WP_088865981_1 DUF63 --I---------------------------------L-P-----Q--NPL--------I--LTPGIFF-------------------------------- 98
WP_013906334_1 DUF63 ---------------------------------VLEPNP-----W-------------I--LTPGIFF-------------------------------- 94
WP_088862658_1 DUF63 --I---------------------------------L-P-----E--NPL--------I--LTPGIVF-------------------------------- 97
WP_088882956_1 DUF63 --V---------------------------------L-P-----K--HPL--------L--LTPGVFF-------------------------------- 97
WP_012571938_1 DUF63 --I---------------------------------L-P-----K--NPL--------I--LTPGIFF-------------------------------- 94
WP_068663933_1 DUF63 --V---------------------------------L-P-----K--NPL--------I--LTPGIFF-------------------------------- 94
WP_074631153_1 DUF63 --V---------------------------------L-P-----K--NPL--------I--LTPGIFF-------------------------------- 94
WP_058939074_1 DUF63 --V---------------------------------L-P-----K--NPL--------I--LTPGIFF-------------------------------- 98
WP_088854681_1 DUF63 --V---------------------------------L-P-----K--NPL--------I--LTPGVFF-------------------------------- 98
WP_088885426_1 DUF63 ---------------------------------VLKPNP-----W-------------I--LTPGIFF-------------------------------- 94
WP_011249694_1 DUF63 --V---------------------------------L-P-----Q--HPL--------I--LTPGIFF-------------------------------- 96
WP_062386972_1 DUF63 --V---------------------------------L-P-----Q--HPL--------I--LTPGIFF-------------------------------- 96
WP_010478887_1 DUF63 --V---------------------------------LKP--------NFW--------I--LTPGIFF-------------------------------- 94
WP_088858605_1 DUF63 ---------------------------------VLEPNP-----W-------------I--LTPGIFF-------------------------------- 94
WP_042690028_1 DUF63 --I---------------------------------L-P-----K--NPL--------I--LTPGIFF-------------------------------- 96
WP_048150762_1 DUF63 --V---------------------------------L-P-----E--NPL--------I--LTPGIFF-------------------------------- 94
WP_048811076_1 DUF63 --V---------------------------------L-P-----Q--HPL--------L--LTPGIFF-------------------------------- 94
WP_050003256_1 DUF63 --V---------------------------------L-P-----K--HPL--------I--LTPGIFF-------------------------------- 96
WP_062372100_1 DUF63 --V---------------------------------L-P-----Q--HPL--------I--LTPGIFF-------------------------------- 94
WP_048165284_1 DUF63 --------------------------------KVLNPNP-----L-------------I--LTPGIFF-------------------------------- 96
WP_048148750_1 DUF63 --------------------------------KVLSPNP-----W-------------I--LTPGIFF-------------------------------- 96
OYT53462_1 hypotheti ---------------------------------FFER-S-----Q--IWN-----------ITPGVYL-------------------------------- 74
KYC51152_1 hypotheti --H------------------------------YERS-------K--FWN-----------ITPGVYV-------------------------------- 74
KYC46003_1 hypotheti -----------------------------------HYER-----S--KFW--------N--ITPGVYV-------------------------------- 74
KYC48643_1 hypotheti -----------------------------------HYER-----S--KFW--------N--ITPGVYV-------------------------------- 74
KYC55321_1 hypotheti -KL-----------------------------------------W-------------N--ITPGVYV-------------------------------- 74
KYC57927_1 hypotheti SKL-----------------------------------------W-------------N--ITPGVYV-------------------------------- 74
KYC57171_1 hypotheti SKL-----------------------------------------W-------------N--ITPGVYV-------------------------------- 74
OIO20701_1 hypotheti --VMGKYVMENGGSAVAGAYSAILSSHLYDYKQVSNP-------LDIVFA--------L--HSPGIYF-------------------------------- 126
PIT83986_1 hypotheti LLVYSASGAQGGNAGAYLGSGCAWFFGGIARWLYPEILNSGIFAY--SFL--------T--VSPGMYL-------------------------------- 143
OIO24760_1 hypotheti VTV-----------------------------SGFTFYP-------------------F--VTPVVYV-------------------------------- 103
OIO24701_1 hypotheti ------------------------------------TLP-----RAVDFF--GLELFPF--VTPGIYL-------------------------------- 102
OIO27021_1 hypotheti --I---------------------------------L-P-----R--TVYLAGVELHPF--ISPGIYF-------------------------------- 99
OIO26762_1 hypotheti --I---------------------------LPRQIELLG-----WTLYPF-----------VTPGIYF-------------------------------- 100
PIN95811_1 hypotheti ---------------------------------ILPR-QIELLGW--TLY-------PF--VTPGIYF-------------------------------- 100
PIO01637_1 hypotheti ---------------------------------ILPR-QIELLGW--TLY-------PF--VTPGIYF-------------------------------- 100
PIO02820_1 hypotheti --I---------------------------------L-P-----R--TVYLAGVELHPF--ISPGIYF-------------------------------- 99
PJD01038_1 hypotheti --I------------------------------L----P-----R--TVYLAGVELHPF--ISPGIYF-------------------------------- 99
PIZ91366_1 hypotheti ---------------------------------ILPR-QIELLGW--TLY-------PF--VTPGIYF-------------------------------- 100
WP_013100570_1 DUF63 ------------------------------------YIE-----R--SFL--------T--ITPGIVF-------------------------------- 93
```

```
WP_004590770_1 DUF63 ------------------------------------YIE-----R--SFL--------T--ITPGIVF------------------------------- 93
WP_048196979_1 DUF63 ------------------------------------YIE-----R--SFL--------T--ITPGIIF------------------------------- 93
WP_015791538_1 DUF63 ------------------------------------HIE-----R--SFL--------T--ITPGIVF------------------------------- 93
WP_048202292_1 DUF63 ------------------------------------YIE-----R--SFL--------T--ITPGIVF------------------------------- 93
WP_012981280_1 DUF63 ------------------------------------YIE-----R--SFL--------T--ITPGIVF------------------------------- 93
WP_064496496_1 DUF63 ------------------------------------YIE-----R--SFL--------T--ITPGIVF------------------------------- 93
WP_011972741_1 DUF63 --V------------------------------L----P-----H--TFY--------T--VTPGIVV------------------------------- 95
WP_013798289_1 DUF63 ---------------------------------VIER----------SFF--------T--ITPGIVI------------------------------- 93
WP_013181100_1 DUF63 ---------------------------------VIER----------LFF--------T--VTPGIVI------------------------------- 95
WP_013867628_1 DUF63 ------------------------------------YIP-----H--SYY--------T--VTPGIVI------------------------------- 95
WP_018153391_1 DUF63 --V------------------------------I----P-----R--LYY--------T--VTPGIVV------------------------------- 95
WP_011170272_1 DUF63 --L---------------------------------F-P-----R--LYY--------T--VTPGIVV------------------------------- 95
WP_012066151_1 DUF63 ------------------------------------FFP-----R--LYY--------T--VTPGIVV------------------------------- 95
ODS43013_1 hypotheti --L-----------------------GIYSSAGVYPENP-----W--AFF--------V--VSPGIFL------------------------------- 99
OIQ06085_1 hypotheti --I---------------------------------Y-PHRPDVL--AFL--------F--IAPGIYF------------------------------- 96
PIN67069_1 hypotheti --I---------------------------------Y-PHRPDVL--AFL--------F--IAPGIYF------------------------------- 96
PIV28099_1 hypotheti --I---------------------------------Y-PHRPDVL--AFL--------F--IAPGIYF------------------------------- 96
PJC13070_1 hypotheti --I---------------------------------Y-PHRPDVL--AFL--------F--IAPGIYF------------------------------- 96
PIZ29927_1 hypotheti --I---------------------------------Y-PHRPDVL--AFL--------F--IAPGIYF------------------------------- 96
PKP60688_1 hypotheti --I---------------------------------Y-PNRPDVL--AFF--------F--IAPGIYF------------------------------- 96
WP_012956954_1 DUF63 --V---------------------------------Y-P-----K--TVF--------L--ITPGLYI------------------------------- 89
WP_067148430_1 DUF63 --V---------------------------------Y-P-----K--TVF--------L--ITPGLYI------------------------------- 89
WP_080460538_1 DUF63 --I---------------------------------Y-P-----Y--NWF--------L--ITPGIYF------------------------------- 88
WP_010877090_1 DUF63 --I---------------------------------Y-P-----L--THL--------L--VTPGIYI------------------------------- 89
WP_013294901_1 DUF63 --I---------------------------------Y-P-----L--THL--------L--VTPGIYV------------------------------- 89
WP_010877090_1 DUF63 --I---------------------------------Y-P-----L--THL--------L--VTPGIYI------------------------------- 89
BAZ99473_1 hypotheti --I---------------------------------Y-P-----L--THL--------L--VTPGIYI------------------------------- 89
PKL66404_1 hypotheti ---------------------------------LYPLIL-----W-------------L--VTPGIYV------------------------------- 97
WP_023991171_1 DUF63 --I---------------------------------Y-P-----L--TYA--------L--VTPGIYI------------------------------- 90
WP_048072898_1 DUF63 --I---------------------------------Y-P-----L--TYA--------L--VTPGIYI------------------------------- 90
WP_100905180_1 hypot --I---------------------------------Y-P-----L--TYL--------L--VTPGIYL------------------------------- 90
WP_100907231_1 hypot --I---------------------------------Y-P-----L--TYL--------L--VTPGIYL------------------------------- 90
WP_100907231_1 hypot --I---------------------------------Y-P-----L--TYL--------L--VTPGIYL------------------------------- 90
WP_048081925_1 DUF63 --I---------------------------------Y-P-----L--VLC--------L--VTPGIYI------------------------------- 89
WP_048081925_1 DUF63 --I---------------------------------Y-P-----L--VLC--------L--VTPGIYI------------------------------- 89
WP_069585525_1 DUF63 --I---------------------------------Y-P-----L--VLW--------L--VTPGIYI------------------------------- 89
WP_069585525_1 DUF63 --I---------------------------------Y-P-----L--VLW--------L--VTPGIYI------------------------------- 89
WP_013643671_1 DUF63 --I---------------------------------Y-P-----L--TYL--------L--VTPGIYL------------------------------- 89
WP_013824571_1 DUF63 --I---------------------------------Y-P-----L--TYI--------L--VTPGIYL------------------------------- 89
WP_048192110_1 DUF63 --I---------------------------------Y-P-----L--TYI--------L--VTPGIYI------------------------------- 93
WP_081810122_1 DUF63 ---------------------------------ILNP-P-----V--KYF--------F--ITPLIYF------------------------------- 117
WP_097298965_1 DUF63 ---------------------------------ILKP-P-----V--KYF--------F--ITPLIFF------------------------------- 93
PIO00292_1 hypotheti --I------------------------------L----P-----R--TFF--------L--VTPSIYL------------------------------- 94
WP_011833679_1 DUF63 ---------------------------------MVPE-P-----W--WIL--------F--VTPQVYF------------------------------- 89
WP_042698332_1 DUF63 --M--------------------------------VPGP-----W--WIL--------F--VTPQVYI------------------------------- 89
WP_042698332_1 DUF63 ---------------------------------MVPG-P-----W--WIL--------F--VTPQVYI------------------------------- 89
WP_011448164_1 DUF63 --Y------------------------------ITSD-------L--HVI--------F--TTPLIFF------------------------------- 92
WP_007314214_1 DUF63 --I------------------------------VPVG-------W--NYL--------L--ITPLIYF------------------------------- 97
WP_042705632_1 DUF63 ---------------------------------FIPF-P-----Y--NVL--------F--ITPLIYF------------------------------- 92
WP_013329717_1 DUF63 ---------------------------------IIPY-P-----Y--YVL--------L--ITPLIYF------------------------------- 92
WP_004077722_1 DUF63 --I-------------------------------IPY-P-----W--YIL--------L--ITPVIFF------------------------------- 92
WP_048150927_1 DUF63 ---------------------------------IIPY-P-----Y--YVL--------L--ITPLIYF------------------------------- 92
WP_012107531_1 DUF63 --M------------------------------VGGD-------W--QYL--------I--VTPPIYF------------------------------- 98
PKL70081_1 hypotheti --M------------------------------ITSD-------L--QFL--------L--VTPPIYV------------------------------- 92
WP_015285007_1 DUF63 --M------------------------------ITGD-------F--KYL--------L--ITPLIYF------------------------------- 98
PKL64570_1 hypotheti ---------------------------------MITG-P-----L--KFL--------L--ITPLIYF------------------------------- 98
WP_015286404_1 DUF63 --F------------------------------ITSD-------I--RFL--------L--ITPLIYF------------------------------- 92
WP_012617618_1 DUF63 --M------------------------------ITSD-------L--QFL--------L--VTPLIFF------------------------------- 92
WP_014867374_1 DUF63 --M------------------------------ITSD-------F--RFL--------L--ITPLIFF------------------------------- 92
CVK33108_1 conserved --M------------------------------ITSD-------L--RFL--------L--ITPLIFF------------------------------- 92
WP_066956331_1 DUF63 --M------------------------------ITSD-------L--RFL--------L--ITPLIFF------------------------------- 92
WP_011844879_1 DUF63 ---------------------------------MITS-D-----L--RFL--------L--ITPLIFF------------------------------- 92
WP_048181868_1 DUF63 --------------------------------MITSD-------L--RFL--------L--ITPLIFF------------------------------- 92
WP_067073014_1 DUF63 --M------------------------------IVSD-------L--RFL--------L--ITPLIFF------------------------------- 92
PKL62929_1 hypotheti --M------------------------------ITSD-------L--HFL--------L--ITPLIFF------------------------------- 92
WP_004039890_1 DUF63 --M------------------------------IASD-------L--RIL--------L--ITPIIFF------------------------------- 92
WP_067049225_1 DUF63 --M------------------------------ITSD-------A--HIL--------L--ITPIIFF------------------------------- 92
KYK37884_1 hypotheti --V-----------------------------------S-----E--SYI--------L--VTPLIFF------------------------------- 88
KYK28019_1 hypotheti --V-----------------------------------S-----E--SYI--------L--VTPLIFF------------------------------- 88
OYT57835_1 hypotheti --I-----------------------------------L-----K--SAL--------F--VTPLIYF------------------------------- 92
OYT33350_1 hypotheti ---------------------------------FLNP-P-----I--SYF--------F--MTPLIYV------------------------------- 95
WP_012964745_1 DUF63 ---------------------------------FLQP-P-----I--SYV--------F--MSPMIYV------------------------------- 92
WP_048091573_1 DUF63 ---------------------------------FVKP-P-----F--SYI--------L--MTPFIYI------------------------------- 92
WP_048096399_1 DUF63 ---------------------------------FVQP-P-----Y--SYL--------L--MTPFIYI------------------------------- 92
WP_012940099_1 DUF63 ---------------------------------FLKP-P-----L--SYF--------F--MSPFIYV------------------------------- 91
WP_010877969_1 DUF63 ---------------------------------FLTP-P-----I--SYI--------F--MTPFIYI------------------------------- 104
WP_013682839_1 DUF63 ---------------------------------FLNP-P-----V--SYF--------F--MTPLIYV------------------------------- 92
WP_015591141_1 DUF63 ---------------------------------FLKP-P-----I--SYF--------F--MTPFIYI------------------------------- 92
WP_012035887_1 DUF63 --------------------------------QLVQP-P-----L--SYM--------F--ITPFVYV------------------------------- 99
WP_014404626_1 DUF63 --------------------------------HLLSP-P-----L--SYM--------F--ITPLVYV------------------------------- 93
BAI60262_1 conserved --V-----------------------------HLLSP-P-----L--SYM--------F--ITPLVYV------------------------------- 93
WP_042684156_1 DUF63 ---------------------------------IVLP-P-----L--SYL--------L--ITPLIYF------------------------------- 101
OFV68024_1 membrane  ---------------------------------LINP-P-----L--SYL--------L--VTPLIFF------------------------------- 59
WP_013720253_1 DUF63 --E------------------------------IVQP-P-----W--SYL--------L--ITPLIFF------------------------------- 94
WP_014587756_1 DUF63 ---------------------------------LVAA-P-----L--KYL--------L--ITPLIYF------------------------------- 91
ABK14947_1 Protein o ---------------------------------LIDP-P-----L--SYL--------L--ITPLIYI------------------------------- 98
OKY79140_1 putative  --I-------------------------------FSP-P-----V--KYL--------F--ITPLIYF------------------------------- 95
WP_086637003_1 DUF63 --V-------------------------------VHE-P-----L--NYL--------L--ITPLIYF------------------------------- 95
WP_048089097_1 DUF63 ---------------------------------IFTP-P-----L--RYL--------F--ITPIIYF------------------------------- 96
WP_097298272_1 DUF63 ---------------------------------IFTP-P-----L--KYL--------F--VTPIIYF------------------------------- 97
WP_013897788_1 DUF63 --V-------------------------------VDP-P-----L--SYL--------L--ITPNIYF------------------------------- 98
WP_015323982_1 DUF63 ---------------------------------AIQQ-P-----F--NLL--------L--ITPIIYF------------------------------- 98
WP_011499115_1 DUF63 ---------------------------------VFDA-P-----L--SYL--------F--ITPNIYF------------------------------- 98
WP_013037846_1 DUF63 --A------------------------------FSA--P-----L--KYL--------I--ITPNIYF------------------------------- 98
WP_048205640_1 DUF63 ---------------------------------IFER-P-----L--SYL--------F--ITPNIYF------------------------------- 98
```

```
WP_072561629_1 DUF63 ---------------------------------VFSA-P-----L--KYL--------F--ITPNIYF-------------------------------- 98
WP_072360082_1 DUF63 ---------------------------------VFSA-P-----L--KYL--------F--ITPNIYF-------------------------------- 98
WP_096711793_1 DUF63 ---------------------------------VFSA-P-----L--KYL--------F--ITPNIYF-------------------------------- 98
ODV50598_1 hypotheti ---------------------------------VFSA-P-----L--KYL--------F--ITPNIYF-------------------------------- 98
WP_013194042_1 DUF63 --I-------------------------------FNA-P-----V--NYL--------L--ITPIIYF-------------------------------- 98
WP_048178067_1 DUF63 --A-----------------------------GIINP-P-----F--SYL--------L--ITPNIYF-------------------------------- 100
WP_048127111_1 DUF63 --A-----------------------------DIFHP-P-----Y--SYV--------L--ITPNIYF-------------------------------- 100
WP_011023594_1 DUF63 ---------------------------------IFHP-P-----F--SYL--------L--ITPNIYF-------------------------------- 104
WP_048184587_1 DUF63 -------------------------------SDIFHP-P-----L--SYM--------L--ITPNIYF-------------------------------- 100
WP_011032542_1 DUF63 ---------------------------------IFHP-P-----F--SYL--------L--ITPNIYF-------------------------------- 100
WP_011032542_1 DUF63 ---------------------------------IFHP-P-----F--SYL--------L--ITPNIYF-------------------------------- 100
WP_048129141_1 DUF63 ---------------------------------IFHP-P-----F--SYL--------L--ITPNIYF-------------------------------- 100
WP_048129141_1 DUF63 ---------------------------------IFHP-P-----F--SYL--------L--ITPNIYF-------------------------------- 100
WP_048169891_1 DUF63 ---------------------------------IFHP-P-----F--SYL--------L--ITPNIYF-------------------------------- 100
WP_048137175_1 DUF63 ---------------------------------IFHP-P-----F--SYL--------L--ITPNIYF-------------------------------- 100
WP_048137175_1 DUF63 ---------------------------------IFHP-P-----F--SYL--------L--ITPNIYF-------------------------------- 100
WP_048137175_1 DUF63 ---------------------------------IFHP-P-----F--SYL--------L--ITPNIYF-------------------------------- 100
WP_048137694_1 DUF63 ---------------------------------IFHP-P-----F--SYL--------L--ITPNIYF-------------------------------- 100
WP_048168126_1 DUF63 --A-----------------------------GIFHP-P-----F--SYL--------L--ITPNIYF-------------------------------- 100
WP_048118082_1 DUF63 ---------------------------------IFHP-P-----F--SYL--------L--ITPNIYF-------------------------------- 100
WP_048158120_1 DUF63 ---------------------------------IFHP-P-----F--SYL--------L--ITPNIYF-------------------------------- 100
WP_011305329_1 DUF63 ---------------------------------IFHP-P-----F--SYL--------L--ITPNIYF-------------------------------- 100
WP_054298619_1 DUF63 ---------------------------------IFHP-P-----F--SYL--------L--ITPNIYF-------------------------------- 100
ALK05385_1 hypotheti ---------------------------------IFHP-P-----F--SYL--------L--ITPNIYF-------------------------------- 100
WP_015052897_1 DUF63 ---------------------------------VLEP-P-----V--SYL--------F--ITPNIYF-------------------------------- 98
WP_023846134_1 DUF63 ---------------------------------ILTH-P-----L--SYL--------F--ITPNIYF-------------------------------- 97
OIN88451_1 hypotheti --I-----------------------------------------I-SGYL--------F--VTPNIWL-------------------------------- 89
PIX50278_1 hypotheti --------------------------------------------IISGYL--------F--VTPNIWL-------------------------------- 89
PIW41402_1 hypotheti --------------------------------------------IISGYL--------F--VTPNIWL-------------------------------- 89
PIY35178_1 hypotheti --------------------------------------------IISGYL--------F--VTPNIWL-------------------------------- 89
PJB74886_1 hypotheti --------------------------------------------IISGYL--------F--VTPNIWL-------------------------------- 89
PIZ33651_1 hypotheti --------------------------------------------IISGYL--------F--VTPNIWL-------------------------------- 89
WP_048165029_1 DUF63 ------------------------------------IIP-----R--TFL--------T--VTPGIYF-------------------------------- 91
WP_042681046_1 DUF63 --I---------------------------------Y-P-----R--TYL--------T--VTPGIYV-------------------------------- 91
WP_013467319_1 DUF63 --I---------------------------------Y-P-----R--TYL--------T--VTPGIYV-------------------------------- 91
WP_048152160_1 DUF63 --I---------------------------------I-P-----R--TYL--------T--VTPGIYI-------------------------------- 91
WP_015849008_1 DUF63 ------------------------------------IIP-----Q--SYL--------T--VTPGIYF-------------------------------- 91
WP_004069276_1 DUF63 ------------------------------------IIP-----R--TYL--------T--VTPGIYF-------------------------------- 91
WP_058946638_1 DUF63 ------------------------------------IIP-----R--TYL--------T--VTPGIYF-------------------------------- 91
WP_042701551_1 DUF63 ------------------------------------IIP-----R--TYL--------T--VTPGIYF-------------------------------- 91
WP_055282692_1 DUF63 ------------------------------------IIP-----R--TYL--------T--VTPGIYI-------------------------------- 91
WP_013906023_1 DUF63 --V---------------------------------Y-P-----R--TYL--------T--VSPGIYF-------------------------------- 90
WP_011013158_1 DUF63 --V---------------------------------F-P-----R--TYL--------T--VSPGIYF-------------------------------- 90
WP_014733166_1 DUF63 --V---------------------------------F-P-----R--TYI--------T--VSPGIYF-------------------------------- 90
WP_068322889_1 DUF63 --V---------------------------------Y-P-----R--TYI--------T--VSPGIYF-------------------------------- 90
WP_068575625_1 DUF63 --V---------------------------------Y-P-----R--TYI--------T--VSPGIYF-------------------------------- 90
WP_010885960_1 DUF63 --V---------------------------------F-P-----R--TYI--------T--VSPGIYF-------------------------------- 90
WP_010867401_1 DUF63 --V---------------------------------F-P-----R--TYI--------T--VSPGIYF-------------------------------- 90
WP_013747987_1 DUF63 --V---------------------------------F-P-----R--TYI--------T--VSPGIYF-------------------------------- 91
WP_068664049_1 DUF63 --I---------------------------------Y-P-----R--TYI--------T--VTPGIYF-------------------------------- 91
WP_011249739_1 DUF63 ------------------------------------LLP-----R--TYL--------T--VSPGGYF-------------------------------- 91
WP_062386854_1 DUF63 ------------------------------------LLP-----R--TYL--------T--VSPGGYF-------------------------------- 91
WP_050003781_1 DUF63 ------------------------------------LLP-----R--TYL--------T--VSPGGYF-------------------------------- 91
WP_042690699_1 DUF63 ------------------------------------LLP-----R--TYL--------T--VSPGGYF-------------------------------- 91
WP_088885212_1 DUF63 -----------------------------------ML-P-----R--TYL--------T--VSPGGYF-------------------------------- 91
WP_010478519_1 DUF63 ----------------------------------M-L-P-----R--TYL--------T--VSPGGYF-------------------------------- 91
WP_088858240_1 DUF63 ------------------------------------MLP-----R--TYL--------T--VSPGGYF-------------------------------- 91
WP_015859016_1 DUF63 -----------------------------------ML-P-----R--TYL--------T--VSPGGYF-------------------------------- 91
WP_014121940_1 DUF63 -----------------------------------ML-P-----R--TYL--------T--VSPGGYF-------------------------------- 91
WP_062373911_1 DUF63 ------------------------------------MLP-----R--TYL--------T--VSPGAYF-------------------------------- 91
WP_088882924_1 DUF63 --I---------------------------------L-P-----R--TYL--------T--VSPGAYF-------------------------------- 91
WP_088862911_1 DUF63 --I---------------------------------L-P-----R--TYL--------T--VSPGGYF-------------------------------- 91
WP_012572006_1 DUF63 --I---------------------------------L-P-----R--TYL--------T--VSPGGYF-------------------------------- 91
WP_088854707_1 DUF63 --I---------------------------------L-P-----R--TYL--------T--VSPGGYF-------------------------------- 91
WP_014789474_1 DUF63 --I---------------------------------L-P-----R--TYL--------T--VSPGGYF-------------------------------- 91
WP_088180728_1 DUF63 --I---------------------------------L-P-----R--TYL--------T--VSPGGYF-------------------------------- 91
WP_088864937_1 DUF63 --I---------------------------------L-P-----R--TYL--------T--VSPGGYF-------------------------------- 91
WP_055429686_1 DUF63 --I---------------------------------L-P-----R--TYL--------T--VSPGGYF-------------------------------- 91
WP_088865932_1 DUF63 --I---------------------------------L-P-----R--TYL--------T--VSPGGYF-------------------------------- 91
WP_014013642_1 DUF63 --V---------------------------------L-P-----R--TYL--------T--VSPGGYF-------------------------------- 91
WP_088856804_1 DUF63 --I---------------------------------L-P-----R--TYL--------T--VSPGGYF-------------------------------- 91
WP_058939475_1 DUF63 --V---------------------------------L-P-----R--TYL--------T--VSPGGYF-------------------------------- 91
EHR77276_1 conserved --F------------------------------FSAE-------M--DVL--------F--ISPLIHL-------------------------------- 148
OUV40025_1 hypotheti ------------------------------------FFP-----SSIDWL--------L--ISPIIHL-------------------------------- 149
MBJ52984_1 hypotheti --F------------------------------FKEN-------I--DIL--------F--ISPVIHF------------------------HLAIWLIF 150
PDH23744_1 hypotheti --L-----------------------------------------A--PYF-----------VSPGIHF-------------------------------- 141
PDH25468_1 hypotheti --M------------------------------FGEV-L-----S--AWF-----------VSPGVHFQTAGWVILSGAAGYSIINNNENSIPKFKRVNT 173

WP_048201856_1 hypot -------------------------------VINIIFISSLKE-------------------------STKS---------------------------- 96
WP_048150344_1 hypot -------------------------------VVGIITVASLLV-------------------------NIFT----YEL--------------------- 104
AOV95360_1 hypotheti -------------------------------TLTAITLLTFII-------------------------THVY---RQRK--------------------- 89
EOD42420_1 Uncharact -------------------------------WILISFLPVLFL-------------------------Y------------------------------- 106
AOV95164_1 hypotheti -------------------------------VIAGLFLASLLI-------------------------SSRM---------------------------- 122
EGQ40074_1 putative  -------------------------------VVAGLFVACLAA-------------------------ARRF---------------------------- 126
KYK23067_1 hypotheti -------------------------------QTAFYVLIFLFL-------------------------GYYL---QKKA--------------------- 162
WP_084383883_1 DUF63 -------------------------------VVTAVTVLALGI-------------------------LITY---------------------------- 163
WP_049984677_1 DUF63 -------------------------------VVTAITILALGV-------------------------GALA----RNR--------------------- 167
WP_103428047_1 hypot -------------------------------VVTAVTVLALGV-------------------------GARA----RDR--------------------- 166
WP_004048647_1 DUF63 -------------------------------VVTAVTVLALGI-------------------------GALA----RDR--------------------- 166
WP_004594600_1 DUF63 -------------------------------VVTAVTILALGV-------------------------GALL-----RD--------------------- 165
WP_004594600_1 DUF63 -------------------------------VVTAVTILALGV-------------------------GALL-----RD--------------------- 165
WP_050050397_1 DUF63 -------------------------------VVTAVTILALGV-------------------------GALL-----RD--------------------- 165
WP_004594600_1 DUF63 -------------------------------VVTAVTILALGV-------------------------GALL-----RD--------------------- 165
WP_050050397_1 DUF63 -------------------------------VVTAVTILALGV-------------------------GALL-----RD--------------------- 165
WP_049983690_1 DUF63 -------------------------------VVTAVTVLSLGV-------------------------GALA----RNR--------------------- 166
```

```
WP_103428162_1 hypot ------------------------------VVTAVTVLSLGV------------------------GALA----RDR-------------------- 166
WP_009378268_1 DUF63 ------------------------------VVTAVTVLSLGV------------------------GALA----RDR-------------------- 166
WP_015763176_1 DUF63 ------------------------------TLFVLTLGSLVV------------------------VRRL----ERG-------------------- 198
WP_018259238_1 DUF63 ------------------------------TLFVLTLGSLVA------------------------VRRL----ERG-------------------- 198
WP_004592615_1 DUF63 ------------------------------TVFLLTLLVLVV------------------------CVGL----ERR-------------------- 192
WP_004518092_1 DUF63 ------------------------------TVFLLTLLVLVV------------------------CVDL----EQR-------------------- 192
WP_014040450_1 DUF63 ------------------------------TVFLLTLLVLVV------------------------CVDL----EQR-------------------- 192
WP_014040450_1 DUF63 ------------------------------TVFLLTLLVLVV------------------------CVDL----EQR-------------------- 192
WP_008309018_1 DUF63 ------------------------------TVFLLTLLVLVV------------------------CVDL----EQR-------------------- 192
WP_014040450_1 DUF63 ------------------------------TVFLLTLLVLVV------------------------CVDL----EQR-------------------- 192
WP_014040450_1 DUF63 ------------------------------TVFLLTLLVLVV------------------------CVDL----EQR-------------------- 192
WP_005534235_1 DUF63 ------------------------------TVFLLTLLVLVA------------------------CVDL----ERR-------------------- 192
WP_004961153_1 DUF63 ------------------------------TVFLLTLLVLVV------------------------CVDL----EQR-------------------- 192
WP_004961153_1 DUF63 ------------------------------TVFLLTLLVLVV------------------------CVDL----EQR-------------------- 192
WP_101350151_1 hypot ------------------------------TVFLLTLLVLVA------------------------CVDL----EQR-------------------- 192
WP_053968273_1 DUF63 ------------------------------TVFLLTLLVLVA------------------------CVDL----EQR-------------------- 192
WP_058995687_1 DUF63 ------------------------------TVFLLTLLVLVA------------------------CVDL----EER-------------------- 192
WP_015790673_1 DUF63 ------------------------------TVFGFAIAALVG------------------------SLRL----ADR-------------------- 193
WP_008524115_1 DUF63 ------------------------------TVFGIAIAALVG------------------------SLRL----ADR-------------------- 193
WP_075936143_1 DUF63 ------------------------------TVFFITLATVVA------------------------SVLA----ERR-------------------- 186
WP_020446311_1 DUF63 ------------------------------TVFLLAVIAITA------------------------SVWL----ERD-------------------- 188
WP_049898450_1 DUF63 ------------------------------TVFVVTLVALLV------------------------ALRL----ARR-------------------- 190
WP_006077061_1 DUF63 ------------------------------TVFLVTLVALLA------------------------ALWL----SRR-------------------- 190
WP_049996700_1 DUF63 ------------------------------TVFVVTLAAILL------------------------SLWL----ARR-------------------- 190
EMA38628_1 hypotheti ------------------------------TVFVVTLAALLV------------------------SLRL----ARR-------------------- 190
WP_049992561_1 DUF63 ------------------------------TVFFVTLGALLS------------------------SVAL----SRR-------------------- 191
WP_010903025_1 DUF63 ------------------------------VMFAVTLVAVVA------------------------AVAV---ARRT-------------------- 189
WP_009760947_1 DUF63 ------------------------------VMFAITLVVLVA------------------------AVYL---ARRT-------------------- 188
WP_059057097_1 DUF63 ------------------------------VMFVITLAALVA------------------------CVFL---ARRT-------------------- 188
WP_058983522_1 DUF63 ------------------------------VMFAITLAALVA------------------------CVSL---ARRT-------------------- 188
WP_071932813_1 DUF63 ------------------------------TVFVLAMAALLV------------------------SLRL----RNQ-------------------- 184
WP_050048584_1 DUF63 ------------------------------TVFFFTLAALLA------------------------SVWL----DRK-------------------- 184
WP_014051341_1 DUF63 ------------------------------TVFVVAVAAILV------------------------SIWL----TRQ-------------------- 187
WP_079233299_1 DUF63 ------------------------------TIFVIALACILA------------------------ALGL----SRQ-------------------- 186
WP_053948572_1 DUF63 ------------------------------TLFVIALACILL------------------------SLWL----ARQ-------------------- 186
WP_049980835_1 DUF63 ------------------------------TVFVIALASILL------------------------ALWL----ARR-------------------- 186
KPN31749_1 hypotheti ------------------------------TIFVIALGSILL------------------------ALWL----SRR-------------------- 136
AGB16040_1 putative  ------------------------------VVALIALVSLVI------------------------AVWL----ERN-------------------- 185
WP_007696755_1 DUF63 ------------------------------VVAVIALVSLVI------------------------AVWL----ERT-------------------- 185
WP_015322904_1 DUF63 ------------------------------TVFLVALVAIVA------------------------GVWL----ERK-------------------- 185
WP_076581869_1 DUF63 ------------------------------TVALIALIAVVA------------------------SVWL----ERN-------------------- 185
WP_005559622_1 DUF63 ------------------------------TVFLVALVAIVA------------------------GVWL----ERK-------------------- 185
WP_006067640_1 DUF63 ------------------------------TVTLIALVALVG------------------------AVFL----ERN-------------------- 180
WP_008164554_1 DUF63 ------------------------------TVALIALVALVG------------------------AVSL----ERN-------------------- 185
WP_006088112_1 DUF63 ------------------------------TVALIAFIAVVG------------------------SVWL----ERN-------------------- 185
WP_012943247_1 DUF63 ------------------------------TVALLTVITVVV------------------------AVWL----DRT-------------------- 185
WP_008895026_1 DUF63 ------------------------------TVALLTVIAVVA------------------------SVWL----DRT-------------------- 185
WP_098727043_1 DUF63 ------------------------------TVFFVALLAVVS------------------------SVWL----ERN-------------------- 185
WP_049990861_1 DUF63 ------------------------------TVFFIALFAVVL------------------------SVWL----DRR-------------------- 183
WP_008013001_1 DUF63 ------------------------------TVFFIALAALVS------------------------SIWL----DRN-------------------- 185
WP_006180566_1 DUF63 ------------------------------TVFFIALFAVVT------------------------SVWL----ERN-------------------- 185
WP_006650865_1 DUF63 ------------------------------TVFFIALFAVVT------------------------SIWL----ERN-------------------- 185
WP_066301295_1 DUF63 ------------------------------TVFFIALFAVVT------------------------SVWL----ERN-------------------- 185
WP_049966804_1 DUF63 ------------------------------TVFFIALFAVVT------------------------SIWL----ERN-------------------- 185
WP_076145380_1 DUF63 ------------------------------TVFFIALFAVVT------------------------SVWL----ERN-------------------- 185
WP_097378845_1 DUF63 ------------------------------TVFFIALFAVVF------------------------SVWL----DRN-------------------- 185
WP_008452261_1 DUF63 ------------------------------TVFFIALFAVVT------------------------SVWL----DRN-------------------- 185
WP_008452261_1 DUF63 ------------------------------TVFFIALFAVVT------------------------SVWL----DRN-------------------- 185
WP_006432643_1 DUF63 ------------------------------TVFFIALVSVIA------------------------SVWL----DRN-------------------- 185
WP_008452261_1 DUF63 ------------------------------TVFFIALFAVVT------------------------SVWL----DRN-------------------- 185
WP_007109662_1 DUF63 ------------------------------TVFFIALFAVVM------------------------SVWL----DRN-------------------- 185
WP_049952821_1 DUF63 ------------------------------TVFFFTLFAVLV------------------------SVWL----ERN-------------------- 185
WP_086889541_1 DUF63 ------------------------------LVFLITLVAVVI------------------------SVWL----DRR-------------------- 188
WP_005580009_1 DUF63 ------------------------------TVTMLAVVAIVI------------------------AVWL----ERN-------------------- 188
WP_049927356_1 DUF63 ------------------------------VVFVVALVALVV------------------------SIWL----ERR-------------------- 185
WP_087714455_1 DUF63 ------------------------------VVFIIAAIAVTV------------------------SVWL----ERE-------------------- 185
WP_013878438_1 DUF63 ------------------------------TVFAVTAVAVVV------------------------SVWL----ERN-------------------- 185
WP_049921111_1 DUF63 ------------------------------TVFFVTLAAVVV------------------------SVWL----DRN-------------------- 185
WP_007142789_1 DUF63 ------------------------------TVFLVTLFAVVG------------------------SVWL----DRN-------------------- 184
WP_006651941_1 DUF63 ------------------------------VVFFIAFVAVVV------------------------SVWL----ERN-------------------- 185
WP_004216437_1 DUF63 ------------------------------VVFFIAFVAVVV------------------------SVWL----ERN-------------------- 185
WP_071402513_1 DUF63 ------------------------------VVFFVAFVAVVV------------------------SVWL----ERN-------------------- 185
WP_006666462_1 DUF63 ------------------------------TVFFMALLAVVV------------------------SVWL----ERN-------------------- 185
WP_049904535_1 DUF63 ------------------------------TVFFLALLAVVA------------------------SVWL----ERN-------------------- 185
WP_006824405_1 DUF63 ------------------------------TVFFLTLLAVVV------------------------SVWL----ERN-------------------- 185
WP_011323802_1 DUF63 ------------------------------VMFGFTLAALVG------------------------GVWL----EGR-------------------- 186
WP_015409719_1 DUF63 ------------------------------VMFAFTLGAVVL------------------------WILL----SRQ-------------------- 187
WP_006883947_1 DUF63 ------------------------------TVFLITLAALLA------------------------SLWL----ESR-------------------- 186
WP_077207545_1 DUF63 ------------------------------TVFFVTLASLLV------------------------TIYL----RRK-------------------- 189
ESS12903_1 putative  ------------------------------TVFAVALAAFVG------------------------SVAA----ARA-------------------- 189
WP_008417361_1 DUF63 ------------------------------TVFFVTLGALLV------------------------SVAA----ERS-------------------- 188
WP_066381608_1 DUF63 ------------------------------TVFFVTLAAVLV------------------------SVAA----ARG-------------------- 183
WP_049947642_1 DUF63 ------------------------------TVFAVALAAVVG------------------------CVRL----ARN-------------------- 189
ESS05953_1 putative  ------------------------------TVFVVALAALLT------------------------AVAV----ARR-------------------- 183
WP_096390051_1 DUF63 ------------------------------TVFAITLGCVVL------------------------AYAL----AAR-------------------- 195
WP_021072749_1 DUF63 ------------------------------TVFAITLGCVVL------------------------AYAL----AAR-------------------- 195
WP_049982791_1 DUF63 ------------------------------TVFGVTLGCVVA------------------------AYGL----AGR-------------------- 193
WP_008585990_1 DUF63 ------------------------------TVFLFTLACVIA------------------------AYAL----ERR-------------------- 195
WP_006628721_1 DUF63 ------------------------------TVFAFTLACVIA------------------------AYAL----ERR-------------------- 193
WP_006113218_1 DUF63 ------------------------------TVFLFTLACVLA------------------------AYGL----ERR-------------------- 194
WP_049930034_1 DUF63 ------------------------------TVFLFTLACVLA------------------------AYGL----ERR-------------------- 195
WP_049906047_1 DUF63 ------------------------------TIFLFTLACVLA------------------------AYGL----ERR-------------------- 195
WP_044965494_1 DUF63 ------------------------------TVFLFTLACVLA------------------------AYGL----ERR-------------------- 195
WP_096393195_1 DUF63 ------------------------------TVFLFTLACVLA------------------------AYGL----ERR-------------------- 195
WP_049908400_1 DUF63 ------------------------------TVFLFTLACVLA------------------------AYGL----ERR-------------------- 195
```

```
WP_049908585_1 DUF63 -------------------------------TVFLFTLACVLA-------------------------AYGL----ERR--------------------- 195
WP_049902631_1 DUF63 -------------------------------TVFLFTLACVLA-------------------------AYGL----ERR--------------------- 195
WP_049983668_1 DUF63 -------------------------------TVFLFTLACVLA-------------------------AFGL----ERR--------------------- 194
WP_053772267_1 DUF63 -------------------------------TVFLFTLACVLT-------------------------AYGL----ERR--------------------- 194
WP_007999151_1 DUF63 -------------------------------TVFGFTLLCVLA-------------------------AYAL----ANR--------------------- 193
WP_015909917_1 DUF63 -------------------------------IVFGVTLACVLA-------------------------AYSL----EGR--------------------- 194
WP_004050594_1 DUF63 -------------------------------TVFGVTLACVLA-------------------------AYSL----AGR--------------------- 194
WP_095636035_1 DUF63 -------------------------------TVFGVTLACVLA-------------------------AYAL----ADR--------------------- 194
WP_008003811_1 DUF63 -------------------------------TVFGVTLACVIA-------------------------AYSL----ADR--------------------- 194
WP_066416034_1 DUF63 -------------------------------TVFGVTLLCVMA-------------------------AYAL----EDR--------------------- 195
ESS03170_1 putative  -------------------------------TVFAVTLVCVVG-------------------------AYAA----ERQ--------------------- 194
WP_089671383_1 DUF63 -------------------------------TMFVLTLVAVLV-------------------------AIRL----ESS--------------------- 193
ERH07656_1 putative  -------------------------------TVFVVTLAAVGA-------------------------GVSL----ERR--------------------- 193
ESS10096_1 putative  -------------------------------TVFVVTLAAVGA-------------------------GVSL----ERR--------------------- 193
ESS07824_1 putative  -------------------------------PLSAAIISPFIY--------------------------------------------------------- 173
ERH05689_1 putative  -------------------------------TMFLLTLAAIGV-------------------------GIGL----ASR--------------------- 193
ERH02266_1 putative  -------------------------------TMFLLTLAAIGV-------------------------GIGL----ANR--------------------- 193
ESS07825_1 putative  -------------------------------TVNLQVLAA------------------------------------------------------------ 30
WP_049970017_1 DUF63 -------------------------------TVFAITLLAVVV-------------------------SVTA----ARK--------------------- 187
WP_007979289_1 DUF63 -------------------------------TVFFVTLAGLVV-------------------------SISL----ARR--------------------- 187
WP_007977745_1 DUF63 -------------------------------TVFAITLAALVV-------------------------SVAL----ARR--------------------- 187
WP_014556445_1 DUF63 -------------------------------TVFAVTLVAVGF-------------------------TVWA----ERR--------------------- 188
WP_049935775_1 DUF63 -------------------------------TVFAVTLLAVVA-------------------------SVAL----ARR--------------------- 186
WP_008325149_1 DUF63 -------------------------------TVFAITLVAVVG-------------------------SVTA----ERS--------------------- 188
WP_007543631_1 DUF63 -------------------------------TVFAITLVAVVG-------------------------SVTA----ERS--------------------- 188
WP_049905041_1 DUF63 -------------------------------TVFAITLVAVVA-------------------------SVFA----ERR--------------------- 188
WP_049913563_1 DUF63 -------------------------------TVFAITLVAVVG-------------------------SVFA----ERQ--------------------- 188
WP_049967851_1 DUF63 -------------------------------TVFAITLVAVVG-------------------------SVFA----ERQ--------------------- 188
WP_049914905_1 DUF63 -------------------------------TVFAITLVAVVG-------------------------SVLA----ERR--------------------- 188
WP_049916430_1 DUF63 -------------------------------TVFAITLVAVVG-------------------------SVLA----ERR--------------------- 188
WP_049896892_1 DUF63 -------------------------------TVFAITLVAVVG-------------------------SVLA----ERR--------------------- 188
WP_049896892_1 DUF63 -------------------------------TVFAITLVAVVG-------------------------SVLA----ERR--------------------- 188
WP_049896892_1 DUF63 -------------------------------TVFAITLVAVVG-------------------------SVLA----ERR--------------------- 188
WP_058828253_1 DUF63 -------------------------------TVFAITLVAVVG-------------------------SVFA----ERR--------------------- 188
WP_058568959_1 DUF63 -------------------------------TVFAITLVAVVG-------------------------SVLA----ERR--------------------- 188
WP_049917947_1 DUF63 -------------------------------TVFAITLVAVVG-------------------------SVFA----ERR--------------------- 188
WP_049920348_1 DUF63 -------------------------------TVFAITLVAVVG-------------------------SVLA----ERR--------------------- 188
WP_089777545_1 DUF63 -------------------------------TVFAITLVAVVG-------------------------SVLA----ERR--------------------- 188
WP_008320633_1 DUF63 -------------------------------TVFAITLVAVVG-------------------------AVVA----ERR--------------------- 188
WP_004060355_1 DUF63 -------------------------------TVFAITLVAVVG-------------------------SVVA----ERE--------------------- 188
WP_103426078_1 hypot -------------------------------TVFAVTLVAVVV-------------------------AVLL----ERQ--------------------- 188
WP_009367433_1 DUF63 -------------------------------TVFVITLGAVVG-------------------------SVAL----ERN--------------------- 187
WP_013440552_1 DUF63 -------------------------------TVFLVTLGAIAV-------------------------AVAL----EHT--------------------- 188
WP_049916626_1 DUF63 -------------------------------TVFLITLGAVVA-------------------------AVAL----ERT--------------------- 188
WP_058582837_1 DUF63 -------------------------------TVFFITLAAVVV-------------------------TVTL----SRN--------------------- 190
WP_101298124_1 hypot -------------------------------TVFFVTLAVVVA-------------------------SVWL----DRR--------------------- 190
PIN95153_1 hypotheti -------------------------------VLVLIVFYYLIK-------------------------EITM---------------------------- 66
PIU22108_1 hypotheti -------------------------------LLVAFALLLLGI-------------------------TCFL----KKK--------------------- 119
AAR39201_1 NEQ352    -----------------------------------LGIITYFI-------------------------ARKI---------------------------- 80
OIR14399_1 hypotheti -------------------------------TLALYSIGALWV-------------------------GKLI---KKVK--------------------- 145
OIR20963_1 hypotheti -------------------------------TITGIALFFIAL-------------------------GVWL----SKS--------------------- 149
OIR22371_1 hypotheti -------------------------------ILVLYSLLIIAL-------------------------GVWL----SKN--------------------- 160
EGQ43935_1 putative  -------------------------------LLILLGTLMIYG-------------------------SKKI----EAV--------------------- 114
MAG21679_1 hypotheti -------------------------------LVATITIATLVI-------------------------AKYL---------------------------- 116
PIN85618_1 hypotheti -------------------------------LTAGITIAALLI-------------------------AKKI----AKA--------------------- 119
PIN99249_1 hypotheti -------------------------------FIGLLTIFSLVL-------------------------SFFL----SKH--------------------- 123
AJF59838_1 hypotheti -------------------------------AVGLLAIAALIF-------------------------SIWL----AKK--------------------- 119
WP_042682145_1 DUF63 -------------------------------TAFFLIVPALIA-------------------------DAKL---------------------------- 110
WP_013468033_1 DUF63 -------------------------------TAFFLIVPALIA-------------------------DAKL---------------------------- 110
WP_055281581_1 DUF63 -------------------------------TAFILIVPALFA-------------------------DSKL---------------------------- 109
WP_048160382_1 DUF63 -------------------------------TAFFLILPALFA-------------------------DSRL---------------------------- 109
WP_042701216_1 DUF63 -------------------------------TAFLLILPALFA-------------------------DSRL---------------------------- 109
WP_004068659_1 DUF63 -------------------------------TAFLLILPALLV-------------------------DSRL---------------------------- 109
WP_058946665_1 DUF63 -------------------------------TAFLLILPALFA-------------------------DSRL---------------------------- 109
WP_014835499_1 DUF63 -------------------------------TAFFLIVPVLVV-------------------------DVKG---------------------------- 109
WP_010884243_1 DUF63 -------------------------------TAFFLILPVLFA-------------------------DVKL---------------------------- 112
WP_013748129_1 DUF63 -------------------------------TAFFLILPALIL-------------------------DVKF---------------------------- 108
WP_010867251_1 DUF63 -------------------------------TAFFLILPALFL-------------------------DVKL---------------------------- 108
WP_014733359_1 DUF63 -------------------------------TAFFLIVPVLIA-------------------------DVKL---------------------------- 109
WP_068319891_1 DUF63 -------------------------------TAFFLIVPVLFA-------------------------DVKL---------------------------- 109
WP_068576049_1 DUF63 -------------------------------TAFFLIVPVLFA-------------------------DVKL---------------------------- 109
WP_014013692_1 DUF63 -------------------------------TTFFIMLPAIVL-------------------------DAKL---------------------------- 114
WP_014789520_1 DUF63 -------------------------------TTFFVMLPAIVL-------------------------DAKL---------------------------- 114
WP_088180676_1 DUF63 -------------------------------TTFFIMLPAIVI-------------------------DAKL---------------------------- 114
WP_088864885_1 DUF63 -------------------------------TTFFVMLPAIVL-------------------------DAKL---------------------------- 114
WP_088856755_1 DUF63 -------------------------------TTFFIMLPVIVL-------------------------DAKL---------------------------- 114
WP_088865981_1 DUF63 -------------------------------TTFFIMLPAIVI-------------------------DAKL---------------------------- 114
WP_013906334_1 DUF63 -------------------------------TAFFLILPALIA-------------------------DAKL---------------------------- 110
WP_088862658_1 DUF63 -------------------------------TAFLLIVPAIMA-------------------------DAKL---------------------------- 113
WP_088882956_1 DUF63 -------------------------------TAFFLILPALVA-------------------------DARL---------------------------- 113
WP_012571938_1 DUF63 -------------------------------TAFVLIVPALVA-------------------------DAKL---------------------------- 110
WP_068663933_1 DUF63 -------------------------------TAFVLIVPALIA-------------------------DAKL---------------------------- 110
WP_074631153_1 DUF63 -------------------------------TAFILILPALAA-------------------------DAKM---------------------------- 110
WP_058939074_1 DUF63 -------------------------------TAFVLILPALAA-------------------------DAKM---------------------------- 114
WP_088854681_1 DUF63 -------------------------------TAFFLIVPAIAL-------------------------DAKL---------------------------- 114
WP_088885426_1 DUF63 -------------------------------TAFFLILPALFA-------------------------DAKL---------------------------- 110
WP_011249694_1 DUF63 -------------------------------TAFFLILPAIYA-------------------------DSKL---------------------------- 112
WP_062386972_1 DUF63 -------------------------------TAFFLILPAIYA-------------------------DSKL---------------------------- 112
WP_010478887_1 DUF63 -------------------------------TAFILIVPALIA-------------------------DAKL---------------------------- 110
WP_088858605_1 DUF63 -------------------------------TAFILIVPALIA-------------------------DAKL---------------------------- 110
WP_042690028_1 DUF63 -------------------------------TAFILIVPAIVA-------------------------DAKL---------------------------- 112
WP_048150762_1 DUF63 -------------------------------TAFVLIVPALVA-------------------------DAKL---------------------------- 110
WP_048811076_1 DUF63 -------------------------------TAFFLIVPALVA-------------------------DAKL---------------------------- 110
WP_050003256_1 DUF63 -------------------------------TAFFLILPSLVA-------------------------DARL---------------------------- 112
WP_062372100_1 DUF63 -------------------------------TAFFLIVPALVA-------------------------DAKL---------------------------- 110
```

```
WP_048165284_1 DUF63 -------------------------------TAFFLILPALVA-------------------------DVKL---------------------------- 112
WP_048148750_1 DUF63 -------------------------------TAFFLIVPALAV-------------------------DVKL---------------------------- 112
OYT53462_1 hypotheti -------------------------------LCVAVGLMGILL-------------------------GLLL----EKT--------------------- 93
KYC51152_1 hypotheti -------------------------------VTFILAFFTILL-------------------------GYQL----EKR--------------------- 93
KYC46003_1 hypotheti -------------------------------VTFILAFFTILL-------------------------GYQL----EKR--------------------- 93
KYC48643_1 hypotheti -------------------------------VTFILAFFTILL-------------------------GYQL----EKR--------------------- 93
KYC55321_1 hypotheti -------------------------------ITFILAFFMILI-------------------------GYQL----EKR--------------------- 93
KYC57927_1 hypotheti -------------------------------ITFILAFFMILI-------------------------GYQL----EKR--------------------- 93
KYC57171_1 hypotheti -------------------------------ITFILAFFMILI-------------------------GYQL----EKR--------------------- 93
OIO20701_1 hypotheti -------------------------------MVGGLFLLTIAV-------------------------CCRF---------------------------- 142
PIT83986_1 hypotheti -------------------------------AVAGMFFAAMYL-------------------------EKKM---------------------------- 159
OIO24760_1 hypotheti -------------------------------LVFLAVVAVLAI-------------------------AKSR---------------------------- 119
OIO24701_1 hypotheti -------------------------------ITFGLVALSIDA-------------------------ALAV----EKK--------------------- 121
OIO27021_1 hypotheti -------------------------------VTFAVILAGFGV-------------------------ARLL---------------------------- 115
OIO26762_1 hypotheti -------------------------------VTFALLAACFIA-------------------------FRKE---------------------------- 116
PIN95811_1 hypotheti -------------------------------VTFALLAACFIA-------------------------FRKE---------------------------- 116
PIO01637_1 hypotheti -------------------------------VTFALLAACFIA-------------------------FRKE---------------------------- 116
PIO02820_1 hypotheti -------------------------------VTFAVILAGFGV-------------------------ARLL---------------------------- 115
PJD01038_1 hypotheti -------------------------------VTFAVILAGFGV-------------------------ARLL---------------------------- 115
PIZ91366_1 hypotheti -------------------------------VTFALLAACFIA-------------------------FRKE---------------------------- 116
WP_013100570_1 DUF63 -------------------------------LVGGFYILTILV-------------------------TGYF---------------------------- 109
WP_004590770_1 DUF63 -------------------------------VIGGLFIITILT-------------------------TAYV---------------------------- 109
WP_048196979_1 DUF63 -------------------------------LIGGFFILTILT-------------------------SAII---------------------------- 109
WP_015791538_1 DUF63 -------------------------------LVGGFFIITILI-------------------------TGII---------------------------- 109
WP_048202292_1 DUF63 -------------------------------LIGGFFIATILI-------------------------TGLI---------------------------- 109
WP_012981280_1 DUF63 -------------------------------LIGGFFIATILI-------------------------TGVV---------------------------- 109
WP_064496496_1 DUF63 -------------------------------LIGGFFILTILT-------------------------TGLV---------------------------- 109
WP_011972741_1 DUF63 -------------------------------VLGIYYIFSIIL-------------------------TGYF---------------------------- 111
WP_013798289_1 DUF63 -------------------------------LIGSYYMASILI-------------------------SGVL---------------------------- 109
WP_013181100_1 DUF63 -------------------------------LIGAYYLLSLII-------------------------SGAL---------------------------- 111
WP_013867628_1 DUF63 -------------------------------LVGLYYMSSIII-------------------------SGIL---FKKD--------------------- 115
WP_018153391_1 DUF63 -------------------------------LVGLYYMLSIIG-------------------------SGAL---------------------------- 111
WP_011170272_1 DUF63 -------------------------------AVGIYYMLSIII-------------------------SGIL----LKK--------------------- 114
WP_012066151_1 DUF63 -------------------------------FVGIYYMLSIII-------------------------SGTI---------------------------- 111
ODS43013_1 hypotheti -------------------------------TMFALTLAALAA-------------------------AKYL---QKKK--------------------- 119
OIQ06085_1 hypotheti -------------------------------TIFLIAVASLFI-------------------------GLYI---------------------------- 112
PIN67069_1 hypotheti -------------------------------TIFLIAVASLFI-------------------------GLYI---------------------------- 112
PIV28099_1 hypotheti -------------------------------TIFLIAVASLFI-------------------------GLYI---------------------------- 112
PJC13070_1 hypotheti -------------------------------TIFLIAVASLFI-------------------------GLYI---------------------------- 112
PIZ29927_1 hypotheti -------------------------------TIFLIAVASLFI-------------------------GLYI---------------------------- 112
PKP60688_1 hypotheti -------------------------------TIFLITIASLLI-------------------------GLYI---------------------------- 112
WP_012956954_1 DUF63 -------------------------------LVGLITIASLLF-------------------------SLFL----YNR--------------------- 108
WP_067148430_1 DUF63 -------------------------------LVGLLTILSFLF-------------------------SVFL---FNKR--------------------- 109
WP_080460538_1 DUF63 -------------------------------IVAGIVVLSLFL-------------------------GIFI----QKK--------------------- 107
WP_010877090_1 DUF63 -------------------------------LVGLTTIATLIA-------------------------AVKL---EETR--------------------- 109
WP_013294901_1 DUF63 -------------------------------LVGLTAIATLLV-------------------------SVKL----EKL--------------------- 108
WP_010877090_1 DUF63 -------------------------------LVGLTTIATLIA-------------------------AVKL---EETR--------------------- 109
BAZ99473_1 hypotheti -------------------------------LVGLTTIATLIA-------------------------AVKL---EETR--------------------- 109
PKL66404_1 hypotheti -------------------------------LTGLITILTLLG-------------------------SIYL----EKK--------------------- 116
WP_023991171_1 DUF63 -------------------------------LTGLGTVFTLLA-------------------------SVLI----ERK--------------------- 109
WP_048072898_1 DUF63 -------------------------------LTGLTAIFTLLA-------------------------SVLI----ERK--------------------- 109
WP_100905180_1 hypot -------------------------------LTGISAILTLLG-------------------------SVLI----ERK--------------------- 109
WP_100907231_1 hypot -------------------------------LTGISAILTLLG-------------------------SVLI----ERK--------------------- 109
WP_100907231_1 hypot -------------------------------LTGISAILTLLG-------------------------SVLI----ERK--------------------- 109
WP_048081925_1 DUF63 -------------------------------LTGFAAIATLLL-------------------------SVYV----ERK--------------------- 108
WP_048081925_1 DUF63 -------------------------------LTGFAAIATLLL-------------------------SVYV----ERK--------------------- 108
WP_069585525_1 DUF63 -------------------------------LTGFAAIATLLI-------------------------SVYV----EKK--------------------- 108
WP_069585525_1 DUF63 -------------------------------LTGFAAIATLLI-------------------------SVYV----EKK--------------------- 108
WP_013643671_1 DUF63 -------------------------------LTGFIAIITVLS-------------------------AYFI----EKK--------------------- 108
WP_013824571_1 DUF63 -------------------------------LTGFIAIASVLS-------------------------AVLI----ERK--------------------- 108
WP_048192110_1 DUF63 -------------------------------FTGLMAIFTVLA-------------------------SVYI----EKK--------------------- 112
WP_081810122_1 DUF63 -------------------------------VIFAICFSILLL-------------------------TLYL----ERT--------------------- 136
WP_097298965_1 DUF63 -------------------------------VIFAICFGTLLV-------------------------ARYL----EKR--------------------- 112
PIO00292_1 hypotheti -------------------------------VIGAIIIGCFFI-------------------------SQKT---------------------------- 110
WP_011833679_1 DUF63 -------------------------------LTMFFAMAMLFI-------------------------SYQL----QKK--------------------- 108
WP_042698332_1 DUF63 -------------------------------LTLIFAIVMLVI-------------------------SYQL----QKN--------------------- 108
WP_042698332_1 DUF63 -------------------------------LTLIFAIVMLVI-------------------------SYQL----QKN--------------------- 108
WP_011448164_1 DUF63 -------------------------------LIFALAMPVLFI-------------------------ANIC----EKR--------------------- 111
WP_007314214_1 DUF63 -------------------------------VMFLYTAGALFV-------------------------SATL----QEH--------------------- 116
WP_042705632_1 DUF63 -------------------------------VIFFYTIFALLI-------------------------SRKL----QNA--------------------- 111
WP_013329717_1 DUF63 -------------------------------VIFFYAIVVLVL-------------------------SRAL----EGR--------------------- 111
WP_004077722_1 DUF63 -------------------------------AIFFYAIIVLII-------------------------SRTL----ESK--------------------- 111
WP_048150927_1 DUF63 -------------------------------VIFFYAIVVLVL-------------------------SRAL----EGR--------------------- 111
WP_012107531_1 DUF63 -------------------------------VLFFFTLAMIFI-------------------------GGTL----KKN--------------------- 117
PKL70081_1 hypotheti -------------------------------LLFFYAVIILFV-------------------------SKYL----QEQ--------------------- 111
WP_015285007_1 DUF63 -------------------------------VLFFFTIGMLFL-------------------------SRYL----TLQ--------------------- 117
PKL64570_1 hypotheti -------------------------------VLFAFTVSMMFL-------------------------SLFL----VKR--------------------- 117
WP_015286404_1 DUF63 -------------------------------VLFFFTVAALLL-------------------------SRTL----ERA--------------------- 111
WP_012617618_1 DUF63 -------------------------------TIFFFTIACLFV-------------------------SRLA----EHY--------------------- 111
WP_014867374_1 DUF63 -------------------------------TVFVVAAVALFA-------------------------GKLA----ENA--------------------- 111
CVK33108_1 conserved -------------------------------TVFVVAAVALFA-------------------------GKLA----ENA--------------------- 111
WP_066956331_1 DUF63 -------------------------------VIFVIAVVALFA-------------------------GKFA----ENA--------------------- 111
WP_011844879_1 DUF63 -------------------------------SIFAVAAIALFA-------------------------GKLA----ENA--------------------- 111
WP_048181868_1 DUF63 -------------------------------TIFAVAAIALFA-------------------------GKLA----ENA--------------------- 111
WP_067073014_1 DUF63 -------------------------------TIFAIAGVALFA-------------------------GKLA----ENA--------------------- 111
PKL62929_1 hypotheti -------------------------------TIFAVAASALFL-------------------------GKLA----ENA--------------------- 111
WP_004039890_1 DUF63 -------------------------------VVFFITVGALFL-------------------------SRLL----EVK--------------------- 111
WP_067049225_1 DUF63 -------------------------------VVFFITVIALFF-------------------------SRIL----EVK--------------------- 111
KYK37884_1 hypotheti -------------------------------AIFGITFTALIV-------------------------TVHI---------------------------- 104
KYK28019_1 hypotheti -------------------------------LIFGITFTSLIV-------------------------TVRI---------------------------- 104
OYT57835_1 hypotheti -------------------------------VTALVAIVALFV-------------------------SRLT----EKS--------------------- 111
OYT33350_1 hypotheti -------------------------------IVFLICFPTLVI-------------------------SIKT---------------------------- 111
WP_012964745_1 DUF63 -------------------------------LVFSIAFPTLLI-------------------------ALKM---------------------------- 108
WP_048091573_1 DUF63 -------------------------------LIFLITFSSLMI-------------------------ALRT---------------------------- 108
WP_048096399_1 DUF63 -------------------------------LTFIITFAALII-------------------------SLRT---------------------------- 108
```

```
WP_012940099_1 DUF63 ------------------------------VVFCIAFPSLLL-------------------------NLKL---------------------------- 107
WP_010877969_1 DUF63 ------------------------------LIFAIAFPTLLL-------------------------SLRF---------------------------- 120
WP_013682839_1 DUF63 ------------------------------VIFCIAFPTLVL-------------------------SIRL---------------------------- 108
WP_015591141_1 DUF63 ------------------------------VIFLIAFPTLVI-------------------------SLRK---------------------------- 108
WP_012035887_1 DUF63 ------------------------------LAFVITAAVLLI-------------------------CLGL----QRA--------------------- 118
WP_014404626_1 DUF63 ------------------------------LAFLITLLALLV-------------------------CLYL----ERS--------------------- 112
BAI60262_1 conserved ------------------------------LAFLITLAVLLI-------------------------CVYL----ERA--------------------- 112
WP_042684156_1 DUF63 ------------------------------VVFAVAIVLLAA-------------------------CVFV---------------------------- 117
OFV68024_1 membrane  ------------------------------LVFVVVLTILLV-------------------------SIRL---------------------------- 75
WP_013720253_1 DUF63 ------------------------------LVFLVTAGSLTL-------------------------TRRI---------------------------- 110
WP_014587756_1 DUF63 ------------------------------LAAAGTMTALLL-------------------------CRQA---------------------------- 107
ABK14947_1 Protein o ------------------------------LVASVTLLALFI-------------------------SRRL---------------------------- 114
OKY79140_1 putative  ------------------------------LIAGIVISILLA-------------------------KYLI----RKT--------------------- 114
WP_086637003_1 DUF63 ------------------------------LVAFITLLVLVV-------------------------VSFS---RKKE--------------------- 115
WP_048089097_1 DUF63 ------------------------------SMFAVTVILLTL-------------------------AVTL----ERK--------------------- 115
WP_097298272_1 DUF63 ------------------------------FMFAVTVSILAI-------------------------AIYL----ERK--------------------- 116
WP_013897788_1 DUF63 ------------------------------LVFIVTLTFLVI-------------------------SKKM----EIM--------------------- 117
WP_015323982_1 DUF63 ------------------------------VVFLITLLCLIV-------------------------AKML----SKK--------------------- 117
WP_011499115_1 DUF63 ------------------------------FVFVVTVFCLLL-------------------------SRSI----ASS--------------------- 117
WP_013037846_1 DUF63 ------------------------------VVFVVTLACLVV-------------------------SKKL----YDL--------------------- 117
WP_048205640_1 DUF63 ------------------------------VVFVATIICLLF-------------------------SRWL----YSS--------------------- 117
WP_072561629_1 DUF63 ------------------------------VVFVVTLACLVV-------------------------SKKL----YDL--------------------- 117
WP_072360082_1 DUF63 ------------------------------VVFVVTLACLVV-------------------------SKKL----YDL--------------------- 117
WP_096711793_1 DUF63 ------------------------------VVFVVTLACLVV-------------------------SKKL----YDL--------------------- 117
ODV50598_1 hypotheti ------------------------------VVFVVTLACLVV-------------------------SKKL----YDL--------------------- 117
WP_013194042_1 DUF63 ------------------------------LVFAIVILTLVV-------------------------SKWL----YNK--------------------- 117
WP_048178067_1 DUF63 ------------------------------LVFAVTVFCLWL-------------------------AIRL----EKA--------------------- 119
WP_048127111_1 DUF63 ------------------------------LVFGITVACLWL-------------------------SIRL----QKA--------------------- 119
WP_011023594_1 DUF63 ------------------------------LVFGVTVICLWL-------------------------SIRL----QKA--------------------- 123
WP_048184587_1 DUF63 ------------------------------LVFGVTIICLWL-------------------------SIKL----QKA--------------------- 119
WP_011032542_1 DUF63 ------------------------------LVFGITVICLWL-------------------------SIRL----QKA--------------------- 119
WP_011032542_1 DUF63 ------------------------------LVFGITVICLWL-------------------------SIRL----QKA--------------------- 119
WP_048129141_1 DUF63 ------------------------------LVFGVTVICLWL-------------------------SIRL----QKA--------------------- 119
WP_048129141_1 DUF63 ------------------------------LVFGVTVICLWL-------------------------SIRL----QKA--------------------- 119
WP_048169891_1 DUF63 ------------------------------LVFGITVVCLWL-------------------------SIRL----QKA--------------------- 119
WP_048137175_1 DUF63 ------------------------------LVFGITVVCLWL-------------------------SIRL----QKA--------------------- 119
WP_048137175_1 DUF63 ------------------------------LVFGITVVCLWL-------------------------SIRL----QKA--------------------- 119
WP_048137175_1 DUF63 ------------------------------LVFGITVVCLWL-------------------------SIRL----QKA--------------------- 119
WP_048137694_1 DUF63 ------------------------------LVFGITVVCLWL-------------------------SIRI----QKA--------------------- 119
WP_048168126_1 DUF63 ------------------------------LVFAITVGCLWI-------------------------SIRL----QKA--------------------- 119
WP_048118082_1 DUF63 ------------------------------LVFAITISCLWI-------------------------SIRM----QKA--------------------- 119
WP_048158120_1 DUF63 ------------------------------LVFAITISCLWI-------------------------SIRM----QKA--------------------- 119
WP_011305329_1 DUF63 ------------------------------MVFAITVGCLWI-------------------------SIRM----QKA--------------------- 119
WP_054298619_1 DUF63 ------------------------------LVFAITVGCLWI-------------------------SIRL----QKA--------------------- 119
ALK05385_1 hypotheti ------------------------------LVFAITVGCLWI-------------------------SIRL----QKA--------------------- 119
WP_015052897_1 DUF63 ------------------------------LVFFVTVIFLII-------------------------AKLV----SKV--------------------- 117
WP_023846134_1 DUF63 ------------------------------VVFAITVFFLVL-------------------------SRWI----VKL--------------------- 116
OIN88451_1 hypotheti -------------------------------LFVFLINSLLV-------------------------ASKL---IEKK--------------------- 108
PIX50278_1 hypotheti -------------------------------LFVFLINSLLV-------------------------ASKL----IEK--------------------- 107
PIW41402_1 hypotheti -------------------------------LFVFLINSLLV-------------------------ASKL----IEK--------------------- 107
PIY35178_1 hypotheti -------------------------------LFVFLINSLLV-------------------------ASKL----IEK--------------------- 107
PJB74886_1 hypotheti -------------------------------LFVFLINSLLV-------------------------ASKL----IEK--------------------- 107
PIZ33651_1 hypotheti -------------------------------LFVFLINSLLV-------------------------ASKL----IEK--------------------- 107
WP_048165029_1 DUF63 ------------------------------MTFIITFSALLI-------------------------AIKA---------------------------- 107
WP_042681046_1 DUF63 ------------------------------LTFAITFTALLI-------------------------THKL---------------------------- 107
WP_013467319_1 DUF63 ------------------------------VTFAITFTALLI-------------------------THKL---------------------------- 107
WP_048152160_1 DUF63 ------------------------------STFIITFSALLI-------------------------THRA---------------------------- 107
WP_015849008_1 DUF63 ------------------------------LVFMITFSSLLI-------------------------THKF---------------------------- 107
WP_004069276_1 DUF63 ------------------------------LVFAITFSALLI-------------------------THKF---------------------------- 107
WP_058946638_1 DUF63 ------------------------------LVFAITFSALLI-------------------------THKF---------------------------- 107
WP_042701551_1 DUF63 ------------------------------LVFAITFSALLI-------------------------THKL---------------------------- 107
WP_055282692_1 DUF63 ------------------------------LVFTITFSALLI-------------------------THKL---------------------------- 107
WP_013906023_1 DUF63 ------------------------------LVFVITFSALLV-------------------------THRL---------------------------- 106
WP_011013158_1 DUF63 ------------------------------LVFVIAFSALLV-------------------------SHKF---------------------------- 106
WP_014733166_1 DUF63 ------------------------------LVFSIAFPALII-------------------------CKKF---------------------------- 106
WP_068322889_1 DUF63 ------------------------------LVFSIAFPALII-------------------------CKKS---------------------------- 106
WP_068575625_1 DUF63 ------------------------------LVFSIAFPALII-------------------------SKKF---------------------------- 106
WP_010885960_1 DUF63 ------------------------------LVSLIATPALLI-------------------------PYKF---------------------------- 106
WP_010867401_1 DUF63 ------------------------------LVFSIAFPALLI-------------------------SHRF---------------------------- 106
WP_013747987_1 DUF63 ------------------------------LVFSIAFPALII-------------------------SHRF---------------------------- 107
WP_068664049_1 DUF63 ------------------------------MVTAFTSLALYV-------------------------SHRH---------------------------- 107
WP_011249739_1 DUF63 ------------------------------VIAAFAIASLFA-------------------------VWRH---------------------------- 107
WP_062386854_1 DUF63 ------------------------------VIAAFAIASLFA-------------------------VWRH---------------------------- 107
WP_050003781_1 DUF63 ------------------------------VIAGFAIASLLT-------------------------VWRH---------------------------- 107
WP_042690699_1 DUF63 ------------------------------VIAGFAIASLFA-------------------------VWRH---------------------------- 107
WP_088885212_1 DUF63 ------------------------------VIAGFAIAALFI-------------------------VWRH---------------------------- 107
WP_010478519_1 DUF63 ------------------------------VIAGFTIAALFV-------------------------VWRH---------------------------- 107
WP_088858240_1 DUF63 ------------------------------VIAAFAIAALFV-------------------------VWRH---------------------------- 107
WP_015859016_1 DUF63 ------------------------------VIAAFAIASLLV-------------------------VWRH---------------------------- 107
WP_014121940_1 DUF63 ------------------------------VIAAFAIASLLI-------------------------VWRH---------------------------- 107
WP_062373911_1 DUF63 ------------------------------VIAAFAIAALLM-------------------------VWRH---------------------------- 107
WP_088882924_1 DUF63 ------------------------------VIAAFAIASLY--------------------------AVWI---------------------------- 106
WP_088862911_1 DUF63 ------------------------------VIATFAIAALF--------------------------AVWI---------------------------- 106
WP_012572006_1 DUF63 ------------------------------VIAAFALASLYV-------------------------VWRH---------------------------- 107
WP_088854707_1 DUF63 ------------------------------VIAAFAIASLYV-------------------------VWKH---------------------------- 107
WP_014789474_1 DUF63 ------------------------------VIAAFAIASLFA-------------------------VWKH---------------------------- 107
WP_088180728_1 DUF63 ------------------------------VIAAFAIASLYV-------------------------VWRH---------------------------- 107
WP_088864937_1 DUF63 ------------------------------VIAAFAIASLYV-------------------------VWRH---------------------------- 107
WP_055429686_1 DUF63 ------------------------------VIAAFAIASLYV-------------------------VWRH---------------------------- 107
WP_088865932_1 DUF63 ------------------------------VIAAFAIASLYV-------------------------VWRH---------------------------- 107
WP_014013642_1 DUF63 ------------------------------VIAAFAIASLYI-------------------------VWRH---------------------------- 107
WP_088856804_1 DUF63 ------------------------------VIATFAIASLYV-------------------------VWRH---------------------------- 107
WP_058939475_1 DUF63 ------------------------------VIAAFAIASLYI-------------------------VWRH---------------------------- 107
EHR77276_1 conserved ------------------------------HLAAWLIGIAIL-------------------------SQWL---AGKWDGATTDRDEQKVRLALLPSLM 189
OUV40025_1 hypotheti ------------------------------HLAVWLIGIGIV-------------------------SHLV---------------------------- 165
```

```
MBJ52984_1 hypotheti SGSIGIVATKNLTDKSNNEIFSRMINRIRFCTLFMLLLFYLIIFEPSLVNHSNLTWFSIPITIIVSVAVAWFSPSIPKL--------------------- 229
PDH23744_1 hypotheti -------------------------------QAAFWVVLAGAR-------------------------GISL----DRA--------------------- 160
PDH25468_1 hypotheti ISMVLISSQFLIYGISISGSNTVESKEIDLTLFAIFGILGVLT-------------------------PIWL---KDIG--------------------- 224

WP_048201856_1 hypot ---------------------------------------------------------------------------------------------------- 96
WP_048150344_1 hypot -----------------------------------------GHKKSYRE--------------------------------------------------- 112
AOV95360_1 hypotheti ---------------------------------------------AYLF---------------------------------EA--VTGL---------- 99
EOD42420_1 Uncharact ---------------------------------------------KYRK----------------------------------Y--YLYI---------- 115
AOV95164_1 hypotheti -----------------------------------------ASESDCTR----------------------------------EELLKNL---------- 137
EGQ40074_1 putative  --------------------------------------------GSQDR----------------------------------A--LGSM---------- 136
KYK23067_1 hypotheti --------FSPKKTLLILLFIFLLIDSVYSILWVLGFDYGASIIEPWFF---------------------------------CILSFFAFLPLFYRFLKK 221
WP_084383883_1 DUF63 -----------------------------------------YSFEAYRR-------------------------------------TGTY---------- 175
WP_049984677_1 DUF63 -----------------------------------------GFLSIPGT-------------------------------------VGVV---------- 179
WP_103428047_1 hypot -----------------------------------------GLVSIPQV-------------------------------------VGAV---------- 178
WP_004048647_1 DUF63 ------------------------------------------NTGSIPS----------------------------------T--VGLI---------- 178
WP_004594600_1 DUF63 -----------------------------------------QNIGSIPL----------------------------------T--VGLV---------- 178
WP_004594600_1 DUF63 -----------------------------------------QNIGSIPL----------------------------------T--VGLV---------- 178
WP_050050397_1 DUF63 -----------------------------------------QNIGSIPL----------------------------------T--VGLV---------- 178
WP_004594600_1 DUF63 -----------------------------------------QNIGSIPL----------------------------------T--VGLV---------- 178
WP_050050397_1 DUF63 -----------------------------------------QNIGSIPL----------------------------------T--VGLV---------- 178
WP_049983690_1 DUF63 ------------------------------------------NIGSIPS----------------------------------I--VGLV---------- 178
WP_103428162_1 hypot ------------------------------------------DIGSVPS----------------------------------T--VGLV---------- 178
WP_009378268_1 DUF63 ------------------------------------------DIGSIPS----------------------------------T--VGLV---------- 178
WP_015763176_1 DUF63 -----------------------------------------GQIDDGNR----------------------------------A--LGVV---------- 211
WP_018259238_1 DUF63 -----------------------------------------GQIDDGNR----------------------------------A--LGVV---------- 211
WP_004592615_1 DUF63 -----------------------------------------GVVDSYYR----------------------------------T--TGAI---------- 205
WP_004518092_1 DUF63 -----------------------------------------GVVDSYYR----------------------------------T--TGAI---------- 205
WP_014040450_1 DUF63 -----------------------------------------GIVDSYYR----------------------------------T--TGAV---------- 205
WP_014040450_1 DUF63 -----------------------------------------GIVDSYYR----------------------------------T--TGAV---------- 205
WP_008309018_1 DUF63 -----------------------------------------GVVDSYYW----------------------------------T--TGAV---------- 205
WP_014040450_1 DUF63 -----------------------------------------GIVDSYYR----------------------------------T--TGAV---------- 205
WP_014040450_1 DUF63 -----------------------------------------GIVDSYYR----------------------------------T--TGAV---------- 205
WP_005534235_1 DUF63 -----------------------------------------GIIDSYYR----------------------------------T--TGAI---------- 205
WP_004961153_1 DUF63 -----------------------------------------GVVDSYYR----------------------------------T--TGAI---------- 205
WP_004961153_1 DUF63 -----------------------------------------GVVDSYYR----------------------------------T--TGAI---------- 205
WP_101350151_1 hypot -----------------------------------------GVIDSYYR----------------------------------T--TGAI---------- 205
WP_053968273_1 DUF63 -----------------------------------------GVIDSYYR----------------------------------T--TGAI---------- 205
WP_058995687_1 DUF63 -----------------------------------------GVVDSYYR----------------------------------T--TGAI---------- 205
WP_015790673_1 DUF63 -----------------------------------------GVVASAER----------------------------------A--LGTV---------- 206
WP_008524115_1 DUF63 -----------------------------------------GVVDSADR----------------------------------T--LGTV---------- 206
WP_075936143_1 DUF63 -----------------------------------------GYVERYGR----------------------------------P--LFAT---------- 199
WP_020446311_1 DUF63 -----------------------------------------GRVDRWER----------------------------------P--FGAA---------- 201
WP_049898450_1 DUF63 -----------------------------------------GMVAEYYG----------------------------------A--LAAF---------- 203
WP_006077061_1 DUF63 -----------------------------------------DIVEEYYG----------------------------------A--LAAF---------- 203
WP_049996700_1 DUF63 -----------------------------------------DVVSEFEP----------------------------------P--LAAM---------- 203
EMA38628_1 hypotheti -----------------------------------------GIVGRYED----------------------------------A--LAGF---------- 203
WP_049992561_1 DUF63 -----------------------------------------DYVDSYPK----------------------------------T--LGII---------- 204
WP_010903025_1 DUF63 -----------------------------------------DAVETYHR----------------------------------P--LAGI---------- 202
WP_009760947_1 DUF63 -----------------------------------------DIAEDYYR----------------------------------P--LAAA---------- 201
WP_059057097_1 DUF63 -----------------------------------------TVVEDYYR----------------------------------P--LAGA---------- 201
WP_058983522_1 DUF63 -----------------------------------------TVFEDYYR----------------------------------P--LAAT---------- 201
WP_071932813_1 DUF63 -----------------------------------------GHADSFEW----------------------------------P--LFGF---------- 197
WP_050048584_1 DUF63 -----------------------------------------GTTETYEW----------------------------------P--LAGF---------- 197
WP_014051341_1 DUF63 -----------------------------------------GYTDRYER----------------------------------P--LAAI---------- 200
WP_079233299_1 DUF63 -----------------------------------------GYVDRYEY----------------------------------P--LGAM---------- 199
WP_053948572_1 DUF63 -----------------------------------------EYVDDYVR----------------------------------P--LTAM---------- 199
WP_049980835_1 DUF63 -----------------------------------------GIVDDYVR----------------------------------P--LAAM---------- 199
KPN31749_1 hypotheti -----------------------------------------GYVDDYVR----------------------------------P--LTAM---------- 149
AGB16040_1 putative  -----------------------------------------EYVSGYEY----------------------------------P--LAGI---------- 198
WP_007696755_1 DUF63 -----------------------------------------GVIPGYEY----------------------------------G--LAGI---------- 198
WP_015322904_1 DUF63 -----------------------------------------EYVSGYEY----------------------------------P--LFGV---------- 198
WP_076581869_1 DUF63 -----------------------------------------DYVSGYEY----------------------------------P--LFGI---------- 198
WP_005559622_1 DUF63 -----------------------------------------EYVSGYEY----------------------------------P--LFGV---------- 198
WP_006067640_1 DUF63 -----------------------------------------DSVSRYEY----------------------------------P--LFGI---------- 193
WP_008164554_1 DUF63 -----------------------------------------GSVSRYEY----------------------------------P--LFGI---------- 198
WP_006088112_1 DUF63 -----------------------------------------DHVSGYEY----------------------------------P--LFGV---------- 198
WP_012943247_1 DUF63 -----------------------------------------DRVSGYEY----------------------------------P--LAAI---------- 198
WP_008895026_1 DUF63 -----------------------------------------DRVSGYEY----------------------------------P--LAAI---------- 198
WP_098727043_1 DUF63 -----------------------------------------DYVSGYEY----------------------------------P--LAGI---------- 198
WP_049990861_1 DUF63 -----------------------------------------DYVSGYEY----------------------------------P--LAGV---------- 196
WP_008013001_1 DUF63 -----------------------------------------DYVSRYEY----------------------------------P--LAGI---------- 198
WP_006180566_1 DUF63 -----------------------------------------DYVSGYEY----------------------------------P--LAGI---------- 198
WP_006650865_1 DUF63 -----------------------------------------DYVSGYEY----------------------------------P--LAGI---------- 198
WP_066301295_1 DUF63 -----------------------------------------DYVSGYEY----------------------------------P--LAGI---------- 198
WP_049966804_1 DUF63 -----------------------------------------DYVSGYEY----------------------------------P--LAGI---------- 198
WP_076145380_1 DUF63 -----------------------------------------DYVSGYEY----------------------------------P--LAGI---------- 198
WP_097378845_1 DUF63 -----------------------------------------DYVSGYEY----------------------------------P--LAGI---------- 198
WP_008452261_1 DUF63 -----------------------------------------GYVSGYEY----------------------------------P--LAGI---------- 198
WP_008452261_1 DUF63 -----------------------------------------GYVSGYEY----------------------------------P--LAGI---------- 198
WP_006432643_1 DUF63 -----------------------------------------GHVSGYEY----------------------------------P--LAGI---------- 198
WP_008452261_1 DUF63 -----------------------------------------GYVSGYEY----------------------------------P--LAGI---------- 198
WP_007109662_1 DUF63 -----------------------------------------GYVSGYEY----------------------------------P--LAGI---------- 198
WP_049952821_1 DUF63 -----------------------------------------GYVSGYEY----------------------------------P--LAGI---------- 198
WP_086889541_1 DUF63 -----------------------------------------GTVSGYEY----------------------------------P--LAAI---------- 201
WP_005580009_1 DUF63 -----------------------------------------DYVPGYEY----------------------------------P--LAAI---------- 201
WP_049927356_1 DUF63 -----------------------------------------DVVSGYEY----------------------------------P--LFGI---------- 198
WP_087714455_1 DUF63 -----------------------------------------EYVSGYEY----------------------------------P--LFGI---------- 198
WP_013878438_1 DUF63 -----------------------------------------DYVSGYEY----------------------------------P--LFAV---------- 198
WP_049921111_1 DUF63 -----------------------------------------GYVSGYEY----------------------------------P--LAAI---------- 198
WP_007142789_1 DUF63 -----------------------------------------DYVSGYEY----------------------------------P--LFGI---------- 197
WP_006651941_1 DUF63 -----------------------------------------DYVPGYEY----------------------------------P--LGAI---------- 198
WP_004216437_1 DUF63 -----------------------------------------DYVPGYEY----------------------------------P--LGAI---------- 198
WP_071402513_1 DUF63 -----------------------------------------DYVPGYEY----------------------------------P--LGAI---------- 198
WP_006666462_1 DUF63 -----------------------------------------DYVSGYEY----------------------------------P--LGAI---------- 198
WP_049904535_1 DUF63 -----------------------------------------DYVSGYEY----------------------------------P--LGAI---------- 198
```

```
WP_006824405_1 DUF63 -----------------------------------------DYVSGYEY----------------------------------P--LGAI---------- 198
WP_011323802_1 DUF63 -----------------------------------------GTLSRSET----------------------------------F--LGGI---------- 199
WP_015409719_1 DUF63 -----------------------------------------GTIDAYEP----------------------------------Y--VAAT---------- 200
WP_006883947_1 DUF63 -----------------------------------------ELVRNRYR----------------------------------A--LAGF---------- 199
WP_077207545_1 DUF63 -----------------------------------------GYVETYAR----------------------------------P--MAAV---------- 202
ESS12903_1 putative  -----------------------------------------GLVSRYEY----------------------------------P--LAAV---------- 202
WP_008417361_1 DUF63 -----------------------------------------GAVSRYEY----------------------------------P--LAAI---------- 201
WP_066381608_1 DUF63 -----------------------------------------DLVERYEY----------------------------------P--LAAI---------- 196
WP_049947642_1 DUF63 -----------------------------------------GRFDGFYR----------------------------------P--LLVV---------- 202
ESS05953_1 putative  -----------------------------------------GLVRSHHR----------------------------------A--LSGI---------- 196
WP_096390051_1 DUF63 -----------------------------------------GLVDDYAR----------------------------------P--LFAM---------- 208
WP_021072749_1 DUF63 -----------------------------------------GLVDDYAR----------------------------------P--LFAM---------- 208
WP_049982791_1 DUF63 -----------------------------------------GIVDDYAR----------------------------------P--LFAM---------- 206
WP_008585990_1 DUF63 -----------------------------------------GVVDDYAR----------------------------------P--LFAS---------- 208
WP_006628721_1 DUF63 -----------------------------------------GVVDDYAR----------------------------------P--LFAS---------- 206
WP_006113218_1 DUF63 -----------------------------------------GIVDDYAR----------------------------------P--LFAS---------- 207
WP_049930034_1 DUF63 -----------------------------------------GAVDDYAR----------------------------------P--LFGS---------- 208
WP_049906047_1 DUF63 -----------------------------------------GVVDDYAR----------------------------------P--LFAS---------- 208
WP_044965494_1 DUF63 -----------------------------------------GVVDDYAR----------------------------------P--LFAS---------- 208
WP_096393195_1 DUF63 -----------------------------------------DIVDDYAR----------------------------------P--LFAS---------- 208
WP_049908400_1 DUF63 -----------------------------------------GVVDDYAR----------------------------------P--LFGS---------- 208
WP_049908585_1 DUF63 -----------------------------------------GVVEDYAR----------------------------------P--LFAS---------- 208
WP_049902631_1 DUF63 -----------------------------------------GVVDDYAR----------------------------------P--LFAS---------- 208
WP_049983668_1 DUF63 -----------------------------------------GVVDDYAR----------------------------------P--LFAS---------- 207
WP_053772267_1 DUF63 -----------------------------------------GVVDDYAR----------------------------------P--LFAL---------- 207
WP_007999151_1 DUF63 -----------------------------------------GVVDDYAR----------------------------------P--LFAF---------- 206
WP_015909917_1 DUF63 -----------------------------------------GVVDDYAR----------------------------------P--LFAM---------- 207
WP_004050594_1 DUF63 -----------------------------------------GVVDDYAR----------------------------------P--LFAM---------- 207
WP_095636035_1 DUF63 -----------------------------------------GVTDDYAR----------------------------------P--LFAM---------- 207
WP_008003811_1 DUF63 -----------------------------------------GVVDDYAR----------------------------------P--LFAM---------- 207
WP_066416034_1 DUF63 -----------------------------------------GIVDDYAP----------------------------------P--LFGA---------- 208
ESS03170_1 putative  -----------------------------------------GLVDDYAK----------------------------------P--LFGS---------- 207
WP_089671383_1 DUF63 -----------------------------------------GYVDDYAY----------------------------------P--LAGI---------- 206
ERH07656_1 putative  -----------------------------------------GVVDAYEY----------------------------------P--VAAI---------- 206
ESS10096_1 putative  -----------------------------------------GVVDAYEY----------------------------------P--VAAI---------- 206
ESS07824_1 putative  ---------------------------------------------------------------------------------------------------- 173
ERH05689_1 putative  -----------------------------------------GLVERYEH----------------------------------A--VFAI---------- 206
ERH02266_1 putative  -----------------------------------------GLVERYEH----------------------------------A--VFAI---------- 206
ESS07825_1 putative  --------------------------------------------------------------------------------------VLGG---------- 34
WP_049970017_1 DUF63 -----------------------------------------GILDGYEY----------------------------------S--LAGI---------- 200
WP_007979289_1 DUF63 -----------------------------------------GVFEDYER----------------------------------P--LAGI---------- 200
WP_007977745_1 DUF63 -----------------------------------------GVVEDYEY----------------------------------P--LAGI---------- 200
WP_014556445_1 DUF63 -----------------------------------------GLVERYDR----------------------------------L--LFIT---------- 201
WP_049935775_1 DUF63 -----------------------------------------GVVERYTR----------------------------------P--LFGS---------- 199
WP_008325149_1 DUF63 -----------------------------------------GLIDDYVK----------------------------------P--LFGA---------- 201
WP_007543631_1 DUF63 -----------------------------------------GLIDDYVK----------------------------------P--LFGA---------- 201
WP_049905041_1 DUF63 -----------------------------------------GLVDDYVK----------------------------------P--LFGA---------- 201
WP_049913563_1 DUF63 -----------------------------------------GLVDDYVK----------------------------------P--LFVA---------- 201
WP_049967851_1 DUF63 -----------------------------------------GLVDDYVK----------------------------------P--LFVA---------- 201
WP_049914905_1 DUF63 -----------------------------------------GLVDDYVK----------------------------------P--LFGA---------- 201
WP_049916430_1 DUF63 -----------------------------------------GLVDDYVK----------------------------------P--LFGA---------- 201
WP_049896892_1 DUF63 -----------------------------------------GIVDDYVK----------------------------------P--LFGA---------- 201
WP_049896892_1 DUF63 -----------------------------------------GIVDDYVK----------------------------------P--LFGA---------- 201
WP_049896892_1 DUF63 -----------------------------------------GIVDDYVK----------------------------------P--LFGA---------- 201
WP_058828253_1 DUF63 -----------------------------------------GLVDDYVK----------------------------------P--LFGA---------- 201
WP_058568959_1 DUF63 -----------------------------------------GLVDDYVK----------------------------------P--LFGA---------- 201
WP_049917947_1 DUF63 -----------------------------------------GLVDDYVK----------------------------------P--LFGA---------- 201
WP_049920348_1 DUF63 -----------------------------------------GLVDDYVK----------------------------------P--LFGA---------- 201
WP_089777545_1 DUF63 -----------------------------------------GLVDDYVK----------------------------------P--LFGA---------- 201
WP_008320633_1 DUF63 -----------------------------------------GVIGDYMK----------------------------------L--LFGV---------- 201
WP_004060355_1 DUF63 -----------------------------------------GIVDNYMK----------------------------------P--LLGA---------- 201
WP_103426078_1 hypot -----------------------------------------GSVDDYQR----------------------------------P--LFVL---------- 201
WP_009367433_1 DUF63 -----------------------------------------GTIDDYTK----------------------------------P--LFGF---------- 200
WP_013440552_1 DUF63 -----------------------------------------DVTDSYER----------------------------------P--LFAI---------- 201
WP_049916626_1 DUF63 -----------------------------------------GTVADYER----------------------------------P--LFGF---------- 201
WP_058582837_1 DUF63 -----------------------------------------GVFDEFYR----------------------------------P--LAVV---------- 203
WP_101298124_1 hypot -----------------------------------------GVIDRYER----------------------------------G--VFWA---------- 203
PIN95153_1 hypotheti -----------------------------------------------------------------------------------VQLMATI---------- 73
PIU22108_1 hypotheti ----------------------------------------YGVNQKY------------------------------------L--LFSI---------- 131
AAR39201_1 NEQ352    -----------------------------------------------------------------------------------P--IIQW---------- 85
OIR14399_1 hypotheti -----------------------------------SKDHKRLVYSTYLL----------------------------------L--IVGY---------- 164
OIR20963_1 hypotheti -------------------------EIDSKTKAIGLISFAIGSYGIWWYFAPGEWIHPTSWVLIVLSAAALTAEFLRSKPLKDPVIFFGIASTLLVILAY 224
OIR22371_1 hypotheti ---------------DLPSLTKGLALVYFTIGGYGLWWYFAPG--DWIH----------------------------------PSSWALI---------- 199
EGQ43935_1 putative  ------------------------------------------TQKPYYK----------------------------------T--LAVA---------- 126
MAG21679_1 hypotheti -----------------------------------------ESKTKYSF----------------------------------HLLFAAT---------- 131
PIN85618_1 hypotheti ------------------------------------------DESKFAK----------------------------------I--FGGI---------- 131
PIN99249_1 hypotheti ------------------------------------------LKKEPLK----------------------------------I--FMGI---------- 135
AJF59838_1 hypotheti ------------------------------------------IKQETIK----------------------------------V--FGAI---------- 131
WP_042682145_1 DUF63 --------------------------------------------KTYPK----------------------------------V--TIAW---------- 120
WP_013468033_1 DUF63 --------------------------------------------KTYPK----------------------------------V--TIAW---------- 120
WP_055281581_1 DUF63 --------------------------------------------KTYPK----------------------------------I--TIGW---------- 119
WP_048160382_1 DUF63 --------------------------------------------KTYPK----------------------------------I--TVGW---------- 119
WP_042701216_1 DUF63 --------------------------------------------RTYPK----------------------------------I--TVAW---------- 119
WP_004068659_1 DUF63 --------------------------------------------KMYPK----------------------------------I--TVAW---------- 119
WP_058946665_1 DUF63 --------------------------------------------KTYPK----------------------------------I--TVAW---------- 119
WP_014835499_1 DUF63 --------------------------------------------RLYPK----------------------------------L--TILW---------- 119
WP_010884243_1 DUF63 --------------------------------------------KLYPK----------------------------------L--TIAW---------- 122
WP_013748129_1 DUF63 --------------------------------------------KLYPK----------------------------------L--TIAW---------- 118
WP_010867251_1 DUF63 --------------------------------------------KLYPK----------------------------------L--TIAW---------- 118
WP_014733359_1 DUF63 --------------------------------------------RLYPK----------------------------------L--TFAW---------- 119
WP_068319891_1 DUF63 --------------------------------------------KTYPK----------------------------------L--TIAW---------- 119
WP_068576049_1 DUF63 --------------------------------------------KTYPK----------------------------------L--TIAW---------- 119
WP_014013692_1 DUF63 --------------------------------------------KTYPK----------------------------------L--TFGW---------- 124
WP_014789520_1 DUF63 --------------------------------------------KTYPK----------------------------------L--TFGW---------- 124
WP_088180676_1 DUF63 --------------------------------------------KTYPK----------------------------------L--TFGW---------- 124
```

WP_088864885_1 DUF63 --------------------------------------------KTYPK----------------------------------L--TFGW---------- 124
WP_088856755_1 DUF63 --------------------------------------------KLYPR----------------------------------L--TFGW---------- 124
WP_088865981_1 DUF63 --------------------------------------------KTYPK----------------------------------L--TLGW---------- 124
WP_013906334_1 DUF63 --------------------------------------------ETYPR----------------------------------I--TIGW---------- 120
WP_088862658_1 DUF63 --------------------------------------------GTYPK----------------------------------V--TIAW---------- 123
WP_088882956_1 DUF63 --------------------------------------------GTYPK----------------------------------I--TTAW---------- 123
WP_012571938_1 DUF63 --------------------------------------------KTYPK----------------------------------I--TVAW---------- 120
WP_068663933_1 DUF63 --------------------------------------------KTYPK----------------------------------I--NVAW---------- 120
WP_074631153_1 DUF63 --------------------------------------------EMYPR----------------------------------I--TIAW---------- 120
WP_058939074_1 DUF63 --------------------------------------------GTYPK----------------------------------I--TTAW---------- 124
WP_088854681_1 DUF63 --------------------------------------------KSYPK----------------------------------V--TIAW---------- 124
WP_088885426_1 DUF63 --------------------------------------------GTYPK----------------------------------I--TIGW---------- 120
WP_011249694_1 DUF63 --------------------------------------------GMYPK----------------------------------I--TVAW---------- 122
WP_062386972_1 DUF63 --------------------------------------------GMYPK----------------------------------I--TVAW---------- 122
WP_010478887_1 DUF63 --------------------------------------------KTYPK----------------------------------V--TVVW---------- 120
WP_088858605_1 DUF63 --------------------------------------------KTYPK----------------------------------I--TIAW---------- 120
WP_042690028_1 DUF63 --------------------------------------------RTYPK----------------------------------V--TIVW---------- 122
WP_048150762_1 DUF63 --------------------------------------------KTYPK----------------------------------I--TIAW---------- 120
WP_048811076_1 DUF63 --------------------------------------------KTYPK----------------------------------I--TIVW---------- 120
WP_050003256_1 DUF63 --------------------------------------------KTYPK----------------------------------I--TIAW---------- 122
WP_062372100_1 DUF63 --------------------------------------------KTYPK----------------------------------I--TIAW---------- 120
WP_048165284_1 DUF63 --------------------------------------------KKYPK----------------------------------I--TVSW---------- 122
WP_048148750_1 DUF63 --------------------------------------------NLYPK----------------------------------L--TVAW---------- 122
OYT53462_1 hypotheti ------------------------------------------TGLGYWI----------------------------------P--PFLL---------- 105
KYC51152_1 hypotheti ------------------------------------------NILSYWK----------------------------------L--PFAV---------- 105
KYC46003_1 hypotheti -----------------------------------------NILSYWKL----------------------------------PFAVGIL---------- 108
KYC48643_1 hypotheti -----------------------------------------NILSYWKL----------------------------------PFAVGIL---------- 108
KYC55321_1 hypotheti ------------------------------------------DILSYWK----------------------------------L--TFAV---------- 105
KYC57927_1 hypotheti ------------------------------------------DILSYWK----------------------------------L--TFAV---------- 105
KYC57171_1 hypotheti ------------------------------------------DILSYWK----------------------------------L--TFAV---------- 105
OIO20701_1 hypotheti ------------------------------------------SRLQYAK-------------------------------------FV------------ 151
PIT83986_1 hypotheti ---------------------------------------------GARY----------------------------------F--AGAC---------- 168
OIO24760_1 hypotheti ------------------------------------------PQNEFHR----------------------------------T--LRAG---------- 131
OIO24701_1 hypotheti -----------------------------------------TGIPLWKT-------------------------------------VGGI---------- 133
OIO27021_1 hypotheti -------------------------------------------KGDWLR----------------------------------N--AFGL---------- 126
OIO26762_1 hypotheti -----------------------------------------------------------------------------------PANAQMA---------- 123
PIN95811_1 hypotheti -----------------------------------------------------------------------------------PANAQMA---------- 123
PIO01637_1 hypotheti -----------------------------------------------------------------------------------PANAQMA---------- 123
PIO02820_1 hypotheti -------------------------------------------KGDWLR----------------------------------N--AFGL---------- 126
PJD01038_1 hypotheti -------------------------------------------KGDWLR----------------------------------N--AFGL---------- 126
PIZ91366_1 hypotheti ---------------------------------------------PANA----------------------------------Q--MAGW---------- 125
WP_013100570_1 DUF63 ------------------------------------------FRDNYYK----------------------------------V--SSVV---------- 121
WP_004590770_1 DUF63 ------------------------------------------FKEKYYK----------------------------------V--SAVI---------- 121
WP_048196979_1 DUF63 ------------------------------------------FKENYYK----------------------------------V--SAII---------- 121
WP_015791538_1 DUF63 ------------------------------------------FKKDYYK----------------------------------V--SAVI---------- 121
WP_048202292_1 DUF63 ------------------------------------------FKEKYYK----------------------------------A--SAII---------- 121
WP_012981280_1 DUF63 ------------------------------------------FKEKYYK----------------------------------V--SAVI---------- 121
WP_064496496_1 DUF63 ------------------------------------------FKEKYYK----------------------------------A--SAVI---------- 121
WP_011972741_1 DUF63 ------------------------------------------FKEKYYK----------------------------------Y---------------- 119
WP_013798289_1 DUF63 ------------------------------------------LRERYYK----------------------------------L---------------- 117
WP_013181100_1 DUF63 ------------------------------------------LKKKYYI----------------------------------G---------------- 119
WP_013867628_1 DUF63 ---------------------------------------------KYHI----------------------------------S---------------- 120
WP_018153391_1 DUF63 ------------------------------------------FKNRYYI----------------------------------S---------------- 119
WP_011170272_1 DUF63 ---------------------------------------------RYYL----------------------------------L---------------- 119
WP_012066151_1 DUF63 ------------------------------------------LKKKYYL----------------------------------L---------------- 119
ODS43013_1 hypotheti -----------------------------------------GI--DYTK----------------------------------T--LFCI---------- 130
OIQ06085_1 hypotheti ------------------------------------------FKKKYYK----------------------------------F--MLVA---------- 124
PIN67069_1 hypotheti ------------------------------------------FKKKYYK----------------------------------F--MLVA---------- 124
PIV28099_1 hypotheti ------------------------------------------FKKKYYK----------------------------------F--MLVA---------- 124
PJC13070_1 hypotheti ------------------------------------------FKKKYYK----------------------------------F--MLVA---------- 124
PIZ29927_1 hypotheti ------------------------------------------FKKKYYK----------------------------------F--MLVA---------- 124
PKP60688_1 hypotheti ------------------------------------------FKKKYYK----------------------------------F--MLVA---------- 124
WP_012956954_1 DUF63 ------------------------------------------KNIDYRY----------------------------------T--LSII---------- 120
WP_067148430_1 DUF63 -----------------------------------------GI--DYRY----------------------------------T--LFYI---------- 120
WP_080460538_1 DUF63 ------------------------------------------TNFDFKY----------------------------------T--LFFI---------- 119
WP_010877090_1 DUF63 -----------------------------------------GW--DYRK----------------------------------L--IFAT---------- 120
WP_013294901_1 DUF63 ------------------------------------------YGWDYRK----------------------------------L--VFAT---------- 120
WP_010877090_1 DUF63 ---------------GW----------------------------DYRK----------------------------------L--IFAT---------- 120
BAZ99473_1 hypotheti ---------------GW----------------------------DYRK----------------------------------L--IFAT---------- 120
PKL66404_1 hypotheti ------------------------------------------TQYDYRY----------------------------------I--IFTV---------- 128
WP_023991171_1 DUF63 ------------------------------------------TGWDYRY----------------------------------I--IFAV---------- 121
WP_048072898_1 DUF63 ------------------------------------------IGWDYRY----------------------------------V--IFAV---------- 121
WP_100905180_1 hypot ------------------------------------------TDRDYRY----------------------------------I--IFMV---------- 121
WP_100907231_1 hypot ------------------------------------------TDRDYRY----------------------------------I--IFMV---------- 121
WP_100907231_1 hypot ------------------------------------------TDRDYRY----------------------------------I--IFMV---------- 121
WP_048081925_1 DUF63 ------------------------------------------TKFDYRY----------------------------------F--ILIV---------- 120
WP_048081925_1 DUF63 ------------------------------------------TKFDYRY----------------------------------F--ILIV---------- 120
WP_069585525_1 DUF63 ------------------------------------------TKFDYRY----------------------------------F--ILIV---------- 120
WP_069585525_1 DUF63 ------------------------------------------TKFDYRY----------------------------------F--ILIV---------- 120
WP_013643671_1 DUF63 ------------------------------------------TQIDYRY----------------------------------I--IFGV---------- 120
WP_013824571_1 DUF63 ------------------------------------------TNFDYRY----------------------------------T--IFAV---------- 120
WP_048192110_1 DUF63 ------------------------------------------TNFDYRY----------------------------------I--IITV---------- 124
WP_081810122_1 DUF63 -----------------------------------------GRIKSYIH----------------------------------M--YAGA---------- 149
WP_097298965_1 DUF63 -----------------------------------------KKIKNYIH----------------------------------T--YAIT---------- 125
PIO00292_1 hypotheti --------------------------------------------RNPEK----------------------------------S--LFIM---------- 120
WP_011833679_1 DUF63 -----------------------------------------KKVVSYTT----------------------------------P--FMLG---------- 121
WP_042698332_1 DUF63 -----------------------------------------KIVVSYTT----------------------------------P--FMLG---------- 121
WP_042698332_1 DUF63 -----------------------------------------KIVVSYTT----------------------------------P--FMLG---------- 121
WP_011448164_1 DUF63 -----------------------------------------GMVQSWKK----------------------------------L--FQWT---------- 124
WP_007314214_1 DUF63 -----------------------------------------GIIRDYHR----------------------------------P--FAGA---------- 129
WP_042705632_1 DUF63 -----------------------------------------GYVDSYHK----------------------------------L--MCGI---------- 124
WP_013329717_1 DUF63 -----------------------------------------KIIADYHK----------------------------------G--FATG---------- 124
WP_004077722_1 DUF63 -----------------------------------------GIIRRYTQ----------------------------------G--FAAG---------- 124
WP_048150927_1 DUF63 -----------------------------------------KIIADYHK----------------------------------G--FATG---------- 124
WP_012107531_1 DUF63 -----------------------------------------GLIKDFLS----------------------------------F--YAFI---------- 130

```
PKL70081_1 hypotheti -----------------------------------------GLCKNYLK----------------------------------L--YGGA---------- 124
WP_015285007_1 DUF63 -----------------------------------------GLTKHYLT----------------------------------F--YFWA---------- 130
PKL64570_1 hypotheti -----------------------------------------GILTSFNA----------------------------------F--YAGA---------- 130
WP_015286404_1 DUF63 -----------------------------------------GIIASYHT----------------------------------G--YAGI---------- 124
WP_012617618_1 DUF63 -----------------------------------------GLLKDYHR----------------------------------G--FAGL---------- 124
WP_014867374_1 DUF63 -----------------------------------------GLVARYSR----------------------------------V--YGGV---------- 124
CVK33108_1 conserved -----------------------------------------GLVARYSR----------------------------------V--YGGV---------- 124
WP_066956331_1 DUF63 -----------------------------------------GLISRYSW----------------------------------F--YGGA---------- 124
WP_011844879_1 DUF63 -----------------------------------------GLVSRYSR----------------------------------L--YGGV---------- 124
WP_048181868_1 DUF63 -----------------------------------------GLVARYSR----------------------------------F--YGSV---------- 124
WP_067073014_1 DUF63 -----------------------------------------GLVPRYSR----------------------------------F--YGGA---------- 124
PKL62929_1 hypotheti -----------------------------------------GLTARYSR----------------------------------V--YGGV---------- 124
WP_004039890_1 DUF63 -----------------------------------------GVVREYSR----------------------------------T--YGLI---------- 124
WP_067049225_1 DUF63 -----------------------------------------GLVADSIR----------------------------------V--YGWI---------- 124
KYK37884_1 hypotheti ------------------------------------------WEENYYK----------------------------------Y--LAAV---------- 116
KYK28019_1 hypotheti ------------------------------------------WRENYYK----------------------------------Y--LAAI---------- 116
OYT57835_1 hypotheti ------------------------------------------KRVSYFK----------------------------------T--WFGI---------- 123
OYT33350_1 hypotheti ------------------------------------------YKREYYK----------------------------------P--YATF---------- 123
WP_012964745_1 DUF63 -----------------------------------------KNLKIYPY----------------------------------T---------------- 117
WP_048091573_1 DUF63 ---------------------------------------------KYRY-------------------------------------YPLP---------- 116
WP_048096399_1 DUF63 ------------------------------------------DYRRYPY----------------------------------P---------------- 116
WP_012940099_1 DUF63 ------------------------------------------RKENYWK----------------------------------H--HFAL---------- 119
WP_010877969_1 DUF63 ------------------------------------------YGEKYYR----------------------------------V--YAVV---------- 132
WP_013682839_1 DUF63 ------------------------------------------RGERYWI----------------------------------H--YGML---------- 120
WP_015591141_1 DUF63 -------------------------------------------KEDYWK----------------------------------Y--YAAV---------- 119
WP_012035887_1 DUF63 -----------------------------------------GVVKDYSK----------------------------------P--YFWT---------- 131
WP_014404626_1 DUF63 -----------------------------------------GMIGDYSK----------------------------------P--FFWA---------- 125
BAI60262_1 conserved -----------------------------------------GRVGDYSK----------------------------------P--LFWT---------- 125
WP_042684156_1 DUF63 ---------------------------------------WGGISRRTRV----------------------------------V--YGLC---------- 132
OFV68024_1 membrane  -------------------------------------------RPQFHT----------------------------------L--FGSF---------- 86
WP_013720253_1 DUF63 ------------------------------------------LGDQFHH----------------------------------G--YAAI---------- 122
WP_014587756_1 DUF63 ------------------------------------------LGDRWLL----------------------------------G--YAAV---------- 119
ABK14947_1 Protein o -------------------------------------------TGGYRL-------------------------------------YSAV---------- 124
OKY79140_1 putative  -----------------------------------------ETDLKTTK----------------------------------M--LFIT---------- 127
WP_086637003_1 DUF63 -----------------------------------------FIEGSQNK----------------------------------T--VFIV---------- 128
WP_048089097_1 DUF63 -----------------------------------------GKIKSYHA----------------------------------F--FGLM---------- 128
WP_097298272_1 DUF63 -----------------------------------------GKIRDYHT------------------------------------FFGYT--------- 129
WP_013897788_1 DUF63 -----------------------------------------GWIRDYRT----------------------------------P--FVLA---------- 130
WP_015323982_1 DUF63 -----------------------------------------WTGNSTET----------------------------------I--FASL---------- 130
WP_011499115_1 DUF63 -----------------------------------------EKVSDWKV----------------------------------P--FALL---------- 130
WP_013037846_1 DUF63 -----------------------------------------GVAGDWKK----------------------------------T--FAAA---------- 130
WP_048205640_1 DUF63 -----------------------------------------GKVTDWHR----------------------------------S--FALL---------- 130
WP_072561629_1 DUF63 -----------------------------------------DVVGDWKK----------------------------------T--FAAA---------- 130
WP_072360082_1 DUF63 -----------------------------------------DVVGDWKK----------------------------------T--FAAA---------- 130
WP_096711793_1 DUF63 -----------------------------------------DVVGDWEK----------------------------------T--FAAA---------- 130
ODV50598_1 hypotheti -----------------------------------------DVVGDWEK----------------------------------T--FAAA---------- 130
WP_013194042_1 DUF63 -----------------------------------------EFIKDYHI----------------------------------L--VASI---------- 130
WP_048178067_1 DUF63 -----------------------------------------GFVRDFYS----------------------------------T--FAGF---------- 132
WP_048127111_1 DUF63 -----------------------------------------GLVKDYHP----------------------------------I--FASF---------- 132
WP_011023594_1 DUF63 -----------------------------------------GFVEDYHP----------------------------------V--FAGF---------- 136
WP_048184587_1 DUF63 -----------------------------------------GFVKDYHP----------------------------------V--FASF---------- 132
WP_011032542_1 DUF63 -----------------------------------------GLVKDYHP----------------------------------A--FAGF---------- 132
WP_011032542_1 DUF63 -----------------------------------------GLVKDYHP----------------------------------A--FAGF---------- 132
WP_048129141_1 DUF63 -----------------------------------------GHVKDFHL----------------------------------V--FAGF---------- 132
WP_048129141_1 DUF63 -----------------------------------------GHVKDFHL----------------------------------V--FAGF---------- 132
WP_048169891_1 DUF63 -----------------------------------------GLVKEYHS----------------------------------V--FAGF---------- 132
WP_048137175_1 DUF63 -----------------------------------------GLVKEYHS----------------------------------I--FAGF---------- 132
WP_048137175_1 DUF63 -----------------------------------------GLVKEYHS----------------------------------I--FAGF---------- 132
WP_048137175_1 DUF63 -----------------------------------------GLVKEYHS----------------------------------I--FAGF---------- 132
WP_048137694_1 DUF63 -----------------------------------------GLVKDYHP----------------------------------V--FAGF---------- 132
WP_048168126_1 DUF63 -----------------------------------------GLIKDFHL----------------------------------I--FASF---------- 132
WP_048118082_1 DUF63 -----------------------------------------GLVKDFHL----------------------------------T--FAGF---------- 132
WP_048158120_1 DUF63 -----------------------------------------GLVKDFHL----------------------------------T--FAGF---------- 132
WP_011305329_1 DUF63 -----------------------------------------GLVKDFHL----------------------------------T--FAGF---------- 132
WP_054298619_1 DUF63 -----------------------------------------GMIKDFHL----------------------------------V--FASF---------- 132
ALK05385_1 hypotheti -----------------------------------------GLIKDFHL----------------------------------I--FASF---------- 132
WP_015052897_1 DUF63 ------------------------------------------RDKDFET----------------------------------V--FALF---------- 129
WP_023846134_1 DUF63 -----------------------------------------KGTGDYRK----------------------------------L--FAGF---------- 129
OIN88451_1 hypotheti ------------------------------------------YKIPYYK----------------------------------I--MFIS---------- 120
PIX50278_1 hypotheti -----------------------------------------KYKIPYYK----------------------------------I--MFIS---------- 120
PIW41402_1 hypotheti -----------------------------------------KYKIPYYK----------------------------------I--MFIS---------- 120
PIY35178_1 hypotheti -----------------------------------------KYKIPYYK----------------------------------I--MFIS---------- 120
PJB74886_1 hypotheti -----------------------------------------KYKIPYYK----------------------------------I--MFIS---------- 120
PIZ33651_1 hypotheti -----------------------------------------KYKIPYYK----------------------------------I--MFIS---------- 120
WP_048165029_1 DUF63 -------------------------------------------SEDWRR----------------------------------T--FLYF---------- 118
WP_042681046_1 DUF63 -------------------------------------------FEDWRK----------------------------------V--FLYF---------- 118
WP_013467319_1 DUF63 -------------------------------------------FEDWRR----------------------------------V--FLYF---------- 118
WP_048152160_1 DUF63 -------------------------------------------FEDWRR----------------------------------A--FLYT---------- 118
WP_015849008_1 DUF63 -------------------------------------------FEDWRK----------------------------------V--FLYF---------- 118
WP_004069276_1 DUF63 -------------------------------------------FEDWRK----------------------------------V--FLYF---------- 118
WP_058946638_1 DUF63 -------------------------------------------FEDWRK----------------------------------V--FLYF---------- 118
WP_042701551_1 DUF63 -------------------------------------------FEDWRK----------------------------------V--FLYF---------- 118
WP_055282692_1 DUF63 -------------------------------------------FEDWQK----------------------------------V--FLYF---------- 118
WP_013906023_1 DUF63 -------------------------------------------FEDWRR----------------------------------V--FLWF---------- 117
WP_011013158_1 DUF63 -------------------------------------------AKDWRG----------------------------------V--FLWF---------- 117
WP_014733166_1 DUF63 -------------------------------------------FKDWRG----------------------------------V--FLWF---------- 117
WP_068322889_1 DUF63 -------------------------------------------FKDWRG----------------------------------V--FLWF---------- 117
WP_068575625_1 DUF63 -------------------------------------------FKDWRG----------------------------------V--FLWF---------- 117
WP_010885960_1 DUF63 -------------------------------------------FKDWRR----------------------------------V--FSLF---------- 117
WP_010867401_1 DUF63 -------------------------------------------FKDWRG----------------------------------V--FLSF---------- 117
WP_013747987_1 DUF63 -------------------------------------------FKDWRG----------------------------------V--FLSF---------- 118
WP_068664049_1 DUF63 -----------------------------------------CRGCDWQR----------------------------------A--FFGF---------- 120
WP_011249739_1 DUF63 ----------------------------------------LGPDERLYP----------------------------------I--YRDF---------- 121
WP_062386854_1 DUF63 ----------------------------------------LGPDERLYP----------------------------------I--YRDF---------- 121
WP_050003781_1 DUF63 ----------------------------------------LGPDEKLYP----------------------------------I--YRDF---------- 121
```

```
WP_042690699_1 DUF63 ----------------------------------------LGPDDRLYP----------------------------------I--YRDF---------- 121
WP_088885212_1 DUF63 ----------------------------------------VGPGEKLYP----------------------------------L--YRDV---------- 121
WP_010478519_1 DUF63 ----------------------------------------VGPGEKFYP----------------------------------L--YRDV---------- 121
WP_088858240_1 DUF63 ----------------------------------------VGPGEKFYP----------------------------------L--YRDV---------- 121
WP_015859016_1 DUF63 ----------------------------------------VGTGEKLYP----------------------------------L--YRDV---------- 121
WP_014121940_1 DUF63 ----------------------------------------VGAGEKLYP----------------------------------L--YRDV---------- 121
WP_062373911_1 DUF63 ----------------------------------------VGPGEKLYP----------------------------------L--YRDV---------- 121
WP_088882924_1 DUF63 ---------------------------------------HLGPEERLYP----------------------------------I--YRDF---------- 121
WP_088862911_1 DUF63 ---------------------------------------HLGPDERLYP----------------------------------I--YRDF---------- 121
WP_012572006_1 DUF63 ----------------------------------------VGSGEKFYP----------------------------------L--YRDF---------- 121
WP_088854707_1 DUF63 ----------------------------------------IGPGEKLYP----------------------------------L--YRDF---------- 121
WP_014789474_1 DUF63 -----------------------------------------CPGERLYP----------------------------------L--YRDF---------- 120
WP_088180728_1 DUF63 ----------------------------------------VGPGEKLYS----------------------------------L--YRDF---------- 121
WP_088864937_1 DUF63 -----------------------------------------CPGERLYP----------------------------------L--YRDF---------- 120
WP_055429686_1 DUF63 ----------------------------------------VGPEERFYP----------------------------------L--YRDF---------- 121
WP_088865932_1 DUF63 ----------------------------------------VGPGEKLYP----------------------------------L--YRDF---------- 121
WP_014013642_1 DUF63 ----------------------------------------VGPGERLYP----------------------------------L--YRDF---------- 121
WP_088856804_1 DUF63 ----------------------------------------VGPGERLYP----------------------------------L--YRDF---------- 121
WP_058939475_1 DUF63 -----------------------------------------CPGEKLYP----------------------------------L--YQDF---------- 120
EHR77276_1 conserved IALMFHWGLLYQPAYIVHDEMGQVWAMGGLLAAFGMLLWSVIKTRHWPA---------------------------------ITRGLFSF---------- 246
OUV40025_1 hypotheti -------------------------------------------GKKWDD----------------------------------V--AGDL---------- 176
MBJ52984_1 hypotheti ------------------------------------------SWSPLER----------------------------------M--LFST---------- 241
PDH23744_1 hypotheti -------EREGHSQKNDLETMATGLILIQFVVYASSISGKEGDLSLWPM----------------------------------L--LGVIAAIAFALWSR 217
PDH25468_1 hypotheti -----------------------------------------DFFDDLQR----------------------------------T--VYFT---------- 237

WP_048201856_1 hypot --------------------------------------NLL----------------------------------------------------------- 99
WP_048150344_1 hypot ------------------------GYYVVILLTL----ALN--------------------------------------------------VHSQIAIVL 134
AOV95360_1 hypotheti ------------------------TAITALWKIT----GLK------------------PDIPLFIQI----------------TG-LAVAVFLPFYLIH 136
EOD42420_1 Uncharact ------------------------LLVILILEIP----FFI----F-------------GRPINLLPS----------------LI---F-ILLFSILIY 150
AOV95164_1 hypotheti ------------------------GYIYLAPFLA----LTV----Y----------TLLGAANLLLLL----------------LP-IGL-ASVLTGIYY 177
EGQ40074_1 putative  ------------------------GLGLLSVAVA----VAA-----------TQVEIAAPAFGAKGLL----------------VP------AALTGVFY 175
KYK23067_1 hypotheti QTMKINTIIFS-------------GGLIVALPCF----FLI--GRWMVGDQWSISTGTYANVFWLIIG----------------LI-SII-IGIVFLIAW 284
WP_084383883_1 DUF63 --------------------------------------YMR--------------------------------------------------NAAIGFGII 187
WP_049984677_1 DUF63 ------------------------GAIWAVGAVG----WAF---LY-----GATTAAPLRLWVPVATT----------------GI-----ALGVTALYY 222
WP_103428047_1 hypot ------------------------GTVWAIGAVG--WAFLY----------GATSPAPLRLWVPVATT----------------GI-----ALGVTGLYY 221
WP_004048647_1 DUF63 ------------------------GSVWAVGAVG----WVL----W----YGLSTSTPLRLWVPVATT----------------GI-----ALGATALYY 221
WP_004594600_1 DUF63 ------------------------GSVWAVGAVW----WAV----W----HGLSTSTPLRLWVPVATT----------------GI-----ALGVTALYY 221
WP_004594600_1 DUF63 ------------------------GSVWAVGAVW----WAV----W----HGLSTSTPLRLWVPVATT----------------GI-----ALGVTALYY 221
WP_050050397_1 DUF63 ------------------------GSVWAVGAVW----WAV----W----HGLSTSTPLRLWVPVATT----------------GI-----ALGVTALYY 221
WP_004594600_1 DUF63 ------------------------GSVWAVGAVW----WAV----W----HGLSTSTPLRLWVPVATT----------------GI-----ALGVTALYY 221
WP_050050397_1 DUF63 ------------------------GSVWAVGAVW----WAV----W----HGLSTSTPLRLWVPVATT----------------GI-----ALGVTALYY 221
WP_049983690_1 DUF63 ------------------------GTVWAIGAVG----WAL----W----YGLSTSAPLRVWVPVATT----------------GI-----ALGVTALYY 221
WP_103428162_1 hypot ------------------------GSVWAVGAVG----WAL----W----YGLSTTAPLRLWVPVATT----------------GI-----ALGVTALYY 221
WP_009378268_1 DUF63 ------------------------GSVWAVGAVG----WAF--------WYGLSTPTTLRLWVPVATT----------------GI-----ALGVTALYY 221
WP_015763176_1 DUF63 ------------------------GWGLLSLNLL----SLL----W-----LSVT-REYVTLYPQMVT----------------IV-VVG-AIALAYVNW 255
WP_018259238_1 DUF63 ------------------------GWGLLTLNLL----YLL----W-----LSVT-REYVTLYPQMVT----------------IV-VVG-AIALAYVNW 255
WP_004592615_1 DUF63 ------------------------GGAFVAITLL----YLT----F-----VALT-REYATLYPSVLI----------------TT-VAL-ASVLSYGLY 249
WP_004518092_1 DUF63 ------------------------GGAFVAVTLL----YLT----F-----VALT-REYATLYPSVLI----------------TT-VVL-ASALSYGLY 249
WP_014040450_1 DUF63 ------------------------GSIFVAVTLL----YLT----F-----VALT-REYATLYPSVLI----------------TT-VAL-ASALSYGLY 249
WP_014040450_1 DUF63 ------------------------GSIFVAVTLL----YLT----F-----VALT-REYATLYPSVLI----------------TT-VAL-ASALSYGLY 249
WP_008309018_1 DUF63 ------------------------GSIFVAVTLL----YLT----V-----VALT-REYATLYPSVLI----------------TT-VAL-ASALSYGLY 249
WP_014040450_1 DUF63 ------------------------GSIFVAVTLL----YLT----F-----VALT-REYATLYPSVLI----------------TT-VAL-ASALSYGLY 249
WP_014040450_1 DUF63 ------------------------GSIFVAVTLL----YLT----F-----VALT-REYATLYPSVLI----------------TT-VAL-ASALSYGLY 249
WP_005534235_1 DUF63 ------------------------GGAFVAVTLL----YLT----F-----VALT-REYATLYPSVLI----------------TT-VAL-ASALSYGLY 249
WP_004961153_1 DUF63 ------------------------GGAFVAVTLL----YLT----F-----VALT-REYATLYPSVLI----------------TT-VAL-ASALAYGLY 249
WP_004961153_1 DUF63 ------------------------GGAFVAVTLL----YLT----F-----VALT-REYATLYPSVLI----------------TT-VAL-ASALAYGLY 249
WP_101350151_1 hypot ------------------------GGAFVAVTLL----YLT----V-----VALT-REYATLYPSVLI----------------TT-VVL-ASALSYGLY 249
WP_053968273_1 DUF63 ------------------------GGAFVAVTLL----YLT----F-----VALT-REYATLYPSVLI----------------TT-VML-ASALSYGLY 249
WP_058995687_1 DUF63 ------------------------GGAFVAVTIL----YLT----F-----VALT-REYATLYPSVLI----------------TT-VAL-ASALSYGLY 249
WP_015790673_1 DUF63 ------------------------GAVAFGVTFG----YLT----------VLGLTVEHVDFLPHVLL----------------II-VGG-ATVAAYGVY 250
WP_008524115_1 DUF63 ------------------------GGVALAITLG----YLV----------VLGLTVEHVAFLPHVLL----------------II-VGL-ATVLAYGVY 250
WP_075936143_1 DUF63 ------------------------GWVLVLCSVG----YLL----YFVPTRLTGT-VDGAGFYPQVTG----------------VT-VGL-SVLFAVSIY 248
WP_020446311_1 DUF63 ------------------------GAVLFLLALG----SLV----Y-----LSVT-TDAVGFNLGVLV----------------VT-VLG-ATLATALAW 245
WP_049898450_1 DUF63 ------------------------GTLALVATLG----YLT----F-----LSIS-TEALGFYPQMLV----------------LT-LGI-ASAIAGGVY 247
WP_006077061_1 DUF63 ------------------------GTLALVAVLG----YLT----Y-----LATA-TEFVGFYPQMLV----------------LT-LGL-AALIAGGVY 247
WP_049996700_1 DUF63 ------------------------GTLALAATLA----YLA----F-----LSYS-TEFVAFYPQMLV----------------IT-LAI-ASGLAAGIY 247
EMA38628_1 hypotheti ------------------------GSLALAVTVG----YLG----Y-----LAAT-TEYLDFYPQMLV----------------AT-LGI-ASAIAVGVY 247
WP_049992561_1 DUF63 ------------------------GAVLFVLTFA----SLI----I-----VSVL-RDSVGFYPSFLL----------------VT-VGL-ASVIAYGIY 248
WP_010903025_1 DUF63 ------------------------GCVLLAATLL----GLG----V-----LAAS-HDYITFIPVFTV----------------LT-LGG-ATVITAVTW 246
WP_009760947_1 DUF63 ------------------------GTVLLVANIL----GLG----Y-----LAAT-RDYVSFIPVFTV----------------LT-LGL-ATVITAPVW 245
WP_059057097_1 DUF63 ------------------------GTALLVATVG----FLV----Y-----LSAT-TEYVSLLPQFTV----------------LT-LLF-ATVITAAVW 245
WP_058983522_1 DUF63 ------------------------GTALLTATVG----FLV----Y-----LSAT-TEYVSLHPQFTL----------------LT-VLF-ATIITAVVW 245
WP_071932813_1 DUF63 ------------------------GSGMLVLALG----YLL----Y-----LSAT-QPQVEFHPAFTI----------------LT-MGG-ATLIAVLGW 241
WP_050048584_1 DUF63 ------------------------GTAMLAATLG----YLV----W-----LSVT-TDFVAFHPIFTI----------------LT-MGL-ALVIAVGLW 241
WP_014051341_1 DUF63 ------------------------GTGVFALSLG----YLV----V-----LAVLPDNPVVFYPQVLA----------------IV-VLL-AVASTALTW 245
WP_079233299_1 DUF63 ------------------------GAVAFLLSIV----YLS----Y-----LAA-TESAVTFYPLVLI----------------AM-VGL-SAAAAAATW 243
WP_053948572_1 DUF63 ------------------------GSGAFLLSVG----YLS----Y-----LAL-TDPQVQFYPLVLV----------------AM-VGL-SAAAAVGTW 243
WP_049980835_1 DUF63 ------------------------GSVAFLLSIA----YLS----M-----LAV-TESRVTFYPLVLV----------------AM-VGL-SAAAAAGTW 243
KPN31749_1 hypotheti ------------------------GSLAFLASVA----YLT----M-----LAA-TEPQVTFYPLVLL----------------AM-VGL-SAAAAAGTW 193
AGB16040_1 putative  ------------------------GTGLLVLSLA----ILA----S-----WAST-GGYG-FFPGLAV----------------TV-LVV-STVITGAVW 241
WP_007696755_1 DUF63 ------------------------GTSLLAVTLV----ILW----Q-----WAAI-GGVG-FFPLITV----------------TT-LVV-ASTLTALVW 241
WP_015322904_1 DUF63 ------------------------GAVALAATLG----FLG----Y-----TAAT-ETYSTFYWTIPA----------------IV-LTA-ATVSTAVVW 242
WP_076581869_1 DUF63 ------------------------GTVVLAVTLG----WLG----Y-----LAAV-TDYVGFVPSIAA----------------IV-LVA-ATVSTAIVW 242
WP_005559622_1 DUF63 ------------------------GAVALAATLG----FLG----Y-----TAAT-QPYSTFYPSIPA----------------IV-LTA-ATVSTALVW 242
WP_006067640_1 DUF63 ------------------------GAVVLAATLG----WLG----Y-----LAVT-TEYASFYPSIPT----------------IV-LTA-ATVSTAIVW 237
WP_008164554_1 DUF63 ------------------------GAAVLAVTLG----WLG----Y-----LAAT-TAYVSFYPSIPA----------------IV-LTA-ATVSTALVW 242
WP_006088112_1 DUF63 ------------------------GAVVLAVTVG----WLG----Y-----LAAT-TEYASFYPSIPA----------------IV-LTA-ATVSTALVW 242
WP_012943247_1 DUF63 ------------------------GTAYLTLTLA----YLG----Y------AST-QPYADFHPLLLL----------------TT-LVG-ATLATAITW 241
WP_008895026_1 DUF63 ------------------------GTTYLTVTLA----YLV----Y------AST-QPYADFHPLLLV----------------TT-LVG-ATLATAVTW 241
WP_098727043_1 DUF63 ------------------------GTGLLTLTVL----FLA----Y-----TAAT-KGYAEFYPLVPL----------------VI-LVG-ATLTTAITW 242
WP_049990861_1 DUF63 ------------------------GTAALTLTIG----YLA----Y-----TAAT-KPYAEFYPLIPL----------------VV-LVG-ATLTTAITW 240
WP_008013001_1 DUF63 ------------------------GTALLTVAIA----YLG----Y-----TAAT-EPYSEFYPLIPL----------------VT-LVG-ATLATAITW 242
WP_006180566_1 DUF63 ------------------------GATVLTVTVA----YLA----Y-----TAAT-EPYAEFYPLIPL----------------VV-LVG-ATVATAITW 242
WP_006650865_1 DUF63 ------------------------GTAALTVTIA----YLA----Y-----TAAT-EPFSEFYPLIPL----------------VV-LVG-ATVATAITW 242
WP_066301295_1 DUF63 ------------------------GTAVLTVTIA----YLA----Y-----TAAT-EPYSEFYPLIPV----------------VI-LIG-ATVATAVTW 242
```

```
WP_049966804_1 DUF63 ------------------------GTAVLTITIA----YLA----Y-----TAAT-EPFSEFYPLIPL----------------VV-LVG-ATVATAITW 242
WP_076145380_1 DUF63 ------------------------GTAVLTVTIA----HLA----Y-----TAAT-EPYSEFYPLIPL----------------VV-LVG-ATVATAITW 242
WP_097378845_1 DUF63 ------------------------GTLAVTITIA----YLA----A-----TAAT-EPYSEFHPLIPL----------------VI-LVG-ATLTTAITW 242
WP_008452261_1 DUF63 ------------------------GTAALTITIA----YLG----F-----VAAT-EPYAAFHPLIPL----------------VV-LVG-ATLTTAITW 242
WP_008452261_1 DUF63 ------------------------GTAALTITIA----YLG----F-----VAAT-EPYAAFHPLIPL----------------VV-LVG-ATLTTAITW 242
WP_006432643_1 DUF63 ------------------------GTTVLTITIA----YLA----Y-----VAAT-EPYAEFYPLIPL----------------VI-LVG-ATLTTAITW 242
WP_008452261_1 DUF63 ------------------------GTAALTITIA----YLG----F-----VAAT-EPYAAFHPLIPL----------------VV-LVG-ATLTTAITW 242
WP_007109662_1 DUF63 ------------------------GTAALTVTIA----YLA----F-----VAAT-EPYAEFHPLIPL----------------VV-LVG-ATLTTALTW 242
WP_049952821_1 DUF63 ------------------------GTLALASALG----YLG----Y-----LAAT-KPYVEFYPGLLL----------------VT-LGG-ATIATAITW 242
WP_086889541_1 DUF63 ------------------------GTGLLAVTVG----YLG----Y-----VAAV-HEYATFYPWLLL----------------TT-LVG-ATIATAITW 245
WP_005580009_1 DUF63 ------------------------GTAYLVITVA----FLG----Y-----TA-V-QEYATFYPWLLL----------------TV-LVG-ATLATWVTW 244
WP_049927356_1 DUF63 ------------------------GATLLATTLA----YLS----Y-----LAAV-KPYAEFYGLLLL----------------TT-LGG-ATLATAIVW 242
WP_087714455_1 DUF63 ------------------------GTALLAVTVG----YLG----Y-----TAAV-HEYATFYPYLLV----------------TT-LVG-ATLSTAIVW 242
WP_013878438_1 DUF63 ------------------------GTGLLALTLA----HLG----Y-----TAAT-EPFSRFYPLLLV----------------TT-LVG-ATAATALTW 242
WP_049921111_1 DUF63 ------------------------GTGLLAVTVG----YLG----F-----TAAT-EPYGTFYPWLLV----------------TT-LVG-ATAATAVTW 242
WP_007142789_1 DUF63 ------------------------GTALLTLTVG----YLG----V-----TAAT-EEYATFYPYLLL----------------TT-LAV-ATLATWLTW 241
WP_006651941_1 DUF63 ------------------------GTVLLTGTVG----YLG----Y-----LAWA-EPYATFYPWLLI----------------VT-LVG-ATVATWLTW 242
WP_004216437_1 DUF63 ------------------------GTVLLTGTVG----YLG----Y-----LAWA-EPYATFYPWLLI----------------VT-LVG-ATVATWLTW 242
WP_071402513_1 DUF63 ------------------------GTVLLTGTIG----YLS----Y-----LAWA-EPYATFYPWLLI----------------VT-LVG-ATVATWLTW 242
WP_006666462_1 DUF63 ------------------------GTTLLVGTVG----YLG----Y-----LSAT-TDYVQFYPLLLL----------------TI-VGG-ATVATAITW 242
WP_049904535_1 DUF63 ------------------------GTTLLVGTVG----YLG----Y-----LSAT-TDYVQFYPLLLL----------------TI-LGG-ATVATAITW 242
WP_006824405_1 DUF63 ------------------------GTTLLAGTVG----YLG----Y-----LSAT-TDYVEFYPLLLL----------------TI-LGG-ATVATAITW 242
WP_011323802_1 DUF63 ------------------------GTAAVVAVVG----WLL----Y-----VAGT-TDIVEFYLLVPT----------------IV-LGG-ATLTTLVFW 243
WP_015409719_1 DUF63 ------------------------GALAVVATVG----GLL----Y-----VSAT-SELVGFFPAVAA----------------IT-LGG-ATLVAALFW 244
WP_006883947_1 DUF63 ------------------------GLGVLALTLG----YLF----F-----VAFT-REYVSVYPQILL----------------VD-VGL-ASVLAYLLY 243
WP_077207545_1 DUF63 ------------------------GSAAFLLTFG----YLT----S-----LAFT-ADYVNFYPQVLL----------------SV-VAI-ATLLAVGIY 246
ESS12903_1 putative  ------------------------GVLGVVASVG----TLG----W-----LAVT-TEYLGFHPVVPV----------------VT-LGT-ATIVAAVAW 246
WP_008417361_1 DUF63 ------------------------GTLVLAVTVG----YLG----W-----LAVT-TDVLVFNPAVAI----------------IT-FGG-ASVVAALAW 245
WP_066381608_1 DUF63 ------------------------GTLAFTVTIG----YLG----W-----LAAS-TSVLEFHPAVAA----------------IT-LVG-ATLVAALAW 240
WP_049947642_1 DUF63 ------------------------GVALNVVTVG----YLF----A-----LSLTPGTPVDFYPQVLV----------------AT-LAL-TAAATAATW 247
ESS05953_1 putative  ------------------------GTGVLLATLA----LLA----A-----VAAD----DAFYPQVFA----------------VV-FVI-ATLAVGAVL 237
WP_096390051_1 DUF63 ------------------------GAVAVVAAVG----YLA----F-----LAAT-TPFVGFNPAVIG----------------SI-LVL-ATATTGVTW 252
WP_021072749_1 DUF63 ------------------------GTAAVVASVG----YLG----Y-----LAAT-TPFVGFNPAVIG----------------SI-LTI-ATATTGVTW 252
WP_049982791_1 DUF63 ------------------------GVGALALALG----YLS----Y-----LAAA-TDYVDFYPIILV----------------ST-LGI-ATLATLVTW 250
WP_008585990_1 DUF63 ------------------------GAAGLALAVG----YLA----V-----LAAT-TGYVTFYPQVLV----------------PT-LVI-ATAATAGTW 252
WP_006628721_1 DUF63 ------------------------GAAGLALAVG----YLS----Y-----LAAT-TGYVTFYPQVLV----------------PT-LAI-ATAATAGTW 250
WP_006113218_1 DUF63 ------------------------GAAGLALAVG----YLA----Y-----LAVA-TDYVTFYPQVLV----------------PT-LVI-ATVTAAITW 251
WP_049930034_1 DUF63 ------------------------GAAGLALAVG----YLA----Y-----LAAT-TGYVTFYPQVLV----------------PT-LAV-ATAATAGTW 252
WP_049906047_1 DUF63 ------------------------GAAGLALAVG----YLS----Y-----LAAT-TGYVTFYPQVLV----------------PT-LAI-ATLATAVTW 252
WP_044965494_1 DUF63 ------------------------GAAGLALAVG----YLS----Y-----LAAT-TGYVTFYPQVLV----------------PT-LVI-ATAATAVTW 252
WP_096393195_1 DUF63 ------------------------GTAGLALAVG----YLS----Y-----LAAT-TDYVTFYPQVLV----------------PT-LVI-ATAATAVTW 252
WP_049908400_1 DUF63 ------------------------GAAGLALAVG----YLS----Y-----LAAT-TGYVTFYPQVLV----------------PT-LVI-ATVATAVTW 252
WP_049908585_1 DUF63 ------------------------GAAGLALAVG----YLS----Y-----LAAT-TGYVTFYPQVLV----------------PT-LVI-ATVATAGTW 252
WP_049902631_1 DUF63 ------------------------GAAGLALAVG----YLS----H-----LAAT-TGYVTFYPQVLV----------------PT-LVI-ATVATAVTW 252
WP_049983668_1 DUF63 ------------------------GAAGLALAVG----YLS----Y-----LAAT-TGYVTFYPQVLV----------------PT-LVI-ATAATAGTW 251
WP_053772267_1 DUF63 ------------------------GTAGLALAVG----YLV----Y-----LAAT-TGYVTFYPQVLV----------------PT-LVI-ATVATAATW 251
WP_007999151_1 DUF63 ------------------------GAVGLAAAVG----YLS----V-----LAAT-TEYVDVFYPIVLV----------------LT-LGI-ATAVTAITW 250
WP_015909917_1 DUF63 ------------------------GAVALTVTFG----YLS----Y-----LSAT-TDYVEFYPIILV----------------LT-LGI-ATAATAVTW 251
WP_004050594_1 DUF63 ------------------------GAVGLAVAFG----YLS----Y-----LSAA-TDYVQFYPIILV----------------LT-LGI-ATVATAVTW 251
WP_095636035_1 DUF63 ------------------------GAVALAVAFG----YLS----Y-----LSAA-TDYVQFYPIVLA----------------LT-LGI-ATAATAVTW 251
WP_008003811_1 DUF63 ------------------------GAVALALAFG----YLS----Y-----LSAA-TDYVQFYPIVLA----------------LT-LGI-ATAATAVTW 251
WP_066416034_1 DUF63 ------------------------GSLALALSVG----YLA----Y-----LAAA-TDYVTFYPQVIG----------------SI-LLI-ASAAAAVTW 252
ESS03170_1 putative  ------------------------GATVLVLTVG----YLS----Y-----LAAA-TDYVTVYPQVIG----------------TV-LVL-ATVSTAATW 251
WP_089671383_1 DUF63 ------------------------GTTLLVGSLG----YLT----------YLGATTEYATIHPQVLL----------------VT-LVG-ATLSAWLTW 250
ERH07656_1 putative  ------------------------GTLLLVTSVG----YLT----------VLAVTTPYVRLRPQVLA----------------VV-VIG-ATLSAAAAW 250
ESS10096_1 putative  ------------------------GTLLLVTSVG----YLT----------VLAVTTPYVRLRPQVLA----------------VV-VIG-ATLSAAVAW 250
ESS07824_1 putative  ------------------------GTMFCSRSQP----SAS--------------------------------------------------GSASPTAAS 195
ERH05689_1 putative  ------------------------GLTLLTLSVG----YLT----------YIGLTTSYATVNLQVLA----------------AV-LGG-ATITAGLTW 250
ERH02266_1 putative  ------------------------GLTLLTLSVG----YLT----------YIGLTTSYATVNLQVLA----------------AV-LGG-ATITAGLTW 250
ESS07825_1 putative  -------------------------------------------------------------------------------------------ATITAGLTW 43
WP_049970017_1 DUF63 ------------------------GTVVLAATIG----YLF----Y-----LTVT-TEVVGFYPQILT----------------VV-LVG-ATVVTALTW 244
WP_007979289_1 DUF63 ------------------------GTLVLAATMG----YLG----V-----LAFT-TRRVGFYPQVLV----------------VV-LVG-ATLVTAVTW 244
WP_007977745_1 DUF63 ------------------------GTVVLAATVG----YLV----M-----LAMT-TQTVRFYPQMLV----------------VV-LGG-ATLVTAVVW 244
WP_014556445_1 DUF63 ------------------------GTGVLALALG----YLI----S-----LVINSAPGVEFYPQVLL----------------TV-VIG-ATAVTGITW 246
WP_049935775_1 DUF63 ------------------------GVVVLGTTLL----YLL----W-----SGVAPDGPGTFYPQVIV----------------VI-LLG-STLAAGGTW 244
WP_008325149_1 DUF63 ------------------------GVAVLAVSLA----YLL----W-----AGLTGSQGATFYPQVLV----------------VI-LLG-STVAAAAAW 246
WP_007543631_1 DUF63 ------------------------GVAVLAVSLA----YLL----W-----AGLTGSQGATFYPQVLV----------------VM-LLG-STVAAAATW 246
WP_049905041_1 DUF63 ------------------------GLAVLAVAVG----YLL----W-----AGLTGAQGATFYPQVLV----------------VI-LVG-ATLVTAATW 246
WP_049913563_1 DUF63 ------------------------GLAVLAVAVG----YLL----W-----AGLTGAQGATFYPQVLA----------------VI-LVG-ATLVTAATW 246
WP_049967851_1 DUF63 ------------------------GLAVLAVAVG----YLL----W-----AGLTGAQGATFYPQVLA----------------VI-LVG-ATLVTAATW 246
WP_049914905_1 DUF63 ------------------------GLAVLAVTLG----YLL----W-----AGLTGAQGATFYPQVLV----------------VI-LVG-ATLTAAATW 246
WP_049916430_1 DUF63 ------------------------GLAVLAVTLG----YLL----W-----AGLTGAQGATFYPQVLV----------------VI-LVG-ATLTAAATW 246
WP_049896892_1 DUF63 ------------------------GLAVLAVTLG----YLL----W-----AGLTGAQGATFYPQVLA----------------VI-LVG-ATLTAAATW 246
WP_049896892_1 DUF63 ------------------------GLAVLAVTLG----YLL----W-----AGLTGAQGATFYPQVLA----------------VI-LVG-ATLTAAATW 246
WP_049896892_1 DUF63 ------------------------GLAVLAVTLG----YLL----W-----AGLTGAQGATFYPQVLA----------------VI-LVG-ATLTAAATW 246
WP_058828253_1 DUF63 ------------------------GLAVLAVAIG----YLL----W-----AGLTGAQGATFYPQVLV----------------VI-LVG-ATLAAAATW 246
WP_058568959_1 DUF63 ------------------------GLAVLAVTLG----YLL----W-----AGLTGAQGATFYPQVLA----------------VI-LVG-ATLTAAATW 246
WP_049917947_1 DUF63 ------------------------GLAVLAVTLG----YLL----W-----AGLTGAQGATFYPQVLV----------------VI-LVG-ATLTAAATW 246
WP_049920348_1 DUF63 ------------------------GLAVLAVTLG----YLL----W-----AGLTGAQGATFYPQVLV----------------VI-LVG-ATLTAAATW 246
WP_089777545_1 DUF63 ------------------------GLAVLAVTLG----YLL----W-----AGLTGAQGATFYPQVLV----------------VI-LVG-ATLAAAATW 246
WP_008320633_1 DUF63 ------------------------GSAVLAVTLG----YLL----W-----AGLTGAQGATFYPQVLL----------------VI-LAG-ATVAAAATW 246
WP_004060355_1 DUF63 ------------------------GTAVLATTLG----YLL----W-----AGITGAQGATFYPQVLL----------------VI-LAG-ATVVATATW 246
WP_103426078_1 hypot ------------------------GVVVLGLTLA----YLA----S-----LAVTGSNSVTFYPQVIL----------------TM-LVG-ATLAALATW 246
WP_009367433_1 DUF63 ------------------------GAAALAVTVG----YLG----V-----LAATGSNGVTFYPQVIT----------------VI-LVG-ATLSAAVTW 245
WP_013440552_1 DUF63 ------------------------GVVVLGLTLA----YLF----S-----LAINGAQGVEFHPQVLV----------------VM-LAG-ATLSAAATW 246
WP_049916626_1 DUF63 ------------------------GVAVLALTLG----YLF----W-----LAASGAEGVEFYPQVLV----------------VM-LLG-ATLAAGATW 246
WP_058582837_1 DUF63 ------------------------GSVVLAVTLA----YLG----Y-----LAAT-TEYVTFYPQVIT----------------VM-LVL-TCVATAITW 247
WP_101298124_1 hypot ------------------------GAAILAVTVG----YLV----A-----LAVT-TEYVTFHPQVLV----------------VI-LVG-ATLSTAVVW 247
PIN95153_1 hypotheti ------------------------GFVSLLIALA----FLQ--------------------WNKEVLT----------------LS-------------- 95
PIU22108_1 hypotheti ------------------------TIPIAIVLFI----FFL----C-------------KMTFVLDFF----------------LI-ILC-ILGISALIY 168
AAR39201_1 NEQ352    --------------------------VLFFATLF----VYA--------------------FYVNIDI-----------------------KVLVLLYF 112
OIR14399_1 hypotheti ---------FLRTEASDQELASLASFALMIGVLF----YLF----N-----FKWGDKPLRDPVLFFKV----------------AN-ITL-LMMALLQLF 224
OIR20963_1 hypotheti LNLSQNELVNP-------------EMLWDTVIIA----SLLTLVLIWLSSWFISNHGIPNMIFVLLFVLFSFNLYLVREVDNNSTMIMFMTIGIIISLIGS 307
OIR22371_1 hypotheti -----------VFSFTALTAKFYRGKPLRDPVLF----FGI--SSALTLILAYLTLAQNEVVYPEILW----------------NT-LIL-ASVLTFVVW 264
EGQ43935_1 putative  ------------------------GTAALFLTLP----FYT----L-------------EEPRALSTF----------------LT-TVPIWAVPGYLIL 164
```

```
MAG21679_1 hypotheti ------------------------GLIPSILIML----FIM--SNF-------------GNWPGFLGS----------------LA-IVAGITAVAVLGF 171
PIN85618_1 hypotheti ------------------------GLVISLPIVL----YEF---------------SIFRAWEGFFIV----------------IG-MVIAITLVVKIIV 171
PIN99249_1 hypotheti ------------------------GIVLCVPVVL----YHL------------LHLTHPIEFFFVFIL-----------------------TGIIVFAVK 172
AJF59838_1 hypotheti ------------------------GTLLAAPFVL----FDF---------------IQFVAWQYFTAI----------------II-----LTAVAVGIS 167
WP_042682145_1 DUF63 ------------------------GTILALYANY----LLV---------------THVKSWKPYELT----------------IF-----WTIVFVLPV 156
WP_013468033_1 DUF63 ------------------------GTILALYANY----LLV---------------THVKSFKPYELT----------------IF-----WTVIFILPV 156
WP_055281581_1 DUF63 ------------------------GSLLALYVNY----LLI---------------INAKSWRPYELT----------------VF-----YTIVFLLPI 155
WP_048160382_1 DUF63 ------------------------GTILALYANY----LLI---------------THVKSWEPYELT----------------LF-----YTVVFFLPI 155
WP_042701216_1 DUF63 ------------------------GTILALYANY----LLV---------------KNVKSWEPYELT----------------LF-----YTVIFLLPI 155
WP_004068659_1 DUF63 ------------------------GTILALYANY----LLI---------------TNVKSWKPYELT----------------IF-----YTLIFLLPI 155
WP_058946665_1 DUF63 ------------------------GTVLALYANY----LLI---------------TNVKSWKPYGLT----------------IF-----YTLIFLLPI 155
WP_014835499_1 DUF63 ------------------------GSILAVYVNY----LLF---------------TNAKCWKPYELT----------------LL-----HTGVSFAVV 155
WP_010884243_1 DUF63 ------------------------GSLLAIYANY----LLI---------------INVKCWKPYELT----------------LL-----HALVSFAVV 158
WP_013748129_1 DUF63 ------------------------GSILAGYSLY----LFI---------------INAKCWKPYELT----------------LL-----HTAISFIAV 154
WP_010867251_1 DUF63 ------------------------GTLLAGYANY----LLI---------------INAKCWKPYELT----------------LL-----HTLVSFVVV 154
WP_014733359_1 DUF63 ------------------------GVALALYANY----LLV---------------MNAKCWKPYELT----------------LL-----HTFISFVAV 155
WP_068319891_1 DUF63 ------------------------GAGLALYANY----ILF---------------KNVKCWEPYELT----------------LL-----HTAVSFGAV 155
WP_068576049_1 DUF63 ------------------------GAGLALYANY----LLF---------------KNVKCWEPYELT----------------LL-----HTAVSFVAV 155
WP_014013692_1 DUF63 ------------------------GALLALWANY----LLI---------------THAKSWEPYGLT----------------LL-----HTFVSWIPA 160
WP_014789520_1 DUF63 ------------------------GALLALGANY----LLV---------------THAKSWEPYELT----------------LI-----HTFVSWIPA 160
WP_088180676_1 DUF63 ------------------------GAVLVLWANY----LLV---------------THAKSWEPYGLT----------------LL-----HTFVSWIPA 160
WP_088864885_1 DUF63 ------------------------GALLALGANY----LLV---------------THAKSWEPYELT----------------LI-----HTFVSWIPA 160
WP_088856755_1 DUF63 ------------------------GLILALAVNY----VLV---------------THAKSWEPYELT----------------LI-----HTAISFIPV 160
WP_088865981_1 DUF63 ------------------------GAVLALWANY----LLV---------------THAKSWEPYGLT----------------LL-----HTFVSWVPA 160
WP_013906334_1 DUF63 ------------------------GTLLALWANY----LLV---------------TNARCWEPYILT----------------LI-----HTAISWAAV 156
WP_088862658_1 DUF63 ------------------------GSVLALWAIY----VWV---------------THARNWEPYALT----------------LL-----HTAVSFAVV 159
WP_088882956_1 DUF63 ------------------------GSLLALWALY----LLV---------------TSAKNWEPYGLT----------------LL-----HTAVSFVAV 159
WP_012571938_1 DUF63 ------------------------GSILALWANY----LLI---------------TNVKSWEPYELT----------------LI-----HTAVSFLAV 156
WP_068663933_1 DUF63 ------------------------GTALALWANY----LLV---------------TNAKSWEPYELT----------------LI-----HTAVSFLAV 156
WP_074631153_1 DUF63 ------------------------GALLALWANY----LLV---------------TNAKSWEPYGLT----------------LI-----HTAVSFAAV 156
WP_058939074_1 DUF63 ------------------------GALLALWANY----LLV---------------INAKSWEPYGLT----------------LI-----HTAVSFAAV 160
WP_088854681_1 DUF63 ------------------------GTILALWANY----LLV---------------THAKNWEPYELT----------------LV-----HTAVSWAVI 160
WP_088885426_1 DUF63 ------------------------GTVLALWANY----LLV---------------THAKSWRPYELT----------------LI-----HTIASWIPV 156
WP_011249694_1 DUF63 ------------------------GTILALWANY----LLV---------------THAKNWEPYELT----------------LI-----HTAVSFAAV 158
WP_062386972_1 DUF63 ------------------------GTILALWANY----LLV---------------THAKSWEPYELT----------------LL-----HTAVSWGVV 158
WP_010478887_1 DUF63 ------------------------GTVLALWANY----LLI---------------TNAKGWRPYELT----------------MI-----HTVISWAAV 156
WP_088858605_1 DUF63 ------------------------GTVLALWANY----LLF---------------ANAKDWRAYELT----------------ML-----HTLASWAVV 156
WP_042690028_1 DUF63 ------------------------GTVLALWANY----VLV---------------THAKDWRPYELT----------------LI-----HTAVSWAVV 158
WP_048150762_1 DUF63 ------------------------GTLLALWANY----QLA---------------IHAKSWEPYELT----------------MI-----HTLISWAVV 156
WP_048811076_1 DUF63 ------------------------GTILALWANY----LLI---------------THAKNWKPYELT----------------MI-----HTLISWAIV 156
WP_050003256_1 DUF63 ------------------------GTVLAAWANY----LLI---------------THAKSWEPYMLT----------------MV-----HTVVSFAVV 158
WP_062372100_1 DUF63 ------------------------GTILAVWANY----LLI---------------THAKNWKPYELT----------------MI-----HTVVSFVAV 156
WP_048165284_1 DUF63 ------------------------GLLLSTYPLY----LFL---------------TNMVRPQAYALT----------------LL-----YTVLFSVPV 158
WP_048148750_1 DUF63 ------------------------GLIVSAYPLY----LFF---------------THIVRPEAYAYT----------------LL-----YTIAFALPV 158
OYT53462_1 hypotheti ------------------------GLAGALYTGF----HLF---TY-------------IAFPERLVY----------------PP-----LMAVAVALF 140
KYC51152_1 hypotheti ------------------------GILGTIYFLA----NLI----P-------------FFNYPLRMA----------------VP-LLM-AFGITLAVY 142
KYC46003_1 hypotheti ------------------------GTIYFLANLI----PFF--------------------NYPLRMA----------------VP-LLM-AFGITLAVY 142
KYC48643_1 hypotheti ------------------------GTIYFLANLI----PFF--------------------NYPLRMA----------------VP-LLM-AFGITLAVY 142
KYC55321_1 hypotheti ------------------------GFIGMLYFLY----NLI-----------------PFFNHPLRIL----------------VP-LSM-ATIITLAVY 142
KYC57927_1 hypotheti ------------------------GFIGMLYFLY----NLI-----------------PFFNHPLRIL----------------VP-LSM-ATIITLAVY 142
KYC57171_1 hypotheti ------------------------GFIGMLYFLY----NLI-----------------PFFNHPLRIL----------------VP-LSM-ATIITLAVY 142
OIO20701_1 hypotheti ------------------------GLALWLPNFL----LLA----------------PMMRYLIYGAV----------------VL------ALVCFAYF 185
PIT83986_1 hypotheti ------------------------GAALCAGCVA----ALL---PL-------------MQYFDYGAI----------------IV-LGAAAVAALMAFI 207
OIO24760_1 hypotheti ------------------------GLVLFLTVFL----PLV---------------PLFHQWVELAQI----------------LV-----LAIIPLALY 167
OIO24701_1 hypotheti ------------------------GLILALGALL----PAL----------------GIAKYWLHGLL----------------AL-----LMGCSGILL 168
OIO27021_1 hypotheti ------------------------GAVLAIIPLA----ALL---------------PFYEHFGHAVAI----------------SV-----APLIALAAY 162
OIO26762_1 hypotheti ------------------------GWTAALLCVA----LLF---------------PLFENWLHAAAI----------------FA-VAF-AALKVFEYA 162
PIN95811_1 hypotheti ------------------------GWTAALLCVA----LLF--PLF-------------ENWLHAAAI----------------FA-VAF-AALKVFEYA 162
PIO01637_1 hypotheti ------------------------GWTAALLCVA----LLF--PLF-------------ENWLHAAAI----------------FA-VAF-AALKVFEYA 162
PIO02820_1 hypotheti ------------------------GAVLAIIPLA----ALL---------------PFYEHFGHAVAI----------------SV-APLIALAAYEFAW 166
PJD01038_1 hypotheti ------------------------GAVLAIIPLA----ALL--PFY-------------EHFGHAVAI----------------SV-APLIALAAYEFAW 166
PIZ91366_1 hypotheti ------------------------TAALLCVALL----FPL----F-------------ENWLHAAAI----------------FA-VAF-AALKVFEYA 162
WP_013100570_1 DUF63 ------------------------GLIPLIYFFI----IFL---------------QHVKTVLPIILS----------------FF-----LVTIFLGIA 157
WP_004590770_1 DUF63 ------------------------GFVIFLYFFS----IFL---------------THIHHLDALIYV----------------AM-----LVAVIYISI 157
WP_048196979_1 DUF63 ------------------------GAIPLLYFLS----IFF------------------KHIHHPEAL----------------LY-----VGILVGGVY 154
WP_015791538_1 DUF63 ------------------------GLTPLLYFLY----IFL---------------KHLVYLEAAIYV----------------GI-----LVGIFYCLA 157
WP_048202292_1 DUF63 ------------------------GVIPLLYFLF----VFL---------------QHIVHLEAILYV----------------GI-----LVGTFYYLV 157
WP_012981280_1 DUF63 ------------------------GLIPLLYFFG----IFL---------------QHITHLEALLYV----------------II-----LVTAFYYLV 157
WP_064496496_1 DUF63 ------------------------GLILLLYFLF----VFL---------------QHITHLEAILYV----------------GI-----LVGIFYYLV 157
WP_011972741_1 DUF63 ------------------------SIIMAVLPML----YFG------------------AIFLTNIVY----------------FG-TLLQILLIIAIIY 156
WP_013798289_1 DUF63 ------------------------AIPFALLPII----YFL-----------PDFLNRIVHWEALLYV----------------SL-----ILFPTYILT 157
WP_013181100_1 DUF63 ------------------------SIVMAVIPIL----YFL-----------YEFVQRIVHIEALTLV----------------AL-----ILAISYILA 159
WP_013867628_1 DUF63 ------------------------AIIMALIPTV----YLL----S-----IFI--NNMVHMNALFYV----------------LG-IV--AIVYGIVIY 162
WP_018153391_1 DUF63 ------------------------ASLMALVPII----YLL--PEF-----LKRITHLEALFYTLVVL----------------FP--------TYFIAI 160
WP_011170272_1 DUF63 ------------------------SIAMAVVPIL----YML----F-----EFSS----RITHPEAIV----------------YV-SGI-LVLSYYLLI 160
WP_012066151_1 DUF63 ------------------------SIIMAIVPIL----YMF-----------FEFSTRITQIESVAYV----------------FG-----ILSTSYLLA 159
ODS43013_1 hypotheti ------------------------GLVPMVYNVF----LIM--ANI-------------ENAAPLFYV----------------SA-----ALVISGAFL 166
OIQ06085_1 hypotheti ------------------------GCLLFGLNLVLIAVFLY-SHSATANFGAFIVFLIFAMWAIIIYA----------------IIKLNPKFSIIGGVLV 183
PIN67069_1 hypotheti ------------------------GCLLFGLNLVLIAVFLY-SHSATANFGAFIVFLIFAMWAIIIYA----------------IIKLNPKFSIIGGVLV 183
PIV28099_1 hypotheti ------------------------GCLLFGLNLVLIAVFLY-SHSATANFGAFIVFLIFAMWAIIIYA----------------IIKLNPKFSIIGGVLV 183
PJC13070_1 hypotheti ------------------------GCLLFGLNLVLIAVFLY-SHSATANFGAFIVFLIFAMWAIIIYA----------------IIKLNPKFSIIGGVLV 183
PIZ29927_1 hypotheti ------------------------GCLLFGLNLVLIAVFLY-SHSATANFGAFIVFLIFAMWAIIIYA----------------IIKLNPKFSIIGGVLV 183
PKP60688_1 hypotheti ------------------------GCLLFGLNLV----LIA---------MFLYSHSATANFGVFVVF----------------LI-FAMCVIVIYALMK 170
WP_012956954_1 DUF63 ------------------------GVILLIPNII----MIP-----------------RLNIIPVIYV----------------LI-----TWIIASSIF 154
WP_067148430_1 DUF63 ------------------------GLLFLLPNII----LFS--NLN-----FTAIFYIFITWIFVSLI----------------FI-IIS-LLVLYIVNY 167
WP_080460538_1 DUF63 ------------------------GLILAIINLI----NIQ----------SINLIAFSQVILSWLVM----------------TV-----IFIIVGIFW 160
WP_010877090_1 DUF63 ------------------------GSVLAAPNLI----SIR----H-------------IDPVPFLSV----------------LG---V-FSVFSAAFY 155
WP_013294901_1 DUF63 ------------------------GAVLSIPNII----SMQ-----------------KINPVPFFSV----------------LG-----VFMASAFLF 154
WP_010877090_1 DUF63 ------------------------GSVLAAPNLI----SIR----H-------------IDPVPFLSV----------------LG---V-FSVFSAAFY 155
BAZ99473_1 hypotheti ------------------------GSVLAAPNLI----SIR----H-------------IDPVPFLSV----------------LG---V-FSVFSAAFY 155
PKL66404_1 hypotheti ------------------------GVLICIPNIL----SLQ-----------------SINFTAMEYV----------------LI-----FWGMFTGIF 162
WP_023991171_1 DUF63 ------------------------GVVLCIPNIY----YTQ-------------------HINPLIVF----------------QV-LGP-WALLTAIFY 156
WP_048072898_1 DUF63 ------------------------GAAMCVPNIY----YAQ-------------------HINPVVVF----------------QV-LAP-WALLTAVFV 156
WP_100905180_1 hypot ------------------------GVVMCLPNLY----YIQ----------HLDLGVVFQVMGSWALL----------------SV-----PFILLGFKW 162
WP_100907231_1 hypot ------------------------GVVMCLPNLY----YIQ----------HLDLGVVFQVMGSWVLV----------------SV-----PFILLGFKW 162
```

```
WP_100907231_1 hypot ------------------------GVVMCLPNLY----YIQ----------HLDLGVVFQVMGSWVLV----------------SV-----PFILLGFKW 162
WP_048081925_1 DUF63 ------------------------GTVLCIPNIL----SIN-----------------HLNWVTFFEV----------------VG----TWAVMTSIFA 155
WP_048081925_1 DUF63 ------------------------GTVLCIPNIL----SIN-----------------HLNWVTFFEV----------------VG----TWAVMTSIFA 155
WP_069585525_1 DUF63 ------------------------GTMLCIPNIL----SIN-----------------HLNWVTFFEV----------------VG----TWAVMTSIFA 155
WP_069585525_1 DUF63 ------------------------GTMLCIPNIL----SIN----H-------------LNWVTFFEV----------------VG-T---WAVMTSIFA 155
WP_013643671_1 DUF63 ------------------------GAVICIPNII----FLG-----------------PINVVAALQV----------------LG---I-WALISSIFV 155
WP_013824571_1 DUF63 ------------------------GAVMCIPNII----YLG----P-------------INWVAFFEV----------------IG---I-WAIISSIFV 155
WP_048192110_1 DUF63 ------------------------GAVICIPNIL----FMG----------------PINIIATLQVV----------------GV-----WALISSIFV 159
WP_081810122_1 DUF63 ------------------------GIILSLAGVI----ILI----Y-----NSDIGFPEPDILLYILL----------------PA-----LALTEFIKR 191
WP_097298965_1 DUF63 ------------------------GLILSLAGVV----ILI--------------FNTSSTWNPGILV-----------------------YALVPAIAL 160
PIO00292_1 hypotheti ------------------------GSVLWLLALF----GFE----L-------------RNPLSLLYI----------------LG-LFS-GILCGLFVL 157
WP_011833679_1 DUF63 ------------------------GIISSLIVAG----FLV----W-----FGFTQATNVDVGAGILV----------------LL-----IAAAAVAL 163
WP_042698332_1 DUF63 ------------------------GVISSLITAG----FLI----W-----FGFARAAEVDILAGLWV----------------FL---I-AAAASIALW 164
WP_042698332_1 DUF63 ------------------------GVISSLITAG----FLI----W-----FGFARAAEVDILAGLWV----------------FL-I---AAAASIALW 164
WP_011448164_1 DUF63 ------------------------GIALTLVTLG----FLT----W-----FGLTNAHIDTLVGLAIL----------------TM-----ATVTTGLLY 166
WP_007314214_1 DUF63 ------------------------GIAGAAVTGL----VLV----------SFGIRNGIIALDVLFFI----------------LL---L-AVATTAAVW 171
WP_042705632_1 DUF63 ------------------------GVAGVAVSLI----PLI----W-----VGITQTQ-INLFVFFAI----------------CA-----MGLATSAAL 165
WP_013329717_1 DUF63 ------------------------GIIACAISLV----PLV----W-----YGLTVTQ-INFFVMFSI----------------LG-I---ACISSLAVW 166
WP_004077722_1 DUF63 ------------------------GIIAVIASLI----PLV----W-----YGITQ---SEIHISVMV----------------II-LAM-AFASSAAVW 166
WP_048150927_1 DUF63 ------------------------GIIACAISLV----PLV----W-----YGLTVTQ-INFFVMFSI----------------LG-I---ACISSLAVW 166
WP_012107531_1 DUF63 ------------------------GCMSVFVVSL----ILL----------SWGMAHAHVDLFILAII----------------LL-----MALAATVLV 171
PKL70081_1 hypotheti ------------------------GLFTSFCVVL----LLT----------SWGLTHTRIDLFVLAII----------------PL---M-AIVATALVF 166
WP_015285007_1 DUF63 ------------------------GVFSVVVASL----ILI----------AWSTTHHGIDFYVLGVI----------------PL---M-AVVATALVW 172
PKL64570_1 hypotheti ------------------------GTFALFVACL----VLV----------AWSTTHHGIDLFILCII----------------PF-----MALCATTIV 171
WP_015286404_1 DUF63 ------------------------GTLACTISAA----FLV----W-----WGLT---YSQVAVSVLL----------------II-MGL-AIVISAGVW 166
WP_012617618_1 DUF63 ------------------------GIIACLISGA----VLC----W-----YGLTVTR-IDFGVLLTI----------------LL---M-AATTTTLVW 166
WP_014867374_1 DUF63 ------------------------GIVACLLSTA----ALV----W-----FGLS-ETTIALDVLVAI----------------LA-L---ASVSSLALW 166
CVK33108_1 conserved ------------------------GIVACLLATA----ALV----W-----FGLS-ETTIALDVLVAI----------------LA-L---ASVSSLALW 166
WP_066956331_1 DUF63 ------------------------GVVACLAAGA----ALI----W-----FGLT-ETQIALNVLVAI----------------LT-L---AAVTSLALW 166
WP_011844879_1 DUF63 ------------------------GIAACLVAAS----ALV----------WFGLTEATIALDVLAVI----------------LV-----FASVTSLAL 165
WP_048181868_1 DUF63 ------------------------GVAACLVAAS----ALV----------WFGLTEATIALDVLAVI----------------LV-----LASVTSLAL 165
WP_067073014_1 DUF63 ------------------------GVVTCLLATA----ALV----W-----FGLT-EATIALDVLAAI----------------LA-L---ASVTSLALW 166
PKL62929_1 hypotheti ------------------------GVVACLLATA----ALV----W-----FGLT-EATIALDVLAVI----------------LA-L---AAVTSLALW 166
WP_004039890_1 DUF63 ------------------------GVALSAAFFV----LLI----------LWGAANTRVDLVVLLAI----------------PA---M-AAVSSAAVF 166
WP_067049225_1 DUF63 ------------------------GIALSVATAL----LLL----------GWGAVNTRIDFVVLFAI----------------PA-M---AAVSTAAVW 166
KYK37884_1 hypotheti ------------------------GIILFLVILG----ILA-------------MNAAVFNSKGFIYI----------------VV-----TTVAVAVIV 154
KYK28019_1 hypotheti ------------------------GLGLFIFILG----ILI----A-----NAQA----FNYKGFIYI----------------VV-----AAVAVSVIV 154
OYT57835_1 hypotheti ------------------------GIILDIIAVS----FLR----I-------------VNLDALLFI----------------LG-----ISLVWLLVV 157
OYT33350_1 hypotheti ------------------------GSVLTIISVV----LIF----------------SNLEIRNFWVI----------------PL-----GFALSLLLT 158
WP_012964745_1 DUF63 ------------------------GIILAAIPPI----ILL-------------TNLRIENWWVFPAS----------------IA-----LALIFTLIF 155
WP_048091573_1 DUF63 ------------------------GAVLSLSVLA----FLL----L-----NLRV----ENWWIFPAA----------------IT-----IAGTAVLIY 154
WP_048096399_1 DUF63 ------------------------GIVLSLIVLA----FLF-------------THLEVENWWVIPTA----------------LT-----IAATITAAY 154
WP_012940099_1 DUF63 ------------------------GVILSVLTLS----FLL-----------LNLEIIHIEVLPISAA-----------------------IATFATALF 157
WP_010877969_1 DUF63 ------------------------GLVPAVGVLI----LLF-------------ANLPVVNWWVIPAG----------------VG-----LAAAFTAVF 170
WP_013682839_1 DUF63 ------------------------GLVLSCGVLV----LLF------------SNLEVVNAWILPSTI----------------IL-----ASLFCSAYR 159
WP_015591141_1 DUF63 ------------------------GVFLSAAVLA----MLF------------SSLEVIHGWLIPVTL----------------TA-----AFLFAAAYH 158
WP_012035887_1 DUF63 ------------------------GVAGIVIVLG----MLF----T----------QEVRLWWAPVLI----------------VL---L-AATFTASIY 169
WP_014404626_1 DUF63 ------------------------GIAGIILVFA----ALL----L-----TQPI----SVWWAPIVV----------------VS---L-AFTFTGAIY 164
BAI60262_1 conserved ------------------------GVAGVIVVWG----TLL----L-----TQPI----RVWWAPIVV----------------FA-L---AFTFTGVVY 164
WP_042684156_1 DUF63 ------------------------GAVLDIGLLS----LLL-------------HTASAEAWVPMAIV----------------GL-----SIVAFLAVF 170
OFV68024_1 membrane  ------------------------GIFWAFLNLL----ILV-------------IAEDVVNLWVLPAV----------------IL-----IALGTITPF 124
WP_013720253_1 DUF63 ------------------------GLAWTLFNLM----LLF----G-----VGLE----NGWVIGAVF----------------AL-----GSALTGGLL 160
WP_014587756_1 DUF63 ------------------------GLLWTAGNLA----LLA----P-----LGVE----RSTIPIAIL----------------AL-----GVAATALVA 157
ABK14947_1 Protein o ------------------------GISWSALNLL----LLI-------------REGVQHAWVPLAVV----------------AL-----GSSITGCVY 162
OKY79140_1 putative  ------------------------GSAIAILNLA----ILT----------SWGINNPGTNPLVPLKI----------------IG-TAI-AVFSPIAAT 171
WP_086637003_1 DUF63 ------------------------GCILALFLTI----YLS----Y-----IAVGSQHTRLSIPFEII----------------VV-----TLTITGLAY 170
WP_048089097_1 DUF63 ------------------------GVVWTVINIA----VLL----S-----AGEI----ENPVPAAVI----------------LI---L-GAVSTYIAY 167
WP_097298272_1 DUF63 ------------------------GAVWGFLNIS----ALL----A-----VGSI----ENPYPAAVI----------------FI-L---AISSTGIVY 168
WP_013897788_1 DUF63 ------------------------GSVWFILNIS----TLF----------FVATVS--VPLAPLFVV----------------IS-----ATAISYAIY 169
WP_015323982_1 DUF63 ------------------------GTLWFVFNIF----LLL---------------SIQKVALPMVLI----------------YI-LGL-GILVTSIVY 169
WP_011499115_1 DUF63 ------------------------GLCWFAINII----TLL------------VVEDITRPFILVAIV----------------LL-----GTVLAGSIY 169
WP_013037846_1 DUF63 ------------------------GGLWFLANLT----ALL----Y-----LEDI----VRPDALVLI----------------LV---I-GTLVAFSIY 169
WP_048205640_1 DUF63 ------------------------GLGWFALNII----TLL------------MVEDIERPFTLIAIL----------------SS-----GTLFAGAVY 169
WP_072561629_1 DUF63 ------------------------GGLWFLSNLY----ALL----Y-----FEDI----VRPDALILI----------------LV-I---GTLVAFSIY 169
WP_072360082_1 DUF63 ------------------------GGLWFLSNLY----ALL----Y-----FEDI----VRPDALVMI----------------LV-I---GTLVAFSIY 169
WP_096711793_1 DUF63 ------------------------GGLWFLFNLY----ALL----Y-----FEDI----VRPDALVLI----------------LV-I---GTLIAFSIY 169
ODV50598_1 hypotheti ------------------------GGLWFLFNLY----ALL----Y-----FEDI----VRPDALVLI----------------LV-I---GTLIAFSIY 169
WP_013194042_1 DUF63 ------------------------GSLWFIFNVS----ILL----Y-----FENV----VNLYVPYFV----------------IG---A-ATLLVFIVY 169
WP_048178067_1 DUF63 ------------------------GLIWFFVNLI----TLL----Y-----FEDV----VAAYVPVFV----------------LV---A-GTGLTFIFY 171
WP_048127111_1 DUF63 ------------------------GLAWFLINLG----VLL----H-----FENI----VVAYVPVFV----------------IG---A-GTGLTFAFY 171
WP_011023594_1 DUF63 ------------------------GLVWFFLNLG----VLL------------HYENVVAAYVPVFVI----------------GV-----GTGLTLVFY 175
WP_048184587_1 DUF63 ------------------------GLVWFFLNLG----TLL------------HFENVVAAYVPVFVI----------------GA-----GTGLTLVFY 171
WP_011032542_1 DUF63 ------------------------GLAWFLLNLG----ALL------------YFEDIVVPYVPVFVL----------------GA-----GTGLTFVFY 171
WP_011032542_1 DUF63 ------------------------GLAWFLLNLG----ALL------------YFEDIVVPYVPVFVL----------------GA-----GTGLTFVFY 171
WP_048129141_1 DUF63 ------------------------GLAWFLLNLG----TLL------------YFENIVAAYVPIFVI----------------GA-----GTGLTLAFY 171
WP_048129141_1 DUF63 ------------------------GLAWFLLNLG----TLL------------YFENIVAAYVPIFVI----------------GA-----GTGLTLAFY 171
WP_048169891_1 DUF63 ------------------------GLAWFLLNLG----TLL------------YYENIVVAYVPVFVL----------------GA-----GTGLTFTFY 171
WP_048137175_1 DUF63 ------------------------GLAWFLLNLG----TLL------------YYENIVVAYVPVFVL----------------GA-----GIGLTFTFY 171
WP_048137175_1 DUF63 ------------------------GLAWFLLNLG----TLL------------YYENIVVAYVPVFVL----------------GA-----GIGLTFTFY 171
WP_048137175_1 DUF63 ------------------------GLAWFLLNLG----TLL------------YYENIVVAYVPVFVL----------------GA-----GIGLTFTFY 171
WP_048137694_1 DUF63 ------------------------GLAWFLLNLG----TLL------------YFENIVVAYIPVFVI----------------GA-----GIGLTFIFY 171
WP_048168126_1 DUF63 ------------------------GLAWFSVNLI----TLL----Y-----FENV----VASYVPLFV----------------LG---A-GIGLTFTFY 171
WP_048118082_1 DUF63 ------------------------GLAWFFINLA----VLL------------HFDNIVYPYVPVFVI----------------VA-----GIGLTFIFY 171
WP_048158120_1 DUF63 ------------------------GLAWFFINLA----VLL------------HFDNIVYPYVPVFVI----------------VA-----GIGLTFIFY 171
WP_011305329_1 DUF63 ------------------------GLALFFINLA----VLL------------HFDTIVYPYVPVFVI----------------VA-----GIGLTFIFY 171
WP_054298619_1 DUF63 ------------------------GLAWFSVNLI----TLL------------YFENVVASYVPLFVL----------------GA-----GIGLTFTFY 171
ALK05385_1 hypotheti ------------------------GLAWFSVNLI----TLL------------YFENVVASYVPLFVL----------------GA-----GIGLTFTFY 171
WP_015052897_1 DUF63 ------------------------GSAWFIANIA----YLL------------HLEDLVLPWVPVFVI----------------VS-----ASALLYILY 168
WP_023846134_1 DUF63 ------------------------GLAWFAVNLS----ILF------------YLEDLTRPWVPIFVL----------------GA-----ASLIVILLK 168
OIN88451_1 hypotheti ------------------------GVMFSGPLLG----FFN-------------IQNALGLWYVLLWL----------------VP-------------W 150
PIX50278_1 hypotheti ------------------------GVMFSGPLLG----FFN-------------IQNALGLWYVLLWL----------------VP-------------W 150
PIW41402_1 hypotheti ------------------------GVMFSGPLLG----FFN-------------IQNALGLWYVLLWL----------------VP-------------W 150
PIY35178_1 hypotheti ------------------------GVMFSGPLLG----FFN-------------IQNALGLWYVLLWL----------------VP-------------W 150
PJB74886_1 hypotheti ------------------------GVMFSGPLLG----FFN-------------IQNALGLWYVLLWL----------------VP-------------W 150
PIZ33651_1 hypotheti ------------------------GVMFSGPLLG----FFN-------------IQNALGLWYVLLWL----------------VP-------------W 150
```

```
WP_048165029_1 DUF63 ------------------------GWALVGLDLL----LLI-----------THLDKVNLTAEPLKYF----------------IP-----FLLVAVGVI 158
WP_042681046_1 DUF63 ------------------------GWALVGFDFV----MLL-----------THLDKVHFTWIVLKYF----------------IP-----AVIIAEGII 158
WP_013467319_1 DUF63 ------------------------GWALVGFDFV----MLL-----------THLDKVHFTWIVLKYF----------------IP-----ALIIAESVI 158
WP_048152160_1 DUF63 ------------------------GWTLVGVEFI----LLL-----------SHLDKIEFNTRVLLYF----------------VP-----LASIALAVI 158
WP_015849008_1 DUF63 ------------------------GWALVGGELL----LLL-----------FNLEKVQFNFDTLKYF----------------IP-----FATIALMTI 158
WP_004069276_1 DUF63 ------------------------GWTLVGGEVL----LLL-----------FNLDKVSFNFEVLKYF----------------IP-----FASLALAVV 158
WP_058946638_1 DUF63 ------------------------GWVLVGGEVL----LLV-----------FNLDKVSFNFEVLKYF----------------IP-----FVSLALSLI 158
WP_042701551_1 DUF63 ------------------------GWGLVGGEVL----LLL-----------INLDKVRFNFEVLKYF----------------IP-----FALLALVVV 158
WP_055282692_1 DUF63 ------------------------GWSLVGMESL----LLL-----------FNIDKVDFNLTVLKYF----------------IP-----FATIALITI 158
WP_013906023_1 DUF63 ------------------------GWGLVAVDLA----ALL------------ARGDVDFNWAVLKYF----------------LP-----ALAVAELAV 156
WP_011013158_1 DUF63 ------------------------GWGLVMVDLA----AMS-----------ANISKINFDFAVLKYF----------------IP-----FLILAELVV 157
WP_014733166_1 DUF63 ------------------------GWGLVMVDFA----AMG-----------ANLNKIHFNLEVLKYF----------------IP-----FVALAEATV 157
WP_068322889_1 DUF63 ------------------------GWGLVMVDFA----AMG-----------ANLEKMNFNFTVLKYF----------------IP-----FLILAELAV 157
WP_068575625_1 DUF63 ------------------------GWGLVMVDFA----AMG-----------ANLEKMHFNFTVLKYF----------------IP-----FLILAELAI 157
WP_010885960_1 DUF63 ------------------------GWGLVLIDVA----ALL-----------VNLQKVTFNFQVLKYF----------------IP-----FVTIAEVVI 157
WP_010867401_1 DUF63 ------------------------GWGLVLVDVA----ALS-----------ANIGKVSFRLEVLKYF----------------IP-----FVIIAELVI 157
WP_013747987_1 DUF63 ------------------------GWGLVLIDVA----ALS-----------ANLQKVSFNLIVLKYF----------------IP-----FLILAELVV 158
WP_068664049_1 DUF63 ------------------------GIVLLLFNVF----VLG-----------ISYSKVSFNWEVLKYF----------------IP-----ALVVAETSI 160
WP_011249739_1 DUF63 ------------------------GWVLVGGLVF----III-----------INWGKLSVRWEYFKYF----------------IP-----SLAVSEAFI 161
WP_062386854_1 DUF63 ------------------------GWVLVGGLVF----III-----------INWGKLSARWEYFKYF----------------IP-----SLAVSEAFI 161
WP_050003781_1 DUF63 ------------------------GWVLVAGLLF----IMV-----------INWNRVSVRWEYFKYF----------------LP-----ALLVSEAVI 161
WP_042690699_1 DUF63 ------------------------GWVLVAGLLF----IMV-----------INWDRVSVRWDYFKYF----------------LP-----SLLVAESFI 161
WP_088885212_1 DUF63 ------------------------GLLLLGGLLF----LLL-----------INLDKTDFRWKYFVYF----------------IP-----SFLAAEGVI 161
WP_010478519_1 DUF63 ------------------------GLLLLGGLLF----LLL-----------INLGRTDFRWEYFVYF----------------IP-----SFLAAEGAI 161
WP_088858240_1 DUF63 ------------------------GLLLVGGLLF----ILI-----------INLDKVDFRWSYFYYF----------------LP-----ALLAAEAFI 161
WP_015859016_1 DUF63 ------------------------GFLLVGGLLF----ILI-----------INLDKVNFRWEYFKYF----------------IP-----SLLVAEAFI 161
WP_014121940_1 DUF63 ------------------------GFLLLAGLVF----ILI-----------INLDKVNFRWDYFKYF----------------IP-----SLLAAEAFI 161
WP_062373911_1 DUF63 ------------------------GFLLVGGLLF----ILV-----------INLDKVNFRWEYFKYF----------------IP-----SFLAAEFFI 161
WP_088882924_1 DUF63 ------------------------GWVLVGGLLF----ILL-----------INHERVNLNLKPLEYF----------------IP-----ALLIAEAFV 161
WP_088862911_1 DUF63 ------------------------GWVLVGGLLF----VLI-----------INLDKVSFNWRVLEYF----------------IP-----ILLLAEASV 161
WP_012572006_1 DUF63 ------------------------GYVLLGGLVF----ILL-----------INLDKVNFRWEVFKYF----------------VP-----ILIAAEAFI 161
WP_088854707_1 DUF63 ------------------------GWILVGGLLF----VLV-----------INLDRVSFNWKALGYF----------------VP-----FLIIAEAFI 161
WP_014789474_1 DUF63 ------------------------GWILVGGLLF----VLV-----------INLDNVSFNPAVFRYF----------------IP-----ALLVSEAFI 160
WP_088180728_1 DUF63 ------------------------GWVLLGGLVF----VLI-----------INLGKVDFNPEVFKYF----------------IP-----ALIIAEAFI 161
WP_088864937_1 DUF63 ------------------------GWVLLGGLVF----VLI-----------INLDKVSFNPEVFRYF----------------IP-----ALVVAEGFI 160
WP_055429686_1 DUF63 ------------------------GWVLLGGLVF----VLV-----------INLGKVNFNTEVFKYF----------------IP-----ALIIAEGFI 161
WP_088865932_1 DUF63 ------------------------GWVLLGGLVF----VLI-----------INRGKVNFNPEVFKYF----------------IP-----ALIIAEGFV 161
WP_014013642_1 DUF63 ------------------------GWVLVGGLLF----LLA-----------INLDKVSFNWEVFKYF----------------IP-----ALVIAEGAV 161
WP_088856804_1 DUF63 ------------------------GWILVGGLLF----VLV-----------INLDKVNFNWEVLKYF----------------VP-----AIVIAEAVI 161
WP_058939475_1 DUF63 ------------------------GWVLVGGLLF----LLV-----------INLDKVSFNWEVFKYF----------------IP-----ALVIAEGAV 160
EHR77276_1 conserved ------------------------GTASVVLGLGHWAQFLA--TPW---AQESGRVLESTPIWPLFVV----------------LG---IPAVVCFFMYR 298
OUV40025_1 hypotheti ------------------------GELNIRIRLV----PLL---------------CLALLFMWALLF----------------RPGYTEHDMGMAWVYI 217
MBJ52984_1 hypotheti ------------------------GNTFCILTLGFWFQFLI--EPW-----GTNSS---QEFWPIVVS----------------LG-IPS-ITVILLYRF 289
PDH23744_1 hypotheti ELSGLFSNVQRMVYLTGVG-----GSLIFFGALA----SFT----I-----DSP--PATDRIWPLVVV----------------IG---V-PAVVCYWMY 277
PDH25468_1 hypotheti ------------------------GVGGSLVLFGGLASYMS----W--------APSDQLNFWPVVVV----------------IG-LPI-VVAYLMIQQ 283

WP_048201856_1 hypot --------------------------------------------------------------------------------------------TSFSILTG 107
WP_048150344_1 hypot --------------------------------------------------------------------------------------------LGAIIV-- 140
AOV95360_1 hypotheti ---------------------LIIGK--RKTS------------------------------------------------------------LYLFPVLT 153
EOD42420_1 Uncharact ---------------------LSYTE----------------------------------------------------------------K-LFYLSIFG 164
AOV95164_1 hypotheti ---------------------LVTRR--NSYSR---------------------------------------------------------K-EYVLIAFS 196
EGQ40074_1 putative  ---------------------AVVRGTEFDRP------------------------------------------------------------VFFIAAFS 194
KYK23067_1 hypotheti ---------------------V-YRK--HESVAV--YKHP----------------------------------------------------LNLSMLGG 306
WP_084383883_1 DUF63 -------------------------------------------------------------------------------------------------TLG 190
WP_049984677_1 DUF63 ---------------------WGVR---FVDVAHLRHP------------------------------------------------------MLLLAVFG 244
WP_103428047_1 hypot ---------------------WGAGVIDVASLRH--------------------------------------------------------P-LFLLAVFG 243
WP_004048647_1 DUF63 ---------------------WGASFVNVTHLRH--------------------------------------------------------P-LILLAVFG 243
WP_004594600_1 DUF63 ---------------------WGASFVNITHLRH--------------------------------------------------------P-LILLAVFG 243
WP_004594600_1 DUF63 ---------------------WGASFVNITHLRH--------------------------------------------------------P-LILLAVFG 243
WP_050050397_1 DUF63 ---------------------WGTSFANITHLRH--------------------------------------------------------P-LILLAVFG 243
WP_004594600_1 DUF63 ---------------------WGASFVNITHLRH--------------------------------------------------------P-LILLAVFG 243
WP_050050397_1 DUF63 ---------------------WGTSFANITHLRH--------------------------------------------------------P-LILLAVFG 243
WP_049983690_1 DUF63 ---------------------WGASLVDIAHLRH--------------------------------------------------------P-LSLLAVFG 243
WP_103428162_1 hypot ---------------------WGTN---FVDGTQ--LRHP----------------------------------------------------LFLLAVFG 243
WP_009378268_1 DUF63 ---------------------WGASVVDVAHLRH--------------------------------------------------------P-LFLLAVFG 243
WP_015763176_1 DUF63 ---------------------KAIEA--AWPELT--DATA--------------------------------------------------Y-IGFFVLWG 279
WP_018259238_1 DUF63 ---------------------KAIEA--AWPELT--DATA--------------------------------------------------Y-IGFFVLWG 279
WP_004592615_1 DUF63 ---------------------RLFES--YRPAWN--DGTG--------------------------------------------------Y-IGLLVIWG 273
WP_004518092_1 DUF63 ---------------------WVFET--YRPALN--DGTG--------------------------------------------------Y-IGLLVIWG 273
WP_014040450_1 DUF63 ---------------------RLFES--YRPALN--DGTG--------------------------------------------------Y-IGLLVIWG 273
WP_014040450_1 DUF63 ---------------------RLFES--YRPALN--DGTG--------------------------------------------------Y-IGLLVIWG 273
WP_008309018_1 DUF63 ---------------------RLFES--YRPALN--DGTG--------------------------------------------------Y-IGLLVIWG 273
WP_014040450_1 DUF63 ---------------------RLFES--YRPALN--DGTG--------------------------------------------------Y-IGLLVIWG 273
WP_014040450_1 DUF63 ---------------------RLFES--YRPALN--DGTG--------------------------------------------------Y-IGLLVIWG 273
WP_005534235_1 DUF63 ---------------------RLFES--YRPAMN--DGTG--------------------------------------------------Y-IGLLVIWG 273
WP_004961153_1 DUF63 ---------------------RLFES--YRPVLN--DGTG--------------------------------------------------Y-IGLLVIWG 273
WP_004961153_1 DUF63 ---------------------RLFES--YRPVLN--DGTG--------------------------------------------------Y-IGLLVIWG 273
WP_101350151_1 hypot ---------------------RLFES--YRPALN--DGTG--------------------------------------------------Y-IGLLVIWG 273
WP_053968273_1 DUF63 ---------------------RLFES--YRPALN--NGTG--------------------------------------------------Y-IGLLVIWG 273
WP_058995687_1 DUF63 ---------------------RLFES--YRPALN--DGTG--------------------------------------------------Y-IGLLVIWG 273
WP_015790673_1 DUF63 ---------------------VLADR--FAPYIN--SGTG--------------------------------------------------Y-VGLVVLWA 274
WP_008524115_1 DUF63 ---------------------ALADR--FYPEIN--SGTG--------------------------------------------------Y-VGLLVIWA 274
WP_075936143_1 DUF63 ---------------------RLAQR--YKPGIN--AGTG--------------------------------------------------Y-VGLVVIFG 272
WP_020446311_1 DUF63 ---------------------TLTEE--YWPEVN--VGTG--------------------------------------------------T-VGLLVVWG 269
WP_049898450_1 DUF63 ---------------------LAIER--IEPAIN--AGTG--------------------------------------------------F-VGLAVIWA 271
WP_006077061_1 DUF63 ---------------------LAIDR--IEPAIN--AGTG--------------------------------------------------F-VGLAVLWA 271
WP_049996700_1 DUF63 ---------------------YAIDR--LAPALN--TGTG--------------------------------------------------F-IGLAILWA 271
EMA38628_1 hypotheti ---------------------LAIDR--LAPALN--AGTG--------------------------------------------------T-IGLAILWA 271
WP_049992561_1 DUF63 ---------------------WLAER--FAPEIN--RGTG--------------------------------------------------Y-IGLVVVWS 272
WP_010903025_1 DUF63 ---------------------AAARR--WLPSVT--SGTE--------------------------------------------------T-VGAVVLWG 270
WP_009760947_1 DUF63 ---------------------WVVRR--SLPTVY--SGTE--------------------------------------------------N-VGALILWG 269
WP_059057097_1 DUF63 ---------------------YYVRR--SLPDVA--SGTE--------------------------------------------------F-AGAVVLWA 269
WP_058983522_1 DUF63 ---------------------YAVRR--YLPDVA--SGTE--------------------------------------------------L-AGAVVLWA 269
WP_071932813_1 DUF63 ---------------------WALRK--YAPAVA--DGTP--------------------------------------------------V-VGAVILWA 265
WP_050048584_1 DUF63 ---------------------VPIRR--WFPWIA--AGTP--------------------------------------------------R-LGAVVLWA 265
```

```
WP_014051341_1 DUF63 ---------------------YVLER--FVPYVN--EGTR--------------------------------------------------G-VGVMVLFA 269
WP_079233299_1 DUF63 ---------------------YAIEA--VAPEIN--EGTG--------------------------------------------------L-AGAMVLVA 267
WP_053948572_1 DUF63 ---------------------YAIET--FAPSIN--EGTG--------------------------------------------------I-AGGMVLVA 267
WP_049980835_1 DUF63 ---------------------YAIEA--FAPEIN--EGTG--------------------------------------------------L-AGAMVLVA 267
KPN31749_1 hypotheti ---------------------YAIEA--FAPKIN--EGTG--------------------------------------------------L-AGAMVLVA 217
AGB16040_1 putative  ---------------------WTIVQ--FAPQLN--EGTG--------------------------------------------------Y-MGIPIIWA 265
WP_007696755_1 DUF63 ---------------------FTIQR--FAPELN--EGTR--------------------------------------------------Y-MGVLIIWA 265
WP_015322904_1 DUF63 ---------------------LAIER--FAPGLN--AGTR--------------------------------------------------Y-MGIVIIWA 266
WP_076581869_1 DUF63 ---------------------AGIQR--FAPSLN--RGTG--------------------------------------------------Y-MGIVVIWA 266
WP_005559622_1 DUF63 ---------------------LGIER--FAPDLN--AGTR--------------------------------------------------Y-MGIVIIWA 266
WP_006067640_1 DUF63 ---------------------MGIQR--FAPELN--RGTG--------------------------------------------------Y-MGIVVIWA 261
WP_008164554_1 DUF63 ---------------------WGITR--FAPELN--RGTE--------------------------------------------------T-MGIVVIWA 266
WP_006088112_1 DUF63 ---------------------WGIQR--FAPGLN--AGTG--------------------------------------------------T-MGIVVIWA 266
WP_012943247_1 DUF63 ---------------------LLIGE--FAPEIN--RGTE--------------------------------------------------Y-MGVVVIWA 265
WP_008895026_1 DUF63 ---------------------LLIGR--FAPEIN--RGTE--------------------------------------------------Y-MGVVVIWA 265
WP_098727043_1 DUF63 ---------------------VALER--FAPELN--RGTG--------------------------------------------------Y-MGIVVIWA 266
WP_049990861_1 DUF63 ---------------------VAIER--YAPELN--RGTR--------------------------------------------------Y-MGIVVIWA 264
WP_008013001_1 DUF63 ---------------------FAIER--FAPELN--RGTR--------------------------------------------------Y-MGLVVIWA 266
WP_006180566_1 DUF63 ---------------------FGIET--YAPELN--RGTR--------------------------------------------------Y-MGIVVIWA 266
WP_006650865_1 DUF63 ---------------------FGLER--YAPELN--RGTR--------------------------------------------------Y-MGIVVIWA 266
WP_066301295_1 DUF63 ---------------------IGIER--YAPELN--RGTR--------------------------------------------------N-MGIVVIWA 266
WP_049966804_1 DUF63 ---------------------FGLER--YAPELN--RGTR--------------------------------------------------Y-MGIVVIWA 266
WP_076145380_1 DUF63 ---------------------VGLKR--YAPELN--RGTR--------------------------------------------------N-MGIVVIWA 266
WP_097378845_1 DUF63 ---------------------VALER--FAPELN--RGTE--------------------------------------------------S-MGIVVIWA 266
WP_008452261_1 DUF63 ---------------------LTLER--VAPELN--RGTR--------------------------------------------------S-MGLVVIWA 266
WP_008452261_1 DUF63 ---------------------LTLER--VAPELN--RGTR--------------------------------------------------S-MGLVVIWA 266
WP_006432643_1 DUF63 ---------------------VALER--FAPELN--RGTE--------------------------------------------------S-MGIVVIWA 266
WP_008452261_1 DUF63 ---------------------LTLER--VAPELN--RGTR--------------------------------------------------S-MGLVVIWA 266
WP_007109662_1 DUF63 ---------------------IALER--VAPELN--RGTR--------------------------------------------------S-MGLVVIWA 266
WP_049952821_1 DUF63 ---------------------LLIQQ--FAPGLN--RGTE--------------------------------------------------Y-MGIVVIWA 266
WP_086889541_1 DUF63 ---------------------VVVDR--FFPSVN--RGTA--------------------------------------------------Y-MGIIVIWA 269
WP_005580009_1 DUF63 ---------------------TAVQR--FVPDVN--RGTR--------------------------------------------------Y-MGLVVIWA 268
WP_049927356_1 DUF63 ---------------------LLIDR--LAPWIN--DGTR--------------------------------------------------Y-MGIVVIWA 266
WP_087714455_1 DUF63 ---------------------YLIVK--LAPGLN--RGTR--------------------------------------------------Y-MGVVVIWA 266
WP_013878438_1 DUF63 ---------------------GAIQR--FAPGIN--RGTE--------------------------------------------------Y-MGIVVIWA 266
WP_049921111_1 DUF63 ---------------------GAIVQ--FAPEIN--RGTR--------------------------------------------------Y-MGIVVIWA 266
WP_007142789_1 DUF63 ---------------------TAIER--FAPEIN--RGTR--------------------------------------------------F-MGAIVIWA 265
WP_006651941_1 DUF63 ---------------------RATHA--FIPGIH--RGTM--------------------------------------------------F-MGAVVIWA 266
WP_004216437_1 DUF63 ---------------------KATHA--FMPGIH--RGTM--------------------------------------------------F-MGAVVIWA 266
WP_071402513_1 DUF63 ---------------------RATHA--FMPGIH--RGTM--------------------------------------------------F-MGAVVIWA 266
WP_006666462_1 DUF63 ---------------------SLIDR--FAPGVN--RGTK--------------------------------------------------F-IGAVVIWA 266
WP_049904535_1 DUF63 ---------------------AVIDR--FAPGIN--RGTK--------------------------------------------------F-IGAVVIWA 266
WP_006824405_1 DUF63 ---------------------AVIDR--FVPGIN--RGTK--------------------------------------------------F-IGAVVIWA 266
WP_011323802_1 DUF63 ---------------------WVTER--FVPGIN--EGTG--------------------------------------------------A-MGALVVWG 267
WP_015409719_1 DUF63 ---------------------WASQR--YAPFIN--EGTG--------------------------------------------------Y-VGAVIVWG 268
WP_006883947_1 DUF63 ---------------------VGADR--YEPEIN--AGTG--------------------------------------------------I-VGLGILWA 267
WP_077207545_1 DUF63 ---------------------LGVDR--FRPEIN--AGTG--------------------------------------------------L-IGLVVLWG 270
ESS12903_1 putative  ---------------------ISTQR--FAPWIN--AGTG--------------------------------------------------F-IGAVIVWG 270
WP_008417361_1 DUF63 ---------------------VGTER--FAPMVN--AATG--------------------------------------------------Y-MGVLVVWG 269
WP_066381608_1 DUF63 ---------------------IGTER--FAPEVN--VATG--------------------------------------------------Y-MGVLVVWG 264
WP_049947642_1 DUF63 ---------------------YATKR--WAAWVH--EGTG--------------------------------------------------I-AGLVVICA 271
ESS05953_1 putative  ---------------------LVLRR--LFPETL--AGTG--------------------------------------------------LTLSGVVLWS 262
WP_096390051_1 DUF63 ---------------------WLIGR--YEPSIN--DGTG--------------------------------------------------Y-IGLVVIWG 276
WP_021072749_1 DUF63 ---------------------WLIER--YEPSVN--DGTG--------------------------------------------------Y-IGLVIIWG 276
WP_049982791_1 DUF63 ---------------------TLATR--YEPSVR--QGTG--------------------------------------------------F-AGILIIWG 274
WP_008585990_1 DUF63 ---------------------ALATR--RIPTIR--QGTG--------------------------------------------------A-AGIVIIWG 276
WP_006628721_1 DUF63 ---------------------ALATR--RIPTIR--QGTG--------------------------------------------------A-AGIVIIWG 274
WP_006113218_1 DUF63 ---------------------AVATR--QIPTIR--QGTG--------------------------------------------------A-AGIVIIWG 275
WP_049930034_1 DUF63 ---------------------ALATR--RIPTIR--QGTG--------------------------------------------------A-AGVVIIWG 276
WP_049906047_1 DUF63 ---------------------AIATR--QIPTLR--QGTG--------------------------------------------------A-AGIVIIWG 276
WP_044965494_1 DUF63 ---------------------AIATR--QISTIR--QGTG--------------------------------------------------A-AGIVIIWG 276
WP_096393195_1 DUF63 ---------------------AIATR--RIPTIR--QGTG--------------------------------------------------A-AGIVIIWG 276
WP_049908400_1 DUF63 ---------------------VIATR--RIPTIR--QGTG--------------------------------------------------A-AGIVIIWG 276
WP_049908585_1 DUF63 ---------------------AIATR--RISTIR--QGTG--------------------------------------------------A-AGIVIIWG 276
WP_049902631_1 DUF63 ---------------------AIATR--RISTIR--QGTG--------------------------------------------------A-AGIVIIWG 276
WP_049983668_1 DUF63 ---------------------ALATR--QIPTIR--QGTG--------------------------------------------------A-AGIVIIWG 275
WP_053772267_1 DUF63 ---------------------ALATR--RIATIR--QGTG--------------------------------------------------A-AGIVIIWG 275
WP_007999151_1 DUF63 ---------------------KLATG--YEPTIR--QGTG--------------------------------------------------V-GGILIIWG 274
WP_015909917_1 DUF63 ---------------------KLATA--YVPDIR--QGTG--------------------------------------------------V-AGIVIIWG 275
WP_004050594_1 DUF63 ---------------------KLATA--YVPEVR--QGTG--------------------------------------------------V-AGIVIIWG 275
WP_095636035_1 DUF63 ---------------------KLATA--SVPEVR--QGTG--------------------------------------------------A-AGIVIIWG 275
WP_008003811_1 DUF63 ---------------------KLATA--YAPEVR--QGTG--------------------------------------------------V-AGIVIIWG 275
WP_066416034_1 DUF63 ---------------------KLIER--FEPSIN--RGTG--------------------------------------------------L-VGLVIIWG 276
ESS03170_1 putative  ---------------------ALIAR--FNPSIN--QGTG--------------------------------------------------I-AGFVIIWG 275
WP_089671383_1 DUF63 ---------------------KLIGT--SYPELN--AGTGF---------------------------------------------------IGFVVIWA 274
ERH07656_1 putative  ---------------------LLIER--AAPAIN--AGTGL---------------------------------------------------VGLVLLWG 274
ESS10096_1 putative  ---------------------LLIER--AAPAIN--AGTGL---------------------------------------------------VGLVLLWG 274
ESS07824_1 putative  ---------------------------------------------------------------------------------------------------- 195
ERH05689_1 putative  ---------------------ALING--VEPSIN--AGTGL---------------------------------------------------MGLVVIWG 274
ERH02266_1 putative  ---------------------ALING--VEPSIN--AGTGL---------------------------------------------------MGLVVIWG 274
ESS07825_1 putative  ---------------------ALING--VEPSIN--AGTGL---------------------------------------------------MGLVVIWG 67
WP_049970017_1 DUF63 ---------------------LAIER--FEPGIN--AGTG--------------------------------------------------A-IGAVIIWG 268
WP_007979289_1 DUF63 ---------------------WGIDR--YAPEIN--AGTG--------------------------------------------------F-IGAVVIWG 268
WP_007977745_1 DUF63 ---------------------WAVNR--YAPEIN--AGTA--------------------------------------------------A-MGAVVIWG 268
WP_014556445_1 DUF63 ---------------------WVIKT--FMPSIN--RGTE--------------------------------------------------F-VGLVILWG 270
WP_049935775_1 DUF63 ---------------------WLIER--YAPAIN--AGTG--------------------------------------------------R-MGFVVIWG 268
WP_008325149_1 DUF63 ---------------------WLIER--FAPDVN--KGTG--------------------------------------------------K-IGLMVLWG 270
WP_007543631_1 DUF63 ---------------------WLIER--FAPDVN--KGTG--------------------------------------------------K-IGLMVLWG 270
WP_049905041_1 DUF63 ---------------------WLVER--FAPEVN--EGTG--------------------------------------------------Y-IGLVVIWG 270
WP_049913563_1 DUF63 ---------------------WLVER--FAPEVN--EGTG--------------------------------------------------Y-IGLVVIWG 270
WP_049967851_1 DUF63 ---------------------WLVER--FAPEVN--EGTG--------------------------------------------------Y-IGLVVIWG 270
WP_049914905_1 DUF63 ---------------------WLVER--FAPEVN--EGTG--------------------------------------------------Y-IGLVVIWG 270
WP_049916430_1 DUF63 ---------------------WLVER--FAPEVN--EGTG--------------------------------------------------Y-IGLVVIWG 270
WP_049896892_1 DUF63 ---------------------WLVER--FAPEVN--EGTG--------------------------------------------------Y-IGLVVIWG 270
WP_049896892_1 DUF63 ---------------------WLVER--FAPEVN--EGTG--------------------------------------------------Y-IGLVVIWG 270
```

```
WP_049896892_1 DUF63 ---------------------WLVER--FAPEVN--EGTG--------------------------------------------------Y-IGLVVIWG 270
WP_058828253_1 DUF63 ---------------------WLVER--FAPEVN--EGTG--------------------------------------------------Y-IGLVVIWG 270
WP_058568959_1 DUF63 ---------------------WLVER--FAPEVN--EGTG--------------------------------------------------Y-IGLVVIWG 270
WP_049917947_1 DUF63 ---------------------WLIER--FAPEIN--EGTG--------------------------------------------------Y-IGLVVIWG 270
WP_049920348_1 DUF63 ---------------------WLVER--FAPEIN--EGTG--------------------------------------------------Y-IGLVVIWG 270
WP_089777545_1 DUF63 ---------------------WLVER--FAPEIN--EGTG--------------------------------------------------Y-IGLVVIWG 270
WP_008320633_1 DUF63 ---------------------WLIER--FKPEIN--AGTG--------------------------------------------------R-IGLVVIWG 270
WP_004060355_1 DUF63 ---------------------WLVER--FAPEVN--EGTG--------------------------------------------------R-IGFVVIWG 270
WP_103426078_1 hypot ---------------------ALVER--FAPHVN--AGTG--------------------------------------------------Y-IGLVVIWG 270
WP_009367433_1 DUF63 ---------------------WLIER--FAPEIN--KGTG--------------------------------------------------T-IGFVIIWG 269
WP_013440552_1 DUF63 ---------------------WLIER--FAPQIN--SGTE--------------------------------------------------T-IGLVIIWG 270
WP_049916626_1 DUF63 ---------------------WLIER--FAPEIN--SGTE--------------------------------------------------T-IGFVILWG 270
WP_058582837_1 DUF63 ---------------------WLVET--YKPELN--AGTV--------------------------------------------------Y-IGLLLIWG 271
WP_101298124_1 hypot ---------------------WLIET--YAPDIN--TGTE--------------------------------------------------Y-IGLVVIWG 271
PIN95153_1 hypotheti ---------------------------------------------------------------------------------------------------- 95
PIU22108_1 hypotheti ---------------------LIFYV--IKFKLN--A-------------------------------------------------------FETLAITA 188
AAR39201_1 NEQ352    ---------------------FGLSLIILCPVFL----------------------------------------------------------IDKKIALA 133
OIR14399_1 hypotheti ---------------------YVYEV--KNPSVI--PDIILSAIGLTVAIFFITYTSPNNYKEYGQ------------------------R-TPLILYFA 274
OIR20963_1 hypotheti ---------------------LIFSHSKWAPAAHMLNP------------------------------------------------------LYLTLYFG 332
OIR22371_1 hypotheti ---------------------FFSWF--IFPLK---------------------------------------------------------S-VYLLLYFG 283
EGQ43935_1 putative  ---------------------KAEKP--GLLNW-----------------------------------------------------------DFALPVSA 182
MAG21679_1 hypotheti ---------------------NKLTK--LKLLRD--------------------------------------------------------K-FNILVTVS 191
PIN85618_1 hypotheti ---------------------ELKEK--KFFSDK----------------------------------------------------------LNILVVAG 190
PIN99249_1 hypotheti ---------------------LITEKLNFNLLSA--------------------------------------------------------K-LNLAVLFG 194
AJF59838_1 hypotheti ---------------------YLIAKKIRPQLVT--------------------------------------------------------DRLNLLAVAG 190
WP_042682145_1 DUF63 ---------------------LVFYK--FKPFEK----------------------------------------------------------LYLYPVLA 175
WP_013468033_1 DUF63 ---------------------LIFYR--FKPFEK----------------------------------------------------------MYLYPVLA 175
WP_055281581_1 DUF63 ---------------------LGYYR--FKPFEK----------------------------------------------------------IYLFPVFA 174
WP_048160382_1 DUF63 ---------------------LVYYK--FRPFEK----------------------------------------------------------LYLLPISA 174
WP_042701216_1 DUF63 ---------------------LAYYK--FRPFEK----------------------------------------------------------LYLFPVFA 174
WP_004068659_1 DUF63 ---------------------LVYYK--FRPFEK----------------------------------------------------------LYLFPVFA 174
WP_058946665_1 DUF63 ---------------------LVYYK--FRPFEK----------------------------------------------------------LYLFSVFA 174
WP_014835499_1 DUF63 ---------------------LAFYK--FKGFDR----------------------------------------------------------LYLYPVLA 174
WP_010884243_1 DUF63 ---------------------LAYYK--FKPFEK----------------------------------------------------------LYLYPTLA 177
WP_013748129_1 DUF63 ---------------------LAYYK--FRPFER----------------------------------------------------------LYLYPVLA 173
WP_010867251_1 DUF63 ---------------------FAFYK--VRPFER----------------------------------------------------------LYLYPVLA 173
WP_014733359_1 DUF63 ---------------------LIFYR--YKPFER----------------------------------------------------------LYLYPVLA 174
WP_068319891_1 DUF63 ---------------------LAFYR--FKPFDR----------------------------------------------------------LYLYPVLA 174
WP_068576049_1 DUF63 ---------------------LAFYR--FKPFDR----------------------------------------------------------LYLYPVLA 174
WP_014013692_1 DUF63 ---------------------LLIYR--YRPFDK----------------------------------------------------------LYLYAVLA 179
WP_014789520_1 DUF63 ---------------------LLIYR--YRPFDR----------------------------------------------------------LYLYAVLA 179
WP_088180676_1 DUF63 ---------------------LLIYK--YRPFDR----------------------------------------------------------LYLYAVLA 179
WP_088864885_1 DUF63 ---------------------LLVYR--YRPFDR----------------------------------------------------------LYLYAVLA 179
WP_088856755_1 DUF63 ---------------------LLLYK--YRPFDR----------------------------------------------------------LYLYATLA 179
WP_088865981_1 DUF63 ---------------------LLVYK--YRPFDR----------------------------------------------------------LYLYAVLA 179
WP_013906334_1 DUF63 ---------------------FTLYR--FRPFEK----------------------------------------------------------LYLYPTLA 175
WP_088862658_1 DUF63 ---------------------LAFYR--WKPFDR----------------------------------------------------------LYLYPTLA 178
WP_088882956_1 DUF63 ---------------------LAFYR--WRPFDR----------------------------------------------------------LYLYPVLA 178
WP_012571938_1 DUF63 ---------------------FAFYR--WKPFER----------------------------------------------------------LYLYPVLA 175
WP_068663933_1 DUF63 ---------------------FAFYR--WRPFER----------------------------------------------------------LYLYPVLA 175
WP_074631153_1 DUF63 ---------------------FAFYR--WRPFER----------------------------------------------------------LYLYPVLA 175
WP_058939074_1 DUF63 ---------------------FAFYR--WRPFDR----------------------------------------------------------LYLYPVLA 179
WP_088854681_1 DUF63 ---------------------LAFYR--WRPFDK----------------------------------------------------------LYLYAVLA 179
WP_088885426_1 DUF63 ---------------------LLYYR--WRPFDK----------------------------------------------------------LYLYAVLA 175
WP_011249694_1 DUF63 ---------------------FIFYR--WRPFDR----------------------------------------------------------LYLYPVLA 177
WP_062386972_1 DUF63 ---------------------LAYYK--WRPFDR----------------------------------------------------------LYLYPVLA 177
WP_010478887_1 DUF63 ---------------------LAYYR--WRPFDK----------------------------------------------------------LYLYAVLA 175
WP_088858605_1 DUF63 ---------------------LAYYK--WRPFDK----------------------------------------------------------LYLYAVLA 175
WP_042690028_1 DUF63 ---------------------LAFYR--WRPFDR----------------------------------------------------------LYLYPVLA 177
WP_048150762_1 DUF63 ---------------------LAFYK--WRPFDK----------------------------------------------------------LYLYAVLA 175
WP_048811076_1 DUF63 ---------------------LAFYK--WRPFDK----------------------------------------------------------LYLYAVLA 175
WP_050003256_1 DUF63 ---------------------LAFYK--WKPFDR----------------------------------------------------------LYLYPALA 177
WP_062372100_1 DUF63 ---------------------LAFYR--WKPFDR----------------------------------------------------------LYLYPVLA 175
WP_048165284_1 DUF63 ---------------------VVYYK--FKPFDR----------------------------------------------------------VFFLATLA 177
WP_048148750_1 DUF63 ---------------------LVFYR--FRPFER----------------------------------------------------------TFLYTVLV 177
OYT53462_1 hypotheti ---------------------FKVSG--INPFNK--------------------------------------------------------P-DNLMLIFA 160
KYC51152_1 hypotheti ---------------------FLSSI--FYKPLK--E-----------------------------------------------------F-ENTAILFG 163
KYC46003_1 hypotheti ---------------------FLSSI--FYKPLK--------------------------------------------------------EFENTAILFG 163
KYC48643_1 hypotheti ---------------------FLSSI--FYKPLK--------------------------------------------------------EFENTAILFG 163
KYC55321_1 hypotheti ----------------------IISPLFYKPIKE---------------------------------------------------------FENIAILFG 163
KYC57927_1 hypotheti ----------------------IISPLFYKPIKE--------------------------------------------------------F-ENIAILFG 163
KYC57171_1 hypotheti ----------------------IISPLFYKPIKE--------------------------------------------------------F-ENIAILFG 163
OIO20701_1 hypotheti ------------------ASKMLFKR--FGIVSS----------------------------------------------------------LAAFVVLA 207
PIT83986_1 hypotheti ---------------------FKLDR----------------------------------------------------------------I-EYVLPVFA 221
OIO24760_1 hypotheti --------------------RFAAPR--FKLDAS--------------------------------------------------------F-TLQLLVFS 188
OIO24701_1 hypotheti ---------------------FKLAAVAFKHRTE--------------------------------------------------------W-MEHAAVFG 190
OIO27021_1 hypotheti ---------------------EFAWK--KRGIKK--------------------------------------------------------DLFERLAIFG 183
OIO26762_1 hypotheti ---------------------WSKRG--FKEGL-----------------------------------------------------------LEKAVIFG 180
PIN95811_1 hypotheti ---------------------WSKRG--FKEGL-----------------------------------------------------------LEKAVIFG 180
PIO01637_1 hypotheti ---------------------WSKRG--FKEGL-----------------------------------------------------------LEKAVIFG 180
PIO02820_1 hypotheti -----------------------KKR--GIKKDL----------------------------------------------------------FERLAIFG 183
PJD01038_1 hypotheti ----------------------------KKRGIK--KDL-----------------------------------------------------FERLAIFG 183
PIZ91366_1 hypotheti ---------------------WSKRG--FKEGL-----------------------------------------------------------LEKAVIFG 180
WP_013100570_1 DUF63 ---------------------YLINK--RFKIFN--------------------------------------------------------SKIDYLTLLG 178
WP_004590770_1 DUF63 ---------------------FYIDRILKLNILQ--------------------------------------------------------NKIDSYAIIG 180
WP_048196979_1 DUF63 ---------------------FVIRF--LDEKLNLNILSS--------------------------------------------------K-IDHYVIIG 180
WP_015791538_1 DUF63 ---------------------KFLDKKLKLNILQ--------------------------------------------------------SKIDSYVLIG 180
WP_048202292_1 DUF63 ---------------------KFLDKNLKLNILQ--------------------------------------------------------SKIDNYVVIG 180
WP_012981280_1 DUF63 ---------------------KFIDKSLKLNILQ--------------------------------------------------------SKIDSYVIVG 180
WP_064496496_1 DUF63 ---------------------KFLDKTLKLNIIQ--------------------------------------------------------SKIDDYVVIG 180
WP_011972741_1 DUF63 -----------------GIISYLLNN--VEYIKNKLKFDK----------------------------------------------------IDKYAILG 185
WP_013798289_1 DUF63 ---------------------VFIIK--KTKIENKIISSK----------------------------------------------------IDKYAIFS 182
WP_013181100_1 DUF63 --------------------IFILTKIKKSKIEK----------------------------------------------------------LDKYTIFS 181
WP_013867628_1 DUF63 ----------------------PIEK--IKILKN--KL-NFD------------------------------------------------T-VDKYAIIS 186
WP_018153391_1 DUF63 ---------------------RILEK--LKNIKKLNIGKI--------------------------------------------------DRIDKYAIFS 187
```

```
WP_011170272_1 DUF63 ---------------------YIVEK--LKKAKI--E----------------------------------------------------R-IDKYAIFS 181
WP_012066151_1 DUF63 --------------------LFIIEKIKKVKLDR---------------------------------------------------------IDKYAIFS 181
ODS43013_1 hypotheti ---------------------FIIMQ--FRGFGF--LYNE---------------------------------------------------KNYLVVLA 189
OIQ06085_1 hypotheti ------------ILDTILIAVFLYSH--FFPNTI-----------------------LDLFVIFLIFSIPLIVAYVIMKMNSKFPDFLKYEKNYIIVLA 245
PIN67069_1 hypotheti ------------ILDTILIAVFLYSH--FFPNTI-----------------------LDLFVIFLIFSIPLIVAYVIMKMNSKFPDFLKYEKNYIIVLA 245
PIV28099_1 hypotheti ------------ILDTILIAVFLYSH--FFPNTI-----------------------LDLFVIFLIFSIPLIVAYVIMKMNSKFPDFLKYEKNYIIVLA 245
PJC13070_1 hypotheti ------------ILDTILIAVFLYSH--FFPNTI-----------------------LDLFVIFLIFSIPLIVAYVIMKMNSKFPDFLKYEKNYIIVLA 245
PIZ29927_1 hypotheti ------------ILDTILIAVFLYSH--FFPNTI-----------------------LDLFVIFLIFSIPLIVAYVIMKMNSKFPDFLKYEKNYIIVLA 245
PKP60688_1 hypotheti ---------------------LNFFK--FSNFLR-------------------------------------------------------YEKNYIIVLA 191
WP_012956954_1 DUF63 ---------------------VLISY--IIPFFK-------------------------------------------------------DRINLSIISA 175
WP_067148430_1 DUF63 ---------------------NRYSN--FYLVLE--KIIN-------------------------------------------------YKINFSIVLA 192
WP_080460538_1 DUF63 ---------------------DLFTL--SSYDGK---------------------------------------------------------INLSIISA 179
WP_010877090_1 DUF63 ---------------------ALSLR--WEFLRG-------------------------------------------------------R-MNLPVIYA 175
WP_013294901_1 DUF63 ---------------------YIIGL--KWELLR-------------------------------------------------------ERMNLYVVYA 175
WP_010877090_1 DUF63 ---------------------ALSLR--WEFLRG-------------------------------------------------------R-MNLPVIYA 175
BAZ99473_1 hypotheti ---------------------ALSLR--WEFLRG-------------------------------------------------------R-MNLPVIYA 175
PKL66404_1 hypotheti ---------------------ALIGR--KWDLLK-------------------------------------------------------EKFNLAVLSA 183
WP_023991171_1 DUF63 ---------------------LIGRK--WSLLKD-------------------------------------------------------K-FNLGVLSA 176
WP_048072898_1 DUF63 ---------------------LIGRK--WSLLKD-------------------------------------------------------K-FNLSILSA 176
WP_100905180_1 hypot --------------------SLLKDK-----------------------------------------------------------------FNLSVLLA 176
WP_100907231_1 hypot --------------------SLLKDK-----------------------------------------------------------------FNLSVLLA 176
WP_100907231_1 hypot --------------------SLLKDK-----------------------------------------------------------------FNLSVLLA 176
WP_048081925_1 DUF63 ---------------------LIGRK--WALLKD-------------------------------------------------------K-FNLGILSA 175
WP_048081925_1 DUF63 ---------------------LIGRK--WALLKD-------------------------------------------------------K-FNLGILSA 175
WP_069585525_1 DUF63 ---------------------LIGRK--WTLLKD-------------------------------------------------------K-FNLGILSA 175
WP_069585525_1 DUF63 ---------------------LIGRK--WTLLKD--K------------------------------------------------------FNLGILSA 175
WP_013643671_1 DUF63 ---------------------LLRNK--WSLLSN-------------------------------------------------------K-INLTVLMA 175
WP_013824571_1 DUF63 ---------------------LLRNK--WWLLKD-------------------------------------------------------K-FNLSVLMA 175
WP_048192110_1 DUF63 ---------------------LLKNK--WSLLKN-------------------------------------------------------K-FNLSVLMA 179
WP_081810122_1 DUF63 ----------------------------ISPAIGMFYLRS-------------------------------------------------R-VYSFVIFS 212
WP_097298965_1 DUF63 --------------------TEIVKK--ISPVIGMTYLRSR--------------------------------------------------MYSFAIFS 187
PIO00292_1 hypotheti ---------------------RKLTG--FLRNW----------------------------------------------------------LDFAALSA 175
WP_011833679_1 DUF63 ---------------------WAVLR--YLFRWEYVNQP----------------------------------------------------LYLVLIGG 187
WP_042698332_1 DUF63 ---------------------A-VLR--YLFRFE--YINE-------------------------------------------------P-LYLVLIAG 187
WP_042698332_1 DUF63 ---------------------AVLRY--LFRFEY--INEP---------------------------------------------------LYLVLIAG 187
WP_011448164_1 DUF63 ---------------------T-LIR--YGFGWH--YMQD-------------------------------------------------R-LYQMLIFG 189
WP_007314214_1 DUF63 ---------------------VVMRR--LLSWEY--VSDP---------------------------------------------------LYITLLFG 194
WP_042705632_1 DUF63 ---------------------WAVMR--YVFKWEFVDEI----------------------------------------------------LYKLLIFG 189
WP_013329717_1 DUF63 ----------------------AFVK--YALKWD--YVDD-------------------------------------------------I-LYRLLIFG 189
WP_004077722_1 DUF63 ---------------------L-VIR--YVFKWD--FASD-------------------------------------------------I-LYKLLIFG 189
WP_048150927_1 DUF63 ----------------------AFVK--YALKWD--YVDD-------------------------------------------------I-LYKLLIFG 189
WP_012107531_1 DUF63 ---------------------WGFMR--YILSWEYVQDP----------------------------------------------------LYIALIFG 195
PKL70081_1 hypotheti ---------------------A-CMR--YVLKWE--YVTD-------------------------------------------------P-LYMTLLFG 189
WP_015285007_1 DUF63 ---------------------A-FMR--YACRWE--YVND-------------------------------------------------P-LYITLLFG 195
PKL64570_1 hypotheti ---------------------WAFMR--YILSWT--YVAD-------------------------------------------------P-LYITLLFG 195
WP_015286404_1 DUF63 ---------------------G-FMR--YVLKWE--YVSD-------------------------------------------------P-LYCVLIFG 189
WP_012617618_1 DUF63 ---------------------A-FLR--YVLKWE--FASD-------------------------------------------------P-IYVLLLLG 189
WP_014867374_1 DUF63 ---------------------ALLV---YGAKWDYASN-I---------------------------------------------------LYKLLIFG 189
CVK33108_1 conserved ---------------------ALLV---YGAKWDYASN-I---------------------------------------------------LYKLLIFG 189
WP_066956331_1 DUF63 ---------------------A-FLV--YVLKWDYASNI----------------------------------------------------LYKLLIFG 189
WP_011844879_1 DUF63 ---------------------WAFLV--YVLKWDYASNI----------------------------------------------------LYKLLIFG 189
WP_048181868_1 DUF63 ---------------------WAFLV--YVLKWDYASNI----------------------------------------------------LYKLLIFG 189
WP_067073014_1 DUF63 ---------------------A-FFV--YVLKWDYASNI----------------------------------------------------LYKLLIFG 189
PKL62929_1 hypotheti ---------------------A-FLV--YVLKWDYASNI----------------------------------------------------LYKLLIFG 189
WP_004039890_1 DUF63 ---------------------A-FLR--YVLGWE--YVSD-------------------------------------------------P-LYLLLIVG 189
WP_067049225_1 DUF63 ----------------------GFLR--YILRWE--YVAD-------------------------------------------------P-LYTLLIAG 189
KYK37884_1 hypotheti ---------------------YYVSR--WFNLGF-------------------------------------------------------ITSNIEIMGA 175
KYK28019_1 hypotheti ---------------------YYVSR--FLNFKF--IT-----------------------------------------------------KNSEIMGG 175
OYT57835_1 hypotheti ---------------------GMIRKI-FHPKLL-------------------------------------------------------SIENSALLMA 179
OYT33350_1 hypotheti ---------------------LVFYC--IVPYCR-------------------------------------------------------NAFSILVFFS 179
WP_012964745_1 DUF63 ---------------------STLSK---------------------------------------------------------------NDESRIAFFS 170
WP_048091573_1 DUF63 ---------------------HMFSD--RLHADT---------------------------------------------------------FSKFVLFG 173
WP_048096399_1 DUF63 ---------------------HLIST--KIEADS---------------------------------------------------------FSKAVVFA 173
WP_012940099_1 DUF63 ---------------------YLPAR--FLGLDK---------------------------------------------------------LSISVFFS 176
WP_010877969_1 DUF63 ---------------------YAIS---LKPMKN-------------------------------------------------------E-LSLLTMFS 189
WP_013682839_1 DUF63 ---------------------LLTSK--FYSPMS-------------------------------------------------------NLLSYMVFFS 180
WP_015591141_1 DUF63 ---------------------FLTRK--FYPKMH-------------------------------------------------------NGLSDLVFFS 179
WP_012035887_1 DUF63 ---------------------V-IAR--HLKLDF--LTNK---------------------------------------------------LNIAILAA 191
WP_014404626_1 DUF63 ---------------------LLVRH--FKIGFL--TM---------------------------------------------------P-MNVAILGA 186
BAI60262_1 conserved ----------------------LIAR--YLKLGF--LTMP---------------------------------------------------LNVAILGA 186
WP_042684156_1 DUF63 ---------------------VGMRP--FTTLLL-------------------------------------------------------SPINAAIVWA 191
OFV68024_1 membrane  ---------------------YLLLR-----TNR---------------------------------------------------------ENMIILTS 140
WP_013720253_1 DUF63 ---------------------L-SRR--AIPSLS--FLDD-------------------------------------------------R-FNLMIVYA 183
WP_014587756_1 DUF63 ---------------------AGSRL--ISSPIS--GG-----------------------------------------------------QSLLILFA 178
ABK14947_1 Protein o ---------------------ILLQILPHSLSMR-------------------------------------------------------R-CALSVILS 184
OKY79140_1 putative  ---------------------LLYYK---KSFIS--DN-----------------------------------------------------IYITILGC 191
WP_086637003_1 DUF63 ---------------------I-IAR--WANYKI--LLDN---------------------------------------------------MYLLIFGS 192
WP_048089097_1 DUF63 ---------------------LISRTLNFTLLTD-------------------------------------------------------K-LNISILFT 189
WP_097298272_1 DUF63 ---------------------FISRQ--LNFTLL--TD---------------------------------------------------R-VNISILFT 190
WP_013897788_1 DUF63 ---------------------WIFCK--SGSVMF--CSG----------------------------------------------------INIALIWS 191
WP_015323982_1 DUF63 ---------------------ITAKKTGFEILTD-------------------------------------------------------R-LNISILGA 191
WP_011499115_1 DUF63 ---------------------LAFNRAGVSFVKD-------------------------------------------------------P-LNMMIIWS 191
WP_013037846_1 DUF63 ---------------------G-LAR--WQGIGL--ITDR---------------------------------------------------LNFTILLA 191
WP_048205640_1 DUF63 ---------------------FMLEKAGWSYVND-------------------------------------------------------R-LNMTIIWA 191
WP_072561629_1 DUF63 ----------------------GLAR--WQGIGL--ITDR---------------------------------------------------LNFTILWA 191
WP_072360082_1 DUF63 ----------------------GLAR--WQGIGL--ITDR---------------------------------------------------LNFTILWV 191
WP_096711793_1 DUF63 ----------------------GLAR--WQGIGL--ITDR---------------------------------------------------LNFTILWV 191
ODV50598_1 hypotheti ----------------------GLAR--WQGIGL--ITDR---------------------------------------------------LNFTILWV 191
WP_013194042_1 DUF63 ---------------------K-ILK--YSGLDI--LSTN---------------------------------------------------LNFAIMWA 191
WP_048178067_1 DUF63 ---------------------RVARY--FKSEIF--TN---------------------------------------------------P-LNLSILMV 193
WP_048127111_1 DUF63 ---------------------LIARR--LKSTIF--TD---------------------------------------------------P-LNLSILLA 193
WP_011023594_1 DUF63 ---------------------LVARH--FKSAIF------------------------------------------------------TNPLNLSILLA 197
WP_048184587_1 DUF63 ---------------------LVARH--FKSSIF------------------------------------------------------TDPLNLSILLA 193
WP_011032542_1 DUF63 ---------------------LVARHLKSAIFTD-------------------------------------------------------P-LNLSILLA 193
WP_011032542_1 DUF63 ---------------------LVARHLKSAIFTD-------------------------------------------------------P-LNLSILLA 193
```

```
WP_048129141_1 DUF63 ---------------------LVARHFKSSIFTD--------------------------------------------------------P-LNLSILLA 193
WP_048129141_1 DUF63 ---------------------LVARH--FKSSIF-------------------------------------------------------TDPLNLSILLA 193
WP_048169891_1 DUF63 ---------------------MVARRLKSSIFTD--------------------------------------------------------P-LNLSILLA 193
WP_048137175_1 DUF63 ---------------------MVARRLKSSIFTD--------------------------------------------------------P-LNLSILLA 193
WP_048137175_1 DUF63 ---------------------MVARRLKSSIFTD--------------------------------------------------------P-LNLSILLA 193
WP_048137175_1 DUF63 ---------------------MVARRLKSSIFTD--------------------------------------------------------P-LNLSILLA 193
WP_048137694_1 DUF63 ---------------------LIALH--FKSSIF-------------------------------------------------------TDPLNLSILLA 193
WP_048168126_1 DUF63 ---------------------LIANH--FKSSIF--TD----------------------------------------------------P-LNLSILMA 193
WP_048118082_1 DUF63 ---------------------LIARH--FKSSIF-------------------------------------------------------TNPLNLSILMA 193
WP_048158120_1 DUF63 ---------------------LIAHH--FKSSIF-------------------------------------------------------TNPLNLSILMA 193
WP_011305329_1 DUF63 ---------------------LIARH--FKSSIF-------------------------------------------------------TNPLNLSILMA 193
WP_054298619_1 DUF63 ---------------------LIAYH--FKSSIF-------------------------------------------------------TDPLNLSILMA 193
ALK05385_1 hypotheti ---------------------LIANH--FKSSIF-------------------------------------------------------TDPLNLSILMA 193
WP_015052897_1 DUF63 ---------------------LVFDRAGSDILKN--------------------------------------------------------K-LNLSILGV 190
WP_023846134_1 DUF63 ---------------------AIFDRIDFDLLKS--------------------------------------------------------N-VNVAILWV 190
OIN88451_1 hypotheti ---------------------LIVLKMVKWPVEN------------------------------------------------------------KIVTAL 169
PIX50278_1 hypotheti ---------------------LIVLKMVKWPVEN------------------------------------------------------------KIVTAL 169
PIW41402_1 hypotheti ---------------------LIVLKMVKWPVEN------------------------------------------------------------KIVTAL 169
PIY35178_1 hypotheti ---------------------LIVLKMVKWPVEN------------------------------------------------------------KIVTAL 169
PJB74886_1 hypotheti ---------------------LIVLKMVKWPVEN------------------------------------------------------------KIVTAL 169
PIZ33651_1 hypotheti ---------------------LIVLKMVKWPVEN------------------------------------------------------------KIVTAL 169
WP_048165029_1 DUF63 ---------------------YALSK--RIRLIE----------------------------------------------------------ENSYIFYA 177
WP_042681046_1 DUF63 ---------------------YLMTK--KIELVR----------------------------------------------------------ENSYLFYA 177
WP_013467319_1 DUF63 ---------------------YLMTK--KIELVR----------------------------------------------------------ENSYLFYA 177
WP_048152160_1 DUF63 ---------------------YFLSR--EFPLIE----------------------------------------------------------RNSYLFYA 177
WP_015849008_1 DUF63 ---------------------YLLSI--KLHLVK----------------------------------------------------------KNSYLFYA 177
WP_004069276_1 DUF63 ---------------------YLLSK--RIRLVR----------------------------------------------------------ENSYLFYA 177
WP_058946638_1 DUF63 ---------------------YLLSK--RIRLVR----------------------------------------------------------ENYYLFYA 177
WP_042701551_1 DUF63 ---------------------YLLST--KIRLVR----------------------------------------------------------ENSYLFYA 177
WP_055282692_1 DUF63 ---------------------YLLSK--KIWLVK----------------------------------------------------------ENSYLFYA 177
WP_013906023_1 DUF63 ---------------------YALSK--KLSLVR----------------------------------------------------------ENSYLFYV 175
WP_011013158_1 DUF63 ---------------------FLLSK--KVSLVR----------------------------------------------------------ENSYLFYA 176
WP_014733166_1 DUF63 ---------------------FLMTK--KLAIVR----------------------------------------------------------ENSYLFYV 176
WP_068322889_1 DUF63 ---------------------YLMTK--KLSIVR----------------------------------------------------------ENSYLFYV 176
WP_068575625_1 DUF63 ---------------------YLMTK--KLSIVR----------------------------------------------------------ENSYLFYV 176
WP_010885960_1 DUF63 ---------------------YLLTK--KLSIVR----------------------------------------------------------DNSYLFYV 176
WP_010867401_1 DUF63 ---------------------YLLSR--RVYLIR----------------------------------------------------------KNSYLFYV 176
WP_013747987_1 DUF63 ---------------------YILSR--KLQIVR----------------------------------------------------------NNSYLFYV 177
WP_068664049_1 DUF63 ---------------------WLLTK--KVKIIA----------------------------------------------------------DNSWLFYA 179
WP_011249739_1 DUF63 ---------------------WALAK--KFKLIV----------------------------------------------------------DNKVLFYT 180
WP_062386854_1 DUF63 ---------------------WALAK--KFKLIA----------------------------------------------------------DNKVLFYT 180
WP_050003781_1 DUF63 ---------------------WALSK--KLELVR----------------------------------------------------------KNRTLFYT 180
WP_042690699_1 DUF63 ---------------------WLLSR--KFELVR----------------------------------------------------------NNKILFYT 180
WP_088885212_1 DUF63 ---------------------WILSR--KFDLIK----------------------------------------------------------NNGLLFYT 180
WP_010478519_1 DUF63 ---------------------WVLSR--KFQLIK----------------------------------------------------------NNGVLFYT 180
WP_088858240_1 DUF63 ---------------------WALSK--KFEIIR----------------------------------------------------------NNRILFYT 180
WP_015859016_1 DUF63 ---------------------WTLSR--KFELIR----------------------------------------------------------NNRLLFYT 180
WP_014121940_1 DUF63 ---------------------WALSR--KFELVR----------------------------------------------------------NNRLLFYT 180
WP_062373911_1 DUF63 ---------------------WLLSK--RFKLVR----------------------------------------------------------NNSVLFYT 180
WP_088882924_1 DUF63 ---------------------WLTSK--FLPVVK----------------------------------------------------------DNSLLFYT 180
WP_088862911_1 DUF63 ---------------------WLLAR--VLPSVR----------------------------------------------------------DNSLLFYT 180
WP_012572006_1 DUF63 ---------------------WVLSK--KVKLVG----------------------------------------------------------DNSLLFYT 180
WP_088854707_1 DUF63 ---------------------WLIAK--KAPLIA----------------------------------------------------------DNKTLFYT 180
WP_014789474_1 DUF63 ---------------------WLLSR--KFALVS----------------------------------------------------------DNSILFYT 179
WP_088180728_1 DUF63 ---------------------WLISK--KLALVR----------------------------------------------------------DNSLLFYT 180
WP_088864937_1 DUF63 ---------------------WLVSK--KLALVR----------------------------------------------------------DNSILFYT 179
WP_055429686_1 DUF63 ---------------------WFVSK--KLALVR----------------------------------------------------------DNSLLFYT 180
WP_088865932_1 DUF63 ---------------------WLVSK--KLALIR----------------------------------------------------------DNSLLFYT 180
WP_014013642_1 DUF63 ---------------------WLLAK--KVKVID----------------------------------------------------------DNSALFYT 180
WP_088856804_1 DUF63 ---------------------WVLTK--KVQLIG----------------------------------------------------------DNKTLFYT 180
WP_058939475_1 DUF63 ---------------------WMLAK--KVPVIA----------------------------------------------------------DNSVLFYT 179
EHR77276_1 conserved ---------------------AGVED--LHQLKL--TGHEAGVLPVGVRLDTWENAGDLTTNHPVQLLSKKGLLAT--------------P-MVLAMVFG 358
OUV40025_1 hypotheti ---------------------GLAIG--FASLIF--------------------------------------------------------S-FHATRGWP 237
MBJ52984_1 hypotheti GIDAQIELEKRGLEPGVLEEKWTVSE--WEKHES--DEKNDIERLMSKAMIA--------------------------------------SPLTLAFTFG 347
PDH23744_1 hypotheti ---------------------S-QGR--SAVAEL--SRQGLVAGILPPGMSDDEYRKSSADDLPEKGIIEPLRSRAIMAQ----------P-LVFLAVAG 340
PDH25468_1 hypotheti GRESSIQLASQGIVAGILPPGMTEEE--YLASES--AEKDLIESLRTKATMA--------------------------------------YPVAFLPVVG 341

WP_048201856_1 hypot FLFNAIVLLL-TLK--F---------------------------HKNEKIRKIWEV-LLAS--I---------------------------LYLMAVSLF 147
WP_048150344_1 hypot --VHIVTTKL-GIQ--W------------------------------------EAI-TLNG------------------------------LFAVTTCIE 168
AOV95360_1 hypotheti HFLDAASTVK-ALE--R--------------------------GLSESRALAQFFI-EFLG--E-----------------------YG--IFVMKALVI 196
EOD42420_1 Uncharact EIYETIFSII-SVY--Y-----F--------------------NLQSEHQLLNILIFQYNN--P-----------------------LL--FYGLKLFLT 209
AOV95164_1 hypotheti QFFGGAASMV-SLS--Q--------------------------GYRQKQLLTQVFT-SVFG--E-----------------------IG--VVILKAGII 239
EGQ40074_1 putative  QFFGGTVSML-AVT--N--------------------------GYPQKQLLAQLST-DILG-------------------------PGG--ILVLKSGLL 237
KYK23067_1 hypotheti HLIDGITSYVSIYD-PL---------------------RMGLPTYIEKHPASNTLM-DI----------------------------WPPLFPLVKFLLI 355
WP_084383883_1 DUF63 VFIEGVLFEFGGLD-------------------------------------------------------------------------------------- 204
WP_049984677_1 DUF63 QLWDAAQNLI-GVT--F-----L--------------------GYSPKMFLTNLIY-QATG--------------------------FPGSTFVLKFLVT 289
WP_103428047_1 hypot QTWDGAQNLI-GVT--F-----L--------------------GYSPKLVVTQFVY-RATG--------------------------FSGSTFVLKLLVT 288
WP_004048647_1 DUF63 QLWDAAQNLI-GVT--F-----L--------------------GYSPKLVVTNLVY-QATG--------------------------FAGSTFVLKLVVT 288
WP_004594600_1 DUF63 QLWDAAQNLI-GVT--F-----L--------------------GYSPKLVVTNLVY-QATG--------------------------FSGSTFVLKLVVT 288
WP_004594600_1 DUF63 QLWDAAQNLI-GVT--F-----L--------------------GYSPKLVVTNLVY-QATG--------------------------FSGSTFVLKLVVT 288
WP_050050397_1 DUF63 QLWDAAQNLI-GVT--F-----L--------------------GYSPKLVVTNLVY-QATG--------------------------FSGSTFVLKLVVT 288
WP_004594600_1 DUF63 QLWDAAQNLI-GVT--F-----L--------------------GYSPKLVVTNLVY-QATG--------------------------FSGSTFVLKLVVT 288
WP_050050397_1 DUF63 QLWDAAQNLI-GVT--F-----L--------------------GYSPKLVVTNLVY-QATG--------------------------FSGSTFVLKLVVT 288
WP_049983690_1 DUF63 QMWDAAQNLI-GVT--F-----L--------------------GYSPKLVVTNLVY-QATG--------------------------FSGSTFVLKLLVT 288
WP_103428162_1 hypot QTWDAAQNLI-GVT--F-----L--------------------GYSPKLVITNFVY-QATG--------------------------FSGSTFVLKLLVT 288
WP_009378268_1 DUF63 QLWDGAQNLI-GVT--F-----L--------------------GYSPKLVVTNLVY-QATG--------------------------FSESTFVLKLLVT 288
WP_015763176_1 DUF63 HAIDGIANVI-TGD--W-----LDAL-----------GIPG--EYFAKHPANRIIV-DVTE--A----I------QPASLSATIGTSWP--FLVVKMLVA 343
WP_018259238_1 DUF63 HAIDGIANVI-TGD--W-----LDAL-----------GIPG--EYFAKHPANRIIV-DVTE--A----L------QPASLSATIGTSWP--FLVVKMLVA 343
WP_004592615_1 DUF63 HAIDGVANVL-LAD--W-----LDAL-----------NVPL--TYYPKHPANEFII-ETTE--A----L------QPAGLSAAIGTSWP--FLIVKLAVA 337
WP_004518092_1 DUF63 HAIDGVANVL-LAD--W-----LDVL-----------NVPL--TYYPKHPANAFII-SATE--S----L------QPASLSAAIGTSWP--FLIVKLAVA 337
WP_014040450_1 DUF63 HAIDGVANVL-LAD--W-----LDAL-----------NVPL--TYYPKHPANAFII-EATE--S----L------QSAGLTAAIGTSWP--FLIVKLAVA 337
WP_014040450_1 DUF63 HAIDGVANVL-LAD--W-----LDAL-----------NVPL--TYYPKHPANAFII-EATE--S----L------QSAGLTAAIGTSWP--FLIVKLAVA 337
WP_008309018_1 DUF63 HAIDGVANVL-LAD--W-----LDAL-----------NVPL--TYYPKHPANAFII-EATE--S----L------QSAGLTAAIGTSWP--FLIVKLAVA 337
WP_014040450_1 DUF63 HAIDGVANVL-LAD--W-----LDAL-----------NVPL--TYYPKHPANAFII-EATE--S----L------QSAGLTAAIGTSWP--FLIVKLAVA 337
WP_014040450_1 DUF63 HAIDGVANVL-LAD--W-----LDAL-----------NVPL--TYYPKHPANAFII-EATE--S----L------QSAGLTAAIGTSWP--FLIVKLAVA 337
```

```
WP_005534235_1 DUF63 HAIDGVANVL-LAD--W-----LDAL-----------NVPL--TYYPKHPANRFII-EATE--A----L------QPAGITAAIGTSWP--FLVVKLAVA 337
WP_004961153_1 DUF63 HAIDGVANVL-LAD--W-----LDAL-----------NVPL--TYYPKHPANEFII-TATE--A----L------QPAGLSAAIGTSWP--FLVVKLAVA 337
WP_004961153_1 DUF63 HAIDGVANVL-LAD--W-----LDAL-----------NVPL--TYYPKHPANEFII-TATE--A----L------QPAGLSAAIGTSWP--FLVVKLAVA 337
WP_101350151_1 hypot HAIDGVANVL-LAD--W-----LDAL-----------NVPL--TYYPKHPANEFII-EATE--A----L------QPAGLSAAIGTSWP--FLIVKLAVA 337
WP_053968273_1 DUF63 HAIDGVANVL-LAD--W-----LDAL-----------NVPL--TYYPKHPANEFII-EATE--A----L------QPASLSAAIGTSWP--FLIVKLAVA 337
WP_058995687_1 DUF63 HAIDGVANVL-LAD--W-----LDAL-----------NVPL--TYYPKHPANEFII-AATE--A----L------QPAGLSAAIGTSWP--FLIVKLAVA 337
WP_015790673_1 DUF63 HAIDGVANVL-ITD--WADALGLPV------------------SYYPKHPANEIIM-NVTE-AV-----------LPAGIFDAIGSAWP--FLVVKLGMA 338
WP_008524115_1 DUF63 HAIDGVANVL-ITD--WADVLALPV------------------SYYPKHPANEIIM-NVTE-AV-----------LPAAVFEAIGSAWP--FLLVKLGVA 338
WP_075936143_1 DUF63 HAIDGVANVI-AAD--W-----TGEL-------GVVDSAGNALQYGAKHPFNDIIV-GVVD--A----I------QPAGLDSVIGLSWG--FLVVKLVAA 342
WP_020446311_1 DUF63 HSIDGFANVI-SLD--W-----SDEF-----------GLAY--DYDPKHVVNSGIQ-DATR--A----I------QPESVSDAIGVTWP--FIPVKVVAA 333
WP_049898450_1 DUF63 QAIDGVANVL-ASD--W-----WDAI-----------GLPF--QYTAKHPANAIIV-GFTE--T----V------FPPSFVAAVGDSWP--FLVVKVALA 335
WP_006077061_1 DUF63 QAVDGVANVL-ASD--W-----WDVI-----------GLPF--QYTAKHPANAIIV-GFTE--T----V------FPPSFVEAVGDSWP--FLVVKVALA 335
WP_049996700_1 DUF63 QAVDGVANVL-ASD--W-----AQAI-----------GLPF--QYSAKHPINAIIV-GATT--A----I------FPPSIIDVIGDSWP--FLLVKLVLA 335
EMA38628_1 hypotheti QGIDGVANVI-ASD--W-----WNAI-----------GLPF--EYTAKHPVNALIV-GFTE--L----V------VPHSVILVIGDSWP--FLVVKIALS 335
WP_049992561_1 DUF63 HAIDGVANVL-ASD--W-----SDVF-----------GLPF--GYVPKHPANATVI-SITD--S----V------LPASVTDVIGTSWP--FLLIKIVVA 336
WP_010903025_1 DUF63 HAIDGVANVV-GLD--W-----GAEL-----------GYPRG-DLISKHPLNAYIV-DATN--A----V------LPQSVTHLIGDTWP--FILLKVAAV 335
WP_009760947_1 DUF63 HAVDGVANVL-GLD--W-----GAEL-----------GYPRG-DLGSKHPLNQYIV-DFTH--S----V------LPESVIHYTGDTWP--FIVVKVLAV 334
WP_059057097_1 DUF63 HAVDGVANVI-GLD--W-----AAEL-----------GLPR--DLVPKHPVNAAVV-DITQ--N----V------LPASVIQYTGDAWP--FLLLKIAAA 333
WP_058983522_1 DUF63 HAVDGVANVI-GLD--W-----AAEL-----------GLPY--DLVAKHPVNAAVV-DITQ--N----V------LPASIIHYTGDAWP--FLLVKIAAA 333
WP_071932813_1 DUF63 HAVDGVANVL-GID--W-----GAEL-----------GLAA--DMVPKHPANRAII-DLTV--A----V------VPEGITAVIGVAWG--FLLVKIVAA 329
WP_050048584_1 DUF63 HAVDGVSNVL-GID--W-----GAEL-----------GLPN--DLVPKHPVNRAVV-DVGE--T----V------IPESIASIIGTAWP--FLLVKIAAA 329
WP_014051341_1 DUF63 HAVDGAANVI-GLD--Y-----MVVL-----------NAGN--NLYPKHPVNKWIV-DTAG--A----A------------------WP--FLVVKMVAA 321
WP_079233299_1 DUF63 HAVDGAANVI-GLD--Y-----LKAL-----------GVPY--NLNPKHPANQFIV-DFFG--A----A------------------WP--FLVVKMAAA 319
WP_053948572_1 DUF63 HAVDGAANVI-GLD--Y-----LMAL-----------GVPY--NLNPKHPANQFIV-DLFG--A----A------------------WP--FLVVKMAAA 319
WP_049980835_1 DUF63 HAVDGAANVI-GLD--Y-----LMAL-----------GVPY--NLTPKHPANRFIV-DFFG--A----A------------------WP--FLVVKMAAA 319
KPN31749_1 hypotheti HAVDGAANVI-GLD--Y-----LMAL-----------GVPY--NLTPKHPANRFIV-DLFG--A----A------------------WP--FLVVKMAAA 269
AGB16040_1 putative  HAVDGTANVI-GLD--W-----ATAF-----------GLPS--NLVPKHPVNAAIQ-RYTG--Q----L------LPESITSVIGDVWP--FLLLKVAAA 329
WP_007696755_1 DUF63 HAIDGVANVI-GLD--W-----AQAF-----------GLPW--DLTPKHPVNAAVQ-RYTG--Q----L------LPESITGVIGDVWP--FLLLKVAAA 329
WP_015322904_1 DUF63 HAVDGFANQL-MLD--W-----SHVW-----------GMT----YTPKHPVNDAIV-TYTS--A----L------LPQS-AEVIGSAWP--FALLKLAAP 327
WP_076581869_1 DUF63 HAVDGFANQL-MLD--W-----SHVW-----------GLA----YAPKHPVNDAIV-THTE--A----VLDWLFFVPSGLTDTIGEAWP--FALLKLAAP 334
WP_005559622_1 DUF63 HAVDGFANQL-MLD--W-----SHVW-----------GMT----YTPKHPLNDAIV-TYTG--A----L------LPQS-AEVIGEAWP--FALLKLAAP 327
WP_006067640_1 DUF63 HAVDGFANQL-MLD--W-----SQVW-----------GLS----YAPKHPVNDAIV-TVTG--S----V------LPASVTEITGAAWP--FALLKLAAP 323
WP_008164554_1 DUF63 HAVDGFANQL-MLD--W-----SHVW-----------GLS----YAPKHPVNDAIV-TYTG--S----I------VPASITSVIGEAWP--FALLKLAAP 328
WP_006088112_1 DUF63 HAVDGFANQL-MLD--W-----SHVW-----------GLG----YSPKHPVNEAIV-TYTG--S----V------VPAGVTEVIGEAWP--FALLKLAAP 328
WP_012943247_1 DUF63 HAVDGVANVI-GLD--W-----ATAF-----------GLEH--NLTPKHPINGAIV-DITG--S----V------LPPNVVDTIGAAWP--FLLVKLVAA 329
WP_008895026_1 DUF63 HAVDGVANVI-GLD--W-----ATAF-----------GLEH--NLSPKHPINGAIV-DITG--S----V------LPPNVVDAIGAAWP--FLLVKLVAA 329
WP_098727043_1 DUF63 HAVDGVANVI-GLD--W-----ATSL-----------GLPA--NLVPKHPINRAIA-NTTA--D----V------LPANIVSITGSAWP--FLLVKIAAA 330
WP_049990861_1 DUF63 HAVDGVANVV-GLD--W-----ATAL-----------GLPA--NLVPKHPINRAID-NTTA--D----V------LPAGVVDIIGSAWP--FLFVKVAAA 328
WP_008013001_1 DUF63 HAVDGVANVV-GLD--W-----ATTL-----------GLPA--NLVPKHPINAAID-STTA--D----V------LPASVVSATGSAWP--FLFVKLAAA 330
WP_006180566_1 DUF63 HAVDGVANVI-GLD--W-----ATTL-----------GLPA--NLVPKHPINRAIA-NTTA--D----V------LPADVVSATGSAWP--FLLVKLAAA 330
WP_006650865_1 DUF63 HAVDGVANVI-GLD--W-----ATTL-----------GLPA--NLVPKHPINRAIA-NTTA--D----V------LPADVVSATGSAWP--FLLVKLAAA 330
WP_066301295_1 DUF63 HAVDGVANVI-GLD--W-----ATTL-----------GLPA--NLVPKHPINRAIA-NTTA--D----V------LPASVVSATGSAWP--FLLVKLAAA 330
WP_049966804_1 DUF63 HAVDGVANVI-GLD--W-----ATTL-----------GLPA--NLVPKHPINRAIA-NTTA--D----V------LPADVVSATGSAWP--FLLVKLAAA 330
WP_076145380_1 DUF63 HAVDGVANVI-GLD--W-----ATTL-----------GLPA--NLVPKHPINRAIA-NTTA--D----V------LPAGVVSATGSAWP--FLLVKLAAA 330
WP_097378845_1 DUF63 HAVDGVANVV-GLD--W-----ATSL-----------GLSA--NLVPKHPINRAIA-NTTA--N----V------LPADVVAVTGSAWP--FLLVKIAAA 330
WP_008452261_1 DUF63 HAVDGVANVV-GLD--W-----ATSL-----------GLPT--NLVPKHPINRAIA-NTTA--D----V------LPANVVAATGSAWP--FLLVKIAAA 330
WP_008452261_1 DUF63 HAVDGVANVV-GLD--W-----ATSL-----------GLPT--NLVPKHPINRAIA-NTTA--D----V------LPANVVAATGSAWP--FLLVKIAAA 330
WP_006432643_1 DUF63 HAVDGVANVI-GLD--W-----ATSL-----------GLPV--NLVPKHPINRAIA-NTTA--D----V------LPADVVAITGSAWP--FLLVKVAAA 330
WP_008452261_1 DUF63 HAVDGVANVV-GLD--W-----ATSL-----------GLPT--NLVPKHPINRAIA-NTTA--D----V------LPANVVAATGSAWP--FLLVKIAAA 330
WP_007109662_1 DUF63 HAVDGVANVV-GLD--W-----ATSL-----------GLPA--NLVPKHPINRAID-NTTA--D----L------LPANIVAVTGSAWP--FLLVKIAAA 330
WP_049952821_1 DUF63 HAVDGVANVV-GLD--W-----ATVF-----------GHSA--NLIPKHPINRWIV-NTTG--S----V------LPADIVAVTGDAWP--FLLVKLAAA 330
WP_086889541_1 DUF63 HAVDGVANVI-GLD--W-----AVAF-----------GHDQ--NLVPKHPVNEWIV-ETTG--S----V------LPADVVAVTGEAWP--FLIVKLIAP 333
WP_005580009_1 DUF63 HAVDGVANVI-GLD--W-----AIAF-----------GHGQ--NLVPKHPVNATIV-DVTG--S----I------LPPEIVDITGEAWP--FLLVKLAAA 332
WP_049927356_1 DUF63 HAVDGVANVI-GLD--W-----ATAF-----------GHAS--NLVPKHPINAAIV-DLTG--S----L------LPQYVTDVTGQAWP--FLLVKLAAA 330
WP_087714455_1 DUF63 HAVDGVANVI-GLD--W-----AVAF-----------GHAN--NLVPKHPVNAAIV-DITG--S----A------LPPEIVDITGAAWP--FLLVKLLAA 330
WP_013878438_1 DUF63 HAVDGVANVI-GLD--W-----ATAF-----------GHSS--NLVPKHPINAAIV-DWTG--S----I------LPANVVAVTGEAWP--FLLVKLAAA 330
WP_049921111_1 DUF63 HAVDGVANVI-GLD--W-----ATAF-----------GHAH--NLRPKHPINEAIV-DATG--S----I------LPENIVAVTGAAWP--FLLVKLAAA 330
WP_007142789_1 DUF63 HAVDGVANVI-GLD--W-----ATAF-----------GHDR--NLAPKHPVNEAIV-DLTA--S----I------LPAEIAEVTGAAWP--FLLVKLAAA 329
WP_006651941_1 DUF63 HAVDGVANVI-GLD--W-----ATVF-----------GHDH--NLLPKHPVNEAIV-ETTR--S----I------LPESVATVTGAAWP--FLFVKLAAA 330
WP_004216437_1 DUF63 HAVDGVANVI-GLD--W-----ATAF-----------GHEH--NLLPKHPVNEAIV-DTTG--S----L------LPADVVAVTGAAWP--FLLVKLAAA 330
WP_071402513_1 DUF63 HAVDGVANVI-GLD--W-----ATVF-----------GHDH--NLLPKHPVNEAIV-ETTR--S----L------LPADVVSVTGAAWP--FLLVKLAAA 330
WP_006666462_1 DUF63 HSIDGVANVV-GLD--W-----ATAF-----------GHDR--NLIPKHPINAAVV-DLTD--S----I------LPESIVAVTGDAWP--FLLVKLAAA 330
WP_049904535_1 DUF63 HSIDGVANVV-GLD--W-----ATAF-----------GHDS--NLIPKHPINAAVV-DLTD--S----I------LPESIVAVTGDAWP--FLLVKLAAA 330
WP_006824405_1 DUF63 HSIDGVANVV-GLD--W-----ATAF-----------GHDR--NLIPKHPINAAVV-DLTD--S----I------LPESIVAVTGDAWP--FLLVKLAAA 330
WP_011323802_1 DUF63 HAVDGVANVL-SLD--W-----GEQL------------VGV--PYGSKHVINEATV-RITG--A----I------QPQWLTDAVGSAWP--FFFIKVIAA 330
WP_015409719_1 DUF63 HSVDGIANVV-SLD--W-----ADAL-----------GIGV--TYGSKHVVNEATV-RITE--S----I------QPAWLSEAIGTAWP--FLGIKIAAA 332
WP_006883947_1 DUF63 HAIDGVANVV-AAD--W-----LPEL------------GHPIDAYGAKHVANRAII-DVTQ--T----I------QPASLSAAIGTSWP--FLFVKLAVA 332
WP_077207545_1 DUF63 HAIDGVANVV-AAD--W-----LGAL-----------GLPAAIRYSPKHPANAFII-SVAE--T----V------LPASLLSTLGSSWP--FLLVKLAVA 336
ESS12903_1 putative  HSIDGVANVL-SLD--W-----AAAI-----------GLPG--RYVPKHVVNRLVI-SLTE--T----V------QPASVSATIGTAWP--FLLVKIAAA 334
WP_008417361_1 DUF63 HTVDGIANVL-SLD--W-----AEEL------------GLP--TYTPKHVVNSAIV-DLTG--A----L------QPPSVSAAIGTAWP--FLLVKIGAA 332
WP_066381608_1 DUF63 HTVDGIANVL-SLD--W-----ASEL------------GLP--SYTPKHVVNSAII-EVTS--A----V------QPPSVSAAIGTAWP--FLLVKIGAA 327
WP_049947642_1 DUF63 HALDGAANVV-GLN--W-----MAALVGDAGLAPAPPGVGG--NLVPKHPVNAAIV-DFSQ--S----V------FPPVLLSTLGDAWP--FLLVKLVAA 346
ESS05953_1 putative  HAVDGAANVV-ALD--W-----MPAL-----------GAGG--NLVPKHPVNRFVV-DVTG--S----L------LPGSVLAVTGDAWP--FLLVKLVAA 326
WP_096390051_1 DUF63 HAIDGVANVI-GLD--W-----MPVL-----------AGSA--NLVPKHVVNAAIV-DWTA--R----L------LPPSVLAVTGDAWP--FLIVKLAAA 340
WP_021072749_1 DUF63 HAIDGVANVI-GLD--W-----MPVL-----------AGSP--NLVPKHVVNAAIV-DWTA--R----L------LPPSVLAVTGDAWP--FLIVKLAAA 340
WP_049982791_1 DUF63 HAVDGVANVI-GLN--W-----MPAL-----------TGTP--NLVPKHVINSLIV-DWTG--R----L------LPDSVVAVTGDAWP--FLLVKLAAA 338
WP_008585990_1 DUF63 HAVDGVANVI-GLD--W-----MPAL-----------TGTA--NLVPKHVVNALIV-DWTG--R----L------LPASVSAVTGDAWP--FLLVKLAAA 340
WP_006628721_1 DUF63 HAIDGVANVI-GLN--W-----MPAL-----------TGTA--NLVPKHVVNALIV-DWTG--R----L------LPGSIVSVTGDAWP--FLLVKLAAA 338
WP_006113218_1 DUF63 HAVDGVANVI-GLD--W-----MPAL-----------TGTA--NLVPKHVVNSLIV-DWTG--R----L------LPASIVSATGDAWP--FLLVKLAAA 339
WP_049930034_1 DUF63 HAVDGVANVI-GLN--W-----MPAL-----------TGTA--NLVPKHVVNALIV-DWTG--R----L------LPDSILAVTGDAWP--FLLVKLAAA 340
WP_049906047_1 DUF63 HAIDGVANVI-GLN--W-----MPAL-----------TDTA--NLVPKHVVNALIV-DWTG--R----L------LPESILSVTGDAWP--FLLVKLAAA 340
WP_044965494_1 DUF63 HAIDGVANVI-GLN--W-----MPAL-----------TGTP--NLVPKHVVNALIV-DWTG--R----L------LPESIVSVTGDAWP--FLLVKLAAA 340
WP_096393195_1 DUF63 HAIDGVANVI-GLN--W-----MPAL-----------TGTP--NLVPKHVVNALIV-DWTG--R----L------LPDSILAVTGDAWP--FLLVKLAAA 340
WP_049908400_1 DUF63 HAIDGVANVI-GLN--W-----MPAL-----------TGTA--NLVPKHVVNALIV-DWTG--R----L------LPDSILAVTGDAWP--FLLVKLAAA 340
WP_049908585_1 DUF63 HAIDGVANVI-GLN--W-----MPAL-----------TGTA--NLVPKHVVNALIV-DWTG--R----L------LPDSILAVTGDAWP--FLLVKLAAA 340
WP_049902631_1 DUF63 HAIDGVANVI-GLN--W-----MPAL-----------TGTA--NLVPKHVVNALIV-DWTG--R----L------LPDSILAVTGDAWP--FLLVKLAAA 340
WP_049983668_1 DUF63 HAIDGVANVI-GLN--W-----MPAL-----------TGTA--NLVPKHVVNALIV-DWTG--R----L------LPASIVSVTGDAWP--FLLVKLAAA 339
WP_053772267_1 DUF63 HAIDGVANVI-GLN--W-----MPAL-----------TGTA--NLVPKHVVNALIV-DWTG--R----L------LPESIVSVTGDAWP--FLLVKLAAA 339
WP_007999151_1 DUF63 HAVDGVANVI-GLD--W-----MPAL-----------TGTP--NLVPKHVVNALIV-DWTG--R----L------LPQSIVSVTGDAWP--FLLVKLAAA 338
WP_015909917_1 DUF63 HAIDGVANVI-GLN--W-----MPAL-----------TGTA--NLVPKHVVNALIV-DWTG--R----L------LPQSIVSVTGDAWP--FLLVKLAAA 339
WP_004050594_1 DUF63 HAVDGVANVI-GLN--W-----MPAL-----------TGTG--NLVPKHVVNELIV-DWTA--R----L------LPQSIVSVTGDAWP--FLLVKLVAA 339
WP_095636035_1 DUF63 HAIDGVANVI-GLN--W-----MPAL-----------TGTA--NLVPKHVVNALIV-DWTA--R----L------LPGSIVSVTGDAWP--FLLVKLAAA 339
WP_008003811_1 DUF63 HAIDGVANVI-GLN--W-----MPAL-----------TGTA--NLVPKHVVNALIV-DWTG--R----L------LPQSIVSVTGDAWP--FLLVKLAAA 339
WP_066416034_1 DUF63 HAIDGVANVI-GLD--W-----MPAL-----------TGTP--NLVPKHVVNELVV-NVTG--R----V------LPESVLAVTGDAWP--FLLVKLAAA 340
ESS03170_1 putative  HAVDGVANVI-GLD--W-----MPAL-----------TGTA--NLVPKHVVNELVV-EWTA--R----L------LPDPILAVTGNAWP--FLLVKLGAA 339
```

```
WP_089671383_1 DUF63 HAVDGMANVV-GLN--W-----MPAL-----------GAGA--NLIPKHPVNAAIV-EYTS-LL-----------LPDSILAVTGDAWP--FLLVKLVAA 338
ERH07656_1 putative  HSVDGVANVV-GLD--W-----MPAL-----------GAGA--NLVPKHPVNAAVV-DITG-AV-----------LPPSVLAITGDTWP--FLLLKLAAA 338
ESS10096_1 putative  HSVDGVANVV-GLD--W-----MPAL-----------GAGA--NLVPKHPVNAAVV-DITG-AV-----------LPPSVLAITGDTWP--FLLLKLAAA 338
ESS07824_1 putative  -----------SSD-------------------------------------------TSTP-S------------------------------------- 203
ERH05689_1 putative  HAVDGVANVI-GLD--W-----MPAL-----------GAGR--NLVPKHPVNAAVV-DITG-SV-----------LPSSVLAVTGDTWP--FLVLKLAAA 338
ERH02266_1 putative  HAVDGVANVI-GLD--W-----MPAL-----------GAGR--NLVPKHPVNAAVV-DITG-SV-----------LPSSVLAVTGDTWP--FLVLKLAAA 338
ESS07825_1 putative  HAVDGVANVI-GLD--W-----MPAL-------------GAGRNLVPKHPVNAAVV-DITGSV------------LPSSVLAVTGDTWP--FLVLKLAAA 131
WP_049970017_1 DUF63 HAVDGVANVA-VLD--W-----ALEL-----------GLAG--EYTAKHPVNQGIV-DITS--S----V------LPSSITSVIGTAWP--FLLVKIVAA 332
WP_007979289_1 DUF63 HSVDGVANVV-VLD--W-----ASAL-----------GLKG--EYGAKHPVNAAII-DVTS--T----V------LPESVAAAIGTVWP--FLLVKIVAA 332
WP_007977745_1 DUF63 HAVDGVANVA-VLD--W-----AGAL-----------GLAG--QYGAKHPVNRAII-DITT--N----V------FPASVTDVIGTAWP--FLFVKIAAA 332
WP_014556445_1 DUF63 HAVDGVANVI-GLD--W-----MRAI-----------GAGP--NLVPKHPVNQFVV-DVTG--S----V------LPTSVLAITGDTWP--FLLIKLLAA 334
WP_049935775_1 DUF63 HAVDGVANVV-GLD--W-----MVAL-----------GAGP--NLVPKHPVNQAVV-DITA--A----T------LPASALAITGDTWP--FLVVKLVAA 332
WP_008325149_1 DUF63 HAVDGVANVV-GLN--W-----MTAI-----------GAGN--NLIPKHPVNQAVV-DITA--S----T------LPDSVLAITGDAWP--FLLVKLVAA 334
WP_007543631_1 DUF63 HAVDGVANVI-GLD--W-----MVAL-----------GAGN--NLIPKHPVNQAVV-DITA--S----T------LPDSVLAITGDAWP--FLLVKLVAA 334
WP_049905041_1 DUF63 HAIDGVANVI-GLD--W-----MVAL-----------GAGN--NLVPKHPVNQAVV-DFTA--S----T------LPESVLAVTGDAWP--FLLVKLVAA 334
WP_049913563_1 DUF63 HAIDGVANVI-GLD--W-----MVAL-----------GAGN--NLVPKHPVNQAVV-DFTA--S----T------LPESVLAVTGDAWP--FLLVKLVAA 334
WP_049967851_1 DUF63 HAIDGVANVI-GLD--W-----MVAL-----------GAGN--NLVPKHPVNQAVV-DFTA--S----T------LPESVLAVTGDAWP--FLLVKLVAA 334
WP_049914905_1 DUF63 HAIDGVANVI-GLD--W-----MTDL-----------GAGH--NLVPKHPVNQAVV-DFTS--S----T------LPESVLAVTGDAWP--FLLVKLVAA 334
WP_049916430_1 DUF63 HAIDGVANVI-GLD--W-----MTDL-----------GAGH--NLVPKHPVNQAVV-DFTS--S----T------LPESVLAVTGDAWP--FLLVKLVAA 334
WP_049896892_1 DUF63 HAIDGVANVI-GLD--W-----MTDL-----------GAGN--NLVPKHPVNQAVV-DFTS--S----T------LPESVLAVTGDAWP--FLLVKLVAA 334
WP_049896892_1 DUF63 HAIDGVANVI-GLD--W-----MTDL-----------GAGN--NLVPKHPVNQAVV-DFTS--S----T------LPESVLAVTGDAWP--FLLVKLVAA 334
WP_049896892_1 DUF63 HAIDGVANVI-GLD--W-----MTDL-----------GAGN--NLVPKHPVNQAVV-DFTS--S----T------LPESVLAVTGDAWP--FLLVKLVAA 334
WP_058828253_1 DUF63 HAVDGVANVV-GLN--W-----MVAL-----------GAGN--NLVPKHPVNQAVV-DFTA--S----T------LPESVLAVTGDAWP--FLLVKLVAA 334
WP_058568959_1 DUF63 HAVDGVANVI-GLD--W-----MVAL-----------GAGN--NLVPKHPVNQAVV-DFTA--S----T------LPESVLAVTGDAWP--FLLVKLVAA 334
WP_049917947_1 DUF63 HAVDGVANVI-GLD--W-----MVAL-----------GAGN--NLVPKHPVNQAVV-DFTS--S----T------LPESVLAVTGDAWP--FLLVKLVAA 334
WP_049920348_1 DUF63 HAVDGVANVI-GLD--W-----MTAL-----------GAGN--NLVPKHPVNQAVV-DFTA--S----T------LPASVLAVTGDAWP--FLLVKLVAA 334
WP_089777545_1 DUF63 HAIDGVANVV-GLN--W-----MVAL-----------GAGN--NLVPKHPVNQAVV-DFTA--S----T------LPESVLAVTGDAWP--FLLVKLVAA 334
WP_008320633_1 DUF63 HAIDGVANVV-GLD--W-----MTAL-----------GAGN--NLIPKHPVNKAVV-DFTV--S----T------LPEPIFAITGGAWP--FLLVKLVAA 334
WP_004060355_1 DUF63 HAIDGVANVV-GLD--W-----MTAL-----------GAGN--NLIPKHPVNKAVV-DLTA--S----T------LPESVLAITGDAWP--FLLVKLVAA 334
WP_103426078_1 hypot HAVDGVANVI-GLD--W-----MTAL-----------GAGP--NLVPKHPVNQFVV-DFTA--S----T------LPPSILAVTGDAWP--FLLVKLVAA 334
WP_009367433_1 DUF63 HAIDGVANVV-GLD--F-----MTAL-----------GAGR--NLVPKHPVNKAIV-DFTG--A----T------LPESVLAITGDAWP--FLLVKLLAA 333
WP_013440552_1 DUF63 HAVDGVANVI-GLD--W-----MTEL-----------GAGP--NLVPKHPVNQFVV-DFTA--S----T------LPAPIHAITGDAWP--FLLVKLVAA 334
WP_049916626_1 DUF63 HAVDGVANVV-GLD--W-----MTAL-----------GAGP--NLVPKHPVNQFVV-DFTA--S----T------LPASVLAVTGDAWP--FLLVKLVAA 334
WP_058582837_1 DUF63 HALDGVANVV-GLD--W-----MPAL-----------NAGP--NLVPKHPVNQGIV-DITG--A----V------LPASVLAVTGDAWP--FLLVKLAAA 335
WP_101298124_1 hypot HAVDGVANVV-GLD--W-----MGAL-----------GAGP--NLVPKHPVNAAIV-DLTG--A----V------LPESVLAVTGDAWP--FLAVKLVAA 335
PIN95153_1 hypotheti -----------------------------------------------KKILKDKLS---LG------------------------------TLIYLLLWL 115
PIU22108_1 hypotheti QLTDGIATFS-ALT--F-----YP-------------------YFREQHVVSNFVI-QSIG--N-----------------------WV--FPLIKLILV 233
AAR39201_1 NEQ352    HSLDIATTFV-GLT--Y-----L--------------------GLKEKHVLS-----SLIG--E----------------------PFF--IGFFKMVVL 174
OIR14399_1 hypotheti HLLDAKATWL-GIQ-EY--------------------------GYAEKHVLPTFLI-EEFG--T----------------------AFV--MIPLKIIVV 319
OIR20963_1 hypotheti HFVDGSATYL-GID-NY--------------------------GYVEKHVLPTWFI-ETFG--T----------------------AIV--MLPLKFLVV 377
OIR22371_1 hypotheti HFVDGGATFL-GID-VY--------------G------------YTEKHVLPDFFI-EYFG--S-------------------A---IV--MLPLKFLVV 328
EGQ43935_1 putative  HLLDATSTFT-ALR--Y--------------------------GADEKHVLGRIAV-DVFG--D-----------------------WG--LFPLKLGVI 225
MAG21679_1 hypotheti QLIGSVPTFI-AVE--F-----F--------------------GYGEQHPISDIIL-KFFP--------------------------FS--FVLIKLGLV 234
PIN85618_1 hypotheti QVLDGSATFV-ATN--I-----Y--------------------TCGEQHPLSDAIL----G-------------------------VNPGLFILVKIALA 233
PIN99249_1 hypotheti QTLDGVATFT-ALS--F-----FP-------------------SYSEQHVLSDFVI-QNFS-------------------------PAA--FILVKVLLS 239
AJF59838_1 hypotheti QALDGSATFI-ATQ--F-----L--------------------NCGEQHPLSEAIL----G-------------------------VFPAAFILVKVALA 233
WP_042682145_1 DUF63 HLFDIGSTVV-AIH--Y-----Y--------------------GYREVHWLENILV-TKFG-------------------------AFI--YYPWILFIL 219
WP_013468033_1 DUF63 HLFDIGSTVV-AIH--Y-----Y--------------------GYREVHWLENILV-TKFG-------------------------AFI--YYPWILFIL 219
WP_055281581_1 DUF63 HIFDIGSTIV-AIH--Y-----Y--------------------GYREVHWLENILV-QKFG-------------------------AFV--YYPWIVFIL 218
WP_048160382_1 DUF63 HIFDIGSTVV-AIH--Y-----Y--------------------GYREVHWLENILV-QKFG-------------------------AFV--YYPWIVFIL 218
WP_042701216_1 DUF63 HIFDIGSTVV-AIH--Y-----Y--------------------GYREVHWIENILV-QKFG-------------------------AFV--YYPWILFIL 218
WP_004068659_1 DUF63 HIFDIGSTVV-AIH--Y-----Y--------------------GYREVHWIENILV-QKFG-------------------------AFV--YYPWIVVIL 218
WP_058946665_1 DUF63 HIFDIGSTVV-AIH--Y-----Y--------------------GYREVHWIENILV-QKFG-------------------------AFV--YYPWIVLIL 218
WP_014835499_1 DUF63 HYYDIASTVV-AIH--F-----Y--------------------NYREVHWVEHHLT-SIFG-------------------------AYV--YYPWITLIL 218
WP_010884243_1 DUF63 HYFDIASTVV-AIH--F-----Y--------------------GYREVHWVEKYLV-GMFG-------------------------AYA--YYPWITIIL 221
WP_013748129_1 DUF63 HYFDIASTVV-AIH--F-----Y--------------------GYREVHWIENHLV-SMFG-------------------------AYV--YYPWITLIL 217
WP_010867251_1 DUF63 HYFDIASTVV-AIH--F-----Y--------------------SYMEVHWVERHLV-NAFG-------------------------AYA--YYPWITVIL 217
WP_014733359_1 DUF63 HYFDIASTVV-AIH--F-----Y--------------------HYREVHWIESHLV-SWLG-------------------------AYA--YYPWITVIL 218
WP_068319891_1 DUF63 HYFDIASTVV-AIH--F-----Y--------------------NYMEVHWIERHLV-SWFG-------------------------AYA--YYPWITIIL 218
WP_068576049_1 DUF63 HYFDIASTVV-AIH--F-----Y--------------------HYMEVHWVERHLV-SWFG-------------------------AYS--YYPWITIIL 218
WP_014013692_1 DUF63 HLYDMGSTVV-AIH--F-----Y--------------------GYREVHWIENILV-QHFG--A-----------------------YF--YYPWITFIL 223
WP_014789520_1 DUF63 HLYDMGSTVV-AIH--F-----Y--------------------GYREVHWIENILV-QHFG--A-----------------------YF--YYPWITFIL 223
WP_088180676_1 DUF63 HLYDMGSTVV-AIH--F-----Y--------------------GYREVHWIENILV-QHFG--A-----------------------YF--YYPWITIIL 223
WP_088864885_1 DUF63 HLYDMGSTVV-AIH--F-----Y--------------------GYREVHWIENILV-QHFG-------------------------AYF--YYPWITFIL 223
WP_088856755_1 DUF63 HYYDIASTVV-AIH--F-----Y--------------------GYREVHWIENILV-QHFG--A-----------------------YF--YYPWITIIL 223
WP_088865981_1 DUF63 HLYDMGSTVV-AIH--F-----Y--------------------GYREVHWIENILV-QHFG-------------------------AYF--YYPWITIIL 223
WP_013906334_1 DUF63 HYFDIASTVV-AID--I-----Y--------------------GYREVHWIENHLV-RALG-------------------------PYA--YYPWITLIL 219
WP_088862658_1 DUF63 HYFDVASTVV-AIH--Y-----Y--------------------GYREVHWIENILV-NHFG--A-----------------------YF--YYPWITVIL 222
WP_088882956_1 DUF63 HYFDIASTVV-AIH--F-----Y--------------------GYREVHWIESMLV-NRFG-------------------------AYF--YYPWITIIL 222
WP_012571938_1 DUF63 HYFDIASTVV-AIH--F-----Y--------------------GYREVHWIENILV-THLG-------------------------AYF--YYPWITFIL 219
WP_068663933_1 DUF63 HYFDIASTVV-AIH--F-----Y--------------------GYREVHWIENILV-THLG-------------------------AYF--YYPWITFIL 219
WP_074631153_1 DUF63 HYFDIASTVV-AIH--F-----Y--------------------GYREVHWIENILV-NYLG-------------------------AYS--YYPWITLIL 219
WP_058939074_1 DUF63 HYFDIASTVV-AIH--F-----Y--------------------GYREVHWIENILV-NHLG-------------------------AYS--YYPWITLIL 223
WP_088854681_1 DUF63 HFYDMGSTVV-AIQ--F-----Y--------------------GYREVHWIENILV-NHFG--A-----------------------YF--YYPWITIIL 223
WP_088885426_1 DUF63 HLFDMGSTVA-GIQ--F-----Y--------------------NYREVHWIEHYLV-QWFG-------------------------PYI--YYPWIALIL 219
WP_011249694_1 DUF63 HYFDVASTVV-AIH--F-----Y--------------------GYREVHWIENHLV-NWFG-------------------------AYI--YYPWITLIL 221
WP_062386972_1 DUF63 HFYDIASTVV-AIH--F-----Y--------------------GYREVHWIENYLV-NWFG-------------------------AYI--YYPWITLIL 221
WP_010478887_1 DUF63 HMYDMGSTVV-GIH--Y-----Y--------------------GYHEVHWIENHLV-QWFG-------------------------AYF--YYPWITLIL 219
WP_088858605_1 DUF63 HMYDMGSTVV-GIH--Y-----Y--------------------GYREVHWIEHHLV-QWFG-------------------------PYF--YYPWITVIL 219
WP_042690028_1 DUF63 HYFDIASTVV-AIH--F-----Y--------------------GYREVHWIESHLV-SWFG-------------------------AYI--YYPWITLIL 221
WP_048150762_1 DUF63 HYYDMGSTVV-GIH--Y-----Y--------------------GYREVHWIEHHLV-QWFG-------------------------AYI--YYPWITIIL 219
WP_048811076_1 DUF63 HYYDMGSTVV-GIH--Y-----Y--------------------GYREVHWIEHHLV-QWFG-------------------------AYI--YYPWITVIL 219
WP_050003256_1 DUF63 HYFDVASTVV-AIH--F-----Y--------------------GYREVHWIEHHLV-QWFG-------------------------AYI--YYPWITLIL 221
WP_062372100_1 DUF63 HYFDMASTVV-GIH--F-----Y--------------------GYREVHWIEHHLV-QWFG-------------------------AYI--YYPWITLIL 219
WP_048165284_1 DUF63 HMYDLASTVV-AIY--Y-----Y--------------------NHYEVHWIEGILS-EHFG-------------------------PFI--LYPWKLLIL 221
WP_048148750_1 DUF63 HTFDLGSTVV-AIH-RY--------------------------NHYEVHWIEGILA-NHFG-------------------------PFI--LYPWKLLIL 221
OYT53462_1 hypotheti HMLDASSTFI-AYD--F-----Y--------------------GFQEEHLLPRFLIQSLGG--S----------------------AFV--MIPAKLVIV 206
KYC51152_1 hypotheti HLLDGCATFI-AID--Y-----Y--------G------------FGEEHLLPLFLI-NLSG--T----------------------ALV--MIPLKLVLI 208
KYC46003_1 hypotheti HLLDGCATFI-AID--Y-----Y--------------------GFGEEHLLPLFLI-NLSG--T----------------------ALV--MIPLKLVLI 208
KYC48643_1 hypotheti HLLDGCATFI-AID--Y-----Y--------------------GFGEEHLLPLFLI-NLSG--T----------------------ALV--MIPLKLVLI 208
KYC55321_1 hypotheti HLLDGCATFI-AID--Y-----YG--------------------FGEEHLLPLFLI-NISG--T----------------------ALI--MIPLKLILI 208
KYC57927_1 hypotheti HLLDGCATFI-AID--Y-----Y--------------------GFGEEHLLPLFLI-NISG--T----------------------ALI--MIPLKLILI 208
KYC57171_1 hypotheti HLLDGCATFI-AID--Y-----Y--------------------GFGEEHLLPLFLI-NISG--T----------------------ALI--MIPLKLILI 208
OIO20701_1 hypotheti HSIDGAATFT-VID--I-----YNKFEPACTLLNK--------CYFEQHVLSNAIG-QIFA---------------------FTGVGFL-LYFIVKAAFS 268
PIT83986_1 hypotheti HALDGAATWV-SHD-IF--------------------GPAHSIAYFEQHVFPSAIG-EGTP-FG----------------------YFA--FFAIKAVFA 273
```

```
OIO24760_1 hypotheti QAFDGGATLT-GIG--A--------------------------GYFEQHVLGGALV-AISP--------------------------WL--FYLIKVAFA 230
OIO24701_1 hypotheti QCFDGAATFV-GIG--I---------------------GTPSAQYFEQHAVGGTLI-EAAG-------------------------PLA--FYALKVVFA 238
OIO27021_1 hypotheti QVLDGSASFI-GIQ--F---------------------GTPAQSYFEQHVVGGAIM-------S----------------------ASPALFLLVKLAFA 230
OIO26762_1 hypotheti QSLDGAASCI-GIA--Y---------------------PPAGTSYFEQHVVGGALM-GAFS-------------------------PFA--FLALKLAFA 228
PIN95811_1 hypotheti QSLDGAASCI-GIA--Y---------------------PPAGTSYFEQHVVGGALM-GAFS--P-----------------------FA--FLALKLAFA 228
PIO01637_1 hypotheti QSLDGAASCI-GIA--Y---------------------PPAGTSYFEQHVVGGALM-GAFS--P-----------------------FA--FLALKLAFA 228
PIO02820_1 hypotheti QVLDGSASFI-GIQ--F---------------------GTPAQSYFEQHVVGGAIM-SASP-------------------------AL---FLLVKLAFA 230
PJD01038_1 hypotheti QVLDGSASFI-GIQ--F---------------------GTPAQSYFEQHVVGGAIM-SASP-AL---------------------------FLLVKLAFA 230
PIZ91366_1 hypotheti QSLDGAASCI-GIA--Y---------------------PPAGTSYFEQHVVGGALM-GAFS--P-----------------------FA--FLALKLAFA 228
WP_013100570_1 DUF63 QLVDGAATTI-GIG--I-----Y--------------------GYLEQHPIPRFLM-EHFT-------------------------PIS--FIVVKFTIT 222
WP_004590770_1 DUF63 QAIDASATAV-GLA--F-----Y--------------------GYWEQHPIPRFFM-EHFG-------------------------AFS--FIPIKILAV 224
WP_048196979_1 DUF63 QLIDASSTTV-GIG--I-----Y--------------------GYWEQHPIPRFLM-EHFG--V-----------------------YA--FIPFKILVI 224
WP_015791538_1 DUF63 QLIDASATTV-GIG--I-----Y--------------------SYWEQHPIPRFLM-ETFG-------------------------VYS--FIPFKLLVV 224
WP_048202292_1 DUF63 QLIDASATTI-GIG--I-----Y--------------------GYWEQHPIPRFLM-ETFG-------------------------VYS--FIPFKLLVV 224
WP_012981280_1 DUF63 QLIDASATTI-GIG--I-----Y--------------------GYWEQHPIPRFLM-ETFG-------------------------VYS--FIPFKLLVI 224
WP_064496496_1 DUF63 QLIDASATTI-GIG--V-----Y--------------------GYWEQHPIPRFLM-ETFG-------------------------VYA--FIPFKLLII 224
WP_011972741_1 DUF63 QLTDASATAV-GIA--S-----H--------------------GYWEQHPIPRFFM-DTFG--A-----------------------YS--MIPLKLALV 229
WP_013798289_1 DUF63 QLVDASATAV-GIG--I-----Y--------------------GYWEQHPIPRFFM-DMFG-------------------------AYV--MIPLKLAVV 226
WP_013181100_1 DUF63 QLVDASATSV-GIA--I-----Y--------------------GYWEQHPVPRFFM-DLMG-------------------------PYV--MIPLKLIVV 225
WP_013867628_1 DUF63 QLTDASATAV-GIG-MY--------------------------GYWEQHPIPRLFM-GIFG--P-----------------------YI--MIPLKLIVV 230
WP_018153391_1 DUF63 QLVDASATSI-GIG--I-----Y--------------------GYWEQHPVPRFFM-DILG--P-----------------------YI--LIPLKLIVV 231
WP_011170272_1 DUF63 QLVDASATSV-GIG-VF--------------------------GYWEQHPVPRFFM-DYFG--P-----------------------YS--IIPLKMLVV 225
WP_012066151_1 DUF63 QLVDAAATSV-GIG--I-----Y--------------------GYWEQHPIPRLLM-DYFG-------------------------PYS--IIPVKLMVA 225
ODS43013_1 hypotheti HLFDASATYV-GID-LL--------------------------GGGEQHVLPRFLI-EHFG--T----------------------ALV--MFPLKLIVV 234
OIQ06085_1 hypotheti HFFDTSTTYV-GIA--F-----F--------------------SYFEKHVLPSAAI-EIFS-------------------------PAI--MFVLKFIV- 288
PIN67069_1 hypotheti HFFDTSTTYV-GIA--F-----F--------------------SYFEKHVLPSAAI-EIFS-------------------------PAI--MFVLKFIV- 288
PIV28099_1 hypotheti HFFDTSTTYV-GIA--F-----F--------------------SYFEKHVLPSAAI-EIFS-------------------------PAI--MFVLKFIV- 288
PJC13070_1 hypotheti HFFDTSTTYV-GIA--F-----F--------------------SYFEKHVLPSAAI-EIFS-------------------------PAI--MFVLKFIV- 288
PIZ29927_1 hypotheti HFFDTSTTYV-GIA--F-----F--------------------SYFEKHVLPSAAI-EIFS-------------------------PAI--MFVLKFIV- 288
PKP60688_1 hypotheti HLFDASTTYI-GIA--F-----F--------------------GYFEKHVLPSAAI-EIFS-------------------------PAI--MFVLKLIV- 234
WP_012956954_1 DUF63 HMFDASTTFV-AVE--F-----F--------------------NYSEQHVLANTLY-QLFD--T----------------------SIT--MFPMKIIVI 220
WP_067148430_1 DUF63 HLFDASTTFV-ALE--Y-----F--------------------NYSEQHVLPNALN-QLFD--T-----------------------YIT-IFPMKIIVI 237
WP_080460538_1 DUF63 HLFDASSTFI-AVD--Y-----Y--------------------GYFEQHVLPGSIY-NLAG--T----------------------AIT--MFPLKIIVI 224
WP_010877090_1 DUF63 HLFDASSTYV-AVD--W-----Y--------G------------YIEQHVLPSALT-GLTG--T----------------------AMV--MFPLKMAVI 220
WP_013294901_1 DUF63 HLFDASSTYV-AVD--L-----Y--------------------GYAEQHVLPSAFT-ALTG--T----------------------ALV--MFPLKIAVI 220
WP_010877090_1 DUF63 HLFDASSTYV-AVD--W-----Y--------G------------YIEQHVLPSALT-GLTG--T-------------A---------MV--MFPLKMAVI 220
BAZ99473_1 hypotheti HLFDASSTYV-AVD--W-----Y--------G------------YIEQHVLPSALT-GLTG--T-------------A---------MV--MFPLKMAVI 220
PKL66404_1 hypotheti HFFDAASTYV-AVD--Y-----Y--------------------GYWEQHVLPSALT-NLWG--T----------------------AFV--MFPLKILVI 228
WP_023991171_1 DUF63 HLLDASSTFI-AVD--Y-----Y--------------------GYGEQHVLPNALT-QLAD--S----------------------AIV--MYPLKIAVI 221
WP_048072898_1 DUF63 HLLDASSTFI-AVD--Y-----Y--------------------GYGEQHVLPNALT-QLAD--S----------------------AII--MYPLKIAVI 221
WP_100905180_1 hypot HLLDASSTFI-AVD--Y-----Y--------------------GYGEQHVLPNALT-QMAD--S----------------------AIV--MFPLKIAVI 221
WP_100907231_1 hypot HLLDASSTFI-AVD--Y-----Y--------------------GYGEQHVLPNALT-QMAD--S----------------------AIV--MFPLKIAVI 221
WP_100907231_1 hypot HLLDASSTFI-AVD--Y-----Y--------------------GYGEQHVLPNALT-QMAD--S----------------------AIV--MFPLKIAVI 221
WP_048081925_1 DUF63 HLFDASATFI-AID--F-----Y--------------------GYGEQHVLPTALM-NITG--T----------------------AAV--MFPLKIGVI 220
WP_048081925_1 DUF63 HLFDASATFI-AID--F-----Y--------------------GYGEQHVLPTALM-NITG--T----------------------AAV--MFPLKIGVI 220
WP_069585525_1 DUF63 HLFDASATFI-AID--F-----Y--------------------GYGEQHVLPTALM-NITG--T----------------------AAV--MFPLKIGVI 220
WP_069585525_1 DUF63 HLFDASATFI-AID--F-----Y--------------------GYGEQHVLPTALM-NITG--T----------------------AAV--MFPLKIGVI 220
WP_013643671_1 DUF63 HLFDASSTYV-AVD--F-----Y--------------------GYSEQHVLPSALT-GLVG--T----------------------AAV--MFPLKIAVI 220
WP_013824571_1 DUF63 HLFDATSTFV-AVD--F-----Y--------------------GYGEQHVLPTFLT-NLTG--T----------------------AFV--MYPLKIVVI 220
WP_048192110_1 DUF63 HLFDASSTYI-AVD--F-----Y--------------------GYSEQHVLPSALT-GLVG--S----------------------AIV--MFPLKIAVI 224
WP_081810122_1 DUF63 FLLDSLTTYIGTVQ-----------------------------GYTNKHPFSSFLA-SIFG--T----------------------GII--LVPLSLILV 256
WP_097298965_1 DUF63 FLLDSFTTYI-GVD--L-----L--------------------GYTNKHPFSSFLT-SIVG--T----------------------GAV--LIPLSLVLV 232
PIO00292_1 hypotheti HFLDATATFI-TLD-VF-----------------------AYLGYWEQHPLTRLIG-GLGG--S-----------------------YV--WFYVAKLYV 222
WP_011833679_1 DUF63 HMLDASATSY-ALA--F-----K--------------------GYIEQHVLGSTLI-DLTG--T----------------------PYV--MFALKIVVL 232
WP_042698332_1 DUF63 HMLDASATSY-ALE--F-----K--------G------------YIEQHVLGSTLI-ELTG--T----------------------PYV--MFALKIVVL 232
WP_042698332_1 DUF63 HMLDASATSY-ALE--F-------------------------KGYIEQHVLGSTLI-ELTG-TP-----------------------YV--MFALKIVVL 232
WP_011448164_1 DUF63 HMLDASATSF-GID--------LHSV-----------------PYIEQHVVGSALI-DLTG--T----------------------AFV--MFPLKLLVL 236
WP_007314214_1 DUF63 QLLDASATSY-GID--------LHPL-----------------HYVEVHVVGSHLI-EWTG--T----------------------AFS--MFALKLLVL 241
WP_042705632_1 DUF63 QLFDASATSF-GLE--------LHEM-----------------KYVEQHVVGGALI-DATG--T----------------------AFS--MYLLKLLVL 236
WP_013329717_1 DUF63 HMLDASATSF-GID--------LHEV-----------------TYVEQHVVGSALI-EATG--T----------------------AFS--MFALKLVVI 236
WP_004077722_1 DUF63 HLLDASATSF-GID--------LHEM-----------------AYVEQHVVGSALI-DATG--T----------------------AFS--MFGLKLVVV 236
WP_048150927_1 DUF63 HMLDASATSF-GID--------LHEV-----------------TYVEQHVVGSALI-EATG--T----------------------AFS--MFALKLVVI 236
WP_012107531_1 DUF63 QMLDASATSY-GLT--F-----HPAV-----------------HYIEQHVVGSNLI-AMTG--T----------------------AFV--MYPLKLVIL 243
PKL70081_1 hypotheti QLLDASATSY-GID--L-----HPSI-----------------KYIEQHVVGSFLI-DWTG--T----------------------AFV--MFPLKLLVL 237
WP_015285007_1 DUF63 QLLDASATSF-GID--L-----HPSL-----------------QYIEQHVVGSHLI-DLTG--T----------------------AFV--MYPLKLIVL 243
PKL64570_1 hypotheti QLLDASATSF-GIE--L-----HPSV-----------------QYVEQHVVGSALI-DATG--T----------------------AFV--MFPLKLVVL 243
WP_015286404_1 DUF63 QLLDASATSY-GID--------LHSM-----------------AYNEAHVVGGALI-AWTG--T----------------------AFS--MFPLKLVIL 236
WP_012617618_1 DUF63 HLLDASSTSY-GIS--L-----HPLH------------------YVEQHVVGSSLI-AATG--T----------------------GFV--MFPLKLAVL 236
WP_014867374_1 DUF63 HMLDASATSY-GID--I---------------------HPIH--YVEQHVVGSNLI-EATG--T----------------------AFS--MFLLKIAVL 236
CVK33108_1 conserved HMLDASATSY-GID--I---------------------HPIH--YVEQHVVGSNLI-EATG--T----------------------AFS--MFLLKIAVL 236
WP_066956331_1 DUF63 HMLDASATSY-GID------------------------LHPI-HYVEQHVVGSNLI-EITG--T----------------------AFS--MFLLKIAVL 236
WP_011844879_1 DUF63 HMLDASATSY-GID--------IHPV-----------------GYVEQHVVGAALI-DATG--T----------------------AFS--MFLLKVAVL 236
WP_048181868_1 DUF63 HMLDASATSY-GID--------IHPV-----------------HYVEQHVVGAALI-DATG--T----------------------AFS--MFLLKVAVL 236
WP_067073014_1 DUF63 HMLDASATSY-GID--I---------------------HPVH--YVEQHVVGAALI-DATG--T----------------------AFS--MFLLKLAVL 236
PKL62929_1 hypotheti HMLDASATSY-GID--I---------------------HPVH--YVEQHVVGSGLI-EATG--T----------------------AFS--MYLLKLAVL 236
WP_004039890_1 DUF63 HLFDASATSF-GID--------LHSL-----------------HYVEQHVVGSALI-EWSG--T----------------------AFS--MFPLKLAVI 236
WP_067049225_1 DUF63 HMLDASATSF-GID--------LHPM-----------------GYVEQHVVGSHLI-EWTG--T----------------------AFS--MFPLKLAVI 236
KYK37884_1 hypotheti HILDASATSF-GLA--M-----F--------------------GYFEQHMVPGLFI-QFFG--T----------------------PFV--MFPLKIAVV 220
KYK28019_1 hypotheti HILDASATSF-GLA--M-----F--------------------GYFEQHMVPGLFI-NLFG-TP-----------------------FV--MFPLKIAVV 220
OYT57835_1 hypotheti HIFDATSTFV-SLR--F-----F--------------------GYAEQHVISNIVI-NALG--P-----------------------AG--MYLLKIPIV 223
OYT33350_1 hypotheti HMFDGIETYI-GTK--Y-----L--------------------GYIEIHVIPRILI-ENLG-------------------------PIS--LPLAKFFVF 223
WP_012964745_1 DUF63 QMLDGSASFI-GIQ--F-----L--------------------GYWELHVLPRFLI-SLTG-------------------------PWI--MIPLKFVIF 214
WP_048091573_1 DUF63 QLLDGASSFL-GIQ--Y-----L--------------------GYWELHVLPRFFI-DLAG--P-----------------------WV--MIPLKLAVI 217
WP_048096399_1 DUF63 QLMDGSTSFL-GIQ--F-----L--------------------GYWELHVVPRYFI-SLFG-------------------------PWI--MVPLKLAVI 217
WP_012940099_1 DUF63 HMLDASATFY-GIT--Y-----L--------------------NYWELHVIPRLLI-NQFG-------------------------AWV--LIPTKFVVF 220
WP_010877969_1 DUF63 HMLDGFETFF-GIS--Y-----L--------------------GYWELHVLPRYLI-DKFG-------------------------PIS--LPAAKFLVF 233
WP_013682839_1 DUF63 HMVDGWATFF-GIQ--F-----L--------------------GYWELHVLPRFLI-STFG-------------------------PWI--MIPAKVIVF 224
WP_015591141_1 DUF63 HMVDGFATFL-GIQ--F-----L--------------------GYWELHVLPRFLI-DTFG-------------------------AWV--MVPSKIIVF 223
WP_012035887_1 DUF63 HMFDASSTFT-AID--L-----V--------G-----------GWYEEHVVPVFFI-DLFG--T----------------------SFV--MYALKLAVF 237
WP_014404626_1 DUF63 HMFDASSTFT-AID--I-----V--------G-----------GFAEKHVVPVFFI-GMFN--T----------------------AFI--MYALKLAVF 232
BAI60262_1 conserved HMFDASSTFT-AIDIVT--------------------------GFAEKHVVPVFFI-DIFG--S----------------------AFV--MYALKLAVF 232
WP_042684156_1 DUF63 HMFDASSTFV-GID--F-----L--------------------GYVEKHVLPTLII-EHVH--T----------------------ALV--MYPLKLGIY 236
OFV68024_1 membrane  HLLDAASTFI-GVD--Y-----L--------------------SYHPKHVIERLLI-EHTG--T----------------------AFS--MIPLKLTIL 185
WP_013720253_1 DUF63 HMLDASSTYL-GVD--W-----F--------------------GYYEKHVVPTFLI-DLAG--T----------------------AAV--MYPLKLIVL 228
WP_014587756_1 DUF63 HMFDAASTYV-GVD--W-----F--------------------GYVEKHVVPSLLI-EAAG--T----------------------ALV--MFPLKLLIL 223
```

```
ABK14947_1 Protein o HMLDASSTYI-GVD--W-----L--------------------GYEEKHVVPTLLI-HTAG--S----------------------AAV--MYPLKLIVL 229
OKY79140_1 putative  HIFDASSTYI-GYQ--L--------------------------GAKEKHVLPSLLI-EWSG-QP-----------------------AI--MFPLKLTVI 235
WP_086637003_1 DUF63 HIFDASSTFV-GIE-RL--------------------------GATEKHVLPAYLI-ELTG--T----------------------AGV--MFPLKILVI 237
WP_048089097_1 DUF63 HLLDASSTFI-GMD--W-----L--------------------GYYEKHVVPTFFI-NLAG--N-----------------FVDSPSLV--MYPLKLLVF 239
WP_097298272_1 DUF63 HLLDASSTFI-GMD--W-----L--------------------NYYEKHVAPTFFI-DIAS--N-----------------YTDHPALV--MFPLKLLVF 240
WP_013897788_1 DUF63 HLLDASSTYI-GID--F-----F--------------------GYQEKHVVPAYLI-EVTN--T----------------------ALV--MYPLKIFTI 236
WP_015323982_1 DUF63 HMLDASSTFI-GVD--M-----L--------------------GYYEKHVLPSYLI-DLTG--T----------------------ALV--MYPLKLAIF 236
WP_011499115_1 DUF63 HLLDASSTFI-GVD--I-----L--------------------GYYEKHVVPAYLI-DLTG--T----------------------ALV--MFPLKLAIF 236
WP_013037846_1 DUF63 HLMDASSTFI-GID--M-----L--------G------------YYEKHVVPAYLI-DLTG--T----------------------ALV--MYPLKLAIF 236
WP_048205640_1 DUF63 HLLDASSTFI-GVD--T-----L--------------------GYYEKHVVPAYLI-DLTG--T----------------------ALV--MYPLKMAIF 236
WP_072561629_1 DUF63 HLMDASSTFI-GID--M-----L--------------------GYYEKHVVPAYLI-DLTG--T----------------------ALV--MYPLKLAIF 236
WP_072360082_1 DUF63 HLMDASSTFI-GID--M-----L--------------------GYYEKHVVPAYLI-DLTG--T----------------------ALV--MYPLKLAIF 236
WP_096711793_1 DUF63 HLMDASSTFI-GID--M-----L--------------------GYYEKHVVPSYLI-DLTG--T----------------------ALV--MYPLKLAIF 236
ODV50598_1 hypotheti HLMDASSTFI-GID--M-----L--------------------GYYEKHVVPSYLI-DLTG--T----------------------ALV--MYPLKLAIF 236
WP_013194042_1 DUF63 HLLDASSTIV-GID-LL--------------------------GYHEKHVVPSYLI-ELTN--T----------------------ATV--MYPLKILIF 236
WP_048178067_1 DUF63 HLMDASSTYI-GVD-KL--------------------------GYFEKHVLPYYLI-ELTG--T----------------------ALV--MYPLKLIIF 238
WP_048127111_1 DUF63 HMMDASSTYI-GID-KL--------------------------GYFEKHVLPSYLI-KLTG--T----------------------ALV--MYPLKLIIF 238
WP_011023594_1 DUF63 HMMDASSTYI-GIE--K-----L--------------------GYFEKHVLPSYLI-ELTG--T----------------------ALV--MYPLKLIIF 242
WP_048184587_1 DUF63 HMMDASSTYI-GIE--K-----L--------------------GYFEKHVLPSYLI-ELTG--T----------------------ALV--MYPLKLIIF 238
WP_011032542_1 DUF63 HMLDASSTYI-GID--K-----L--------------------GYFEKHVLPSYLI-KLTD--T----------------------ALV--MYPLKIIIF 238
WP_011032542_1 DUF63 HMLDASSTYI-GID--K-----L--------------------GYFEKHVLPSYLI-KLTD--T----------------------ALV--MYPLKIIIF 238
WP_048129141_1 DUF63 HMMDASSTYI-GID--K-----L--------------------GYFEKHVLPSYLI-KLTD--T----------------------ALV--MYPLKLIIF 238
WP_048129141_1 DUF63 HMMDASSTYI-GID--K-----L--------------------GYFEKHVLPSYLI-KLTD--T----------------------ALV--MYPLKLIIF 238
WP_048169891_1 DUF63 HMLDASSTYI-GID--K-----L--------------------GYFEKHVLPSYLI-KLTD--T----------------------ALV--MYPLKLIIF 238
WP_048137175_1 DUF63 HMLDASSTYI-GID--K-----L--------------------GYFEKHVLPSYLI-KLTD--T----------------------ALV--MYPLKLIIF 238
WP_048137175_1 DUF63 HMLDASSTYI-GID--K-----L--------------------GYFEKHVLPSYLI-KLTD--T----------------------ALV--MYPLKLIIF 238
WP_048137175_1 DUF63 HMLDASSTYI-GID--K-----L--------------------GYFEKHVLPSYLI-KLTD--T----------------------ALV--MYPLKLIIF 238
WP_048137694_1 DUF63 HMLDASSTYI-GID--K-----L--------------------GYFEKHVLPSYLI-KLTD--T----------------------ALV--MYPLKIVIF 238
WP_048168126_1 DUF63 HLMDASSTYI-GVD-KL--------------------------GYFEKHVVPAYLI-DLTG--T----------------------ALV--MYPLKLIIF 238
WP_048118082_1 DUF63 HLMDASSTYI-GVD--H-----L--------------------GYFEKHVVPTYLI-NLTG--T----------------------ALV--MYPLKLIIF 238
WP_048158120_1 DUF63 HLMDASSTYI-GVD--H-----L--------------------GYFEKHVVPTYLI-NLTG--T----------------------ALV--MYPLKLIIF 238
WP_011305329_1 DUF63 HLMDASSTYI-GID--H-----L--------------------GYFEKHVVPTYLI-NLTG--T----------------------ALV--MYPLKLIIF 238
WP_054298619_1 DUF63 HLMDASSTYI-GVD--K-----L--------------------GYFEKHVVPAYLI-DLTG--T----------------------ALV--MYPLKLIIF 238
ALK05385_1 hypotheti HLMDASSTYI-GVD--K-----L--------------------GYFEKHVVPAYLI-DLTG--T----------------------ALV--MYPLKLIIF 238
WP_015052897_1 DUF63 HLLDASSTFA-GVD--F-----L--------------------GYYEKHVVPAFLI-DLTG--T----------------------ALV--MYPLKLLIF 235
WP_023846134_1 DUF63 HLMDASSTIA-GID--F-----L--------------------GYYEKHVVPSYLI-DLTG--T----------------------AFV--MYPLKLSIF 235
OIN88451_1 hypotheti HLFDATTTFV-SLQ--Y-----F--------------------NYFEQHVLPRTII-ELTG--T----------------------PFS--FVVVKLIVV 214
PIX50278_1 hypotheti HLFDATTTFV-SLQ--Y-----F--------------------NYFEQHVLPRTII-ELTG--T----------------------PFS--FVVVKLIVV 214
PIW41402_1 hypotheti HLFDATTTFV-SLQ--Y-----F--------------------NYFEQHVLPRTII-ELTG--T----------------------PFS--FVVVKLIVV 214
PIY35178_1 hypotheti HLFDATTTFV-SLQ--Y-----F--------------------NYFEQHVLPRTII-ELTG--T----------------------PFS--FVVVKLIVV 214
PJB74886_1 hypotheti HLFDATTTFV-SLQ--Y-----F--------------------NYFEQHVLPRTII-ELTG--T----------------------PFS--FVVVKLIVV 214
PIZ33651_1 hypotheti HLFDATTTFV-SLQ--Y-----F--------------------NYFEQHVLPRTII-ELTG--T----------------------PFS--FVVVKLIVV 214
WP_048165029_1 DUF63 HFYDATTTFV-GVD--F-----M--------------------GYWEQHVLPRYLI-DLTG--T----------------------AAV--MYLLKFTVL 222
WP_042681046_1 DUF63 HFYDATTTFV-GIQ--F-----M--------------------GYWEQHVLPRFLI-NLTG--T----------------------AAV--MYLLKFVTL 222
WP_013467319_1 DUF63 HFYDATTTFV-GIQ--F-----M--------------------GYWEQHVLPRFLM-NLTG--T----------------------AAV--MYLLKFVTL 222
WP_048152160_1 DUF63 HFYDATTTFV-GVD--F-----L--------------------GYWEQHVLPRYLI-DLTG--T----------------------AAV--MYLLKFTVL 222
WP_015849008_1 DUF63 HFYDATTTFV-GVD--F-----M--------------------GYWEQHVLPRYLM-DITG--T----------------------AAV--MYFLKFAVL 222
WP_004069276_1 DUF63 HFYDATTTFV-GVD--F-----L--------------------GYWEQHVLPRYLM-SLTG--T----------------------AAV--MYLLKFSVL 222
WP_058946638_1 DUF63 HFYDATTTFV-GVD--F-----L--------------------GYWEQHVLPRYLM-GLTG--T----------------------AAV--MYLLKFSVL 222
WP_042701551_1 DUF63 HFYDATTTFV-GVD--F-----L--------------------GYWEQHVLPRYLI-DLTG--T----------------------AAV--MYLLKFSVL 222
WP_055282692_1 DUF63 HFYDATTTFV-GVD--F-----L--------------------GYWEQHVVPRYLM-ELTG--T----------------------AAV--MYLLKFAVL 222
WP_013906023_1 DUF63 HFYDATTTFV-GVD--F-----M--------------------NYWEQHVLPRFLM-ALTG--T----------------------AAV--MYPLKFTVL 220
WP_011013158_1 DUF63 HFYDATTTFV-GVD--F-----M--------------------GYWEQHVLPRLLI-GLTG--T----------------------AAV--MYVLKFIIL 221
WP_014733166_1 DUF63 HFYDATTTFV-GVD--F-----M--------------------GYWEQHVLPRLLI-GLTG--T----------------------AGV--MYLLKFIVL 221
WP_068322889_1 DUF63 HFYDATTTFV-GVD--F-----M--------------------GYWEQHVLPRLLI-GLTG--T----------------------AAV--MYVLKFTVL 221
WP_068575625_1 DUF63 HFYDATTTFV-GVD--F-----M--------------------GYWEQHVLPRLLI-GLTG--T----------------------AAV--MYLLKFTVL 221
WP_010885960_1 DUF63 HFYDATTTFV-GVD--F-----M--------------------GYWEQHVVPRFLI-GLTG--T----------------------AAV--MYVLKFVVL 221
WP_010867401_1 DUF63 HFYDATTTFV-GVD--F-----M--------------------GYWEQHVVPRLLI-GLTG--T----------------------AAV--MYVLKFSIL 221
WP_013747987_1 DUF63 HFYDATTTFV-GVD--F-----M--------------------GYWEQHVVPGLLI-GLTG--T----------------------AAV--MYVLKFVVL 222
WP_068664049_1 DUF63 HFYDATTTFV-GVD--F-----M--------------------GYWEQHVLSRFFI-EHTG--T----------------------AAT--MYILKFVML 224
WP_011249739_1 DUF63 HFYDATTTFV-GIQ--F-----F--------------------GFWEQHVLARTLI-DLTG--T----------------------PAV--MYLEKLLVL 225
WP_062386854_1 DUF63 HFYDATTTFV-GIQ--F-----F--------------------GFWEQHVLARTLI-DLTG--T----------------------PAV--MYLEKLLVL 225
WP_050003781_1 DUF63 HFYDATTTFV-GIQ--F-----F--------------------GFWEQHVLARTLM-NLLG--T----------------------PAV--MYLEKLAVI 225
WP_042690699_1 DUF63 HFYDATTTFV-GIQ--F-----F--------------------GFWEQHVLARTLM-NLFG--T----------------------PAV--MYLEKLVII 225
WP_088885212_1 DUF63 HFYDATTTFV-GIQ--F-----F--------------------GYWEQHVLARTLI-DLTG--T----------------------PAV--IYLEKLVIL 225
WP_010478519_1 DUF63 HFYDATTTFV-GIQ--F-----F--------------------GYWEQHVLARTLM-DLTG--T----------------------PAV--IYLEKLLVL 225
WP_088858240_1 DUF63 HFYDATTTFV-GIQ--F-----F--------------------GYWEQHVLARTLI-DMTG--T----------------------PAV--MYLEKFVIL 225
WP_015859016_1 DUF63 HFYDATTTFV-GIQ--F-----F--------------------GYWEQHVLARTLI-DLTG--T----------------------AAV--MYLEKLIIL 225
WP_014121940_1 DUF63 HFYDATTTFV-GIQ--F-----F--------------------GYWEQHVLARTLI-NLTG--T----------------------AAV--MYLEKLLIL 225
WP_062373911_1 DUF63 HFYDATTTFV-GIQ--F-----F--------------------GYWEQHVLARTLI-NLTG--T----------------------AAV--MYLIKLAVL 225
WP_088882924_1 DUF63 HFYDATTTFV-GLQ--F-----F--------------------GFWEQHVLARWLI-NLSG--T----------------------PAV--MYVEKFLVL 225
WP_088862911_1 DUF63 HFYDATTTFV-GIQ--F-----F--------------------GFWEQHVLARWLM-STFG--T----------------------PAV--MYAEKFFIL 225
WP_012572006_1 DUF63 HFYDATTTFV-GIQ--F-----F--------------------GFWEQHVLARKLM-EFSG--T----------------------PAV--IYPEKLLIL 225
WP_088854707_1 DUF63 HFYDATTTFV-GIQ--F-----F--------------------GFWEQHVLARWLM-DTLG--T----------------------PAV--MYLEKFFIL 225
WP_014789474_1 DUF63 HFYDATTTFV-GIQ--F-----L--------------------GFWEQHVLARWLI-DAFG--T----------------------AAV--MYLEKFLIL 224
WP_088180728_1 DUF63 HFYDATTTFV-GIQ--F-----L--------------------GFWEQHVLARWLM-DTFG--T----------------------PAV--IYAEKFLIL 225
WP_088864937_1 DUF63 HFYDATTTFV-GIQ--F-----L--------------------GFWEQHVLARWLM-DTFG--T----------------------PAV--IYAEKFLIL 224
WP_055429686_1 DUF63 HFYDATTTFV-GIQ--F-----L--------------------GFWEQHVLARWLM-DAFG--T----------------------PAV--IYPEKFLIL 225
WP_088865932_1 DUF63 HFYDATTTFV-GIQ--F-----L--------------------GFWEQHVLARWLM-DTFG--T----------------------PAV--IYAEKFLIL 225
WP_014013642_1 DUF63 HFYDATTTFV-GIQ--F-----L--------------------GYWEQHVLARWLI-DAFG--T----------------------AAV--MYLEKFLIL 225
WP_088856804_1 DUF63 HFYDATTTFV-GIQ--F-----L--------------------GYWEQHVLARWLI-DTFG--T----------------------AAV--MYLEKFLIL 225
WP_058939475_1 DUF63 HFYDATTTFI-GIQ--F-----F--------------------GYWEQHVLARWLI-DTFG--T----------------------AAI--MYLEKFLIL 224
EHR77276_1 conserved QLCDGFATMV-GID-SF--------------------------GYGEKHPVSDAVI-QFGG-RL-----------NDSFGIEYGEGAWL--FTLVKVGLI 415
OUV40025_1 hypotheti TITRGLLSFAVGAC--------FVGL-----------------GHWAQLAATPWL--QESG-------------------------RLPNEVVFWPSLIV 285
MBJ52984_1 hypotheti QLCDGLATWI-GID--F-------------------------GNYTEKHILGTKIM-EIGS--T----------------IIEGDGAWL--FLIVKIILT 398
PDH23744_1 hypotheti QIMDGLATWI-GID-GF-----P--------------------GLGEKHVASQRVI-DAGL--WVNEGLGIEHGMLDEGV-------WL--FALVKFALA 401
PDH25468_1 hypotheti QFLDGIATYI-GID--F-----F--------------------NYSEKHVLSAAVI-ELFD--T----------------------AAT--FAVLKLAIG 386

WP_048201856_1 hypot ILILLVFDIWDF---------------------------LAVFLVFHFF-FCFANTLRQ--------------------------FYVYVLSSN-- 187
WP_048150344_1 hypot FHSFS----------GESCS-------------------LAWIVALLYSSLALIILKRH----------------------------SQHNGN--- 204
AOV95360_1 hypotheti VPVALYIEREVE------GD-------------------ISREVLFMIGVYGLVLGLRN--------------------------YFLLLL----- 236
EOD42420_1 Uncharact LFIALYTKYYLK------DD-------------------LRNVIYLMLFYLGFLPGLRT--------------------------SLIDILI---- 250
AOV95164_1 hypotheti GLAVYMLEKDIE-----EER-------------------MKALALIVLYSIGLATGLRV--------------------------LLRLSMGI--- 282
EGQ40074_1 putative  GAGLFAARD------TEDRF-------------------TEVALLLALVVVGLATGLRV--------------------------YLRVLSGL--- 279
KYK23067_1 hypotheti ILVIFVFDVFYKKELERYPR-------------------LINLLKIGIFILGVSPGLRD--------------------------LLRVTMGV--- 403
```

```
WP_084383883_1 DUF63 ---------------------------------------LTLVHIIESVAIGLG-------------------------------FVVLLISLRR- 229
WP_049984677_1 DUF63 GGIVWYLADAKP---EMNRT-------------------WWWLMTFFIGAIGLPMGVRG--------------------------SLRMMLGV--- 334
WP_103428047_1 hypot GGIVWYLADAKP---EMNRT-------------------WWWLMTFFIGAIGLPMGVRG--------------------------SLRMMLGV--- 333
WP_004048647_1 DUF63 VGIVWYLADAKE---EMNHT-------------------WWWLMTFFIGAIGLPMGVRG--------------------------SLRMLLGA--- 333
WP_004594600_1 DUF63 VGIVWYLADAKE---EMNHT-------------------WWWLMTFFIGAIGLPMGVRG--------------------------SLRMLLGA--- 333
WP_004594600_1 DUF63 VGIVWYLADAKE---EMNHT-------------------WWWLMTFFIGAIGLPMGVRG--------------------------SLRMLLGA--- 333
WP_050050397_1 DUF63 VGIVWYLADAKE---EMNHT-------------------WWWLMTFFIGAIGLPMGVRG--------------------------SLRMLLGA--- 333
WP_004594600_1 DUF63 VGIVWYLADAKE---EMNHT-------------------WWWLMTFFIGAIGLPMGVRG--------------------------SLRMLLGA--- 333
WP_050050397_1 DUF63 VGIVWYLADAKE---EMNHT-------------------WWWLMTFFIGAIGLPMGVRG--------------------------SLRMLLGA--- 333
WP_049983690_1 DUF63 GGIVWYLADAKE---EMNHT-------------------WWWLMTFFIGAIGLPMGVRG--------------------------SLRMMLGV--- 333
WP_103428162_1 hypot GGIVWYLADAKD---EMSHT-------------------WWWLMTFFIGAIGLPMGVRG--------------------------SLRMMLGV--- 333
WP_009378268_1 DUF63 GGIVWYLADAKA---EMNHT-------------------WWWLMTFFVGAIGLPMGVRG--------------------------SLRMMLGV--- 333
WP_015763176_1 DUF63 SLVLWLFTEEFV---EESQR-------------------YALLLLVAVTAVGLGPGTRD--------------------------MLRVTFGI--- 388
WP_018259238_1 DUF63 SLVLWLFTEEFV---EESQR-------------------YALLLLVAVTAVGLGPGTRD--------------------------MLRVTFGI--- 388
WP_004592615_1 DUF63 SIVVSLFNQEFI---DESPR-------------------YALLLLIAVTAVGLGPGTRD--------------------------MLRATFGI--- 382
WP_004518092_1 DUF63 SLVVSLFNQEFI---DDSPR-------------------YALLLLIAVTAVGLGPGTRD--------------------------MLRATFGI--- 382
WP_014040450_1 DUF63 SLVVSLFNQEFI---DDSPR-------------------YALLLLIAVTAVGLGPGTRD--------------------------MLRATFGI--- 382
WP_014040450_1 DUF63 SLVVSLFNQEFI---DDSPR-------------------YALLLLIAVTAVGLGPGTRD--------------------------MLRATFGI--- 382
WP_008309018_1 DUF63 SLVVSLFNQEFI---DDSPR-------------------YALLLLIAVTAVGLGPGTRD--------------------------MLRATFGI--- 382
WP_014040450_1 DUF63 SLVVSLFNQEFI---DDSPR-------------------YALLLLIAVTAVGLGPGTRD--------------------------MLRATFGI--- 382
WP_014040450_1 DUF63 SLVVSLFNQEFI---DDSPR-------------------YALLLLIAVTAVGLGPGTRD--------------------------MLRATFGI--- 382
WP_005534235_1 DUF63 SLVVSLFNQEFI---DDSPR-------------------YALLLLIAVTAVGLGPGTRD--------------------------MLRATFGI--- 382
WP_004961153_1 DUF63 SLVVSLFNQEFI---DDSPR-------------------YALLLLIAVTAVGLGPGSRD--------------------------MLRATFGI--- 382
WP_004961153_1 DUF63 SLVVSLFNQEFI---DDSPR-------------------YALLLLIAVTAVGLGPGSRD--------------------------MLRATFGI--- 382
WP_101350151_1 hypot SLVVSLFNQEFI---DDSPR-------------------YALLLLIAVTAVGLGPGTRD--------------------------MLRATFGI--- 382
WP_053968273_1 DUF63 SLVVSLFNQEFI---DDSPR-------------------YALLLLIAVTAVGLGPGTRD--------------------------MLRATFGI--- 382
WP_058995687_1 DUF63 SLVVSLFNQEFI---DDSPR-------------------YALLLLIAVTAVGLGPGSRD--------------------------MLRATFGI--- 382
WP_015790673_1 DUF63 LGIVWLFDEAFI---EENPR-------------------YSYVLLVGVVAVGLGPGSRD--------------------------MLRAAFGI--- 383
WP_008524115_1 DUF63 VAIVWLFNEAFI---EENPR-------------------YSYVLLVGVTAVGLGPGTRD--------------------------MLRAAFGI--- 383
WP_075936143_1 DUF63 TAVVWVFDDTIF---DDEPR-------------------YAILLLVAILAVGLGPGTRD--------------------------MLRVTFGI--- 387
WP_020446311_1 DUF63 VLVLWLFNDEIL---EESPR-------------------YAVLMLLAVLAVGLGPGTRD--------------------------VLRATFGV--- 378
WP_049898450_1 DUF63 VGILWLFDDRII---EESPR-------------------YSLLLLLAVIVVGLGPGTRD--------------------------MLRATFGI--- 380
WP_006077061_1 DUF63 VGILWLFDDRII---EESPR-------------------YSLLLLLAVIVVGLGPGTRD--------------------------MLRATFGI--- 380
WP_049996700_1 DUF63 VGVIWLFDEQIF---EESPR-------------------YSYLLMLAVLVVGLGPGTRD--------------------------MLRATFGI--- 380
EMA38628_1 hypotheti VGVIWLFEEAIF---EDSPR-------------------YAYLLMLAIIVVGLGPGTRD--------------------------MLRATFGI--- 380
WP_049992561_1 DUF63 VAIVWLFTDDFM---EESPR-------------------YAIVMLMAIAAVGLGPGTRD--------------------------MIRATFGI--- 381
WP_010903025_1 DUF63 LFVLSLFNEELR---ADAPR-------------------YTTLMLVAVLAVGLGPGTRD--------------------------MIRATFGI--- 380
WP_009760947_1 DUF63 LLVLSLFDEELR---EEAPR-------------------YFTLMLVAVLAVGLGPGTRD--------------------------MLRATFGI--- 379
WP_059057097_1 DUF63 LFVLWLFNAELR---EEAPR-------------------YFTLMLVAVVAVGLGPGSRD--------------------------MLRATFGI--- 378
WP_058983522_1 DUF63 LFVLWLFDDQLR---AEAPR-------------------YFTLMLVAVIAVGLGPGSRD--------------------------MLRATFGI--- 378
WP_071932813_1 DUF63 TIVTWLFDEQMI---TEGPR-------------------FSTLLLIAVVAVGLGPGVRD--------------------------MLRATFGV--- 374
WP_050048584_1 DUF63 LIIIALFDEEMY---DEGPR-------------------FTTLLLIAIIAVGLGPGTRD--------------------------MLRATFGV--- 374
WP_014051341_1 DUF63 AFVLWIFEPELY---DESPR-------------------YTTLLLVAVASVGLGPGTRD--------------------------LFRSMFGV--- 366
WP_079233299_1 DUF63 TFLIWVFEPDLV---EETPR-------------------YTTLLLIAVSAVGLGPGSRD--------------------------LLRSVFGV--- 364
WP_053948572_1 DUF63 TFLIWVFEPELV---EETPR-------------------YTTLLLIAVSAVGLGPGSRD--------------------------LLRSVFGV--- 364
WP_049980835_1 DUF63 TFLIWVFEPELV---EETPR-------------------YTTLLLIAVSAVGLGPGSRD--------------------------LLRSVFGV--- 364
KPN31749_1 hypotheti TFLIWVFEPELV---EETPR-------------------YTTLLLIAVSAVGLGPGTRD--------------------------LLRSVFGV--- 314
AGB16040_1 putative  VFIIWVFNEEVF---EESPR-------------------FTIMLMLTVVAVGLGPGTRD--------------------------MLRATFGV--- 374
WP_007696755_1 DUF63 VFIIWVFNEEVF---EESPR-------------------FTIMLMLTVVAVGLGPGTRD--------------------------MLRATFGV--- 374
WP_015322904_1 DUF63 VAIIWLFNEEIF---EESPQ-------------------FAYLLLVAVVAVGLGPGTRD--------------------------MLRATFGV--- 372
WP_076581869_1 DUF63 VFIIWIFNEEIF---EESPR-------------------FAILMMITVVAVGLGPGTRD--------------------------MLRATFAI--- 379
WP_005559622_1 DUF63 VFIIWLFNEEIF---EESPQ-------------------FAYLLLVAVVAVGLGPGSRD--------------------------MLRATFGV--- 372
WP_006067640_1 DUF63 VFIIWIFNEEIF---DENPR-------------------FAILMMITVVAVGLGPGTRD--------------------------MLRATFGI--- 368
WP_008164554_1 DUF63 VFIIWIFNEEIF---DENPR-------------------FAILMMITVVAVGLGPGTRD--------------------------MLRATFGI--- 373
WP_006088112_1 DUF63 VFIIWIFNEEIF---EESPR-------------------FAILMMITVVAVGLGPGTRD--------------------------MLRATFGV--- 373
WP_012943247_1 DUF63 TFVVWVFNEEIF---DEQPR-------------------FAILMLITVVAVGLGPGTRD--------------------------MLRATFGV--- 374
WP_008895026_1 DUF63 TFVVWVFNEEIF---DEQPR-------------------FAILMLITVVAVGLGPGTRD--------------------------MLRATFGV--- 374
WP_098727043_1 DUF63 VFVIWIFDETVF---EDSPR-------------------YAVLLMITVVAVGLGPGTRD--------------------------MLRATFGV--- 375
WP_049990861_1 DUF63 VFVIWIFDETVF---EDSPR-------------------YAILLMITVVAVGLGPGTRD--------------------------MLRATFGV--- 373
WP_008013001_1 DUF63 VFVIWIFDETVF---EDNPR-------------------YAILLMITVVAVGLGPGTRD--------------------------MLRATFGV--- 375
WP_006180566_1 DUF63 VFVIWIFDETVF---EDSPR-------------------YAVLLMITVVAVGLGPGTRD--------------------------MLRATFGV--- 375
WP_006650865_1 DUF63 VFVIWIFDETVF---EDSPR-------------------YAVLLMITVVAVGLGPGTRD--------------------------MLRATFGV--- 375
WP_066301295_1 DUF63 VFVIWIFDETVF---EDSPR-------------------YAVLLMITVVAVGLGPGTRD--------------------------MLRATFGV--- 375
WP_049966804_1 DUF63 VFVIWIFDETVF---EDSPR-------------------YAVLLMITVVAVGLGPGTRD--------------------------MLRATFGV--- 375
WP_076145380_1 DUF63 VFVIWIFDETVF---EDSPR-------------------YAVLLMITVVAVGLGPGTRD--------------------------MLRATFGV--- 375
WP_097378845_1 DUF63 VFVIWIFDETVF---EDSPR-------------------YAIMLMITVVAVGLGPGTRD--------------------------MLRATFGV--- 375
WP_008452261_1 DUF63 VFVIWIFDETVF---EDSPR-------------------YAVLLMITVVAVGLGPGTRD--------------------------MLRATFGV--- 375
WP_008452261_1 DUF63 VFVIWIFDETVF---EDSPR-------------------YAVLLMITVVAVGLGPGTRD--------------------------MLRATFGV--- 375
WP_006432643_1 DUF63 IFVIWIFDETVF---EDSPR-------------------YAILLMITVVAVGLGPGTRD--------------------------MLRATFGV--- 375
WP_008452261_1 DUF63 VFVIWIFDETVF---EDSPR-------------------YAVLLMITVVAVGLGPGTRD--------------------------MLRATFGV--- 375
WP_007109662_1 DUF63 VFVIWIFDETVF---EDSPR-------------------YAVLLMITVVAVGLGPGTRD--------------------------MLRATFGV--- 375
WP_049952821_1 DUF63 VFVIWIFDETVF---EDSPR-------------------YAILLMITVVAVGLGPGTRD--------------------------MIRATFGV--- 375
WP_086889541_1 DUF63 VIVLWIFNEEVF---EESPQ-------------------YTYLLLITVVAVGLGPGTRD--------------------------MLRATFGV--- 378
WP_005580009_1 DUF63 VVVVWIFNEEVF---DESPR-------------------YAILLLITVVAVGLGPGTRD--------------------------MLRATFGI--- 377
WP_049927356_1 DUF63 VVVVWIFNEEVF---EESPR-------------------YAYLLMITVVAVGLGPGTRD--------------------------MLRATFGV--- 375
WP_087714455_1 DUF63 VVVIWIFNEEVF---EDSPR-------------------YAILLMITVVAVGLGPGTRD--------------------------MLRATFGV--- 375
WP_013878438_1 DUF63 VVVVWIFDETVF---EDSPR-------------------YAILLMITVVAVGLGPGTRD--------------------------MLRATFGV--- 375
WP_049921111_1 DUF63 VLVVWIFDETVF---EESPR-------------------YAILLMITVVAVGLGPGTRD--------------------------MLRATFGV--- 375
WP_007142789_1 DUF63 VFVVWIFNEEIF---EESPR-------------------YAILLMITVVAVGLGPGTRD--------------------------MLRATFGV--- 374
WP_006651941_1 DUF63 VFVIWIFDETVF---EESPR-------------------YTILLLITVVAVGLGPGTRD--------------------------MLRATFGV--- 375
WP_004216437_1 DUF63 VFVIWIFDETVF---EESPR-------------------YTILLLITVVAVGLGPGTRD--------------------------MLRATFGV--- 375
WP_071402513_1 DUF63 VFVIWIFDETVF---EESPR-------------------YTILLLITVVAVGLGPGTRD--------------------------MLRATFGV--- 375
WP_006666462_1 DUF63 VFVVWIFDDTIF---EESPR-------------------YAILLLITVVAVGLGPGTRD--------------------------MLRATFGI--- 375
WP_049904535_1 DUF63 VFVVWIFDDTIF---EESPR-------------------YAILLLITVVAVGLGPGTRD--------------------------MLRATFGI--- 375
WP_006824405_1 DUF63 VFVVWIFDDTIF---EESPR-------------------YAILLLITVVAVGLGPGTRD--------------------------MLRATFGI--- 375
WP_011323802_1 DUF63 VVVVWVFNEEIF---EESPR-------------------YAYLLFIAVLAVGLGPGTRD--------------------------MLRATLAI--- 375
WP_015409719_1 DUF63 VFVVWVFNDEIF---EDSPR-------------------YAYLLLIAILAVGLGPGTRD--------------------------MLRATLAI--- 377
WP_006883947_1 DUF63 LGIVWLFDERIF---DESPR-------------------YAVLLLVAAAAVGLGPGTRD--------------------------ILRVTFAI--- 377
WP_077207545_1 DUF63 LAVVWIFDREIF---EESPR-------------------YALLLLTAIVAVGLGPGTRD--------------------------MIRATFGI--- 381
ESS12903_1 putative  LFVVWVFDETVF---EDSPR-------------------FTMMLIVTVLAVGLGPGTRD--------------------------MLRATFGI--- 379
WP_008417361_1 DUF63 LFVVWVFNDEIF---EESPQ-------------------YSMLLLVAILAVGLGPGTRD--------------------------TLRATFGI--- 377
WP_066381608_1 DUF63 LFVVWVFNDEIF---EESPR-------------------YSMLLLVAILAVGLGPGTRD--------------------------FLRATFGV--- 372
WP_049947642_1 DUF63 TAVVAIFEREMF---EESPR-------------------YAVLLLIAVTAVGMGPGTRD--------------------------MLRATFGV--- 391
ESS05953_1 putative  ALVVAVFDERTF---EESPR-------------------YAVLLLVAIVAVGLGPGTRD--------------------------LLRATLGV--- 371
WP_096390051_1 DUF63 TFVVWVFNDEMF---AESPR-------------------YTVLLLVTVLAVGLGPGTRD--------------------------MLRATFGI--- 385
```

```
WP_021072749_1 DUF63 TFVVWVFNDEMF---AESPR-------------------YTVLLLITVLAVGLGPGTRD--------------------------MLRATFGI--- 385
WP_049982791_1 DUF63 TFVVWVFNGEMF---EESPR-------------------YTLMLLITVLAVGLGPGSRD--------------------------MLRATFGI--- 383
WP_008585990_1 DUF63 TFVVWVFNGEMY---DESPR-------------------YSLLLLITVLAVGLGPGTRD--------------------------MLRATFGV--- 385
WP_006628721_1 DUF63 TFVVWIFNGEMY---DESPR-------------------YTLLLLITVLAVGLGPGTRD--------------------------MLRATFGV--- 383
WP_006113218_1 DUF63 TFVVWVFNGELF---EESPR-------------------YTLLLLITVLAVGLGPGTRD--------------------------MLRATFGV--- 384
WP_049930034_1 DUF63 TFVVWVFNGEMK---AESPR-------------------YTLLLLITVLAVGLGPGTRD--------------------------MLRATFGV--- 385
WP_049906047_1 DUF63 TFVVWVFNGEMY---EESPR-------------------YTLLLLITVLAVGLGPGTRD--------------------------MLRATFGV--- 385
WP_044965494_1 DUF63 TFVVWVFNGEMY---EESPR-------------------YTLLLLITVLAVGLGPGTRD--------------------------MLRATFGV--- 385
WP_096393195_1 DUF63 TFVVWVFNGEMY---EESPR-------------------YTLLLLITVLAVGLGPGTRD--------------------------MLRATFGV--- 385
WP_049908400_1 DUF63 TFVVWVFNGEMF---EESPR-------------------YTLLLLITVLAVGLGPGTRD--------------------------MLRATFGV--- 385
WP_049908585_1 DUF63 TFVVWVFNGEMF---EESPR-------------------YTLLLLITVLAVGLGPGTRD--------------------------MLRATFGV--- 385
WP_049902631_1 DUF63 TFVVWVFNGEMF---EESPR-------------------YTLLLLITVLAVGLGPGTRD--------------------------MLRATFGV--- 385
WP_049983668_1 DUF63 TFVVWVFNGEMY---EESPR-------------------YTLLLLITVLAVGLGPGTRD--------------------------MLRATFGV--- 384
WP_053772267_1 DUF63 TFVVWVFNGEMY---EESPR-------------------YTLLLLITVLAVGLGPGTRD--------------------------MLRATFGV--- 384
WP_007999151_1 DUF63 TFVVWVFNGEMF---EESPR-------------------YTLMLLITVLAVGLGPGTRD--------------------------MLRATFGI--- 383
WP_015909917_1 DUF63 TFVVWVFNGEMF---DESPR-------------------YTLLLLITVLAVGLGPGSRD--------------------------MLRATFGI--- 384
WP_004050594_1 DUF63 TFVVWVFNGEMF---DESPR-------------------YTLLLLITVLAVGLGPGSRD--------------------------MLRATFGI--- 384
WP_095636035_1 DUF63 TFVVWVFNGEMF---DESPR-------------------YTLLLLITVLAVGLGPGTRD--------------------------MLRATFGI--- 384
WP_008003811_1 DUF63 TFVVWVFNGELF---DESPR-------------------YTLLLLITVLAVGLGPGSRD--------------------------MLRATFGI--- 384
WP_066416034_1 DUF63 TFVVWVFNGELY---EESPR-------------------YTLLLLITVLAVGLGPGTRD--------------------------MLRATFGV--- 385
ESS03170_1 putative  TFVVWVFNGELF---EESPR-------------------YTLLLLITVLAVGLGPGTRD--------------------------MLRATFGV--- 384
WP_089671383_1 DUF63 TFLVWIFNDELF---EESPR-------------------YAILLLVAVVAVGLGPGTRD--------------------------MLRATFGV--- 383
ERH07656_1 putative  TFVIWVFEPDLL---EESPR-------------------YSILLLIAVLAVGLGPGTRD--------------------------VLRATFGV--- 383
ESS10096_1 putative  TFVIWVFEPDLL---EESPR-------------------YSILLLIAVLAVGLGPGTRD--------------------------VLRATFGV--- 383
ESS07824_1 putative  -----------------SPS-------------------------------G-------------------------------------------- 207
ERH05689_1 putative  TFVVWVFEPELF---DETPR-------------------YSILLLIAVLAVGLGPGTRD--------------------------MLRATFGV--- 383
ERH02266_1 putative  TFVVWVFEPELF---DETPR-------------------YSILLLIAVLAVGLGPGTRD--------------------------MLRATFGV--- 383
ESS07825_1 putative  TFVVWVFEPELF---DETPR-------------------YSILLLIAVLAVGLGPGTRD--------------------------MLRATFGV--- 176
WP_049970017_1 DUF63 VAVVWVFDEQIF---EESPR-------------------YAILLLIAILAVGLGPGTRD--------------------------MLRAAFGI--- 377
WP_007979289_1 DUF63 TAVVWVFDEQIF---EESPR-------------------YAILLLIAILAVGLGPGTRD--------------------------MLRATFGI--- 377
WP_007977745_1 DUF63 LAVVWVFDEQIF---EESPR-------------------YAILLLIAIVAVGLGPGTRD--------------------------MLRATFGI--- 377
WP_014556445_1 DUF63 TFVIWVFEPEIF---EESPR-------------------YTMLLLIAVLAVGLGPGTRD--------------------------MLRATFGV--- 379
WP_049935775_1 DUF63 TLVVWLFDAQIF---EDSPR-------------------YAVLLMIAILAVGLGPGTRD--------------------------MLRATIGV--- 377
WP_008325149_1 DUF63 TAVIWVFDEQIF---EESPR-------------------YAILLLIAVLAVGLGPGTRD--------------------------MLRATFGV--- 379
WP_007543631_1 DUF63 TAVIWVFDEQIF---EESPR-------------------YAILLLIAVLAVGLGPGTRD--------------------------MLRATFGV--- 379
WP_049905041_1 DUF63 TAVVWVFDEQIF---EDSPR-------------------YAILLLIAVLAVGLGPGSRD--------------------------MLRATFGV--- 379
WP_049913563_1 DUF63 TAVVWVFDEQIF---EDSPR-------------------YAILLLIAVLAVGLGPGSRD--------------------------MLRATFGV--- 379
WP_049967851_1 DUF63 TAVVWVFDEQIF---EDSPR-------------------YAILLLIAVLAVGLGPGSRD--------------------------MLRATFGV--- 379
WP_049914905_1 DUF63 TAVVWVFDEQIF---EDSPR-------------------YAILLLIAVLAVGLGPGSRD--------------------------MLRATFGV--- 379
WP_049916430_1 DUF63 TAVVWVFDEQIF---EDSPR-------------------YAILLLIAVLAVGLGPGSRD--------------------------MLRATFGV--- 379
WP_049896892_1 DUF63 TAVVWVFDEQIF---EDSPR-------------------YAILLLIAVLAVGLGPGSRD--------------------------MLRATFGV--- 379
WP_049896892_1 DUF63 TAVVWVFDEQIF---EDSPR-------------------YAILLLIAVLAVGLGPGSRD--------------------------MLRATFGV--- 379
WP_049896892_1 DUF63 TAVVWVFDEQIF---EDSPR-------------------YAILLLIAVLAVGLGPGSRD--------------------------MLRATFGV--- 379
WP_058828253_1 DUF63 TAVVWVFDEQIF---EDSPR-------------------YAILLLIAVLAVGLGPGSRD--------------------------MLRATFGV--- 379
WP_058568959_1 DUF63 TAVVWVFDEQIF---EDSPR-------------------YAILLLIAVLAVGLGPGSRD--------------------------MLRATFGV--- 379
WP_049917947_1 DUF63 TAVVWVFDEQIF---EDSPR-------------------YAILLLIAVLAVGLGPGSRD--------------------------MLRATFGV--- 379
WP_049920348_1 DUF63 TAVVWVFDEQIF---EDSPR-------------------YAILLLIAVLAVGLGPGSRD--------------------------MLRATFGV--- 379
WP_089777545_1 DUF63 TAVVWVFDEQIF---EDSPR-------------------YAILLLIAVLAVGLGPGSRD--------------------------MLRATFGV--- 379
WP_008320633_1 DUF63 TAVVWVFDEQIF---EESPR-------------------YAILLLIAVLAVGLGPGSRD--------------------------MLRATFGV--- 379
WP_004060355_1 DUF63 TAVVWVFDEQIF---EDSPR-------------------YAILLLIAVLAVGLGPGTRD--------------------------MLRATFGV--- 379
WP_103426078_1 hypot TFVVWVFEEGVF---EESPR-------------------YTMLLLVAVLAVGLGPGTRD--------------------------MLRATLGV--- 379
WP_009367433_1 DUF63 TLVVWVFEEEIF---EESPR-------------------YTILLLIAILAVGLGPGTRD--------------------------MLRATFGV--- 378
WP_013440552_1 DUF63 TFVVWVFEEGIF---EESPR-------------------YTMLLLIAVLAVGLGPGTRD--------------------------MLRATFGV--- 379
WP_049916626_1 DUF63 TFVVWVFEEGIF---EESPR-------------------YTMLLLVAVLAVGLGPGTRD--------------------------MLRATFGV--- 379
WP_058582837_1 DUF63 VFIIWIFEPEIF---EESPQ-------------------YAMLLLIAALAVGLGPGTRD--------------------------MLRATFGI--- 380
WP_101298124_1 hypot TFVVWVFEDEIM---EESPR-------------------YAILLLIAILAVGLGPGTRD--------------------------MLRATFGI--- 380
PIN95153_1 hypotheti VLMLWTIYEIVV------------------------------------------------------------------------------------ 127
PIU22108_1 hypotheti IVVIIILRNSNM-----SEN-------------------KKNYILLFITLFGFATGLRD--------------------------LFSIVNHLI-- 277
AAR39201_1 NEQ352    YVLLKTY--------------------------------NKPYIRASIFVLGLLTGGRN--------------------------FLIEIALK--- 209
OIR14399_1 hypotheti TAALWMLFSIKE--DDDDPQ-------------------AITLLMMFLITLGLAPGVRD--------------------------MLRIALGT--- 365
OIR20963_1 hypotheti TGVIIALENEEH--KEDQKQ-------------------MISLLILFLLALGLGPGTRD--------------------------ILRIVFGT--- 423
OIR22371_1 hypotheti TGVITALELEKS--NDEDPS-------------------MVALLLMFLLALGLGPGTRD--------------------------ILRIMFGT--- 374
EGQ43935_1 putative  IPVVHYIRESFD------GE-------------------EKAFYIFIITALGLGIATRN--------------------------TLQTLAIS--- 267
MAG21679_1 hypotheti LLILHYVDIEIE-----NEN-------------------LRGFIKIVVLILGLATGLRD--------------------------LFRLGIPA--- 277
PIN85618_1 hypotheti LLIIHYIDQDVK-----DKN-------------------FANFVKIAVIILGFATGGRD--------------------------LLTLGAGTCL- 278
PIN99249_1 hypotheti LAILWYVEKEIQ-----DEK-------------------LKNFIKLFVIIIGFAPGIRD----------------------------AFSLGLTLS 283
AJF59838_1 hypotheti LLIIYYVDKEVK-----DKN-------------------MAGFIKVFIIIMGFAPGIRD--------------------------AFTVGVGTCF- 278
WP_042682145_1 DUF63 IVVYYGLKWLVP-----DGE-------------------ERRYWYLAIYILGLGPAIRD--------------------------PAQMILQLAG- 264
WP_013468033_1 DUF63 IVVYYGLKWLVP-----DEE-------------------ERRYWYLAIYILGLGPAIRD--------------------------PAQMILQLAG- 264
WP_055281581_1 DUF63 VVVYYGLRYLVP-----DEE-------------------ERKFWYLAVYVLGLGPAIRD--------------------------PAQMILQIVS- 263
WP_048160382_1 DUF63 VVVYYGLRYLVP-----DEE-------------------ERKYWYLAIYILGLGPAIRD--------------------------PAQMVLQIVG- 263
WP_042701216_1 DUF63 GVVYYGLKYLVP-----DEE-------------------ERKFWYLAVYILGLGPAIRD--------------------------PAQMILQIG-- 262
WP_004068659_1 DUF63 VAVYYGLKYLVP-----DEE-------------------ERSFWYLAVYILGLGPAIRD--------------------------PAQMILQIG-- 262
WP_058946665_1 DUF63 IVVYYGLRYLVP-----DEE-------------------ERRFWYLAVYILGLGPAIRD--------------------------PAQMILQIG-- 262
WP_014835499_1 DUF63 VAVYYILKYFVP-----DQD-------------------ERNYWYLAIYILGLGPAIRD--------------------------PAQMVLQIVCR 264
WP_010884243_1 DUF63 IIVYYVLKELVL-----DPE-------------------ERYYWYLAVYILGLGPGIRD--------------------------PAQMVLQIPCR 267
WP_013748129_1 DUF63 VVVYYILRNLVQ-----DPE-------------------ERYYWYLAVYILGLGPGIRD--------------------------PTQMVLQIVCR 263
WP_010867251_1 DUF63 FLVYYILRNMVP-----DGD-------------------ERNYWYLAIYILGLGPAIRD--------------------------PAQMVLQIACQ 263
WP_014733359_1 DUF63 IAVYYILTKMVP-----EEE-------------------ERNYWYLAIYILGLGPGIRD--------------------------PAQMVLQIVCQ 264
WP_068319891_1 DUF63 VAVYYILKTMVP-----DEE-------------------ERNYWYLAIYILGLGPAIRD--------------------------PAQMILQIVCQ 264
WP_068576049_1 DUF63 VAVYYILKTMVP-----DEE-------------------ERNYWYLAIYILGLGPAIRD--------------------------PAQMVLQIACQ 264
WP_014013692_1 DUF63 VIVYYGLQKLVD-----DEE-------------------ERRLWYLMVYVLGLGPAIRD--------------------------PAQLVLQIGG- 268
WP_014789520_1 DUF63 VVVYYGLQKLVP-----DEE-------------------ERRLWYLMVYVLGLGPAIRD--------------------------PAQLVLQIGG- 268
WP_088180676_1 DUF63 VAVYYGLQKLVP-----DEE-------------------ERHLWYLMVYVLGLGPAIRD--------------------------PAQLVLQIGG- 268
WP_088864885_1 DUF63 VVVYYGLQKLVP-----DEE-------------------ERRLWYLMVYVLGLGPAIRD--------------------------PAQLVLQIGG- 268
WP_088856755_1 DUF63 LVVYYGLKYLVQ-----DEE-------------------ERHLWYLMVYVLGLGPAIRD--------------------------PAQLVLQIGG- 268
WP_088865981_1 DUF63 VAVYYGLQKLVP-----DEE-------------------ERRLWYLMVYVLGLGPAIRD--------------------------PAQLVLQIGG- 268
WP_013906334_1 DUF63 VTVYYGLKYLVP-----DKE-------------------ERNFWYLAIYILGLGPAIRD--------------------------PAQMVLQIVCG 265
WP_088862658_1 DUF63 IVVYYGLKYLVP-----DEE-------------------ERRFWYLAVYILGLGPAIRD--------------------------PAQLVLQL--- 265
WP_088882956_1 DUF63 VVVYYGLKYLVP-----DEE-------------------ERRYWYLAVYILGLGPAIRD--------------------------PAQMILQL--- 265
WP_012571938_1 DUF63 VIVYYGLKYLVP-----DEE-------------------ERYFWYLAIYILGLGPAIRD--------------------------PAQLVLQIG-- 263
WP_068663933_1 DUF63 VIVYYGLKYLVP-----DEE-------------------ERYFWYLAIYILGLGPAIRD--------------------------PAQLVLQIG-- 263
WP_074631153_1 DUF63 VVVYYGLRYLVP-----DEE-------------------ERYFWYLAIYILGLGPAIRD--------------------------PAQLVLQV--- 262
WP_058939074_1 DUF63 VVVYYGLKYLVP-----DEE-------------------ERRFWYLAIYILGLGPAIRD--------------------------PAQLVLQIGG- 268
WP_088854681_1 DUF63 VVVYYGLKYLVP-----EEE-------------------ERRFWYLAIYILGLGPAIRD--------------------------PAQLVLQIPG- 268
```

```
WP_088885426_1 DUF63 IVVYYGLKYLVT-----DEE-------------------ERRFWYLAIYILGLGPAIRD--------------------------PAQMILQISG- 264
WP_011249694_1 DUF63 VVVYYALKELVT-----DEE-------------------ERRFWYLAIYILGLGPAIRD--------------------------PAQMILQL--- 264
WP_062386972_1 DUF63 VVVYYALKELVT-----DEE-------------------ERRFWYLAIYILGLGPAIRD--------------------------PAQMVLQL--- 264
WP_010478887_1 DUF63 IVVYYGLQRLVP-----DEE-------------------ERRFWYLAIYVLGLGPAVRD--------------------------PAQMVLQV--- 262
WP_088858605_1 DUF63 IAVYYGLKYLVP-----DEE-------------------ERRFWYLAIYILGLGPAVRD--------------------------PAQMVLQL--- 262
WP_042690028_1 DUF63 IVVYYALQELVQ-----DEE-------------------ERRFWYLAVYVLGLGPAIRD--------------------------PAQMILQV--- 264
WP_048150762_1 DUF63 IAVYYGLKHLVP-----DEE-------------------ERRFWYLAIYVLGLGPAIRD--------------------------PAQMVLQI--- 262
WP_048811076_1 DUF63 IAVYYGLKYLVP-----DEE-------------------ERRFWYLAIYVLGLGPAIRD--------------------------PAQMVLQI--- 262
WP_050003256_1 DUF63 IAVYYALRELVP-----DEE-------------------ERHFWYLAIYILGLGPAIRD--------------------------PAQMILQIGG- 266
WP_062372100_1 DUF63 IAVYYALKYLVP-----DEE-------------------ERKFWYLAIYVLGLGPAIRD--------------------------PAQMILQIGG- 264
WP_048165284_1 DUF63 GIVYYGVKYLIP-----NEN-------------------ERKYWYFATFVLGFAPGVRD--------------------------PTQLTLGA--- 264
WP_048148750_1 DUF63 AVVYYGVRYFVK-----DEN-------------------ERRYWYFAMFVLGLGPGFRD--------------------------PAQLVLGA--- 264
OYT53462_1 hypotheti LLVIYALEKYNT--EEEDEI-------------------LYRMLKVLVFILGIGPGTRN--------------------------TMLPALSL--- 252
KYC51152_1 hypotheti LVVIYLLDTLYKEERKNFKEKEVTIFRYRFKVTTLMMNNFYLSIKVLIFILGFGPGFRN-------------------------TLLPSLF----- 274
KYC46003_1 hypotheti LVVIYLLDTLYKEERKNFKEKEVTIFRYRFKVTTLMMNNFYLSIKVLIFILGFGPGFRN-------------------------TLLPSLF----- 274
KYC48643_1 hypotheti LVVIYLLDTLYKEERKNFKEKEVTIFRYRFKVTTLMMNNFYLSIKVLIFILGFGPGFRN-------------------------TLLPSLF----- 274
KYC55321_1 hypotheti LIVIYLLDTLYKEEQNNFKEKEIKISKYRLKITAFMINNFYLSIKVLIFILGFGPGFRN-------------------------TLLPSLF----- 274
KYC57927_1 hypotheti LIVIYLLDTLYKEEQNNFKEKEIKISKYRLKITAFMINNFYLSIKVLIFILGFGPGFRN-------------------------TLLPSLF----- 274
KYC57171_1 hypotheti LIVIYLLDTLYKEEQNNFKEKEIKISKYRLKITAFMINNFYLSIKVLIFILGFGPGFRN-------------------------TLLPSLF----- 274
OIO20701_1 hypotheti FVAAYFICREAQ-----DEN-------------------ERNYLFLLIIVFGLAPGIRD--------------------------VLRMVCGA--- 311
PIT83986_1 hypotheti AVVVSVIRR------EEDRR-------------------LVLYLLLVILIIGLAPGLRD--------------------------VLRMVAGT--- 315
OIO24760_1 hypotheti FVVAYLLSR------EKDGE-------------------EKTFVALLITIFGLAPGVRD--------------------------ALRILAGV--- 272
OIO24701_1 hypotheti LAAVTIVRNEFI--KKEDAE-------------------KKTYVLLLLTIFGLAPGARD--------------------------VLRIMAGV--- 284
OIO27021_1 hypotheti FAVVELLRREDK-----DSQ-------------------ERRFILLLVAIFGFAPGLRD--------------------------LLRIVCGV--- 273
OIO26762_1 hypotheti FAAIWLLRREKE-------E-------------------ERNFVLILIAVFGLAPGLRD--------------------------ALRIAAGV--- 269
PIN95811_1 hypotheti FAAIWLLRREKE-------E-------------------ERNFVLILIAVFGLAPGLRD--------------------------ALRIAAGV--- 269
PIO01637_1 hypotheti FAAIWLLRREKE-------E-------------------ERNFVLILIAVFGLAPGLRD--------------------------ALRIAAGV--- 269
PIO02820_1 hypotheti FAVVELLRREDK-----DSQ-------------------ERRFILLLVAIFGFAPGLRD--------------------------LLRIVCGV--- 273
PJD01038_1 hypotheti FAVVELLRREDK-----DSQ-------------------ERRFILLLVAIFGFAPGLRD--------------------------LLRIVCGV--- 273
PIZ91366_1 hypotheti FAAIWLLRREKE-------E-------------------ERNFVLILIAVFGLAPGLRD--------------------------ALRIAAGV--- 269
WP_013100570_1 DUF63 ILILYILNREEM-----DPN-------------------LRNIIKLIIMALGLAPGLRN--------------------------LFRTIMGV--- 265
WP_004590770_1 DUF63 LLALYVVNREVE-----DEN-------------------IKNIIKLCIMALGMAPGMRD--------------------------LFRMIMGV--- 267
WP_048196979_1 DUF63 LLALYILNREVE-----DEN-------------------IRNIIKLCIMALGLAPGLRD--------------------------LFRTVMGV--- 267
WP_015791538_1 DUF63 LLVLYILNREIE-----DEN-------------------IKNIIKLCIMALGLAPGLRN--------------------------LFRTIMGV--- 267
WP_048202292_1 DUF63 LLVLYVLNREIE-----DEN-------------------IRNIIKLCIMALGLAPGLRN--------------------------LFRTVMGV--- 267
WP_012981280_1 DUF63 LLALYILNREVK-----DEN-------------------IKNIIKLCIMALGLAPGLRD--------------------------LFRTVMGV--- 267
WP_064496496_1 DUF63 LLALYILNKEVE-----DEN-------------------IKNIIKLCIMALGLAPGLRD--------------------------LFRTVMGV--- 267
WP_011972741_1 DUF63 FSVLYVLNHEVD-----NKN-------------------LRNIIKITIMALGFAPGLRN--------------------------LFRIVMGV--- 272
WP_013798289_1 DUF63 LIALYLINEEVD-----NKD-------------------LKNILKITIMCLGLAPGLRN--------------------------LLRTIMGV--- 269
WP_013181100_1 DUF63 VLVLELFNREVE-----NKN-------------------LRNILKIAVLSLGLAPGLRN--------------------------LFRTIMGV--- 268
WP_013867628_1 DUF63 IAVLYIINKEIN-----NRD-------------------LRNILKITIMALGLAPGLRD--------------------------LFRTIMGV--- 273
WP_018153391_1 DUF63 VLVLYIINKEID-----NKD-------------------LRNILKITIMSLGLAPGLRN--------------------------LFRIVMGV--- 274
WP_011170272_1 DUF63 LLVLDIFNKEVK-----DDN-------------------LRNILKITVMALGLAPGLRN--------------------------LFRTVMGV--- 268
WP_012066151_1 DUF63 YFVLDVFNKEVK-----DDN-------------------LRNILKITVMALGLAPGLRN--------------------------LFRMIMGV--- 268
ODS43013_1 hypotheti VAVLYLIDKSVE-----DAL-------------------ERRFIKFVILVLGLGPAIRD--------------------------VALMVL----- 275
OIQ06085_1 hypotheti IPALYFIDK------IDDKT-------------------TRRMIKIVIFILGISPAIRN--------------------------FTRILLGV--- 330
PIN67069_1 hypotheti IPALYFIDK------IDDKT-------------------TRRMIKIVIFILGISPAIRN--------------------------FTRILLGV--- 330
PIV28099_1 hypotheti IPALYFIDK------IDDKT-------------------TRRMIKIVIFILGISPAIRN--------------------------FTRILLGV--- 330
PJC13070_1 hypotheti IPALYFIDK------IDDKT-------------------TRRMIKIVIFILGISPAIRN--------------------------FTRILLGV--- 330
PIZ29927_1 hypotheti IPALYFIDK------IDDKT-------------------TRRMIKIVIFILGISPAIRN--------------------------FTRILLGV--- 330
PKP60688_1 hypotheti IPALYFIDQ------IDDKT-------------------TRRMIKIVIFILGISPAIRN--------------------------FTRILLGV--- 276
WP_012956954_1 DUF63 VAVLYIIDQYFD-----DET-------------------IKSLLKLTVFVLGLAPGLRN--------------------------FLTMAIGV--- 263
WP_067148430_1 DUF63 VAVLYIIDRYFE-----DTT-------------------VKNLLKLTVFVLGLAPGLRN--------------------------ILTLAIATI-- 281
WP_080460538_1 DUF63 CISLYIIDKYVD-----DEI-------------------LSGTLKLAIFILGLAPGVRN--------------------------FLSLAIGTWI- 269
WP_010877090_1 DUF63 LTALYAIDRYVE--SDMDSE----------------------ALKLVIFILGLAPGLRN--------------------------FLSLSMAV--- 263
WP_013294901_1 DUF63 LLGLYAIDRYVE-----DPG-------------------DSALLKLVIFILGLAPGLRN--------------------------FLSLSMAV--- 263
WP_010877090_1 DUF63 LTALYAIDRYVE--SDMDSE----------------------ALKLVIFILGLAPGLRN--------------------------FLSLSMAV--- 263
BAZ99473_1 hypotheti LTALYAIDRYVE--SDMDSE----------------------ALKLVIFILGLAPGLRN--------------------------FLSLSMAV--- 263
PKL66404_1 hypotheti LSALYIIDTSIE-----DKT-------------------IKNTLKLSIFILGLAPGLRN--------------------------FLSLIMGV--- 271
WP_023991171_1 DUF63 LPALYIIDTYVE-----DKT-------------------IKNMLKLAIFILGLAPGIRN--------------------------FLSLAMGT--- 264
WP_048072898_1 DUF63 LPALYIIDTYVE-----DKT-------------------IRNMLKLAIFILGLAPGIRN--------------------------FLSLAMGT--- 264
WP_100905180_1 hypot LPSLYVIDTYVE-----DKT-------------------IRNMLKLAIFILGLAPGLRN--------------------------FLSLAMGT--- 264
WP_100907231_1 hypot LPSLYVIDTYVE-----DKT-------------------IRNMLKLAIFILGLAPGLRN--------------------------FLSLAMGT--- 264
WP_100907231_1 hypot LPSLYVIDTYVE-----DKT-------------------IRNMLKLAIFILGLAPGLRN--------------------------FLSLAMGT--- 264
WP_048081925_1 DUF63 LAALYVVDEYVE-----DNT-------------------IKNMLKLAIFILGLAPGLRD--------------------------FLSLSIGT--- 263
WP_048081925_1 DUF63 LAALYVVDEYVE-----DNT-------------------IKNMLKLAIFILGLAPGLRD--------------------------FLSLSIGT--- 263
WP_069585525_1 DUF63 LAAIYVIDEYIE-----DDT-------------------IKNMLKLAIFILGLAPGLRD--------------------------FLSLSIGT--- 263
WP_069585525_1 DUF63 LAAIYVIDEYIE-----DDT-------------------IKNMLKLAIFILGLAPGLRD--------------------------FLSLSIGT--- 263
WP_013643671_1 DUF63 LAALYVIDSYIE-----DKT-------------------IANMLKLAIFILGFAPGLRN--------------------------FLSLIMGTAL- 265
WP_013824571_1 DUF63 LSALYVIDSYVE-----DKT-------------------IRNMLKLAIFILGLAPGLRN--------------------------FLSIIMGTF-- 264
WP_048192110_1 DUF63 LGALYIIDSYIE-----DRT-------------------IANMLKLAIFILGLAPGLRN--------------------------FLSLIMGT--- 267
WP_081810122_1 DUF63 VLIIMMLERESR--GDKTMD-------------------EKYMLILTLIVLGLSMGARN--------------------------LLVIAFGI--- 302
WP_097298965_1 DUF63 MLIILMLEKDST--REKNTD-------------------EKYMLTLTLIVLGFSMGARN--------------------------LLAMVFGV--- 278
PIO00292_1 hypotheti LVVIYVINKDVK-----DEG-------------------FKNLLKFAILTLGLAPGVRD--------------------------LLRLAMLV--- 265
WP_011833679_1 DUF63 IPAIWILEKFRK--EEGMEA-------------------IWHLVLLAMIVVGLAPGLRD--------------------------VLRMVLGI--- 278
WP_042698332_1 DUF63 IPAIWVLEKFRK--EEGMES-------------------IWHLVLLAMIVVGLAPGLRD--------------------------VLRMVLGI--- 278
WP_042698332_1 DUF63 IPAIWVLEKFRK--EEGMES-------------------IWHLVLLAMIVVGLAPGLRD--------------------------VLRMVLGI--- 278
WP_011448164_1 DUF63 IPGIYILELYRR---EGASG-------------------IWYLILLAMIMVGLAPGIRD--------------------------MMRMVLYV--- 281
WP_007314214_1 DUF63 FPGIYILQRFRS---EGPQA-------------------LWHLILLAMITVGLAPGIRD--------------------------MVRMVLYV--- 286
WP_042705632_1 DUF63 IPGIYILEMYRR--DGGNQA-------------------LWHMILFAMIVVGLAPGIRD--------------------------MVRMVLYV--- 282
WP_013329717_1 DUF63 IPAIYILQIYRQ---EGNKA-------------------LFHLIILAMIMVGLAPGIRD--------------------------MVRMILYV--- 281
WP_004077722_1 DUF63 IPAIYILEMYRK---EGNKA-------------------LWHLIILAMIMVGMAPGIRD--------------------------MVRMILYV--- 281
WP_048150927_1 DUF63 IPAIYILQIYRQ---EGNKA-------------------LFHLIILAMIMVGLAPGIRD--------------------------MVRMILYV--- 281
WP_012107531_1 DUF63 FPAIYIMQLYRK---EANAT-------------------FWHLVLLAMIVVGLAPGIRD--------------------------MIRMVLYV--- 288
PKL70081_1 hypotheti FPAIYIMEQYRK---EANPA-------------------FWHLVLLAMITVGLAPGIRD--------------------------MTRMVLYV--- 282
WP_015285007_1 DUF63 FPAIYIMQLYRK---EANTA-------------------FWHLVLLAMIVVGFAPGVRD--------------------------MVRMVLYV--- 288
PKL64570_1 hypotheti FPAIYIMELYRK---EANTA-------------------FWHLVLLAMIVVGFAPGIRD--------------------------MVRMVLYV--- 288
WP_015286404_1 DUF63 FPAIYILQQYRK---EAAPV-------------------FWHLVLLAMIIVGFAPGVRD--------------------------MVRMILFT--- 281
WP_012617618_1 DUF63 IPGIYILELYRK---EGASA-------------------FWHLVVFAMIVVGLAPGLRD--------------------------MIRMVLYV--- 281
WP_014867374_1 DUF63 IPAIYILEMYKK---EGSPG-------------------LWHLILLAMIVVGMAPGIRD--------------------------MVRMVLYV--- 281
CVK33108_1 conserved IPAIYILEMYKK---EGSPG-------------------LWHLILLAMIVVGMAPGIRD--------------------------MVRMVLYV--- 281
WP_066956331_1 DUF63 IPAIYVLELYRR---EGNPD-------------------LWHLILLAMIVVGMAPGIRD--------------------------LVRMVLYV--- 281
WP_011844879_1 DUF63 IPAVYILEMYRR---EGDPG-------------------LWHLILLAMIVVGMAPGIRD--------------------------LARMVLHV--- 281
WP_048181868_1 DUF63 VPAVYVLEMYRR---EGDSG-------------------LWHLILLAMIVVGMAPGIRD--------------------------LARMVLYV--- 281
WP_067073014_1 DUF63 VPAVYVLEMYRR---EGSPG-------------------LWHLILLAMIVVGMAPGIRD--------------------------LVRMVLYV--- 281
```

```
PKL62929_1 hypotheti IPAIYVLEMYRR---EGNPD-------------------LWHLILLAMIVVGMAPGIRD--------------------------LVRMVLYV--- 281
WP_004039890_1 DUF63 VPAIYVLELYRK---EGNVA-------------------FWHLVLLAMIVVGMAPGVRD--------------------------MMRMVIYV--- 281
WP_067049225_1 DUF63 IPAIYVLEMYRK---EGNPA-------------------FWHLVVLAMIVVGLAPGIRD--------------------------MMRMVIYV--- 281
KYK37884_1 hypotheti AVVLSAVEDVKE------EG-------------------YKNFIKLVILILGAAPGMRD--------------------------LLRIFFMT--- 262
KYK28019_1 hypotheti GIALSAVEDVKE------EG-------------------YKTFIKLVILILGAAPGIRD--------------------------LLRIFFMT--- 262
OYT57835_1 hypotheti LLVLYFVDKERD------IN-------------------MRNLIKLAILVLGIAPGLRN--------------------------TLRLAMGV--- 265
OYT33350_1 hypotheti LGVLYLIDTSKE-----PEK-------------------LKNYLKLILTILGLAPGLRD--------------------------GLRMTFGV--- 266
WP_012964745_1 DUF63 FLILLYLDKVDE-----EEN-------------------LKGFIKFVIMVLGLAPGIRN--------------------------ALRMTFGV--- 257
WP_048091573_1 DUF63 VPILYLIDSERT---KEDEN-------------------LLGFIKFVVFTLGFAPGIRN--------------------------GLRMMFGV--- 262
WP_048096399_1 DUF63 IPILYLLDREEE-----EES-------------------LAGFIKFVLFTLGFAPGIRN--------------------------GLRMTFGV--- 260
WP_012940099_1 DUF63 TLVLWILEKERD------VQ-------------------LKNFIKFVLLVLGLAPATRD--------------------------ITRMIFYV--- 262
WP_010877969_1 DUF63 YALLYLLDTSEE-----DKK-------------------LRNYVKFVLVVLGLAPGLRD--------------------------GIRMMFGT--- 276
WP_013682839_1 DUF63 AAILYVLDSSRD---EEDTE-------------------LINFVKFVLIVLGLAPGLRD--------------------------ALRMTFQV--- 269
WP_015591141_1 DUF63 LAVLYLLDTSED-----DIH-------------------FVNFIKFVLVVLGLAPGMRD--------------------------ALRMCFAT--- 266
WP_012035887_1 DUF63 IPVIYIIEKYFK----DDPQ-------------------MYYGIKFVLLVLGAGPGIRN--------------------------TLHMTFVSA-- 282
WP_014404626_1 DUF63 IPVVYLIERYFV----EEKD-------------------LYYVLKFVLLVLGFGPGIRN--------------------------TLELVFVHS-- 277
BAI60262_1 conserved IPVIYLIEKYFI----EDKD-------------------LYYVLKFVLLVLGFGPGIRN--------------------------TLELVFIHG-- 277
WP_042684156_1 DUF63 LPVLYLIDVHMR---DEGHD-------------------FIQLLKLTLLVLGLAPAMRN--------------------------TLRMTLGI--- 281
OFV68024_1 membrane  IPLLYLLDRELH--DAADAD-------------------IKGILILTLIVVGLAPAIRN--------------------------TTRMVFGI--- 231
WP_013720253_1 DUF63 LPALFMIDKAID-----DRS-------------------LRNLTKLALITLGLAPAVRN--------------------------TLRLALGV--- 271
WP_014587756_1 DUF63 IPVLAVVDGALR----DDPS-------------------LKNLTLFALLTLGLAPAVRN--------------------------TLRLTLGI--- 267
ABK14947_1 Protein o LPLLVLLDESLR----NDRD-------------------IMGILVLTLLVLGLAPATRN--------------------------TLRMMLGI--- 273
OKY79140_1 putative  ILSIYLIEKTFL----KEDE-------------------FKVLLYLVLIVLGLAPALRN--------------------------TLRLTLGV--- 279
WP_086637003_1 DUF63 LTVIWLVDSVFKEGKEEDVE-------------------LKTLVLLAILVLGLAPAIRN--------------------------TLRLTLGV--- 285
WP_048089097_1 DUF63 IPVLYMLDHQFD--NDKEKK-------------------LINLMKLAILVLGLSPAVRN--------------------------TLRLLMGV--- 285
WP_097298272_1 DUF63 IPVLYMLDNKLE--DEKEKK-------------------LIGIIKLAILVLGLSPAVRN--------------------------TLRILMGV--- 286
WP_013897788_1 DUF63 ILIIYVLDTYIS--EKEEAE-------------------LKNIVKMVIILGLSPAIRN--------------------------TLRITFGI--- 282
WP_015323982_1 DUF63 IPVLYLLDTNFN-EDEESRN-------------------LRTFVKLVILVLGLSPACRN--------------------------TIRMVFGI--- 283
WP_011499115_1 DUF63 LPVIYIIDTQFV-ADEESKR-------------------LRTFVKMVIIILGLSPACRN--------------------------TIRMAFGI--- 283
WP_013037846_1 DUF63 IPVIYVLDTQFN-DSEESIS-------------------LRTFVKMVIIVLGLSPATRN--------------------------TIRMALGI--- 283
WP_048205640_1 DUF63 LPVIYLLDTQFT-ADEESRN-------------------LRTFVKMVIIVLGLSPACRN--------------------------TIRMALGI--- 283
WP_072561629_1 DUF63 IPVIYVLDSQFN-DSEESIS-------------------LRTFVKMVIIVLGLSPATRN--------------------------TIRMALGI--- 283
WP_072360082_1 DUF63 IPVIYVLDSQFN-DSEESIS-------------------LRTFVKMVIIVLGLSPATRN--------------------------TIRMALGI--- 283
WP_096711793_1 DUF63 IPVIYVLDTQFN-DSEESIS-------------------LRTFVKMVIIVLGLSPATRN--------------------------TIRMALGI--- 283
ODV50598_1 hypotheti IPVIYVLDTQFN-DSEESIS-------------------LRTFVKMVIIVLGLSPATRN--------------------------TIRMALGI--- 283
WP_013194042_1 DUF63 IPVLYLLDSYFN-EDRESRN-------------------LKTFVKIVILILGLAPACRN--------------------------TIRMTLGI--- 283
WP_048178067_1 DUF63 IGVLYVLDTQFE-DDEESVN-------------------LKMLIKMVILILGLSPATRN--------------------------TIRMMLGI--- 285
WP_048127111_1 DUF63 VGVLYALDTQFE-KDEESNN-------------------LKMLIKMVILVLGLSPATRN--------------------------TIRMMLGI--- 285
WP_011023594_1 DUF63 VGVIYALDTQFE-KDEESEN-------------------LKMLIKMVILILGLSPATRN--------------------------TIRMMLGI--- 289
WP_048184587_1 DUF63 LGVIYALDTQFE-KDEESEN-------------------LKMLIKMVILILGLSPATRN--------------------------TIRMMLGI--- 285
WP_011032542_1 DUF63 VGVLYALDTQFE-KDDESTN-------------------LKMLIKMVILILGLSPATRN--------------------------TIRMMLGI--- 285
WP_011032542_1 DUF63 VGVLYALDTQFE-KDDESTN-------------------LKMLIKMVILILGLSPATRN--------------------------TIRMMLGI--- 285
WP_048129141_1 DUF63 VGVLYALDTQFE-KDDESEN-------------------LKMLIKMVILILGLSPATRN--------------------------TIRMMLGI--- 285
WP_048129141_1 DUF63 VGVLYALDTQFE-KDDESEN-------------------LKMLIKMVILILGLSPATRN--------------------------TIRMMLGI--- 285
WP_048169891_1 DUF63 VGVLYALDTQFE-KDEESNN-------------------LKMLIKMVILILGLSPATRN--------------------------TIRMMLGI--- 285
WP_048137175_1 DUF63 VGVLYALDTQFE-KDEESNN-------------------LKMLIKMVILILGLSPATRN--------------------------TIRMMLGI--- 285
WP_048137175_1 DUF63 VGVLYALDTQFE-KDEESNN-------------------LKMLIKMVILILGLSPATRN--------------------------TIRMMLGI--- 285
WP_048137175_1 DUF63 VGVLYALDTQFE-KDEESNN-------------------LKMLIKMVILILGLSPATRN--------------------------TIRMMLGI--- 285
WP_048137694_1 DUF63 VGVLYALDTQFE-KDEESTN-------------------LKMLIKMVILILGLSPATRN--------------------------TIRMMLGI--- 285
WP_048168126_1 DUF63 IGVLYVLDTQFE-DDERSSN-------------------LKVLIKLVILILGLSPATRN--------------------------TVRMMLGI--- 285
WP_048118082_1 DUF63 VGVLYVLDTQFE-DDERSLN-------------------LKMLIKMVILILGLSPATRN--------------------------TIRMMMGI--- 285
WP_048158120_1 DUF63 VGVLYVLDTQFE-DDERSLN-------------------LKMLIKMVILILGLSPATRN--------------------------TIRMMMGI--- 285
WP_011305329_1 DUF63 VGVLYVLDTQFE-DDERSLN-------------------LKVLIKMVILILGLSPATRN--------------------------TIRMMMGI--- 285
WP_054298619_1 DUF63 IGVLYVLDTQFE-DDERSLN-------------------LKVLIKLVILILGLSPATRN--------------------------TVRMMLGI--- 285
ALK05385_1 hypotheti IGVLYVLDTQFE-DDERSSN-------------------LKVLIKLVILILGLSPATRN--------------------------TVRMMLGI--- 285
WP_015052897_1 DUF63 LPVIYILDTQFN-EDEESRT-------------------LKTFLKLVIIMLGLAPAVRN--------------------------TIRMMLGV--- 282
WP_023846134_1 DUF63 IPVLYVLDNHFD-EDEESET-------------------LKTFLKLVIIVLGLSPACRN--------------------------TIRMAFGV--- 282
OIN88451_1 hypotheti VAVLNILDRYSD-----DKE-------------------FTNYLKIMIGVLGFVTGLRD--------------------------LLRLVLLV--- 257
PIX50278_1 hypotheti VAVLNILDRYSD-----DKE-------------------FTNYLKIMIGVLGFVTGLRD--------------------------LLRLVLLV--- 257
PIW41402_1 hypotheti VAVLNILDRYSD-----DKE-------------------FTNYLKIMIGVLGFVTGLRD--------------------------LLRLVLLV--- 257
PIY35178_1 hypotheti VAVLNILDRYSD-----DKE-------------------FTNYLKIMIGVLGFVTGLRD--------------------------LLRLVLLV--- 257
PJB74886_1 hypotheti VAVLNILDRYSD-----DKE-------------------FTNYLKIMIGVLGFVTGLRD--------------------------LLRLVLLV--- 257
PIZ33651_1 hypotheti VAVLNILDRYSD-----DKE-------------------FTNYLKIMIGVLGFVTGLRD--------------------------LLRLVLLV--- 257
WP_048165029_1 DUF63 LLALYLMREIQE--SESDKE-------------------LMDFIKMVIFILGFAPGTRN--------------------------LLRMLMGV--- 268
WP_042681046_1 DUF63 MIVLYLMKELQE--TESDKE-------------------LMDFIKTVVFILGFAPGTRN--------------------------LLRMLMGV--- 268
WP_013467319_1 DUF63 MVALYLMKELQE--TESDKE-------------------LMDFIKTVIFILGFAPGTRN--------------------------LLRMLMGV--- 268
WP_048152160_1 DUF63 AVALYLMDEIEE--TEADRE-------------------LMDFIKTVMFILGFAPGTRN--------------------------LLRMLMGV--- 268
WP_015849008_1 DUF63 MVALYLMEYLQE--TESDKE-------------------LMDFIKMVMFILGFAPGTRN--------------------------LLRMLMGV--- 268
WP_004069276_1 DUF63 IVALYLMEELQE--SESEKE-------------------LMDFIKMVMFILGFAPGTRN--------------------------LLRMLMGV--- 268
WP_058946638_1 DUF63 MVALYLMEELQE--SESEKE-------------------LMDFIKMVMFILGFAPGTRN--------------------------LLRMLMGV--- 268
WP_042701551_1 DUF63 MVALYLMEELEE--SESDKE-------------------LMDFIKMVMFILGFAPGTRN--------------------------LLRMLMGV--- 268
WP_055282692_1 DUF63 MIALYLMEKLEE--TESDKE-------------------LMDFIKMVMFILGFAPGTRN--------------------------LLRMLMGV--- 268
WP_013906023_1 DUF63 YVALWLMRELER--EGEDKE-------------------LLDFVRMVIFILGFAPGTRN--------------------------LLRMLMGV--- 266
WP_011013158_1 DUF63 FAAVWLLKELEK--EGEEKE-------------------LLDFIKMVIFILGFAPGTRN--------------------------LLRMLMGV--- 267
WP_014733166_1 DUF63 YAALWIMKKLEE--EGEEKE-------------------LLNFIKMVIFILGFAPGTRN--------------------------LLRMLMGA--- 267
WP_068322889_1 DUF63 YLALWIMKKLEE--EGEDKE-------------------LLDFIRMVIFILGFAPGTRN--------------------------LLRMLLGA--- 267
WP_068575625_1 DUF63 YLALWIMKKLEE--EGEDRE-------------------LLDFIRMVIFILGFAPGTRN--------------------------LLRMLLGA--- 267
WP_010885960_1 DUF63 YTALWIMVKLEE--EGEEEE-------------------LLDFIKMVIFILGFAPGTRN--------------------------LLRMLMGV--- 267
WP_010867401_1 DUF63 FIALWIMEKLEK--EGEDKE-------------------LLDFIRMVIFILGFAPGTRN--------------------------LLRMLMGV--- 267
WP_013747987_1 DUF63 YTALWIMLKLEE--EGEEKE-------------------LLDFIRMVIFILGFAPGTRN--------------------------LLRMLMGV--- 268
WP_068664049_1 DUF63 LLVVYLIDHVVE--NEAERD-------------------MGEFVKLVIFLLGFAPGTRN--------------------------LLRMFMGV--- 270
WP_011249739_1 DUF63 IPVVYLLDRMME---DEDPE-------------------LINFVKLAIFVLGFGPGTRN--------------------------LLIALLG---- 269
WP_062386854_1 DUF63 IPVVYLLDRMME---DEDPE-------------------LINFVKLVIFILGFGPGTRN--------------------------LLITLLG---- 269
WP_050003781_1 DUF63 TIVVYVLDKFMT---DEDPE-------------------LINFVKLTVFILGFGPGTRN--------------------------VLITLLG---- 269
WP_042690699_1 DUF63 TLVIYVLDRLMT---DEDPE-------------------LINFVKLTVFILGFGPGTRN--------------------------LLITLLR---- 269
WP_088885212_1 DUF63 VPIVLILDRYME---DEDPD-------------------LINFVKLAMFILGFGPGTRN--------------------------LLIALMGA--- 270
WP_010478519_1 DUF63 IPIVWILDRHME---GEDPE-------------------LINFVKLAMFILGFGPGTRN--------------------------LLITLMGA--- 270
WP_088858240_1 DUF63 LPVVWILDRMME---DEDPD-------------------LINFVKLAMFILGFGPGTRN--------------------------LLITLMGV--- 270
WP_015859016_1 DUF63 LPIVWILDVEMK---DEDPD-------------------LINFVKLAMFILGFGPGTRN--------------------------LLITLMGVGR- 272
WP_014121940_1 DUF63 LPIVWILDVEMK---DEDPD-------------------LINFVKLAMFILGFGPGTRN--------------------------LLITLMGVGR- 272
WP_062373911_1 DUF63 IPVVWILDVEMK---DEDPD-------------------LINFVKLAMFILGFGPGTRN--------------------------LLITLMGVGT- 272
WP_088882924_1 DUF63 LPIVWVLDGWME---EEDRD-------------------LINFVKLTVFVLGFGPGTRN--------------------------LLIMLMGG--- 270
WP_088862911_1 DUF63 WPIVWILDNWME---DEDPD-------------------LINFVKLTIFVLGFGPGTRN--------------------------LLIALMGG--- 270
WP_012572006_1 DUF63 LPIVWILDREMK---DEDPE-------------------LINFVKLAMFILGFGPGTRN--------------------------LLITLMG---- 269
WP_088854707_1 DUF63 LPVVWILDREMK---DEDPD-------------------LINFVKLTVFILGFGPGTRN--------------------------LLIMLMGGG-- 271
```

```
WP_014789474_1 DUF63 LPVVWILDREMK---EEDPD-------------------LINFVKLTIFILGFGPGTRN--------------------------LLIMLMGG--- 269
WP_088180728_1 DUF63 LPIVWILDRAMA---DEDPD-------------------LINFVKLTMFILGFGPGTRN--------------------------LLIMLMGG--- 270
WP_088864937_1 DUF63 LPIVWILDRAMA---DEDPD-------------------LINFVKLTMFILGFGPGTRN--------------------------LLIMLMGG--- 269
WP_055429686_1 DUF63 LPIVWILDRWMK---DEDPD-------------------LINFVKLTMFILGFGPGMRN--------------------------LLIMLMGG--- 270
WP_088865932_1 DUF63 LPIVWILDRAMA---DEDPD-------------------LINFVKLTMFILGFGPGTRN--------------------------LLIMLMGG--- 270
WP_014013642_1 DUF63 LPVVWILDRWMA---DEDPD-------------------LINFVKLTVFILGFGPGTRN--------------------------LLIMLMGG--- 270
WP_088856804_1 DUF63 LPVVWILDRGMG---DEDPD-------------------LINFVKLTIFILGFGPGTRN--------------------------LLIMLMGG--- 270
WP_058939475_1 DUF63 LPVVWILDRWMA---DEDPD-------------------LINFVKLTVFILGFGPGTRN--------------------------LLIMLMGG--- 269
EHR77276_1 conserved GAIVWLFSEMKV--EQRQRH-------------------LRLLIVLAVLIVGLAPGLRD--------------------------IGRLTLGV--- 461
OUV40025_1 hypotheti LGIPGIVCVVLYRIGRDDAR-------------------QLKLTGFEAGVLPEGISIKSWETEEKVVANHPIEQLSNKALLASPLVLAMIFGQL-- 360
MBJ52984_1 hypotheti GVLILVFSKVRI--EQNQKH-------------------LRLLIVLALLAVGLAPGLRN--------------------------LGRNVLGV--- 444
PDH23744_1 hypotheti GLIYYVFYSLNF--ERREQH-------------------LRLLIGLAMLVVGMAPGLRD--------------------------VGRLTLGV--- 447
PDH25468_1 hypotheti GIIFWFYSLANF--EHRQQH-------------------LRLLIGLALMVVGMAPGLRD--------------------------VGRLMLGV--- 432
```
