## Supplementary Data 5 for "Beyond bacterial paradigms: uncovering the functional significance and first biogenesis machinery of archaeal lipoproteins"

**Supplementary Data 5. Multiple alignment of arCOG02178 proteins.** The alignment was colored using http://www.bioinformatics.org/sms2/color_align_cons.html tool with default parameters for amino acid grouping and consensus 90%. The characterized protein from *Hfx. volcanii* (WP_004042703, HVO_2611, AliB) is highlighted with red font, and potential active-site residues in that protein are highlighted in yellow.

```
WP_006882700_1 DUF63 ------------------------------------------------------MQILPSGLAIPPLPYVITLVVLTAAAVAGLYLLDPPVTSRQVVALA 46
WP_015790720_1 DUF63 ------------------------------------------------------MALLPSGLVVPALPYLLVLGLGTLLTTLLLLGFEPPVGKLHVLALL 46
WP_008524992_1 DUF63 -------------------------------------------------------MVLPSGFVVPALPYLVVLGLGLLLTTLLLLGLEPPVGKGHVLALL 45
WP_015762485_1 DUF63 ------------------------------------------------------MQILPTSFALPPLPYLVVLLGGAGLVTALLSAAEPEIDQRAVVSLL 46
WP_018257825_1 DUF63 ------------------------------------------------------MQILPTSFALPPLPYLVVLLGGAGLVTALLSAAEPEIDQRAVVSLL 46
WP_101349367_1 hypot -------------------------------------------------------MVLPSGLALPPLEYTVALLAGTLVVTALLYALEPPIDQRTVVALT 45
WP_004518630_1 DUF63 -------------------------------------------------------MVLPSGLALPPLEYTVALLAGTLVVTALLYALEPPIDQRTVVALT 45
WP_004963804_1 DUF63 -------------------------------------------------------MVLPSGLALPPLEYTVALLAGTLVVTALLYALEPPINQRTVVALT 45
WP_007189047_1 DUF63 -------------------------------------------------------MVLPSGLALPPLEYTVALLAGTLVVTALLYALEPPINQRTVVALT 45
WP_004593738_1 DUF63 -------------------------------------------------------MVLPSGLALPPLEYTVALLAGTLVVTALLYALEPPIGQRTVVALT 45
WP_005535867_1 DUF63 -------------------------------------------------------MVLPSGLALPPLEYTVALLAGTLVVTALLYALEPPIDQRTVVALT 45
WP_053967006_1 DUF63 -------------------------------------------------------MILPSGLALPPLEYTVALLAGTLVVTALLYALEPPIDQRTVVALT 45
WP_008310684_1 DUF63 -------------------------------------------------------MVLPSGLALPPLEYTVALLAGTLVVTALLYALEPPIDQRTVVALT 45
WP_058997315_1 DUF63 -------------------------------------------------------MVLPSGLALPPLEYTVALLAGTLVVTALLYALEPPIDQRTVVALT 45
WP_050037596_1 DUF63 -------------------------------------------------------MVLPSGLALPPLEYTVALLAGTLVVTALLYALEPPIDQRTVVALT 45
WP_050037596_1 DUF63 -------------------------------------------------------MVLPSGLALPPLEYTVALLAGTLVVTALLYALEPPIDQRTVVALT 45
WP_014039614_1 DUF63 -------------------------------------------------------MVLPSGLALPPLEYTVALLAGTLVVTALLYALEPPIDQRTVVALT 45
WP_064288256_1 DUF63 -------------------------------------------------------MVLPSGLALPPLEYTVALLAGTLVVTALLYALEPPIDQRTVVALT 45
WP_049994803_1 DUF63 ------------------------------------------------------MPLLPSGLVVPPLPYVIAVVVAAATVASFLVSLAPRVEQQHVFAFT 46
WP_020445351_1 DUF63 --------------------------------MNPSVTVGAAGSGSLWGPILQAGGLLPEGFGLPPLPYVGALLLGLITVGALLWALSPPVTNATVVAFA 68
WP_083761671_1 DUF63 -------------------------MFGASGQAA----AEPQSQRFPTTTTRTRVVVLPSDFALPALPYLAALALGGVVAVVLLYRRRPPVTERTVAALA 71
WP_084260786_1 DUF63 ----------------MKGIAFVAARVGRSGPAGRVARGRSGPQRFRPRRHRIEPVPLPADFALPPAPYLAALAVGAAVAVAALYRRRPRVTAASVTAFA 84
WP_014556864_1 DUF63 ------------------------------------------------------MVIFPDGFALPAIPYLLTIIFASGVVAYAAIRQRPTVTSSRVLALA 46
WP_070365684_1 DUF63 -------------------------------------------------------MLLPTGSTLPPLPYLVGLAAALLVVSAWLYRRDVALQTRTVLALA 45
WP_049951854_1 DUF63 -------------------------------------------------------MVLPEGFAVPPLELLVPLLCAFVAALALVWKTAPPVTDRTVLAFA 45
WP_015319722_1 DUF63 -------------------------------------------------------MVLPEGFALPPWQLLVPLVLAAVGVGAALWSLDPRVTDRTVLAFA 45
WP_005554260_1 DUF63 -------------------------------------------------------MVLPEGFALPPAQFLVPLVVAAVAVAAALWSADPPVTDRTVLAFV 45
AGB15442_1 putative  -------------------------------------------------------MVLPEGFAIPPWYFTVVLLLGVAGVGALLWAIDPPVTDRTVLAFT 45
WP_007698493_1 DUF63 -------------------------------------------------------MVLPEGFAIPPWYFTVVLLLGIAGVGALLWAIDPPVTDRTVLAFV 45
WP_086886713_1 DUF63 -------------------------------------------------------MVLPEGFVLPPWYLLGPTLVILGGVVAFLWSLDPPVTDRTVLAFT 45
WP_049926795_1 DUF63 -------------------------------------------------------MVLPEGFVLPPWYLLGPTVVILAGVVAFLWSLDPPVSDRTILAFA 45
WP_049922464_1 DUF63 -------------------------------------------------------MVLPEGFVLPPWYLLGPLLVILGGTIAFLWALEPQVTDRTVLAFA 45
WP_008008940_1 DUF63 -------------------------------------------------------MVVPEGFVVPPWYLLVPVVVILGGVIALLWALEPPVTDRTVLSFA 45
WP_098724536_1 DUF63 -------------------------------------------------------MVLPEGFVVPPWYLLVPVIVFLGSVLALLWALEPPVTDRTVLAFA 45
WP_097380374_1 DUF63 -------------------------------------------------------MVLPEGFVVPPWYLLVPVIVVLAGVIALLWTLEPPVTDRTVIAFA 45
WP_006429103_1 DUF63 -------------------------------------------------------MVLPEGFVVPPWYLLGPVIVVLGGVIALLWALEPPVTDRTVIAFA 45
WP_008457054_1 DUF63 -------------------------------------------------------MVLPEGFVVPPWYLLVPVIVILGSIVALLWALEPPVTDRTVIAFA 45
WP_008457054_1 DUF63 -------------------------------------------------------MVLPEGFVVPPWYLLVPVIVILGSIVALLWALEPPVTDRTVIAFA 45
WP_008457054_1 DUF63 -------------------------------------------------------MVLPEGFVVPPWYLLVPVIVILGSIVALLWALEPPVTDRTVIAFA 45
WP_007107690_1 DUF63 -------------------------------------------------------MVLPEGFVVPPWYLLVPVIVILGSIIALLWALEPPVTDRTVIAFA 45
WP_015299220_1 DUF63 -------------------------------------------------------MVLPEGFVVPPWYMVVPAAVILGGVIALLWAIEPPVTDRTVLAFA 45
WP_006649902_1 DUF63 -------------------------------------------------------MVLPEGFVVPPWYMVVPAAVILGGVIALLWAIEPPVTDRTVLAFA 45
WP_066299764_1 DUF63 -------------------------------------------------------MVLPEGFVVPPWYMVVPAVVILGGVIALLWAVEPPVTDRTVLAFA 45
WP_049965539_1 DUF63 -------------------------------------------------------MVLPEGFVVPPWYMVVPAAVILGGVIALLWAIEPPVTDRTVLAFA 45
WP_049989654_1 DUF63 -------------------------------------------------------MVLPEGFVVPPWYLLVPAVVILGGVVALLWAIDPPVTDRSVLAFA 45
WP_076147241_1 DUF63 -------------------------------------------------------MVLPEGFVVPPWYMVVPAVVILGGVIALLWAIEPPVTDRTVLAFA 45
WP_012941789_1 DUF63 -------------------------------------------------------MVLPEGFVLPPWYLLVPLLVILAGIVALLWVLEPPVTDRTVLAFA 45
WP_008895944_1 DUF63 -------------------------------------------------------MVLPEGFVLPPWYLLVPLLVILAGIVALLWVLEPPVTDRTVLAFA 45
WP_007142166_1 DUF63 -------------------------------------------------------MVLPEGFVLPPWYFLVAILVVLGGVLAMLWALEPPVTDRTVVAFV 45
WP_013879207_1 DUF63 -------------------------------------------------------MVLPEGFALPPWYLLVPLLVVLGGTLALLWALEPPVTDRTVISFA 45
WP_006088775_1 DUF63 -------------------------------------------------------MVLPEGFVLPPWYLLVPILVVLAGVIALLWALEPPVTDRTVLAFV 45
WP_076581448_1 DUF63 -------------------------------------------------------MVVPEGFVLPPWYLLVPLLVVLVGIVALLWALGPPVTDRTVLAFA 45
WP_006064996_1 DUF63 -------------------------------------------------------MVLPEGFVLPPWYLLVPLLVVLAGVVALLWALDPPVTDRTVLAFA 45
WP_049890149_1 DUF63 -------------------------------------------------------MVVPEGFVLPPWYLLVPILVVLAGIIALLWALEPPVTDRTVLAFA 45
WP_005580668_1 DUF63 -------------------------------------------------------MVLPEGFVLPPWYLLVPILAVLGGVLALLWELEPPVTDRTVVAFV 45
WP_087715216_1 DUF63 -------------------------------------------------------MVVPEGFVLPPWYLLVPLLVVLGGVLALLWTLEPPVTDRTVIAFA 45
WP_006664680_1 DUF63 -------------------------------------------------------MVLPEGFVLPPWYLLVPLLAVLGGVVALLWALDPPVTDRTVLALT 45
WP_006110228_1 DUF63 -------------------------------------------------------MVLPEGFVLPPWYLLVPLLAVLGGVVALLWALDPPVTDRTVLALT 45
WP_006826566_1 DUF63 -------------------------------------------------------MVLPEGFVLPPWYLLVPLLAVLGGVVALLWALDPPVTDRTVLALT 45
WP_004214806_1 DUF63 -------------------------------------------------------MVLPEGFVLPAWYLLVPLLAVLGGVVALLWTLGPPVTDRTVLAFA 45
WP_006653721_1 DUF63 -------------------------------------------------------MVLPEGFVLPAWYLLVPLLAVLGGVVALLWTLGPPVTDRTVLAFA 45
WP_006653721_1 DUF63 -------------------------------------------------------MVLPEGFVLPAWYLLVPLLAVLGGVVALLWTLGPPVTDRTVLAFA 45
WP_050049052_1 DUF63 -------------------------------------------------------MVLPAGSTVPPIPQTAALLAALGAVLWGLRRDGVRVTDHTILALS 45
WP_075938119_1 DUF63 -----------------------------------------------------MSPLLPAGTALPPLPYLLAVLLGLGAVGLGLRRRRPAVTPAVVGGFA 47
WP_077206978_1 DUF63 ----------------------------------------------------MLLQVLPEGLVVPALPYLFGLAVAGIAVGAVLSIRRPPVTQRLVVAFT 48
WP_049892606_1 DUF63 ------------------------------------------------------MAVLPSGFALPPLPHLLAVAALAAAVLVGLRRDPPTVDGRAVLALA 46
WP_009487551_1 DUF63 -------------------------------------------------------MVLPEGFALPPLPYLAAVAVAVVTVAWLLARESPPVDDLTILAFG 45
WP_059057568_1 DUF63 -------------------------------------------------------MVLPSGFAPPPLPYLVAVAAAVLAVGWLLVRESPPVTDRTVLAFA 45
WP_058984064_1 DUF63 -------------------------------------------------------MVLPSGFALPSLPYLAVVAVAVVAVGWLLVRESPPVTDRTVLAFG 45
EMA41075_1 hypotheti ---------------------------------------------------MQILQVLPEGSTLPPVAYLVVLVLALAGVALALRRVGPRVTDRVVVSFA 49
WP_049998124_1 DUF63 --------------------------------------------------MQVPAQLLPAGSALPPVGYLLVLGVVFVAVALSLRRLGPRVTDRVVVAFA 50
WP_049902415_1 DUF63 ---------------------------------------------------MVPVQLLPEGSTLPPPGVLLVLATALVGVVLALRRVAPQVTDQVVVAFG 49
WP_006079317_1 DUF63 ---------------------------------------------------MVPVQVLPEGTTLPSPGVLLALAIGFAGVVVALRRIAPRVTDRVVVAFA 49
WP_080509272_1 DUF63 MRAANSARVNSGGTTRPVGRRGGKAIARPRRRAPASATLSPPPASDQPVYDSLQLQVLPEGFALPPLGYLLVLAVALAAVGYVLRARDPPFGERHVLALV 100
WP_008000170_1 DUF63 --------------------------------------MAVALAALAALAAPATLPFLPAGTTLPPAPYLLAVLLVAGVVAVVLRRRPPTVTDRHVLALT 62
WP_049983070_1 DUF63 --------------------------------------------------MMGGLPLVPAGTTLPPLPYLIAVSIVVGAVGAGLGRRRPTVTDRHVLALA 50
WP_095637763_1 DUF63 --------------------------------------------------MAGGLPFLPAGTTIPPAPYLVGVLLAAGAVAVALRRRRPRVTGRRVLALA 50
WP_015910634_1 DUF63 --------------------------------------------------MVGDLPFLPAGTTLPPAPYLLVVLLATGGVVAALRRRRPRVTGRRVLALA 50
WP_008006417_1 DUF63 --------------------------------------------------MVGGLPFVPEGTTLPPVPYLLVVLLATAGVVAALRRRRPRVTGRRVLALA 50
WP_008583911_1 DUF63 ----------------------------------------------------MVGGLLPAGTTLPPLPHLLVVLLAAGGVAAALRRRRPEVTGRRVLALA 48
WP_006627969_1 DUF63 ----------------------------------------------------MVGGLLPAGTTLPPLPHLLVVLLAAGGVAAALRRRRPRVTGRSVLALA 48
WP_053772749_1 DUF63 ----------------------------------------------------MVGALLPAGTTLPPLPHLVVLLLATAGVVAALRRRRPAVTGRRVLALA 48
```

```
WP_004046581_1 DUF63 --------------------------------------------------MVGGLPFLPAGTTLPPAPYLLAVLLATGGVVAALRRRRPRVTGRRVLALA 50
WP_006112271_1 DUF63 ----------------------------------------------------MVGGLLPAGTTLPPLPHLLVVLLATAGVVAALRRRRPRVTGRRVLALA 48
WP_008443330_1 DUF63 ----------------------------------------------------MVGGLLPAGTTLPPLPHLLVVLLATGGVVAALRRRRPRVTARRVLALA 48
WP_008367944_1 DUF63 ----------------------------------------------------MVGGLLPAGTTLPPLPHLLVVLLATGAVVAALRRRRPRVTARRVLALA 48
WP_007345917_1 DUF63 ----------------------------------------------------MVGGLLPAGTTLPPAPHLLVVLLATGAVVAALRRRRPRVTARRVLALA 48
WP_004595209_1 DUF63 ----------------------------------------------------MVGGLLPAGTTLPPAPHLLVVLLATGAVVAALRRRRPRVTARRVLALA 48
WP_096395669_1 DUF63 ----------------------------------------------------MVGGLLPAGTTLPPLPHLLVVLLATGGVAAALRRRRPRVTARRVLALA 48
WP_017344445_1 DUF63 ----------------------------------------------------MVGGLLPAGTTLPPLPHLLVVLLATGGVAAALRRRRPRVTARRVLALA 48
ESS03942_1 putative  --------------------------------------------MTGFIAALGGLPFLPEGTTLPPVPYLVAVLLAGGGVAVAFRRRAPEVSAAHVLGLV 56
WP_066418621_1 DUF63 ------------------------------------------MATAPTPATLGDLPFLPAGTTLPPLPYLLVALVAGGGVAVALRRRRPRVAAAHVLALT 58
ESS10517_1 putative  -------------------------------------------------------MVLPEGFALPPLSYLVGLLAAVATVGYTLYRSRPQVGEEHVLGLA 45
ERH00845_1 putative  ------------------------------------------------------MALLPAGFTVPPWPYLVVLALAVGGVGYGLASRRPAVTARHVLGLA 46
ESS08053_1 putative  ------------------------------------------------------MALLPAGFTVPPWPYLVVLALAVGGVGYGLASRRPAVTARHVLGLA 46
WP_103427710_1 DUF63 ---------------------------------------------------MPLLQFLPEGFALPPLPYLIALAVGLGVVAVGATSRDLAVDRRHVVAFV 49
WP_049937371_1 DUF63 ------------------------------------------------------MAVLPEGFALPPLPYLVVLVAGLVAVGTLLRRRRPPVTDATIRAFV 46
WP_049987236_1 DUF63 ---------------------------------------------------MREYGDSPRRFALPPLPYLAALLVAAAVVGREAYRRAPAVTTPVVLAFA 49
WP_006053788_1 DUF63 ------------------------------------------------------MAILPEGFALPPLPYLVGLLVAVGVVVAAVARRRPEVTPQRILAFV 46
WP_008384832_1 DUF63 ------------------------------------------------------MAILPEGFSVPPLPYLAALLVAVGVVAAAAVRRRPAVTPRRVLAFV 46
WP_058582368_1 DUF63 ------------------------------------------------------MQVLPEGFALPPLPYLLALVVGLAAVGYAFYRRRPALTDRHVLALA 46
WP_004060105_1 DUF63 ------------------------------------------------------MAILPEGFALPPLPYLVVLVAGLVGVGVGLVRTRPTVSSAHVLALV 46
WP_008320401_1 DUF63 ------------------------------------------------------MAILPEGFALPPLPYLVVLVVGLVGVGVGLVRTRPTVSSAHVIALV 46
WP_008325536_1 DUF63 ------------------------------------------------------MAILPEGFALPPLPYLAVLVAGLVGVGVGLVRTRPPVSEGHILALV 46
WP_007543205_1 DUF63 ------------------------------------------------------MAILPEGFALPPLPYLAVLVAGLVGVGVGLVRTRPPVSEGHILALV 46
WP_004042703_1 DUF63 ------------------------------------------------------MAILPEGFALPPLPYLVALVAGLVGVGVGLTRTRPTVTPAHVLALV 46
WP_004062371_1 DUF63 ------------------------------------------------------MAILPEGFALPPLPYLVALVAGLVGVGVGLTRTRPTVAPAHVLALV 46
WP_008575916_1 DUF63 ------------------------------------------------------MAILPEGFALPPLPYLVALVAGLVGVGVGLRRTRPTVTPAHVLALV 46
WP_008575916_1 DUF63 ------------------------------------------------------MAILPEGFALPPLPYLVALVAGLVGVGVGLRRTRPTVTPAHVLALV 46
WP_008575916_1 DUF63 ------------------------------------------------------MAILPEGFALPPLPYLVALVAGLVGVGVGLRRTRPTVTPAHVLALV 46
WP_089777339_1 DUF63 ------------------------------------------------------MAILPEGFALPPLPYLVALAAGLVGVGVGLTRTRPTVTPAHVLALV 46
WP_007275294_1 DUF63 ------------------------------------------------------MAILPEGFALPPLPYLVALAAGLVGVGVGLTRTRPTVTPEHVLALV 46
WP_004969536_1 DUF63 ------------------------------------------------------MAILPEGFALPPLPYLVALVAGLVGVGVGLTRTRPTVTPAHVLALV 46
WP_050459703_1 DUF63 ------------------------------------------------------MAILPEGFALPPLPYLVALVAGLVGVGVGLSRTRPTVTPAHVAALV 46
WP_008095201_1 DUF63 ------------------------------------------------------MAILPEGFALPPLPYLVVLVAGLVGVGVGLSRTRPTVTPAHVAALV 46
WP_049967774_1 DUF63 ------------------------------------------------------MAILPEGFALPPLPYLVVLVAGLVGVGVGLSRTRPTVTPAHVAALV 46
WP_058569127_1 DUF63 ------------------------------------------------------MAILPEGFALPPLPYLVALVAGLVGVGVGLRRTRPTVTPAHVAALV 46
WP_058828475_1 DUF63 ------------------------------------------------------MAILPEGFALPPLPYFVALVAGLVGVGIGLRRTRPTVTPAHVAALV 46
WP_009376007_1 DUF63 ------------------------------------------------------MQILPEGFALPPLPYLLVLLVVGVAVAVGVRTRRPAVTGDHVVALA 46
WP_101294394_1 DUF63 ------------------------------------------------------MQLLPEGFALPPAPYLLGLLVGLGAVGWAFARRRPPVTDRHVVALA 46
WP_089671020_1 DUF63 ------------------------------------------------------MVLLPEGFSLPPLPYLGGLLIAVAAVGYGLSRRRPPVTARHILAFA 46
WP_096390099_1 DUF63 ------------------------------------------------------MVLLPEGFGLPPLPHLVGLLVAVALVAAGIARRRPTVSERHVVAFA 46
WP_021073253_1 DUF63 ------------------------------------------------------MVLLPEGFGLPPLPHLVGLLVAVALVAAGIARRRPAVSERHVVAFA 46
WP_007983568_1 DUF63 -------------------------------------------------------MVLPDGFAFPPISYLLPLLLAVGVVIWLLWRSEPTISERTVVSLA 45
WP_049969758_1 DUF63 -------------------------------------------------------MVLPEGFALPSLVYLLPLLLAVGSVAWALWRVKPTISERTVVALA 45
WP_066141551_1 DUF63 -------------------------------------------------------MVLPDGFALPPLLYLLPLLLAVGAVLWLLWEAEPTIGERTVVSLA 45
WP_008414921_1 DUF63 -------------------------------------------------------MVLPDGFSLPPLSYLLVLLVAAAAVATALWRRRPPVREATVLAFV 45
WP_066381248_1 DUF63 -------------------------------------------------------MVLPDGFSLPSIPYLLALLAASAAVAAALWRRRPPIRETTVLAFV 45
WP_014051708_1 DUF63 ------------------------------------------------------MAILPAGFALPPLPYLLVLLLAGAAVAYGLRERRPPVDDRFVRALV 46
WP_049980148_1 DUF63 -------------------------------------------------------MVLPAGFALPPLPYLLALVVAGGALAVALWRRDPAVSDRLVLALV 45
WP_053948907_1 DUF63 -------------------------------------------------------MVLPAGFALPPLPYLLGLLLAGGAVAVGLRRRDPPVSGRLVLALV 45
WP_079233625_1 DUF63 -------------------------------------------------------MVLPAGFALPPLPYLLGLLLAGGAVAAGLWRRDPPVSDRLVLALV 45
KPN32293_1 hypotheti -------------------------------------------------------MVLPAGFALPPLPYLLALLLAGGAVAVALWRRDPAVSDRLVLALV 45

WP_006882700_1 DUF63 PWMIAGGAFHAFEQ-----A------GLFPAVLTPFFAAPAVYVTTFLLAAAVWLPSV-IRAQM----------------VDDVGR---IARNLGVVGTV 115
WP_015790720_1 DUF63 PWMAIGGMAHGFYQ-----I--PQPPGLYPEWVGPLFGAPAVYVTTYVVVAAVWLLLI-LLGTA----------------IGKLDR---VVTYLGATGVG 119
WP_008524992_1 DUF63 PWMAVGGIAHGFYQ-----I--PNPPGLYPEWAGPLFGAPAVYLTTYVLVATVWLLLI-VLGTA----------------TNSLDR---VVTYLGATGLG 118
WP_015762485_1 DUF63 PWMAIGGAFHAFEQ-----L------ALYPPLYEPLFGTPAVYLTTLLLTGGVWLALQ---AVG--------------IVRGNEDT---VVRNLGLVGTA 115
WP_018257825_1 DUF63 PWMAIGGAFHAFEQ-----L------ALYPSLYEPLFGTPAVYLTTFLLTGGVWLALQ---AVG--------------IARGNEDT---VVRNLGLVGTA 115
WP_101349367_1 hypot PWMALGGALHAFHQ---PPI------DAYRPVVAPLFGAPAVYLTTFVTLALVWITLT-LFSVR----------------RGHSET---ISRNLGYIGVG 116
WP_004518630_1 DUF63 PWMALGGALHAFHQ---PPI------EAYRPVVAPLFGAPAVYLTTFVTLGVVWITLT-LFSVR----------------RGHSET---ISRNLGYIGTG 116
WP_004963804_1 DUF63 PWMALGGALHAFHQ---PPI------EAYRPVVTPLFGAPAVYFTTFVTLGVVWVTLT-LFSVR----------------RGHSET---ISRNLGYIGIG 116
WP_007189047_1 DUF63 PWMALGGALHAFHQ---PPI------EAYRPVVTPLFGAPAVYLTTFVTLGVVWVTLT-LFSVR----------------RGHSET---ISRNLGYIGIG 116
WP_004593738_1 DUF63 PWMALGGALHAFHQ---PPI------EAYRPVVAPLFGAPAVYLTTLVTLGVVWITLT-LFSVR----------------RGHSET---ISRNLGLIGTG 116
WP_005535867_1 DUF63 PWMALGGALHAFHQ---PPI------AAYRPVVAPLFGAPAVYLTTFVTLGVVWITLT-LFSVR----------------RGHSET---ISRNLGYIGIG 116
WP_053967006_1 DUF63 PWMALGGALHAFHQ---PPI------EAYRPVVAPLFGAPAVYLTTFVTLGVVWITLT-LFSVR----------------RGHSET---ISRNLGYIGIG 116
WP_008310684_1 DUF63 PWMALGGALHAFHQ---PPI------EAYRPVVAPLFGAPAVYLTTFVTLGVVWITLT-LFSVR----------------RGHSET---ISRNLGYIGIG 116
WP_058997315_1 DUF63 PWMALGGALHAFHQ---PPI------EAYRPVVAPLFGAPAVYLTTFVTLGVVWITLT-LFSVR----------------RGHSET---ISRNLGYIGIG 116
WP_050037596_1 DUF63 PWMALGGALHAFHQ---PPI------EAYRPVVAPLFGAPAVYLTTFVTLGVVWITLT-LFSVR----------------RGHSET---ISRNLGYIGIG 116
WP_050037596_1 DUF63 PWMALGGALHAFHQ---PPI------EAYRPVVAPLFGAPAVYLTTFVTLGVVWITLT-LFSVR----------------RGHSET---ISRNLGYIGIG 116
WP_014039614_1 DUF63 PWMALGGALHAFHQ---PPI------EAYRPVVAPLFGAPAVYLTTFVTLGVVWITLT-LFSVR----------------RGHSET---ISRNLGYIGIG 116
WP_064288256_1 DUF63 PWMALGGALHAFHQ---PPI------EAYRPVVAPLFGAPAVYLTTFVTLGVVWITLT-LFSVR----------------RGHSET---ISRNLGYIGIG 116
WP_049994803_1 DUF63 PWMAVGASAHVFYQ-LEPAV------DVYPAWAEPLFGAPMVYLTTFVALGAVWTFLA-FGSSV---------------NPDENDN---VALFLGLAGTA 120
WP_020445351_1 DUF63 PWMMTGGVLHALDQ-----V------REFPDYVEPLFGTPAVYATVALAAGTVWVITT--LYSA-----------------MTAGT---PDRHLGAIGTG 135
WP_083761671_1 DUF63 PWMAAGGTLYALYQ-----A------GAAPEPLAPLFSAPAVYVSVGVLAGGVWAGVA-DRDGS----------------TWSLTG---AAGPLAASGSV 140
WP_084260786_1 DUF63 PWVAAGGALYALHQ-----V------GVPRDSLDPLFGSPAVYVTVGILAGGVWATVA---DRP--------------AESWSASG---APALLAGAGSA 153
WP_014556864_1 DUF63 PWMALGAALHVQYV-----L------NALPPAIRPLAGTPAVYAIVTIAAIGTWV-SL---DLL-----------------VTSAY---IAQILAGVGAV 111
WP_070365684_1 DUF63 PWMVLGSALYVCYQ-----I------GAAPAALAPLCSSPTVYASTAVLAGLVWALAS------------------------RFEA---TDRWLAGTGVV 107
WP_049951854_1 DUF63 PWMVLGSTFYVLYQ-----V------AAVPEAVSPLFGAPVVYVTTATLAGFVWIGGL-SLHRA-----------------GFSQS---AETVLGAVGAV 113
WP_015319722_1 DUF63 PWMVVGSSLHVLHR-----L------GAYPEDVAVLFGTPSVYLVTGVVAGAVWIGGI-VLERT-----------------GVGIT---TEWPVAVLGGV 113
WP_005554260_1 DUF63 PWMVFGSSLHVLHR-----L------DAYPEDLAVLFGTPSVYVITGIAAGAVWLGGI-ALERT------------------GAAS---SESLVAAVGTL 112
AGB15442_1 putative  PWMMFGSTLYVLGR---DPI------SAFPEPIAPLFEAPSVYVTTAILAGFIWIVAN-ILHAG-----------------GLYRS---IPRFVGICGTA 115
WP_007698493_1 DUF63 PWMMFGSTLYVLGR---DPV------SAFPEPIAPLFEAPSVYVTTAILAGFVWIVAN-ILHAG-----------------GLYRS---IPRFVGICGTA 115
WP_086886713_1 DUF63 PWMMFGSTLHVLHR-----L------EAYPPEIAIMFATPSVYLVSAVVAGLAWIGGI-LLHAA-----------------GLQPT---IERFLGITGTG 113
WP_049926795_1 DUF63 PWMLFGSTLHVLHR-----L------GAFPDSIEILFATPSVYLVTAVIMGLAWIVGI-FLHAA-----------------GLQPT---NERVPLIAGSG 113
WP_049922464_1 DUF63 PWMLFGSTLHVLYR-----I------DAYPDTIAVLFSAPSVYLVTAVIGGLVWVLGL-LLHAA-----------------GLQPT----ERILGVTGTA 112
WP_008008940_1 DUF63 PWMMFGSSLHVLYK-----L------DAFPATIEPLFSAPMVYAVTAVVAGLAWIVAI-FLHVG-----------------GLQPT---VARFVGIAGTA 113
WP_098724536_1 DUF63 PWMMFGASLHVLYKINAFQI------NAFPASIESLFSAPIVYAVTAIIAGIVWIIAI-FLHVG-----------------GLQPT---ISRFVGIAGTA 118
WP_097380374_1 DUF63 PWMMFGATLHVLYK-----M------AAFPASIEPLFTAPMVYAVTAVLAGLVWIVAV-FLHVG-----------------GLQRT---THRFVGIAGTA 113
WP_006429103_1 DUF63 PWMMFGACLHVLYK-----L------GAFPATIEPLFTAPMVYAVTATVAGIVWIISI-FLHVG-----------------GLQQT---IPRFVGIAGTA 113
WP_008457054_1 DUF63 PWMMFGATLHVLYK-----L------AAFPPSIEPLFTAPMVYAVTAIVAGAVWIVAV-FLHVG-----------------GLQRT---IPRFVGIAGTA 113
WP_008457054_1 DUF63 PWMMFGATLHVLYK-----L------AAFPPSIEPLFTAPMVYAVTAIVAGAVWIVAV-FLHVG-----------------GLQRT---IPRFVGIAGTA 113
WP_008457054_1 DUF63 PWMMFGATLHVLYK-----L------AAFPPSIEPLFTAPMVYAVTAIVAGAVWIVAV-FLHVG-----------------GLQRT---IPRFVGIAGTA 113
WP_007107690_1 DUF63 PWMMFGATLHVLYK-----L------AAFPPSIEPLFTAPMVYAVTATVAGAVWIVAV-FLHVG-----------------GLQRT---IPRFVGIAGTA 113
WP_015299220_1 DUF63 PWMMFGATLHVLYK-----M------GAFPPTIEPLFTAPMVYVVTAIIAGLAWVVGV-FLHVG-----------------GLQPT---IPRFVGVAGTA 113
WP_006649902_1 DUF63 PWMMFGATLHVLYK-----M------GAFPSTIEPLFTAPMVYVVTAIIAGLAWVVGV-FLHVG-----------------GLQPT---IPRFVGVAGTA 113
WP_066299764_1 DUF63 PWMMFGATLHVLYK-----M------GAFPPTIEPLFTAPMVYVVTAIIAGLAWVVGV-FLHVG-----------------GLQPT---IPRFVGVAGTA 113
```

```
WP_049965539_1 DUF63 PWMMFGATLHVLYK-----M------GAFPSTIEPLFTAPMVYVVTAIIAGLAWVVGV-FLHVG-----------------GLQPT---IPRFVGVAGTA 113
WP_049989654_1 DUF63 PWMMFGAALHVLYK-----L------DAFPPAIEPLFSAPMVYVVTAIVAGIVWIVAV-FLHIG-----------------GLQPT---IARFVGVAGTA 113
WP_076147241_1 DUF63 PWMMFGATLHVLYK-----M------SAFPPTIEPLFTAPMVYVVTAIIAGLAWIVGV-FLHVG-----------------GLQPT---IPRFVGVAGTA 113
WP_012941789_1 DUF63 PWMMFGSALHVLYQ-----L------GAYPENVETLFATPTVYLVAAAVAGAAWIVAI-FLYAG-----------------GLQPT---IPRFFGIAGTA 113
WP_008895944_1 DUF63 PWMMFGSALHVLYQ-----L------DAYPENVETLFATPTVYLVAAAVAGAAWIVAV-FLYAG-----------------GLQPT---ISRFFGIAGTA 113
WP_007142166_1 DUF63 PWMMFGSALHVLHQ-----L------GAYPDSIAVLFTSPGVYLVTAIVGGLVWILGI-FLEAA-----------------GLQRS---IERFVGVTGTG 113
WP_013879207_1 DUF63 PWMMFGSTLQVLYR-----L------EAFPDSIAVLFTAPSVYLVTAIVAGLVWVVGI-FLQAG-----------------GLQPT---IERVVGVAGTG 113
WP_006088775_1 DUF63 PWMLFGSALHVLHR-----L------EAYPDQFETLFATPTVYLVTAIIAGGSWIIGI-FLYAG-----------------GLQST---IERFVGVAGTA 113
WP_076581448_1 DUF63 PWMMFGSTLHVLHR-----I------DAYPDNLESLFAAPSVYLVTAVIAGSAWIIGI-FLYAG-----------------GLQPT---IERFVGISGTA 113
WP_006064996_1 DUF63 PWMMFGATLHVLHR-----I------EAYPDHIEVLFASPSVYLVTAVVAGAAWIIAS-LLYAG-----------------GLQST---ISRFVGISGTA 113
WP_049890149_1 DUF63 PWMMFGSTLHVLHR-----L------DLYPDQIETLFAAPSVYLVTAIVAGAAWIIAI-FLYAG-----------------GLQPT---IARFVGISGTA 113
WP_005580668_1 DUF63 PWMLFGATLHVLYR-----T------GAYPETVEVLFTSPGVYLITAIVAGAAWILGI-FLQEG-----------------GLQPT---SERVVGITGTG 113
WP_087715216_1 DUF63 PWMMFGSTLHVLHR-----L------EAYPDSIEVLFASPSVYLLTATVAGLVWIVGI-FLYAG-----------------GLQSS---IERVVGIVGTG 113
WP_006664680_1 DUF63 PWMMFGATLHVLHR-----F------EAYPDSIEVLFATPTVYLITATGAGLVWILSI-FIYTG-----------------GLQPS---IERVLGTVGTG 113
WP_006110228_1 DUF63 PWMMFGATLHVLHR-----F------GAYPDSIEVLFATPTVYLITATGAGLVWVLSI-FIYTG-----------------GLQPT---IERVLGTVGTG 113
WP_006826566_1 DUF63 PWMMFGATLHVLHR-----F------EAYPDSIEVLFATPTVYLITATGAGLVWILSI-FIYTG-----------------GLQPT---IERVLGTVGTG 113
WP_004214806_1 DUF63 PWMLFGSTLHVLHR-----T------EAYPDSIEVLFAAPSVYLVTATVAGLVWVLSI-FLYSG-----------------GLQQT---IERPLGTVGTG 113
WP_006653721_1 DUF63 PWMLFGSTLHVLHR-----T------EAYPDSIEVLFAAPSVYLVTATVAGLVWVLSI-FLYSG-----------------GLQPT---IERPLGTVGTG 113
WP_006653721_1 DUF63 PWMLFGSTLHVLHR-----T------EAYPDSIEVLFAAPSVYLVTATVAGLVWVLSI-FLYSG-----------------GLQPT---IERPLGTVGTG 113
WP_050049052_1 DUF63 PWMVAGATVHVVYQ-----L------DVLPSAVAPLASSPAVYGTTAVVAGFVWLLAG------------------------RTTR---PLWWLAGVGVL 107
WP_075938119_1 DUF63 PWMATGAAGYALYQ-----A------EAVPDAVAPLFGNPTVYATTFVVAGALWLAVA---DRP--------------ADAWSLAG---VPGALALAGSA 116
WP_077206978_1 DUF63 PWMTAGAAVHVLYQ-----LQLIPGAQPLPPVIAPLFGAPTVYVTTFLVAGAIWA-GL---AVA-------------------ERA---VPPTLAATGGF 117
WP_049892606_1 DUF63 PWMVAGSSLYACHQ-----L------GLYPSAVAPVFGSPIVYATTAAAAGATWL-AV---RRF-------------------GAR---PQVLLAGVGAV 109
WP_009487551_1 DUF63 PWMALGSALYVCFQ-----L------RLYPESVAPFFGSPIVYATTFAVAGATWLAAR------------------------RTTR---PLAVLAGVGAA 107
WP_059057568_1 DUF63 PWMAFGSTLYVCFQ-----L------GVFPDAVAPFFGSPTVYASTFAAAGVTWLVAR------------------------RTAR---PLVALASVGGA 107
WP_058984064_1 DUF63 PWMVLGSTLYVCFQ-----L------GVFPDAIAPFFGSPIVYASTFAAAGATWLAAR------------------------RTAR---PLLALAATGGV 107
EMA41075_1 hypotheti PWMVVGSSCYVLYQ-----V-----HDTSSFLLKPFFSSPTVYVSVAVLAGAVWAASA---AAG------------LPADRWRPPS---VPSVVGLSGAV 121
WP_049998124_1 DUF63 PWMMLGSSLYVLYQ-----V------DGVPPILRPFFGSPTVYISVAAVAGGVWAVTA---AAE------------LPTDRWRPPS---VPGVVALLGGV 121
WP_049902415_1 DUF63 PWMVAGSSCYVLYQ-----V------EAVPRSVAPFFGSPTVYVSIAVLAGAVWAATA---AAG------------FPADRWRPPS---VPGIVALVGSA 120
WP_006079317_1 DUF63 PWMVAGSSCYVLYQ-----V------EAVPQSVAPFFSSPAVYVSIAVLAGAVWAATA---ATG------------LPADRWRLPS---VPGVVALVGSA 120
WP_080509272_1 DUF63 PWMCVGAGGHVLYV-----L------GALPTPLRPLFGTPSAYLTTAALAGGVWLAAL---SVG---------GDSVGSEAGHGSG---DARTLAVVGTV 174
WP_008000170_1 DUF63 PWVILGSAAHVLYV-----I------DALPPLVRPLAGSPTVYLTVGALAGASWLAADATDAEAADATNAEAAAAADGRGAERDPAGSGVPRALAAVGTL 151
WP_049983070_1 DUF63 PWMALGSATHVLYV-----V------DALPAVLAPFAGSPTVYLTVGSLAGAAWLASV---AIR-------------------PQR---VPTALAASGAC 114
WP_095637763_1 DUF63 PWMALGSAVHVLYV-----V------DALPPLLSPLAGSPTVYLTVGTLAGAAWLAAD---AVR-------------------PDR---VSESLALAGAL 114
WP_015910634_1 DUF63 PWMALGSAAHVLYV-----V------DALPPLLSPFAGSPTVYLTVGSLAGAAWLVAD----------------------ATRPDR---VPAVLAVAGTL 114
WP_008006417_1 DUF63 PWMALGSAAHVLYV-----V------GALPPLLAPFAGSPTVYLTVGSLAGAAWLTAD---AVR-------------------PDR---VPAALAASGAL 114
WP_008583911_1 DUF63 PWMALGSAAHVLYV-----V------DALPPLVAPFAGSPTVYLTVGAAAGAAWL-AV---DAA------------------RPDR---VAATLAAAGAL 112
WP_006627969_1 DUF63 PWMALGSAGHVLYV-----V------DALPPVLAPFAGSPTVYLTVGAVAGAAWL-AV---DAA------------------RPDR---VAAWLAAAGAL 112
WP_053772749_1 DUF63 PWMALGSAAHVLYV-----V------DALQPLLAPFAGSPAVYLTVGALAGAAWL-AA---DAA------------------RPDR---IAATIAAAGAL 112
WP_004046581_1 DUF63 PWMALGSAAHVLYV-----V------DALPPVLAPFAGSPTVYLTVGSLAGAAWL-AA---DAA------------------RPDR---VAETLAVAGAL 114
WP_006112271_1 DUF63 PWMALGSAAHVLYV-----V------DALPPLLSPFAGSPTVYLTVGSLAGAAWL-AV---DAA------------------SPDR---VPATLAAAGGL 112
WP_008443330_1 DUF63 PWMALGSAAHVLYV-----V------DALPPLLAPLAGSPTVYLTVGSLAGAAWLTAD---AAR-------------------PNR---VAAPLAAAGAL 112
WP_008367944_1 DUF63 PWMALGSAAHVLYV-----V------DALPSSLAPFAGSPTVYLTVGTLAGAAWL-AA---DAA------------------RPDR---VAATLAAAGVL 112
WP_007345917_1 DUF63 PWMALGSAAHVLYV-----V------DALPSSLAPFAGSPTVYLTVGTLAGAAWL-AA---DAA------------------RPDR---VAATLAAAGVL 112
WP_004595209_1 DUF63 PWMALGSAAHVLYV-----V------DALPSSLAPFAGSPTVYLTVGTLAGAAWL-AA---DAA------------------RPDR---VAATLAAAGVL 112
WP_096395669_1 DUF63 PWMALGSAAHVLYV-----V------DALPSSLAPFAGSPTVYLTVGVLAGAAWL-AA---DAA------------------RPDR---VAATLAAAGAL 112
WP_017344445_1 DUF63 PWMALGSAAHVLYV-----V------DALPSSLAPFAGSPTVYLTVGVLAGAAWLAADAWLAVD----------------ADRPDR---IAATLAAAGVL 118
ESS03942_1 putative  PWIALGAAFHVLYV-----V------GGLPRPLAPLGGSPTVYLATGTLAGGVWLGAE-AVDGG--------------------DR---TPRLLAAAGTS 121
WP_066418621_1 DUF63 PWIALGSALHVLYV-----V------GTLPGVLAPFGGSPTVYLTVATLAGATWLVAD---AVA-----------------DDAGR---VPALLAAAGTL 124
ESS10517_1 putative  PWIAVGSCLHVHYV-----I------GSLPAALEPLAGTPAVYLTVAALAGGVWL-GI---DRA-------------VTSSPDSRA---VGVALFATGTA 114
ERH00845_1 putative  PWIVVGSALHVLYA-----V------ESLPAVIAPLAGTGAVYVTVAAIAGGGWLLAD---AVR-------------------PRT---VPALLAASGLA 110
ESS08053_1 putative  PWIVVGSALHVLYV-----V------ESLPPVIAPLAGTGAVYVTVAAIAGGGWLLAD---AVR-------------------PRT---VPALLAASGLA 110
WP_103427710_1 DUF63 PWILAGSCAHALYV-----V------DALPALLRPLGGTPAVYATVAVGAGLTWL-AA---DAA-----------------VDAET---TPQVLGYVGLA 114
WP_049937371_1 DUF63 PWMVLGSALHVLYV-----L------GALPSVVRPFAGTPAVYLSVAVVAGAAWL-AV-----------------------GADAG---TARRLRRVGIL 108
WP_049987236_1 DUF63 PWMVLGSALHVLYI-----E------DALPALARPLAGTPAVYVSVAVLAVAIWT-AA---DAL-----------------SDPDR---VAPTLGLVGGL 114
WP_006053788_1 DUF63 PWMILGSAVHVLYV-----V------NALPPVIHPLGGTPAVYLSVGVLGIGTFVAAA----------------------EWAEEA---LVPLLTGVGTV 110
WP_008384832_1 DUF63 PWMVFGSAAHVLYV-----V------DALPPLLRPLGGTAAVYLAVGALGVGTFV-AA---DAR------------------AGDA---VALVLAGLGSL 110
WP_058582368_1 DUF63 PWMVTGSALHVAYA-----L------GVLPAVVDPLMGTPSVYLTVGVVAGGVWV-VA---DAL-------------------NWP---VPTALAVTGVV 109
WP_004060105_1 DUF63 PWMVTGSVLHVLYV-----V------GLLPPVVEPLAGTPSVYVTVAIVAGATWL-VL---DAA-------------------SAA---SPRALAGPGLV 109
WP_008320401_1 DUF63 PWMVTGSVLHVLYV-----V------GLLPPAVEPLAGTPSVYLTVALVAGATWV-AL---DAA-------------------DAA---SPRALAGPGVV 109
WP_008325536_1 DUF63 PWMVSGSALHVFYV-----L------DFLPPALRPLAGTPAVYATVAVLAGATWL-GL---HVG-------------------DVD---TPRNLAVPGVA 109
WP_007543205_1 DUF63 PWMVSGSALHVFYV-----L------DFLPPALRPLAGTPAVYATVAVLAGATWL-GL---HVG-------------------DVD---TPRNLAVPGLA 109
WP_004042703_1 DUF63 PWMVSGSALHVLYV-----V------DALPPAVEPLAGTPAVYLSVAVVAGATWL-AL---DAA-------------------DLA---SPQTLAAPGLL 109
WP_004062371_1 DUF63 PWMVSGSALHVLYV-----V------GALPPAVEPLAGTPAVYLSVAVVAGATWL-AL---DAA-------------------DLA---SPQTLAAPGLL 109
WP_008575916_1 DUF63 PWMVSGSALHVLYV-----V------GALPPAVEPLAGTPAVYLSVAVVAGATWL-AL---DAA-------------------DLA---SPQTLAAPGLL 109
WP_008575916_1 DUF63 PWMVSGSALHVLYV-----V------GALPPAVEPLAGTPAVYLSVAVVAGATWL-AL---DAA-------------------DLA---SPQTLAAPGLL 109
WP_008575916_1 DUF63 PWMVSGSALHVLYV-----V------GALPPAVEPLAGTPAVYLSVAVVAGATWL-AL---DAA-------------------DLA---SPQTLAAPGLL 109
WP_089777339_1 DUF63 PWMVSGSALHVLYV-----V------DALPPAVEPLAGTPSVYLTVAVVAGATWL-AL---DAT-------------------DLA---SPTALAAPGAL 109
WP_007275294_1 DUF63 PWMVAGSALHVLYV-----V------DALPPAVEPLAGTPAVYLSVAVVAGATWL-AL---DAT-------------------DLA---SPPALAAPGAL 109
WP_004969536_1 DUF63 PWMVAGSALHVLYV-----V------DALPPAVEPLAGTPAVYLSVAVVAGASWL-AL---DAA-------------------DLD---SPTTLAAPGLL 109
WP_050459703_1 DUF63 PWMVSGSALHVLYV-----I------DALPPAVEPLAGTPSVYLSVAVAAGATWL-AL---DAA-------------------DLA---SPTALAVPGLL 109
WP_008095201_1 DUF63 PWMVSGSALHVLYV-----I------DALPPAVEPLAGTPAVYLSVAVAAGATWL-AL---DAA-------------------DLA---SPTALAVPGLL 109
WP_049967774_1 DUF63 PWMVSGSALHVLYV-----I------DALPPAVEPLAGTPAVYLSVAVAAGATWL-AL---DAA-------------------DLA---SPTALAVPGLL 109
WP_058569127_1 DUF63 PWMVSGSALHVLYV-----I------DALPPAVEPLAGTPAVYLSVAVVAGASWL-AL---DAA-------------------DLD---SPTALAVPGAL 109
WP_058828475_1 DUF63 PWMVSGSALHVLYV-----I------DALPPAVEPLAGTPAVYLSVAVVAGATWL-AL---DAT-------------------DLD---SPTALAIPGAL 109
WP_009376007_1 DUF63 PWMAVGSVLHVLHV-----V------DALPSILDPLAGTPSVYITVALLVGVVWL-AA---DAL-----------------TDAET---VPVVLAVTGLL 111
WP_101294394_1 DUF63 PWMCTGACLHVLYV-----L------DALPPAVRPLAGTPAVYLTVALVAGAIWI-GA---DEG-------------------TGR---APRALAVTGAF 109
WP_089671020_1 DUF63 PWMVVGSSLHVLHV-----I------GGLPPAVEPFGGTAAVYLTVSAVAGGSWLLAA---STR------------------PRRQ---VPAVIAGTGLL 111
WP_096390099_1 DUF63 PWMAVGSCLHVLYV-----M------GGLPPVVRPLAGTPAVYLSVAAMAGATWLAVD---AAA-----ASLAADDVDAETDAPDR---APTAIAAVGVL 124
WP_021073253_1 DUF63 PWMAVGSCLHVLYV-----M------GGLPPAVRPLAGTPAVYLSVAAIAGATWL-AI---DAGAAALAADAEPDADDADRVATNR---VPKAIAAIGVL 128
WP_007983568_1 DUF63 PWMVVGAGLNALYQ-----L----PESPIPDVFLPFFGTPAVYLTTFAVAGAVWLLSI---HRL-------------------RAP---VPRVLAGTGLV 111
WP_049969758_1 DUF63 PWMVVGAGLHALYQ-----V------RFVPESLAPLLGTPSVYLTTFVVAGVVWLFSL---RRR-------------------KNS---VPQTLAGVGTA 109
WP_066141551_1 DUF63 PWMVVGAGFNALHQ-----L----PNSPVPDVIDPLFGTPSVYLTTFVLAGIVWLLAI---HQL-------------------RTP---VPRALASGGLV 111
WP_008414921_1 DUF63 PWMVAGAACHVLEV-----I------EAVPPAFAPLLGTPSVYVSTFVLAGAIWTVTD--------------------------------DRFLALAGTL 102
WP_066381248_1 DUF63 PWMVAGAAAHVLEV-----I------DAVPPTLAPLFGTPSVYVSTFVLAGAIWAVTD--------------------------------DRILATAGTL 102
WP_014051708_1 DUF63 PWMLVGAWLHVLHI-----V------GAAPAAIDPLLGTPAAYISTAVLAGGVWL-GA---DAS-------------------ERD---TARLLLGVGVA 109
WP_049980148_1 DUF63 LWMLAGAWLHVLHV-----V------GAAPGAIDPLLGTPAAYLTTAILAGAVWL-AV---DAG-------------------AVA---TERAFAAVGAV 108
WP_053948907_1 DUF63 PWMLAGAWLHVLDV-----V------GAVPDAVAPLFGTPAAYVTTAVLAGAVWLAIE------------------------ADDG---NERPFAAAGTG 107
WP_079233625_1 DUF63 PWMLAGAWLHVLDV-----V------GAVPDAVAPLLGTPAAYVTTAILAGTVWLAVN------------------------ADDG---NERPFAATGGV 107
KPN32293_1 hypotheti PWMLAGAWLHVLHV-----V------GAAPEAIDPLLGTPAAYLTTAILAGAVWL-AV---DAG-------------------EIA---TERAFAAAGAA 108

WP_006882700_1 DUF63 LVLALTVVAYAFAA-GRVDSTTALVWLPVGFLGAGVLTAPVYYLLALKWTGAVDRAGAAGPLVVYAHALDGFSTAVGVDVIGTGERTPIPRRIMDFAETL 214
WP_015790720_1 DUF63 VLIVFLGAMVWQGI-GPG-LSFTPLWPTVAFLISLALTAVIYFLVSLRWTGVVARTGIAGAVVLFAHTFDGVITMIGKDVLEGGERTPIPAMIMDVAGSL 217
WP_008524992_1 DUF63 VLLVFLGVIIWQGM-GPE-LSFQPLWPTVAFLVALVLTATTYFLVSLKWTGEVARTGVAGVAVVFAHAFDGVITTVGKDVLGAHERTPIPATIMDFAGQL 216
WP_015762485_1 DUF63 IFVVLVVVRIWQSI-AQP--TFSPVWPAVSLIVALAVTALVVLAISLWRTPVFVRTRYAGPVVVFAHALDGVSTAIGADVVGASEQSPLPRLIMEFAGQL 212
```

```
WP_018257825_1 DUF63 IFVVLVVVRIWQSI-AQP--TFSPVWPAVSLIVALAVTALVVLAISLWRTPVFVRTRYAGPVVVFAHALDGVSTAIGADVVGASEQSPLPRLIMEFAGQL 212
WP_101349367_1 hypot FLTVLLVIAVVMAL-ESG--LLGLIWPTVAVVVATVVTAVTVLAVALWRTPVVVRMRYAAPTVVFAHMLDGVSTAVGADVIGITERTPIPARIMEFAGTL 213
WP_004518630_1 DUF63 LLTVLLVIGVVMAL-RSE--LLALIWPTVAVVVAIAVTAVTVLAVALWRTPVVVRMRYAAPTVVFAHMLDGVSTAVGADVIGITERTPIPARIMEFAGTL 213
WP_004963804_1 DUF63 LLTVLLVIAVVMAL-ESG--LLGLIWPTIAVIVATVVTAVTVLAVALWRTPVVVRMRYAAPTVVFAHMLDGVSTAVGADVIGITERTPIPARIMEFAGTL 213
WP_007189047_1 DUF63 LLTVLLVIAVVMAL-ESG--LLGLIWPTIAVIVATVVTAVTVLAVALWRTPVVVRMRYAAPTVVFAHMLDGVSTAVGADVIGITERTPIPARIMEFAGTL 213
WP_004593738_1 DUF63 LLTVLLVIAVVMAL-ESG--LLGLIWPTVAVVVATVVTAATVLAVALWRTPVVVRMRYAAPTVVFAHMLDGVSTAVGADVIGITERTPIPARIMEFAGTL 213
WP_005535867_1 DUF63 LLTVLLVIAVVMAL-ESG--LLGLIWPTVAVVVAAVVTAVTVLAVALWRTPVVVRMRYAAPTVVFAHMLDGVSTAVGADVIGITERTPIPARIMEFAGTL 213
WP_053967006_1 DUF63 LLTVLLVIAVVMAL-ESG--LLGLIWPTVAVVVATVVTAVTVLAVALWRTPVVVRMRYAAPTVVFAHMLDGVSTAVGADVIGITERTPIPARIMEFAGTL 213
WP_008310684_1 DUF63 LLTVLLVIAVVMAL-ESG--LLGLIWPTVAVVVAIVVTAVTVLAVALWRTPVVVRMRYAAPTVVFAHMLDGVSTAVGADVIGITERTPIPARIMEFADTL 213
WP_058997315_1 DUF63 LLTVLLVIAVVMAL-ESG--LLGLIWPTVAVVVAIVVTAVTVLAVALWRTPVVVRMRYAAPTVVFAHMLDGVSTAVGADVIGITERTPIPARIMEFAGTL 213
WP_050037596_1 DUF63 LLTVLLVIAVVMAL-ESG--LLGLIWPTVAVVVATVVTAVTVLAVALWRTPVVVRMRYAAPTVVFAHMLDGVSTAVGADVIGITERTPIPARIMEFAGTL 213
WP_050037596_1 DUF63 LLTVLLVIAVVMAL-ESG--LLGLIWPTVAVVVATVVTAVTVLAVALWRTPVVVRMRYAAPTVVFAHMLDGVSTAVGADVIGITERTPIPARIMEFAGTL 213
WP_014039614_1 DUF63 LLTVLLVIAVVMAL-ESG--LLGLIWPTVAVVVATVVTAVTVLAVALWRTPVVVRMRYAAPTVVFAHMLDGVSTAVGADVIGITERTPIPARIMELAGTL 213
WP_064288256_1 DUF63 LLTVLLVIAVVVAL-ESG--LLGLIWPTVAVVVATVVTAVTVLAVALWRTPVVVRMRYAAPTVVFAHMLDGVSTAVGADVIGITERTPIPARIMEFAGTL 213
WP_049994803_1 DUF63 VLLGLLGVAAWREA-FVG---VEPFWSAAALVITVPVTIGVYFTIAYTATGVVARAKLLGGLVIFAHALDGITTAVGIDILETSERSPLPRTIMDLAGQL 216
WP_020445351_1 DUF63 TAVTFAIFGLIEAF-NAG--TLDTLPAAITIAVTGVVAAITWAIIGVRFTDSAALASWSGVVVVVAHALDGVSTAVGYDMLDVSERSPVSEAVLEAGAAL 232
WP_083761671_1 DUF63 LLAAALVAAALAETPDTM--GTSAAGSAVILAASIAATAVAWVLLSRTA--EVSATGAVGVLAVFGHTLDGISTAVGYDLLGFGEQTPLSRIILEAGAAL 236
WP_084260786_1 DUF63 LLLAALAAAAFGPTPAGTGGGGTPVASVAIAVVSVVVAATAWGVLRRVR--DVDATGSVGSLAVFGHTLDGVSTAVGYDLLGFGEQTPLSRLIIDAGAAL 251
WP_014556864_1 DUF63 AALIAIAGVIAI---NPT--SLRVTVSIIGIGVSVILSGGVWVMLTRIVP-KTQLTAPMGTFAVFGHTLDGVSTTIGIDLLGFGERTPLSALIIEFASGL 205
WP_070365684_1 DUF63 LSLFPIGVAIAVAL-DAE--SFSPGWSILGAVLAGLLALLIWYVLGQYQPETTAILGPAGGVAVFAHVLDGISTTIGIDVLQFGEQTPLSAALIELGAAL 204
WP_049951854_1 DUF63 CFVGAAGYALRTGL-ETG--TLEAFWPGVSVILAAILTALAWVGVSQWASETAALTGTTGILVVFGHALDGVSTAVGYSVLGASESVPASQLILAAGKAL 210
WP_015319722_1 DUF63 GLVGAAATAVGAGG-GFD----GVFWPLLSVVLTVLVTAAVWIAFGRRYEDVVAVTGATGLLVVFGHTLDGISTAIGFDVLGASEEVPLSLLILEAGHAL 208
WP_005554260_1 DUF63 GFVGAAATAVSVQG-GFG----NVFWPLLSVVLTVLVTAGVWAALGRRYERVAAVTGATGALVVFGHTLDGISTAIGFDVLGASEEVPLSLLILEAGHAL 207
AGB15442_1 putative  FLSVFATAAIMTGI-ESG--TFAPFWPVVAVVITGIVTAIGWLALSLWATEVAAITGLTGALVCFAQVLDGTSTAIGYDVLGAGEQVPLSTAVLELAGSL 212
WP_007698493_1 DUF63 FLSVFATAAIMTGV-EGG--TFTPFWPVVAVVITGIVTAIGWFALSLWATEVAAVTGLTGALVCFGQVLDGTSTAIGYDVLGAGEQVPLSTAILGLAGDL 212
WP_086886713_1 DUF63 FFVVFAMFAIVTSL-EVG--TFQPFWPVIATVVAGIVTALAWLALGLWFTDVAATTGVTGAIVVFGHALDGVTTAVGYDVLGAHEEVPLSRLVLETAGGL 210
WP_049926795_1 DUF63 FFVVFAGFATVTGL-NGG--TFQPFWPVISVVVAGVVTALAWLALGLSFTEVAATTGVTGAIVVFGHALDGVSTAVGYDVMRAGEEVPLSQIVLDTAADL 210
WP_049922464_1 DUF63 FFAVFAMFAIISSW-EQG--TFQPFWPVIAVVVSGVVTSLAWVALSLWFTDVAATTGVTGAIVVFGHVLDGVSTAIGFDVLGGGEDVPLSQLILEAGRSL 209
WP_008008940_1 DUF63 FFTVFAMVAIFQSI-ELG--TFEPFWPVVSVIVAGILTAIAWLVLGLWFTDVAATTGMTGALVVFGHTLDGVSTAVGYDVLGASENVPLSLLILEAGKAL 210
WP_098724536_1 DUF63 FFTVFAMFAVLRSI-EAG--VFEPFWPVIAVVVAGIVTAIAWIALSLWFTDVATTTSTTGALVVFGHTLDGVTTAIGYDVLGAAESVPLSLVLLEAGEAL 215
WP_097380374_1 DUF63 FFSVFAMFALIRSF-EDG--TFEPFWPVISVVVAGIVTAITWVAVSLWFTDIAATTSATGALVVFGHTLDGVTTAVGYDVLAAGETVPVSLLLLEAGAAL 210
WP_006429103_1 DUF63 FFTVFTMFALSVGL-EDG--TLEPFWPVIAVVVAGVATAVAWIALSLWVTDIATTTSATGALVVFGHTLDGVTTAIGYDVLAAGEDVPLSLLILEAGEAL 210
WP_008457054_1 DUF63 FFTVFATGALLLSL-ETG--TIAPFWPVIGVVAAGIVTAVVWLMVSLWVTDIAATTGATGVLVVFGHALDGVTTAIGYDVLTAGEDVPLSLLILEAGEAL 210
WP_008457054_1 DUF63 FFTVFATGALLLSL-ETG--TIAPFWPVIGVVAAGIVTAVVWLMVSLWVTDIAATTGATGVLVVFGHALDGVTTAIGYDVLTAGEDVPLSLLILEAGEAL 210
WP_008457054_1 DUF63 FFTVFATGALLLSL-ETG--TIAPFWPVIGVVAAGIVTAVVWLMVSLWVTDIAATTGATGVLVVFGHALDGVTTAIGYDVLTAGEDVPLSLLILEAGEAL 210
WP_007107690_1 DUF63 FFTVFATGALLLGL-ETG--TIAPFWPVIGVVAAGIVTAVVWLVVSLWVTDIAATTGATGVLVVFGHALDGVTTAIGYDVLAAGEDVPLSLLILEAGEAL 210
WP_015299220_1 DUF63 FFTVFTMFALFRSI-QTG--TFEPFWPVITVVIAGIVTALAWLALSLWFTDVAATTGTTGALVVFAHTLDGVTTAVGYDVLGAGEDVPLSLLLLEAGEAL 210
WP_006649902_1 DUF63 FFTVFTMFALFRSI-QTG--TFDPFWPVISVVIAGIVTALAWLVLSLWFTDVAATTGTTGALVVFAHTLDGVTTAVGYDVLGAGENVPLSLLLLEAGEAL 210
WP_066299764_1 DUF63 FFTVFTMFALFRSI-QMG--TLDPFWPVITVVVAGIVTAVAWLGLSLWFTDVAATTGTTGALVVFAHTLDGVTTAVGYDVLGAGENVPLSLLLLEAGEAL 210
WP_049965539_1 DUF63 FFTVFTMFALFRSI-QTG--TFDPFWPVITVVIAGIVTALAWLVLSLWFTDVAATTGTTGALVVFAHTLDGVTTAVGYDVLGAGENVPLSLLLLEAGEAL 210
WP_049989654_1 DUF63 FFTVFAMVAILQSI-DAG--TFDPFWPVIAVVVTGIVTTIAWIALSLWFTDVAATTGATGVLVVFAHTLDGVTTAIGYDVLAVGEDVPLSLLILEAGESL 210
WP_076147241_1 DUF63 FFTVFTMFALFQSI-QTG--TFEPFWPVITVVVAGIVTALAWLALSLWFTDVAATTSTTGALVVFAHTLDGVTTAVGYDVLGAGENVPLSLLLLEAGEAL 210
WP_012941789_1 DUF63 FFAVFASIILNVSW-AEE--TFNLFWPVIAVIAAGIVTAIAWVALSLWFTDVAKTTSYTGALVVFGHTLDGVSTAIGYDVIGVEESVPLSALLLEAGGSL 210
WP_008895944_1 DUF63 FFAVFASIILNVSW-SEE--TFNLFWPVIAVIAAGIVTAIAWVALSLWVTDIAKTTSYTGALVVFGHTLDGVSTAVGYDVIGVEETVPLSALLLEAGESL 210
WP_007142166_1 DUF63 FFVVFAMFALQLGW-AAG--TFEPFWPVIAVVLSGIVTALAWLALGIWFTDVAAITGMTGVVVVFGHVLDGVSTAVGYDVLGAGENVPLSQLLLEAGEAL 210
WP_013879207_1 DUF63 FFVVFAMFIVLSGW-NAG--SFSPFWPVIAVVVAGIVTALAWVALGLWFTDVAATTGATGALVVFGQTIDGVSTAVGYAVLGSSEEVPLSRAILEVGKSL 210
WP_006088775_1 DUF63 FFAVFTMVILNIGW-EEG--TFELFWPVISVVIAGIVTAVAWIALSLWFTDVAATTSATGALVVFGHTLDGVSTAIGYDVLGAAEDVPLSRLILEAGESL 210
WP_076581448_1 DUF63 FFAVFVMIFLNIGW-AVG--NFELFWPVISVVIAGAVTAIAWIALSLRYTDVAATTSITGVIVVFGHALDGVSTAIGYDVLGAAEEVPLSRLILEAGESL 210
WP_006064996_1 DUF63 FFAVFAMIAFNIGW-ERG--TIDPFWPVIAVVVTGIVTAIAWVALSLSFTDVAATTGLTGALVVFGHTLDGVSTAIGYDILGASEEVPLSRLLLEAGEAL 210
WP_049890149_1 DUF63 FFAVFAMITLNVGW-EQG--TFEPFWPVIAVVATGVVTALAWVVLSLRFTDVAATTGATGVLVVFGHTLDGVSTAIGYDVLGAAEEVPLSRLILEAGETL 210
WP_005580668_1 DUF63 FFAVFAMLAIGRSW-GAG--TFEPFWPVIAVVASGIVTALAWLALGLWFTDVAATTGATGALVVFGHTLDGISTAIGYDVLGAGEEVPLSRMILETGSAL 210
WP_087715216_1 DUF63 FFAVFAMIAVVDGW-EAG--TFEPFFPVISVVIAGIVTALAWLALGLWFTDVAATTSLTGVLVVFGHALDGVSTAIGFDLLGAHEEVPVSRLILEAGEAL 210
WP_006664680_1 DUF63 FFVVFTMFALLQSL-ESG--TFEPFWSVIAVVAAGIATAVAWLVLGLWFTDVAAVTATTGVLVIFGHALDAVSTAIGYDVLNAGEEVPLSALILTVGDEL 210
WP_006110228_1 DUF63 FFVVFTMFAILQSL-ESG--TFEPFWSVIAVVAAGIATAIAWLVLGLWFTDVAAVTATTGVLVIFGHALDAVSTAIGYDVLNAGEEVPLSALILNAGSAL 210
WP_006826566_1 DUF63 FFVVFTMFALLQSI-ESG--TFEPFWSVIAVVAAGIATAIAWLILGLWFTDVAAVTATTGVLVIFGHALDAVSTAIGYDVLNAGEEVPLSALILNAGSAL 210
WP_004214806_1 DUF63 FFAVFTMFLLFESV-ELG--TFEPFWPVIAVVISGVVTALAWLIIGLWFTDVAAVTGATGVLVVFGHVLDGVSTAIGYDILGAGEEVPLSELILDAGHAL 210
WP_006653721_1 DUF63 FFAVFTMFLLFESV-ELG--TFEPFWPVIAVVISGVVTALAWLVLGLWFTDVAAVTGATGVLVVFGHALDGVSTAIGYDILGAGEEVPLSELILDAGHAL 210
WP_006653721_1 DUF63 FFAVFTMFLLFESV-ELG--TFEPFWPVIAVVISGVVTALAWLVLGLWFTDVAAVTGATGVLVVFGHALDGVSTAIGYDILGAGEEVPLSELILDAGHAL 210
WP_050049052_1 DUF63 AVLVPTGVALAVAA-AGG--TLTLTVPLLGVAAGAVMAWVTWRLFGAVRPSDATTVGVAGPVVLFGHALDGATTTLGVDVLGFGEKTPLSRIVMEIAGAL 204
WP_075938119_1 DUF63 CFAVLVGVGIRHAL-TTG--AVTPAGPVRALALSVVVTGGLWLAVRSW----VGVTGLPGVLALFGHTLDGISTAQGI-ALGFGEQTPLSAALIEAGHGL 208
WP_077206978_1 DUF63 ATLAGAALVWRTAT-VQG--TFEPVVPLAALLASVVVAVATYALLDRTRPTVTARTGTLGAVTVFGHALDGVSTAVGVDLLGSGERSPIPAAIMEFAGSL 214
WP_049892606_1 DUF63 AAAVPITAALARGW-TAG--TLTLAWPLGAVGAALVLTAGSWRLLARIRPTDTRAVGVAGVLAVFGHALDATSTAVGVDALGYGEQTPLSTAIIELARAL 206
WP_009487551_1 DUF63 LALVPAAAAINFGL-EHD--SLDLAWPLAAVVAAAVLGHVTWFAVRSWRPEAARLTGVAGALAVFGHVLDGTSTAVGIDVLGFGEQTPLSALVMEFAAHL 204
WP_059057568_1 DUF63 LTVFPAAAAVNYGL-ANG--TLTLAWPLAAVAAAAVLGRVAWWVVERLRPTDAAAVGGAGALAVFAHVLDGTSTAVGVDVLGFGEQTPLSAAIMHFAADL 204
WP_058984064_1 DUF63 LTLAPTAAAVGYAL-AHD--TLTLAWPLAAVVAAAVLGHVAWWVVEQLRPDDVAAVGGAGALAVFAHVLDGTSTAVGVDVLGFGEQTPLSAAIMHAAAEL 204
EMA41075_1 hypotheti LAVVVVGWALAMS--APD---LEPVWPGVALAVAVVCAAAVWVAIRRLVP-ATRVTGAVGGLCVLGHALDGVSTAVGIDVLGFGERSPFSRAIIEFAATL 215
WP_049998124_1 DUF63 LAVVAVGWALSVG--APG---LTVAWPAVGAVVATVLAAVVWGGLRRAVP-KTECAGAVGALAVFGHALDGVSTAVGIDVLGFGERSPVSRAVIEFAAGL 215
WP_049902415_1 DUF63 AAIAAVGWAFAFGVETRQ--GLTVAWPSVGLVIAVVLAAIVWSGLRRVAP-ETRITGGVGALAVFGHALDGVSTAVGIDVLGFGEQSPISRAIITFAADL 217
WP_006079317_1 DUF63 VAIAAVGWAFAFGVETSQ--GLTVAWPAVGLVVAVAVATVVWLLLGRLAP-ETRITGGVGALAVFGHALDGVSTAVGIDVLGFGEQSPISRAIITFAADL 217
WP_080509272_1 DUF63 ALAVVLAGVAAYGA-GRG--TFAPALPGLGLLGGAALGVVAARLLFAAYD-PAAVTGTAGALAVAAHGVDGLTTAVGVDLLGYGERTPLSRLIIEFAAGL 270
WP_008000170_1 DUF63 ALVPVVAVAVAAGI-STD----GAFWSAVSLAAAVPIAALVWTGLTRARP-ETGITGGVGALAVFGHALDGASTAVGTTQLGFGERTPLSRLILEIEG-L 244
WP_049983070_1 DUF63 LLAPVVFVALATGL-SPD----GAFWSGVSLALTLPIAGVAWVGLIRLRP-EAAVTGGVGALAVFAHALDGVSTAIGTTQLGFGERTPVSRLLLELGG-L 207
WP_095637763_1 DUF63 CLAPVVAVAVSTGL-SSE----GARWSTIALALTVPITGAVWVALTRLRP-EAAVTGGVGALAVFGHALDGVSTAVGTAQLGFGERTPVSRILLELGG-L 207
WP_015910634_1 DUF63 LLAPVVAVAVSTGL-SPE----GARWSTIALVLTIPIAGAVWVGLTRLRP-ETAITGGVGALAVFAHALDGVSTAVGTTHLGFGERTPVSRILLEIGG-L 207
WP_008006417_1 DUF63 CLVPVVAAAVSTGL-SPA----GARWSTIALALTLPIAGVVWVGLTRLRP-ETAVTGGVGALAVFGHALDGVSTAVGTTHLGFGERTPVSRILLEFDW-I 207
WP_008583911_1 DUF63 LVVPVVAVAAGTGL-SPE----GARWSAVALAITVPVAGAAWVGLTRFRP-EAAVTGGVGALAVFGHALDGVSTAVGTVQLGFGERTPLSRLLLEVGG-L 205
WP_006627969_1 DUF63 LVVPVVAVAAVTGL-SPE----GARWSAVALAVTLPVAGAAWGGLLRLRP-ETAITGSVGALAVFGHALDGVSTAVGTTQLGFGERTPVSRLLLELGG-L 205
WP_053772749_1 DUF63 CLVPVVAVAVGTGV-SPV----GARWSTLALALTVPVAGATWAGLTRARP-ETRVTGGVGGLAVFGHALDGVSTAVGTTQLGFGERTPLSRILLEVGG-L 205
WP_004046581_1 DUF63 CLAPVVAVAVSTGL-SPA----GARWSTVALALTLPIAGAAWVGLTRFRP-ETAITGGVGALAVFGHALDGVSTAVGTTQLGFGERTPLSRILLEFDW-I 207
WP_006112271_1 DUF63 LLLPVVAVAVGTGL-SPV----GARWSAAALALTVPIAGGAWLGLTRFRP-AAAVTGGVGALAVFGHALDGVSTAVGTTQLGFGERTPVSRILLELGG-L 205
WP_008443330_1 DUF63 ILIPVVAVAVGTGL-SPT----GARWSAGALALTVPIAGAAWVVLTRFRP-EAAVTGGVGALAVFGHALDGVSTAVGTTQLGFGERTPLSRILLEIGG-L 205
WP_008367944_1 DUF63 LLVPVVALAVATGL-SPI----GARWSAVALALAIPIAAAAWVGLTRFRP-EAAVTGSVGALAVFGHALDGVSTAVGTTQLGFGERTPLSRILLEIGG-L 205
WP_007345917_1 DUF63 LLVPVVAVAVATGL-SPI----GARWSAVAVALAIPIAAAAWVGLTRFRP-EAVVTGSVGALAVFGHALDGVSTAVGTTQLGFGERTPLSRILLEIGG-L 205
WP_004595209_1 DUF63 LLVPVVAVAVATGL-SPI----GARWSAVALALAIPIAAAAWVGLTRFRP-EAAVTGSVGALAVFGHALDGVSTAVGTTQLGFGERTPLSRILLEIGG-L 205
WP_096395669_1 DUF63 CLLPVVAVALGAGL-SPA----GARWSAVAFALTLPIAGAAWVGLTRFRP-EAAATGSVGALAVFGHALDGVSTAVGTTQLGFGERTPVSRILLEIGG-L 205
WP_017344445_1 DUF63 LLVPVVAVALGAGV-SPA----GVRWSAVAFALTLPIAGAAWVGLTRFRP-EAAATGSVGALAVFGHALDGVSTAVGTTQLGFGERTPVSRILLEIGG-L 211
ESS03942_1 putative  LLVPVVAVAVAGGVSGPG-----ARWSAVAVVVAAPVAIGAWRALVRINP-AVGVTGGVGGLVVFAHALDGVSTAVGVTQLGFGERSPLSRLLLELGG-L 214
WP_066418621_1 DUF63 VLAPVVAAAVAGGVSGPG-----AYWSAVAAVASVPVAAVVRRALGRVEP-GVAVTGGVGALAVYGHALDGLSTAVGVTQLGFGERSPLSRVILELDW-V 217
ESS10517_1 putative  LLIPVVAAALWVGT-AES--GLTLAWPVAALVASLPITAGVWVGLQKANS-AVQVTGLVGVFAVFGHVLDGVSTAVGIDILGFGERTPLSRLIFDIAGSL 210
ERH00845_1 putative  VAVPLVGTTIIVGF-TQN--TLAPVWPAAGAGMAVLLTAAGWATLRRRRPTATTATGWVGLLAVFGHTLDAVSTTVGVDVLGFGERTPLSRVILDAAAAL 207
ESS08053_1 putative  VAVPLVGTTIIVGF-TQN--TLAPVWPAAGAGMAVLLTAAGWATLRRRRPTATTATGWVGLLAVFGHTLDAVSTTVGVDVLGFGERTPLSRVILDAAAAL 207
WP_103427710_1 DUF63 AVVPTCATGIAVGL-SRG--TFSPLWSTVALGVSLAVGAAAWALLTRLVP-RVEHTGRVGLLALLAHAVDGVSTAVGVDVLGFGERTPLSRYIIEFAAGL 210
WP_049937371_1 DUF63 LALPVVAAVLAVGA-GAG--TLRLGYALGILVGAGVAAAGTWLLLRRLRP-AVTVTGRLGILAVGAHALDGVSTAVGVDLLGFGERTPLSRLVIEFGASL 204
WP_049987236_1 DUF63 LAVAAVVAVLLVGA-SRG--TLSPAVPAAGVLAAVALAGAIWAGLTRALP-EVRKTGLLGAFAVFAHVLDGVTTAVGIDLLGFGERTPLSRLIIEFAAGL 210
WP_006053788_1 DUF63 LSAVAVAAAFLVGA-SEG--TLAPVVPAIGLVVATVVSAGAWVILTRVLP-DARITAPVGILTVFGHALDAVSTAVGIDLLGFSERTPLSRAIIEFAATL 206
```

```
WP_008384832_1 DUF63 LVAVTVADALFVGA-VRG--TLAPLVPAVGLVAAAVVAAAAWAVLIRAYP-AARLTGAVGFLAVFGHALDAVSTAVGVDLLGFGERTPLSRAIIEFAATL 206
WP_058582368_1 DUF63 AAIPAVGLPVSAAL-AAG--SFSPFWSTVALVVSLVLAAVAWTALKTIRP-ETTVTHWAGALAVFGHTLDGVSTAVGLDVLSFGERTPLSRLIFEFAAEL 205
WP_004060105_1 DUF63 VAGGLVAATIAAGL-QRG--TLALFWPGIGLVVSLVISPIAWLVVQRLSP-EAATAGSLGLLAIFGHTLDAVSTAVGIDVLGFGERTPLSQVILEVAAAL 205
WP_008320401_1 DUF63 GGGVLLVSAIRAGV-AAG--TLALFWPGVGLVASVVVSLVAWPALKRFSP-ETAVAGTLGWLAIFGHTLDAVSTAVGIDVLGFGERTPLSKVILEAAAAL 205
WP_008325536_1 DUF63 LAGLLVGFALFVGL-AGG--TLAPLWPGIALVLSLLVGPAAWVAVTHIAP-EATDAGRLGALTIFGHTLDAISTAVGIDVLGFGERTPLSQVILEAAAAL 205
WP_007543205_1 DUF63 LAGLLVGFALFVGL-AGG--TLAPLWPGIALVLSLLVGPVAWVAVTRIAP-EAADAGRLGALTIFGHALDAISTAVGIDVLGFGERTPLSKVILEAAAAL 205
WP_004042703_1 DUF63 VAAGLAGFALFIGL-AAG--TLSVVWPGTALVAALVVTPLTWRVVTRLAP-EAAAAGSLGLLALFGHALDALSTAVGVDVLGFGERTPLSRVILDAAAAL 205
WP_004062371_1 DUF63 VAAGLAGFALFAGL-AAG--TLSVVWPGLALVAAVVVTPLTWRVVTRLAP-EAAAAGSLGLLALFGHALDALSTAVGVDVLGFGERTPLSRVILDAAAAL 205
WP_008575916_1 DUF63 VAAGLAGFALFAGL-AAG--TLSVVWPGLALVAAVVVTPLTWRVVTRLAP-EAAAAGSLGLLAIFGHALDALSTAVGVDVLGFGERTPLSRVILDAAAAL 205
WP_008575916_1 DUF63 VAAGLAGFALFAGL-AAG--TLSVVWPGLALVAAVVVTPLTWRVVTRLAP-EAAAAGSLGLLAIFGHALDALSTAVGVDVLGFGERTPLSRVILDAAAAL 205
WP_008575916_1 DUF63 VAAGLAGFALFAGL-AAG--TLSVVWPGLALVAAVVVTPLTWRVVTRLAP-EAAAAGSLGLLAIFGHALDALSTAVGVDVLGFGERTPLSRVILDAAAAL 205
WP_089777339_1 DUF63 VAAGLSGFALFIGL-AAG--TLSVVWPGFALVAALVVTPLTWRVVTRLAP-EAAAAGSLGLLAIFGHALDALSTAVGVDVLGFGERTPLSRIILDAAAAL 205
WP_007275294_1 DUF63 VAAGLAGFALFIGL-AAG--TLSVVWPGVALVAALVVSPLTWRVVTRLAP-EAAAAGSLGLLAIFGHALDALSTAVGVDVLGFGERTPLSRVILDAAAAL 205
WP_004969536_1 DUF63 VAFGLAGFALFIGL-AAG--TLSVVWPGVALVAALVVSPLTWRVVTRLAP-EAAAAGSLGLLAIFGHALDALSTAVGVDVLGFGERTPLSRVILDAAAAL 205
WP_050459703_1 DUF63 VAAGLAGFALFIGL-AAG--TLSVVWPGVALVAALAVTPLTWRVVTRIDT-DAAAAGSLGLLAIFGHALDALSTAVGIDVLGFGERTPLSRIILEAAAAL 205
WP_008095201_1 DUF63 VAAGLSGFALFIGL-AAG--TLSIVWPGVALVAALAVTPLTWRVVTRVAP-EAAAAGSLGLLAIFGHALDALSTAVGIDVLGFGERTPLSRIILEAAAAL 205
WP_049967774_1 DUF63 VAAGLSGFALFIGL-AAG--TLSIVWPGVALVAALAVTPLTWRVVTRVAP-DAAAAGSLGLLAIFGHALDALSTAVGIDVLGFGERTPLSRIILEAAAAL 205
WP_058569127_1 DUF63 VAAGLAGFALFIGL-AAG--TLSVVWPGVALVAALAATPLTWRIVTRVAP-EAAAAGSLGLLAIFGHALDALSTAVGIDVLGFGERTPLSRVILEAAAAL 205
WP_058828475_1 DUF63 VAAGLAGFALFIGL-AAG--TLSVVWPGFALVAALVATPLTWRVVTRVAP-EAAAAGSLGLLAIFGHALDALSTAVGIDVLGFGERTPLSRIILEAAAAL 205
WP_009376007_1 DUF63 ALVPSLVAALLVGQ-GRG--SLAVTWPAIALVVAVVLTGVTWVVLRQVRP-EVALTGSVGVLALFGHTLDGVSTAVGLEVLGFSERTPLSRLIIELGQAL 207
WP_101294394_1 DUF63 ALLAALGLALAVGS-ARG--TLHPRWPALGLVASAVVAAAAWVGLARLRP-QAAITGGAGALAMFGHVLDGVSTALGVDVLGFGERTPLSRVIIEFGDAL 205
WP_089671020_1 DUF63 ALLPVVGGAVAVGA-QQG--TLAVFWPAVGVVLSVAVTVLAWVGLRRLRPAAASTTGSVGLLALFAHSLDAVSTAVGVDVLGFGERTPASRLIIEFADRL 208
WP_096390099_1 DUF63 ALVPVIATTLAVGA-GAG--SLTPRWPAIALVVSIVVGALAWAGLVRIAP-RASVTGSVGMLAVLGHALDAVSTAVGVDVLGFGERTPLSQLIMDGAAAL 220
WP_021073253_1 DUF63 ALLPVIATALAVGA-DGG--SLTPRWPAIALVGSIVVGAAAWVGLVRLVP-RASITGSVGSLAVLGHALDAVSTAVGVDVLGFGERTPLSRLIMDAAAAL 224
WP_007983568_1 DUF63 GVAVVLLVALWYGV-QHD--SLRLFWPFVALVVAVILTGLLWAATQFFYPDVAIITGAVGVLTLFAHALDSVSTAVGIDVLGFGERTPLSQAILEIAHSL 208
WP_049969758_1 DUF63 AILLVVAVALWYGV-QND--SLRLFWPFVALVVAVVLTGLLWAGTQFFYPHVAVITGAVGVLTLFGHALDSVSTAVGIDVLNFGERTPVSRAILEFAELL 206
WP_066141551_1 DUF63 GVALVLVVALWFGI-EHD--SLRLFWPFVALVVAVILTGMLWAATQFFYPDVAIITGAVGVLTLFAHALDGVSTAVGIDVLGFGEQTPLSRAILEIAHSI 208
WP_008414921_1 DUF63 AALLAVGAAFGVGY-GRG--TITPYWPLVALVLSIVLAAGTWGLFRRALPGVAATTGVAGALVVLAHALDGISTAVGIDVLEVTERSPLPRAIMGVAEGL 199
WP_066381248_1 DUF63 AALLAVAAAFAVGY-TRG--TITPVWPLVALACSIALAAGTWGLFRRAVPGVAATTGAAGALVVLAHALDGVSTAVGIDVLEVTERSPLPRAIMEFAETL 199
WP_014051708_1 DUF63 VAVLVLGVGVQWGA-TSG--RFSPAWPAVAAILGVTLGFALWALLTRVRP-DVLVAGSAGALAVVAHSLDGVSTAVGIDILGATERTPLSRAIIEFGASL 205
WP_049980148_1 DUF63 VAALVLAVGVQWGL-AEG--TFAPAWPAVAAVLGVAVGIAVWLGLRRAYP-GVGDAGAAGALAVVAHSLDGISTAIGIDLLGATERTPLSRAVIEFGATL 204
WP_053948907_1 DUF63 LALIALVVGFRWGF-DEG--TVGVVWPAIAAVFGVAVGVGVWVGLRRVSP-SIGDAGVAGALAIVAHSLDGISTAIGIDVLDATERTPLSRMVIEFGAAL 203
WP_079233625_1 DUF63 LALIALVVGFRWGF-TEG--TVGVVWPAVAAVLGVAVGIGVWVGLQRPYP-GIADAGGAGALAVVAHSLDGVSTAIGIDVLGATERTPLSRAVIEFGATL 203
KPN32293_1 hypotheti VAALVLAVGVQWGL-DEG--TFAPAWPVVAAVLGVGVGIAVWLGLRRAYP-GVGDAGAAGALAIVAHSLDGISTAIGIDLLGATERTPLSRAVIEFGATL 204

WP_006882700_1 DUF63 PTYPYIGKGWLFAVVKLVVVAGIVVLLADLVEDEPTQGNLLFAFVAAVGMGPAANNLFLFLIGPAL------- 280
WP_015790720_1 DUF63 PTAPYLGSGWLFLLVKVLLATLVIITFAEYIEERPARGHLLLVIITAVGAGPAVNNFVLFTLG---------- 280
WP_008524992_1 DUF63 PTAEYIGSGWLFLVVKLLLATAVVVAFAEYIEERPARGSLFLALITAVGIGPAMNNVVIYILG---------- 279
WP_015762485_1 DUF63 PTADLLGVGWLFLLVKLVIAAAIVVSFNRYLADEPTEGALVFSLIAAVGMGPATNNLFLFLAPAF-------- 277
WP_018257825_1 DUF63 PTADLLGVGWLFLLVKLVIAAAIVVSFNRYLADEPAEGALVFSLIAAVGMGPATNNLFLFLAPAF-------- 277
WP_101349367_1 hypot PTAPYLGKGWLFVFVKLLVAVAVVFLLDDYLEEEPIEASLLLALVTAVGIGPATNNIVLFLFSPV-------- 278
WP_004518630_1 DUF63 PTAPYLGKGWLFVFVKLLVAIGVVFLLDDYLEEDPVEASLLLALVTAVGIGPATNNIVLFLFSPV-------- 278
WP_004963804_1 DUF63 PTAPYLGKGWLFVFVKLLVAIGVVFLLDDYLEEDPVEASLLLAVVTAVGIGPATNNIVLFLFSPV-------- 278
WP_007189047_1 DUF63 PTAPYLGKGWLFVFVKLLVAIGVVFLLDDYLEEDPVEASLLLAVVTAVGIGPATNNIVLFLFSPV-------- 278
WP_004593738_1 DUF63 PTASYLGKGWLFVFVKLLVAVAVVFLLDDYLEEDPAEASLLLALVTAVGIGPATNNIVLFLFSPV-------- 278
WP_005535867_1 DUF63 PTASYLGKGWLFVFVKLLVAVAVVFLLDDYLEEDPAEASLLLALVTAVGIGPATNNIVLFLFSPV-------- 278
WP_053967006_1 DUF63 PTASYLGKGWLFVFVKLLVAVAVVFLLDDYLEEEPVEASLLLALVTAVGIGPATNNIVLFLFSPV-------- 278
WP_008310684_1 DUF63 PTAPYLGKGWLFVFVKLLVAIAVVFLLDDYLEEDPVEASLLLALVTAVGIGPATNNIVLFLFSPV-------- 278
WP_058997315_1 DUF63 PTAPYLGKGWLFVFVKLLVAIAVVFLLDDYLEEDPVEASLLLAVVTAVGIGPATNNIVLFLFSPV-------- 278
WP_050037596_1 DUF63 PTALYLGKGWLFVFVKLLVAIAVVFLLDDYLEEDPVEASLLLALVTAVGIGPATNNIVLFLFSPV-------- 278
WP_050037596_1 DUF63 PTALYLGKGWLFVFVKLLVAIAVVFLLDDYLEEDPVEASLLLALVTAVGIGPATNNIVLFLFSPV-------- 278
WP_014039614_1 DUF63 PTAPYLGKGWLFVFVKLLVAIAVVFLLDDYLEEDPVEASLLLALVTAVGIGPATNNIVLFLFSPV-------- 278
WP_064288256_1 DUF63 PTAPYLGKGWLFVFVKLLVAIAVVFLLDDYLEEDPVEASLLLALVTAVGIGPATNNIVLFLFSPV-------- 278
WP_049994803_1 DUF63 PVAETIGVGWLFVLVKLAIAVIVVVAFADYLESDPVRGNLAFAAIIALGLGPAVNNLLLFALRDSAALATVAS 289
WP_020445351_1 DUF63 PTAEFVGAGWLFVLVKLIVATGIVVLMREWIRDAPQEARLVLLLVAAVGLGPGTNNLMLFAVGG--------- 296
WP_083761671_1 DUF63 P--PDIGGGWLFALVKIALGGAIVVLFESYVREAPAEGYLVLGLVAAVGFGPGFHNLVLFALL---------- 297
WP_084260786_1 DUF63 PTAELIGAGWLFAAVKVGLGVAVVALFEEYVRADPTEGYLLLGLVVAVGLGPGVHNVVLFALA---------- 314
WP_014556864_1 DUF63 PTDAIIGSGWLFVVVKLLIASVVVWVFADIADTAPAQSNILLAIISAVGFGPGVHNLLLFSVAG--------- 269
WP_070365684_1 DUF63 PTAPYIGTGWVFALVKIALGVAVVWFIADLVRDDPPLGNGLLLVITAVGLGPATHNLILFAVAGPAGV----- 272
WP_049951854_1 DUF63 PTAEYIGAGWLFVVVKLALALAVVGLFKEYVRDAPRQARLLLAFIAAVGLGPGVHNVLLFTIA---------- 273
WP_015319722_1 DUF63 PTAEYVGGGWLFIVVKIALALVIVGLFREYVEDAPRQGRALLALLAAVGLGPGAHNVLLFLVT---------- 271
WP_005554260_1 DUF63 PTADYIGGGWLFAVVKVALAVVIVGLFRAYVEDAPRQGRALLALLAAVGLGPGAHNVLLFLVT---------- 270
AGB15442_1 putative  PTAEYIGAGWLFVLVKLTLALVVVWLFADLVRDRPRAGRIALGFVAAVGLGPGVHNLLLFTISG--------- 276
WP_007698493_1 DUF63 PTAEYIGAGWLFVLVKLTLALVVVWLFADLVRDRPRAGRIALGFVAAVGLGPGVHNLLLFTISG--------- 276
WP_086886713_1 DUF63 PTAEYVGAGWLFVLVKVVLALVVLGLFREFVREAPRQARLLLAFIAAVGLGPGVHNVLLFMVG---------- 273
WP_049926795_1 DUF63 PTAEYIGAGWLFVLVKVGLAMVVLGLFREFVEEAPRQARLLLAFIAAVGLGPGVHNVLLFMVG---------- 273
WP_049922464_1 DUF63 PTAEYIGAGWLFVLVKVVLALLILGLFREYVQERPRQGRFLLALIAAVGLGPGVHNVLLFVVA---------- 272
WP_008008940_1 DUF63 PTAGVIGAGWLFVLVKVGLAMVILGLFREYVEERPRQARTLLTLVAAVGLGPAVHNTLLFMVG---------- 273
WP_098724536_1 DUF63 PTAEYIGSGWLFVLVKVSLAMVILGLFREYVEEAPRQARTLLALVAAVGLGPATHNTLLFMVS---------- 278
WP_097380374_1 DUF63 PTAEYVGAGWLFVLVKVVLAVVILGLFREYVDERPQQARTILALVAAVGLGPAVHNTLLFTVA---------- 273
WP_006429103_1 DUF63 PTAEYIGAGWLFVLVKVALALVILGLFREYVEEAPRQGRTMLALVAAVGLGPAVHNTLLFAVA---------- 273
WP_008457054_1 DUF63 PTATYIGAGWLFVLVKVVLAVVILGLFREYVEDRPQQARTVLALVAAVGLGPAVHNTLLFAVG---------- 273
WP_008457054_1 DUF63 PTATYIGAGWLFVLVKVVLAVVILGLFREYVEDRPQQARTVLALVAAVGLGPAVHNTLLFAVG---------- 273
WP_008457054_1 DUF63 PTATYIGAGWLFVLVKVVLAVVILGLFREYVEDRPQQARTVLALVAAVGLGPAVHNTLLFAVG---------- 273
WP_007107690_1 DUF63 PTAEYIGAGWLFVLVKVVLAVVILGLFREYVEDRPQQARTVLALVAAVGLGPAVHNTLLFAVG---------- 273
WP_015299220_1 DUF63 PTAEYVGAGWLFVLVKVALALVIVGLFREYVADRPQQGRTVLALVAAVGLGPAVHNVLLFTVM---------- 273
WP_006649902_1 DUF63 PTAEYVGAGWLFVLVKVALALVIVGLFREYVADRPQQGRTVLALVAAVGLGPAVHNVLLFTVM---------- 273
WP_066299764_1 DUF63 PTAEYVGAGWLFVLVKVALALVIVGLFREYVADRPQQGRTILALVAAVGLGPAVHNVLLFTVM---------- 273
WP_049965539_1 DUF63 PTAEYVGAGWLFVLVKVALALVIVGLFREYVADRPQQGRTVLALVAAVGLGPAVHNVLLFTVM---------- 273
WP_049989654_1 DUF63 PTAEYVGAGWLFVLVKVVLGLVILGLFREYVEEQPRQARTLLALVAAVGLGPAVHNVLLFIVT---------- 273
WP_076147241_1 DUF63 PTAEYVGAGWLFVLVKVALALVIVGLFREYVADRPQQGRTVLALVAAVGLGPAVHNVLLFTVM---------- 273
WP_012941789_1 DUF63 PIAEYVGAGWLFVLVKAALALVILGLFKEYVEDAPQQARIVLSGIAAVGLGPGVHNVLLFIAT---------- 273
WP_008895944_1 DUF63 PTAEYVGAGWLFVLVKAALALVILGLFKEYVEDAPQQARIVLAGIAAIGLGPGVHNVLLFIAT---------- 273
WP_007142166_1 DUF63 PTAEYVGGGWVFVVVKAALALAILGLFTEYVEERPGEARTILALVAAVGLGPGTHNVLLFLVA---------- 273
WP_013879207_1 DUF63 PTAEYIGAGWLFVLVKVVLALVVLGLFKEYVEEEPRQARTILGLLAAIGLGPGVHNTLLFLVT---------- 273
WP_006088775_1 DUF63 PTVEYIGAGWLFILVKVVLALVIVGLFKEWVEEQPQQARTVLALVAAVGLGPGVHNVLQFIVT---------- 273
WP_076581448_1 DUF63 PTDEYIGAGWVFIVVKVLLAVVILGLFKEYVDERPQQARVVLALIAAVGLGPGVHNVLQFVIT---------- 273
WP_006064996_1 DUF63 PTEAYIGAGWLFILVKVALAMVILGLFREYVEERPQQARLLLALVAAVGLGPGTHNVLLFLIT---------- 273
WP_049890149_1 DUF63 PTEPYIGAGWLFVLVKVVLALVILGLFKEYVDEQPQQARLILALLAAVGLGPGVHNVLLFIIT---------- 273
WP_005580668_1 DUF63 PTAEYIGGGWLFVLVKVGLALAILGLFREYVEEAPRQARTILALVAAIGLGPGVHNTLLFVVA---------- 273
WP_087715216_1 DUF63 PTAEYIGGGWLFVLVKVALAVVILGLFKEYVEEAPRQARTILALIAAVGLGPGVHNVLLFMVG---------- 273
WP_006664680_1 DUF63 PTAEFIGAGWLFVVVKVALVMGVLGLFREYVEDAPRQARLLLAFIAAVGFGPGIHNVILFAVA---------- 273
WP_006110228_1 DUF63 PTAEFIGAGWLFVVVKVALVMGILGLFREFVEDAPRQARLLLAFIAAVGFGPGIHNVILFAVA---------- 273
WP_006826566_1 DUF63 PTAEFIGAGWLFVVVKVALVMGVLGLFREYVEDAPRQARLLLAFIAAVGFGPGIHNVVLFAVA---------- 273
WP_004214806_1 DUF63 PTADYIGGGWLFVVVKLALAMVILGLFREYVEEAPRQARLLLAFIAAVGLGPGAHNVLLFAVA---------- 273
```

```
WP_006653721_1 DUF63 PTAEYIGGGWLFIVVKVALAMVVLGLFREYVEEAPRQARLVLAFIAAVGLGPGVHNVLLFAVA---------- 273
WP_006653721_1 DUF63 PTAEYIGGGWLFIVVKVALAMVVLGLFREYVEEAPRQARLVLAFIAAVGLGPGVHNVLLFAVA---------- 273
WP_050049052_1 DUF63 PTADVIGVGWLFALVKLGLAVAVLALFAGYVRDEPSEGFGLLGLIAAVGLGPASYNVLLFAVMTPTGF----- 272
WP_075938119_1 DUF63 PPAAVFGEAWLFVLVKIVVATLVLFAFRDYVRDDPGEGSLVLGLVAAVGFGPGVHNLVLFTIA---------- 271
WP_077206978_1 DUF63 PTASVIGKGWLFVLVKLLIAAAILVLFADFVEEDPAQGNAALGVVAAVGLGPGTHNLLLFAAAGMV------- 280
WP_049892606_1 DUF63 PTAPVVGGGWLFVVVKLALAAGVVVLLAESVRDRPQEGTAILGLVAAVGLGPGVHNVVLFIAANPTGF----- 274
WP_009487551_1 DUF63 PTEPYLGVGWLFVLVKVALVAVVVVLLADYVREVPGEGYLLLGLVAAVGLGPGVHNVLLFVASNPTGF----- 272
WP_059057568_1 DUF63 PTASVLGVGWLFVLVKTVLGAAVVLLLAEYVREDPAEGNLLLAVVTAVGLGPGAHNVLLFVAANPAGF----- 272
WP_058984064_1 DUF63 PTAPVIGVGWLFVLVKTALGAAVVLLLAEYVREDPAEGNLLLAVVTAVGLGPGAHNVLLFVAANPAGF----- 272
EMA41075_1 hypotheti PTASLLGAGWLFVVVKLALAAAVLVLLASYVREEPGEGYLLLGAVAAVGLGPGAHNLLLFTIAS--------- 279
WP_049998124_1 DUF63 PTTPFIGAGWLFVLVKLALAAAVVVFLSEYVREEPAEGYLLLGAVAAVGLGPGAHNLLLFTIATP-------- 280
WP_049902415_1 DUF63 PTAPVLGAAWLFVLVKLALAAAVVFLLAGYVRDEPAEGYLLLGAVAAVGLGPGAHNLLLFTIASP-------- 282
WP_006079317_1 DUF63 PTAPVLGTVWLFVLVKLVLASVVVVLFSGYVRDEPTEGYLLLGAVAAVGLGPGVHNLLLFTIASP-------- 282
WP_080509272_1 DUF63 PTEPTLGAGWLFVLVKLAVACVVVAVIAPAAEEAPREGNALLALVTAVGLGPGVHNALLFAVAGG-------- 335
WP_008000170_1 DUF63 PAVPVFGEGWLFVLVKLAVASLVVWLFTPYVRDEPAEGYLLLGFIAAVGLGPAAHNLLLFSVVA--------- 308
WP_049983070_1 DUF63 PAVPVLGEGWLFLLVKLVVASVVTWLFAAYVRETPTEGFLFLGFVASVGLGPAAHNLLLFAVAA--------- 271
WP_095637763_1 DUF63 PSLPVLGEGWLFLLVKLAVASAVVWLFAEYVRETPAEGYLLLGFVAAVGLGPAAHNLLLFAIAG--------- 271
WP_015910634_1 DUF63 PSLPVVGEGWLFLLVKLAVASAVVWLFAGYIRETPGEGYLLLGFVAAVGLGPAAHNLLLFAVAA--------- 271
WP_008006417_1 DUF63 SSVPVLGEGWVFLLVKLAVASAVVWLFAEYVRETPAEGFLLLGFVAAVGLGPAAHNLLLFAVAA--------- 271
WP_008583911_1 DUF63 PPVPVLGEGWLFLLVKLAVATAVVWLFAAYVRETPGEGFLLLGFVAAMGLGPAVHNLVLFSVAA--------- 269
WP_006627969_1 DUF63 PPVPVLGEGWLFLLVKLAVATAVVWLFAAYVRESPGEGFLLLGFVAAMGLGPAVHNLVLFSVAA--------- 269
WP_053772749_1 DUF63 PSLPVIGEGWLFLLVKLAVASAVVWLFAAYVREEPAEGYLLLGFVAAMGLGPAAHNLLLFSVAA--------- 269
WP_004046581_1 DUF63 PSVPVIGEGWLFLLVKLAVASAVVWLFAEYVRDRPTEGYLLLGFVAAMGLGPAAHNLLLFAVAG--------- 271
WP_006112271_1 DUF63 PSVPVLGEGWLFLLVKLVVATAVVWLFAAYVRETPAEGYLLLGFVAAMGLGPAAHNLLLFSVAA--------- 269
WP_008443330_1 DUF63 PSLPVVGEGWLFLLVKLAVASAVVWLFAAYVREEPAEGYLLLGFVAAMGLGPAAHNLLLFSVAA--------- 269
WP_008367944_1 DUF63 PSLPVIGEGGLFLLVKLAVASAVVWLFAAYVREEPAEGYLLLGFVAAMGLGPAAHNLLLFSVAA--------- 269
WP_007345917_1 DUF63 PSLPVIGEGGLFLLVKLAVASAVVWLFAAYVREEPAEGYLLLGFVAAMGLGPAAHNLLLFSVAA--------- 269
WP_004595209_1 DUF63 PSLPVIGEGGLFLLVKLAVASAVVWLFAAYVREEPAEGYLLLGFVAAMGLGPAAHNLLLFSVAA--------- 269
WP_096395669_1 DUF63 PSLPVIGEGWLFLLVKLAVASAVVWLFAAYVREEPTEGYLLLGFVAAMGLGPAAHNLLLFSVAA--------- 269
WP_017344445_1 DUF63 PSLPVIGEGWLFLLVKLAVASAVVWLFAAYIREEPTEGYLLLGFVAAMGLGPAAHNLLLFSVAA--------- 275
ESS03942_1 putative  PSVPVFGEGWVFLLVKLAVAGVAVSLFVPYVRETPREGMLFLGFLAAVGLGPAAHNLLLFTIAG--------- 278
WP_066418621_1 DUF63 PAVPVLGEGWLFLLVKLAVASAVVYLFVPYVREEPSEGLLLLGFVAAVGLGPAVHNLLLFTVTA--------- 281
ESS10517_1 putative  PTADLLGTGWLFVVVKAALAGFVVSLFVEFVREDPSQGYLLLGFIAAVGLGPGVHNLLLFVVAG--------- 274
ERH00845_1 putative  PTADIIGTGWLFILVKVGVVSAVVVLFAEYVAEEPSEGYPLLGVIAAVGLGPGTYNLLLFAITAG-------- 272
ESS08053_1 putative  PTADIIGTGWLFILVKVGVVSAVVVLFAEYVAEEPSEGYPLLGVIAAVGLGPGTYNLLLFAITAG-------- 272
WP_103427710_1 DUF63 PTEPLLGSGWLFVLVKLALVGGVVVLFADLMEEDPREGSLLLGVVAAVGLGPGAHNLLLFAIA---------- 273
WP_049937371_1 DUF63 PTAPAVGGAWLFVLVKLVVASAVVVALTETVREEPTTGRLLLGFVAAVGLGPGAHNVLLFTVAP--------- 268
WP_049987236_1 DUF63 PTEPVLGAGWLFVLVKIAVASVVVWLFADLVESDPAESRLLLGFVAAVGLGPATHNALLFTISAPA------- 276
WP_006053788_1 DUF63 PTEPFFGTVWLFVLVKLVIVSGIVALFADYVREDPTEGNLLLGFVAAVGLGPGVHNLILFTILGGA------- 272
WP_008384832_1 DUF63 PTEPYLGTVWLFVLVKLVIASGIVALFVDYVREDPAEGYALLGFVAAVGLGPGAHNLLLFTVLGGA------- 272
WP_058582368_1 DUF63 PTADIIGTGWLFVLVKLLLAGFVVSIMADYVREEPSEGLLLLTLVAAVGLGPGVHNLILFAVA---------- 268
WP_004060105_1 DUF63 PTAETLGIGWLFVLVKLAVVGLVVTLLADYVREEPTEGALLLGFVAAVGLGPGVHNLLLFAIAGA-------- 270
WP_008320401_1 DUF63 PTADVIGVGWLFVLVKLAVAGLVVTLLADYVREEPTEGSLLLGFVAAVGLGPGVHNLLLFAISGA-------- 270
WP_008325536_1 DUF63 PTAEVIGVGWLFVLVKLIVAGAVVTLLADYVREEPTEGALLLGLVAAVGLGPGVHNLLLFAVSGV-------- 270
WP_007543205_1 DUF63 PTAEVIGVGWLFVLVKLVVAGFVVTLLADYVREEPTEGALLLGLVAAVGLGPGVHNLLLFAVSGV-------- 270
WP_004042703_1 DUF63 PTAEVIGVGWLFVLVKLAVAGFVVALLADYVREDPTEGSLLLGLVAAVGLGPGVHNLLLFAIAG--------- 269
WP_004062371_1 DUF63 PTAEVIGVGWLFVLVKLAVAGFVVALLADYVREDPTEGSLLLGLVAAVGLGPGVHNLLLFAIAG--------- 269
WP_008575916_1 DUF63 PTAEVIGVGWLFVLVKLAVAGFVVALLAGYVREDPTEGSLLLGLVAAVGLGPGVHNLLLFAIAG--------- 269
WP_008575916_1 DUF63 PTAEVIGVGWLFVLVKLAVAGFVVALLAGYVREDPTEGSLLLGLVAAVGLGPGVHNLLLFAIAG--------- 269
WP_008575916_1 DUF63 PTAEVIGVGWLFVLVKLAVAGFVVALLAGYVREDPTEGSLLLGLVAAVGLGPGVHNLLLFAIAG--------- 269
WP_089777339_1 DUF63 PTAEAVGVGWLFVLVKLAVAGFVVALLADYVREDPTEGSLLLGLVAAVGLGPGVHNLLLFAIAG--------- 269
WP_007275294_1 DUF63 PTAEALGVGWLFVLVKLAVAGFVVALLADYVREDPTEGSLLLGLVAAVGLGPGVHNLLLFAIAG--------- 269
WP_004969536_1 DUF63 PTAEALGVGWLFVLVKLAVAGFVVALLADYVREDPTEGSLLLGLVAAVGLGPGVHNLLLFAIAG--------- 269
WP_050459703_1 DUF63 PTAEVIGVGWLFVLVKLVVAGFVVALLADYVREDPTEGSLLLGLVAAVGLGPGVHNLLLFAIAG--------- 269
WP_008095201_1 DUF63 PTAEVIGVGWLFVLVKLVVAGFVVALLADYVREDPTEGSLLLGLVAAVGLGPGVHNLLLFAIAG--------- 269
WP_049967774_1 DUF63 PTAEVIGVGWLFVLVKLVVAGFVVALLADYVREDPTEGSLLLGLVAAVGLGPGVHNLLLFAIAG--------- 269
WP_058569127_1 DUF63 PTAEVIGVGWLFVLVKLVVAGFVVALLADYVREDPTEGALLLGLVAAVGLGPGVHNLLLFAIAG--------- 269
WP_058828475_1 DUF63 PTAEVIGVGWLFVLVKLVVAGFVVALLADYVREDPTEGALLLGLVAAVGLGPGVHNLLLFAIAG--------- 269
WP_009376007_1 DUF63 PTAQYVGGAWLFVLVKLLVAAGVVTLLVDYVREEPAEGYLLLGLVAAVGFGPGVHNLVLFSIA---------- 270
WP_101294394_1 DUF63 PTAELVGAGWLFVLVKLAVASVVVALMADYVREEPTEGYLLLALIAAVGLGPGTHNLVLFSIA---------- 268
WP_089671020_1 DUF63 PTAEALGSGWLFVVVKLVLVGGVVVLFADYIEDEPTEGYLLLGFIAAVGLGPGVHNLLLFAVTGG-------- 273
WP_096390099_1 DUF63 PTAETLGVGWLFVLVKLTLVCGIVVLFADYVEADPTEGYLLLGFVAAVGLGPGAHNLLLFTVVGGA------- 286
WP_021073253_1 DUF63 PTAEALGVGWLFVLVKLALVCGIVVLFADYVEADPTEGYLLLGFIAAVGLGPGAHNLLLFTVAGGA------- 290
WP_007983568_1 DUF63 PTADSFGVGWLFVVVKLAIAEFVVLLFADYVSSEPTEGYLLLGVVAAVGLGPGAHNLILFAVTG--------- 272
WP_049969758_1 DUF63 PTADAIGVGWLFILVKLAIAEFVVILFADYVADEPAEGYLLLGVVAAVGLGPGAHNLILFAITG--------- 270
WP_066141551_1 DUF63 PAGDSLGVGWLFVLVKLAIAEFVVLLFADYVASEPTEGYLLLGVVAAVGLGPGAHNLILFAITG--------- 272
WP_008414921_1 DUF63 PTAETIGVGWLFVLVKLALACVIVWLMAEYVEDEPNEGFLLLALIAAVGFGPGVHNLLLFAVGG--------- 263
WP_066381248_1 DUF63 PTSEAIGVGWLFVLVKLTLACGIVWLMAEYVEDEPSEGYLLLALIAAVGFGPGVHNLLLFAVG---------- 262
WP_014051708_1 DUF63 PTAPYLGSVWLFVVVKAALACGIALLLADSVREDPAEGRLLFVLVAAVGLGPGAHNLLLFTIS---------- 268
WP_049980148_1 DUF63 PTAELLGSAWLFVLVKLALGAGITALLAESVREAPRQGRLLLAFVAAVGLGPGAHNLLLFSIMG--------- 268
WP_053948907_1 DUF63 PTADLFGSAWLFVLVKLALGAGVTALLAESVREAPREGRLLLAFVAAVGLGPGAHNLLLFSVTG--------- 267
WP_079233625_1 DUF63 PTAELIGSAWLFVLVKLALGAGITALLAESVREAPREGRLLLAFVAAVGLGPGAHNLLLFSITA--------- 267
KPN32293_1 hypotheti PTAELLGTAWPFVLVKLALGAGITALLAESVREAPRQGRLLLAFVAAVGLGPGAHNLLLFSIMG--------- 268
```
