## Supplementary Information for "Beyond bacterial paradigms: uncovering the functional significance and first biogenesis machinery of archaeal lipoproteins"

**This PDF file includes:**

Supplementary Tables 1-3

Legend for Supplementary Data 1 to 7

**Other supporting materials for this manuscript include the following:**

Supplementary Data 1 to 7

**Supplementary Table 1. Plasmids used in this study.**

| Name | Relevant characteristic(s) | Source of reference |
|---|---|---|
| pTA131 | Amp$^r$, pBluescript II with BamHI-XbaI fragments from pGB70 harboring *pfdx-pyrE2* | 1 |
| pTA963 | Amp$^r$, *pyrE2* and *hdrB* markers, Tryptophan-inducible promoter (p.*tna*) | 2 |
| pYH1 | pTA131 carrying fragment with ~1500 nucleotides of fused upstream and downstream regions of *aliA* (*hvo_2859*) | This study |
| pYH2 | pTA131 carrying fragment with ~1500 nucleotides of fused upstream and downstream regions of *aliB* (*hvo_2611*) | This study |
| pYH3 | pTA963 carrying C-terminally myc-tagged *hvo_1176* | This study |
| pYH4 | pTA963 carrying C-terminally myc-tagged mutant *hvo_1176* (C21S) | This study |
| pYH5 | pTA963 carrying C-terminally myc-tagged *hvo_B0139* | This study |
| pYH6 | pTA963 carrying C-terminally myc-tagged mutant *hvo_B0139* (C22S) | This study |
| pYH7 | pTA963 carrying C-terminally myc-tagged *hvo_1705* | This study |
| pYH8 | pTA963 carrying C-terminally myc-tagged mutant *hvo_1705* (C31S) | This study |
| pYH9 | pTA963 carrying C-terminally myc-tagged *hvo_0844* | This study |
| pYH10 | pTA963 carrying C-terminally his-tagged *aliB* | This study |
| pYH11 | pTA963 carrying *aliA* | This study |
| pYH12 | pTA963 carrying both C-terminally myc-tagged *hvo_1176* and his-tagged *aliB* | This study |
| pYH13 | pTA963 carrying both C-terminally myc-tagged *hvo_1705* and his-tagged *aliB* | This study |

**Supplementary Table 2. Strains used in this study.**

| Name | Relevant characteristic(s) | Source of reference |
|---|---|---|
| H53 | Δ*pyrE2*/Δ*trpA* | 1 |
| YH_113 | H53 Δ*aliA* | This study |
| YH_57 | H53 Δ*aliB* | This study |
| YH_93 | H53 Δ*aliA*/Δ*aliB* | This study |
| YH_116 | H53 containing pYH3 | This study |
| YH_188 | H53 containing pYH4 | This study |
| YH_73 | H53 containing pYH5 | This study |
| YH_183 | H53 containing pYH6 | This study |
| YH_75 | H53 containing pYH7 | This study |
| YH_184 | H53 containing pYH8 | This study |
| YH_74 | H53 containing pYH9 | This study |
| YH_159 | Δ*aliA* containing pYH9 | This study |
| YH_82 | Δ*aliB* containing pYH9 | This study |
| YH_173 | Δ*aliA*/Δ*aliB* containing pYH9 | This study |
| YH_160 | Δ*aliA* containing pYH3 | This study |
| YH_132 | Δ*aliB* containing pYH3 | This study |
| YH_174 | Δ*aliA*/Δ*aliB* containing pYH3 | This study |
| YH_158 | Δ*aliA* containing pYH5 | This study |
| YH_81 | Δ*aliB* containing pYH5 | This study |
| YH_172 | Δ*aliA*/Δ*aliB* containing pYH5 | This study |
| YH_162 | Δ*aliA* containing pYH7 | This study |
| YH_83 | Δ*aliB* containing pYH7 | This study |
| YH_176 | Δ*aliA*/Δ*aliB* containing pYH7 | This study |
| YH_76 | H53 containing pTA963 | This study |
| YH_157 | Δ*aliA* containing pTA963 | This study |
| YH_230 | Δ*aliA* containing pYH11 | This study |
| YH_233 | Δ*aliA* with pTA963 carrying both C-terminally myc-tagged *hvo_1176* and *aliA* | This study |
| YH_231 | Δ*aliA* with pTA963 carrying both C-terminally myc-tagged *hvo_1705* and *aliA* | This study |
| YH_84 | Δ*aliB* containing pTA963 | This study |
| YH_136 | Δ*aliB* containing pYH10 | This study |
| YH_165 | Δ*aliB* containing pYH12 | This study |
| YH_167 | Δ*aliB* containing pYH13 | This study |

**Supplementary Table 3. Primers used in this study.**

| Name | Description | 5' to 3' sequence | Source of reference |
|---|---|---|---|
| hvo_2859_ups_fwd | forward primer used to amplify approximately 750 bp upstream of *aliA* | ATTATCTAGAGAGGCGGCCCGCGAA | This study |
| hvo_2859_ups_rvr | reverse primer used to amplify approximately 750 bp upstream of *aliA* | GAGACGGCGGGTCGATCGCTCTGCTACCGTTGC | This study |
| hvo_2859_down_fwd | forward primer used to amplify approximately 750 bp downstream of *aliA* | GCAACGGTAGCAGAGCGATCGACCCGCCGTCTC | This study |
| hvo_2859_down_rvr | reverse primer used to amplify approximately 750 bp downstream of *aliA* | AATACTCGAGCACCGCGACGGCGAG | This study |
| hvo_2859_inside_fwd | forward primer used to verify the absence of *aliA* in knockout strains | GGCTTCGTCTGGCACTACTT | This study |
| hvo_2859_inside_rvr | reverse primer used to verify the absence of *aliA* in knockout strains | TCATCCAATCGAGGCCGATG | This study |
| hvo_2611_ups_fwd | forward primer used to amplify approximately 750 bp upstream of *aliB* | ATTATCTAGAGACGACCGCGAGACCG | This study |
| hvo_2611_ups_rvr | reverse primer used to amplify approximately 750 bp upstream of *aliB* | AACAGGGTCGCGGCCGCAGGAGGCGCGGTG | This study |
| hvo_2611_down_fwd | forward primer used to amplify approximately 750 bp downstream of *aliB* | CACCGCGCCTCCTGCGGCCGCGACCCTGTT | This study |
| hvo_2611_down_rvr | reverse primer used to amplify approximately 750 bp downstream of *aliB* | AATACTCGAGCACGTCTGAGACGGGGTC | This study |
| hvo_2611_inside_fwd | forward primer used to verify the absence of *aliB* in knockout strains | ATGGCCATCCTCCCCG | This study |
| hvo_2611_inside_rvr | reverse primer used to verify the absence of *aliB* in knockout strains | TCAGCCGGCGATGGC | This study |
| hvo_1176Myc_oe_fwd | forward primer used to amplify *hvo_1176* with a C-terminal myc tag | TATATTCCATATGATGAACAGGCGACTCCTC | This study |
| hvo_1176Myc_oe_rvr | reverse primer used to amplify *hvo_1176* with a C-terminal myc tag | TATATTGAATTCTCACAGATCCTCCTCGCTGATGAGCTTCTGCTCGCGGTCGTCGTCTTCCA | This study |

| | | | |
|---|---|---|---|
| hvo_1176Myc_C21S_F | forward primer used to amplify *hvo_1176* with a C-terminal myc tag and a C21S mutation | GACCGCCGGGAGTCTCGGCGG | This study |
| hvo_1176Myc_C21S_R | reverse primer used to amplify *hvo_1176* with a C-terminal myc tag and a C21S mutation | ACGAGGAGGAGGGCGAGC | This study |
| hvo_B0139Myc_oe_fwd | forward primer used to amplify *hvo_B0139* with a C-terminal myc tag | TATATTCCATATGATGCTCTCCAGACGAGAG | This study |
| hvo_B0139Myc_oe_rvr | reverse primer used to amplify *hvo_B0139* with a C-terminal myc tag | TATATTGAATTCTCACAGATCCTCCTCGCTGATGAGCTTCTGCTCTGCGACCTTCACCAC | This study |
| hvo_B0139_C22S_F | forward primer used to amplify *hvo_B0139* with a C-terminal myc tag and a C22S mutation | GGTCGCTGGCAGCGGCGGTTC | This study |
| hvo_B0139_C22S_R | reverse primer used to amplify *hvo_B0139* with a C-terminal myc tag and a C22S mutation | CCGCTCACGAGCGCGGCT | This study |
| hvo_1705Myc_oe_fwd | forward primer used to amplify *hvo_1705* with a C-terminal myc tag | TATATTCCATATGATGCACGAACGCGACCGA | This study |
| hvo_1705Myc_oe_rvr | reverse primer used to amplify *hvo_1705* with a C-terminal myc tag | TATATTGAATTCTCACAGATCCTCCTCGCTGATGAGCTTCTGCTCGAGAACGCCCACCTC | This study |
| hvo_1705_C31S_F | forward primer used to amplify *hvo_1705* with a C-terminal myc tag and a C31S mutation | GCCGGGAGTACCGGCGGAAGCGAC | This study |
| hvo_1705_C31S_R | reverse primer used to amplify *hvo_1705* with a C-terminal myc tag and a C31S mutation | CCGGTACTCCCGGCGAGTGCGC | This study |
| hvo_0844Myc_oe_fwd | forward primer used to amplify *hvo_0844* with a C-terminal myc tag | TATATTCCATATGATGAAAAGGCGGGAGTTT | This study |
| hvo_0844Myc_oe_rvr | reverse primer used to amplify *hvo_0844* with a C-terminal myc tag | TATATTGAATTCTTACAGATCCTCCTCGCTGATGAGCTTCTGCTCTTCGCCACCTCCGGT | This study |
| hvo_2859_oe_F | forward primer used to amplify *aliA* | TATATTCCATATGATGGCAACGGTAGCAGAG | This study |
| hvo_2859_oe_R | reverse primer used to amplify *aliA* | TATATTGAATTCTTAGACGCCGAACGTCGCG | This study |
| hvo_2611His_oe_fwd | forward primer used to amplify *aliB* with a C-terminal his tag | TATATTCCATATGATGGCCATCCTCCCCG | This study |

| | | | |
|---|---|---|---|
| hvo_2611His_oe_rvr | reverse primer used to amplify *aliB* with a C-terminal his tag | TATATTGAATTCTCAGTGATGGTGATGGTGATGCGGGCCGCCGCCGGCGATGGCGAA | This study |
| pTA963_dual_F | forward primer used to verify the insertion of lipoprotein encoding genes in the pTA963 plasmid | CCCCGAAAAGTGCCACCTAAA | This study |
| pTA963_dual_R | reverse primer used to verify the insertion of lipoprotein encoding genes in the pTA963 plasmid | CGTCACGTACGAGCGCAAA | This study |
| ga_dual_F | forward primer used to amplify lipoprotein encoding genes from pTA963 overexpression plasmid for the construction of a dual expression vector | CCGAAATCGGCAAAATCCCTTACTGGCGAAAGGGGGATGT | This study |
| ga_dual_R | reverse primer used to amplify lipoprotein encoding genes from pTA963 overexpression plasmid for the construction of a dual expression vector | GCGGAGGAATCCGATGAGTACTGGCACGACAGGTTTCC | This study |
| pTA963_seq_F | forward primer used to verify the insertion of genes in the multi cloning site region | ATGCACACACCAGTCCACGAGC | This study |
| pTA963_seq_R | reverse primer used to verify the insertion of genes in the multi cloning site region | GAACAAAAGCTGGAGCTCCACCG | This study |

**Supplementary Data 1 (separate file). Number of predicted lipoproteins in archaeal genomes from the arCOG database**

**Supplementary Data 2 (separate file). 500 arCOGs with the best lipoprotein co-occurrence scores**

**Supplementary Data 3 (separate file). Number of proteins assigned to arCOG2177 or arCOG02178 in each genome**

**Supplementary Data 4 (separate file). Multiple alignment of arCOG02177 proteins**

**Supplementary Data 5 (separate file). Multiple alignment of arCOG02178 proteins**

**Supplementary Data 6 (separate file). Gene neighborhoods of arCOG02177 and arCOG02178**

**Supplementary Data 7 (separate file). *Hfx. volcanii* lipoproteins predicted with ≥90% confidence using SignalP 6.0**
